# Coheritability and Coenvironmentability as Concepts for Partitioning the Phenotypic Correlation

**DOI:** 10.1101/598623

**Authors:** Jorge Vasquez-Kool

## Abstract

Central to the study of joint inheritance of quantitative traits is the determination of the degree of association between two phenotypic characters, and to quantify the relative contribution of shared genetic and environmental components influencing such relationship. One way to approach this problem builds on classical quantitative genetics theory, where the phenotypic correlation 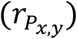 between two traits is modelled as the sum of a genetic component called the coheritability (*h*_*x,y*_), which reflects the degree of shared genetics influencing the phenotypic correlation, and an environmental component, namely the coenvironmentability (*e*_*x,y*_) that accounts for all other factors that exert influence on the observed trait-trait association. Here a mathematical and statistical framework is presented on the partition of the phenotypic correlation into these components. I describe visualization tools to analyze 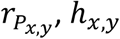 and *e*_*x,y*_ concurrently, in the form of a three-dimensional (3DHER-plane) and a two-dimensional (2DHER-field) plots. A large data set of genetic parameter estimates (heritabilities, genetic and phenotypic correlations) was compiled from an extensive literature review, from which coheritability and coenvironmentability were derived, with the object to observe patterns of distribution, and tendency. Illustrative examples from a diverse set of published studies show the value of applying this partition to generate hypotheses proposing the differential contribution of shared genetics and shared environment to an observed phenotypic relationship between traits.

## Introduction

A fundamental aspect in the study of heredity is to investigate associations among traits at the phenotypic level, and to determine the degree shared genetics and common environmental influences shape such associations. Many traits are analyzed jointly in genetic studies in the hope of providing greater statistical power to detect associations to causal genetic factors (Melton et al. 2010, Cheng et al. 2013, Jia and Jannick 2012). Understanding how ceratin traits relate to disease risk is of primary concern in clinical medicine (Wellman et al. 2013, Oren et al. 2015, Barabási et al. 2010) and in animal and plant breeding (Sölkner et al. 2008). Most phenotypes are the result of a complex interaction of multiple genetic and environmental factors (Lander and Schork 1994), and the ability to assay these characters presents a unique opportunity to explore mechanisms underlying concerted inheritance.

The phenotypic correlation 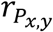 between two traits, say *x* and *y*, has been extensively used to quantify the relationship between observable characters. Its partition between a genetic component and an environmental component was originally worked out by Hazel (1943) as a function of heritabilities (*h*^2^) of the traits and correlations (genetic 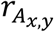, environmental 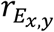) between them (see equation [5] below). The term coheritability, coined by Nei (1960) as a descriptor of the genetic component, is a measure of joint inheritance of two traits that representes the genetic contribution to their phenotypic correlation. This contrasts with coenvironmentability, *e*_*x,y*_ that quantifies the contribution of all other sources of variation, excluding additive genetic factors, influencing the phenotypic correlation. However, the coheritability as a concept and measure has remained largely unrecognized. In the 1996 edition of *Principles of Quantitative Genetics* the term coheritability was reintroduced more formally (Falconer and MacKay 1996, p. 317) in the context of correlated response to selection. Yet a proper treatment of its mathematical and statistical properties is still lacking. Despite its simple formulation, there is a persistent confusion in the scientific literature of what coheritability means or how should it be calculated. Some have denoted coheritability as the mere ratio of additive genetic to phenotypic covariances (de Reggi 1972, Janssens 1979) without sufficient theoretical support, while others employ the term coheritability (or co-heritability) to refer to different *ad hoc* measures of coinheritance (Sae-Lim 2015, Hoskens et al. 2018), or colloquially coinheritance applies to any observed joint transmission of traits (Wambua et al. 2006, Höblinger et al. 2009). Uncertainty is compounded when coheritability is conflated with genetic correlation (Yang et al. 2016, Traglia et al. 2017, Yin et al. 2017), or is referred by various names, such as ‘coefficient of genetic prediction’ (Baradat 1976), ‘correlative heritability’ (Chen et al. 2003), ‘genetic contribution to the phenotypic correlation’ (Posthuma et al. 2003), ‘standardized genetic covariance’ (Rao and Rice 2005), ‘bivariate heritability’ (DeStephano et at. 2009), ‘Endophenotype Ranking Value’ (Glahn et al. 2012), and ‘proportion of phenotypic correlation due to genes’ (Wu et al. 2010, 2013, Muñoz el al. 2018).

This study substantially builds up on previous work (Hazel 1943, Lerner 1950, Searle 1961, Yamada 1968, Bedard et al. 1971, Plomin and DeVries 1979), and connects to modern investigations (Gui et al. 2017, Pick et al. 2016, Gianola et al 2015) in the attempt to clarify the nature of coheritability and coenvironmentability as constituent of the phenotypic correlation. I explore this topic through theoretical arguments and through the analyses of data compiled from a diverse and large number of studies. The objectives of this paper were: (1) to present a theoretical background on the mathematical and statistical properties of phenotypic correlation, coheritability and coenvironmentability. (2) Analyze their distribution, dispersion and tendency. (3) Model the relationship between the phenotypic correlation and (*h*_*x,y*_, *e*_*xy*_) and 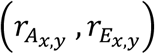. (4) Present illustrative examples on the application of the 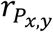 decomposition in order to gain insight in different biological problems. Further statistical and mathematical details as well as illustrative examples are provided in the Supplementary Information (SI) document accompanying this paper.

## Theoretical Background

### The components of the sample phenotypic correlation

In the framework of quantitative genetics, an individual’s phenotypic value *P* is modelled as the sum of an additive genetic value *A* and an environmental value *E*, thus *P* = *A* + *E*. This simple decomposition also applies to the phenotypic covariance 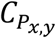, which as a measure of linear association between the phenotypic values of two characters *x* and *y*, results from the sum of an additive genetic covariance 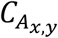, and a term including all residual genetic and non-genetic factors, namely the environmental covariance 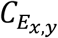, thus

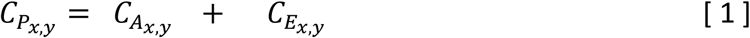

To standardize, both sides of equation [1] are divided by the geometric mean of the phenotypic variances of each trait (i.e., which could be construed as the joint *bivariate phenotypic variability* of the traits *x* and *y*),

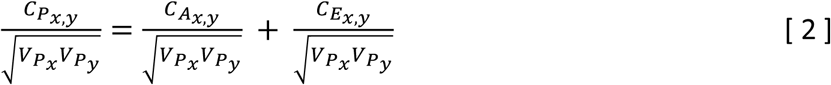

This expression can be summarized as follows:

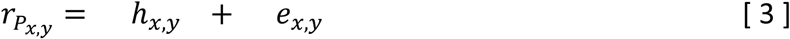

The term 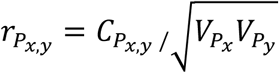 is the phenotypic correlation between the phenotypic values of characters *x* and *y*; it measures the linear association between the two observable characters.

The coheritability, defined by Nei (1960), as

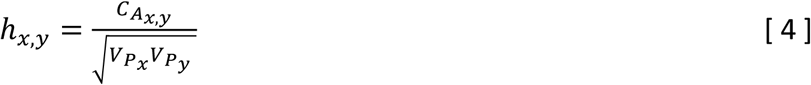

is the ratio of the genetic covariance on the bivariate phenotypic variability, is the component of the phenotypic correlation attributed to shared genetic effects, and thus reflects the extent that joint genetic influences have on the observed association of the characters.

The coenvironmentability (broad-sense), 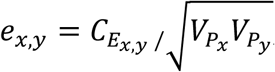, is the component of the phenotypic correlation defined as the ratio of the residual covariance (i.e. non-additive genetic, environmental, GxE interactions) to the bivariate phenotypic variability. It represents the joint influence of all factors that are not accounted by additive genetic factors that exert influence on the observed relationship between the traits.

Since covariances can be expressed in terms of their respective variances *V* and correlations *r*, such that 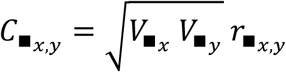 (where ▪ is either of the subscripts P, A or E). Equation [2] can be re-written as:

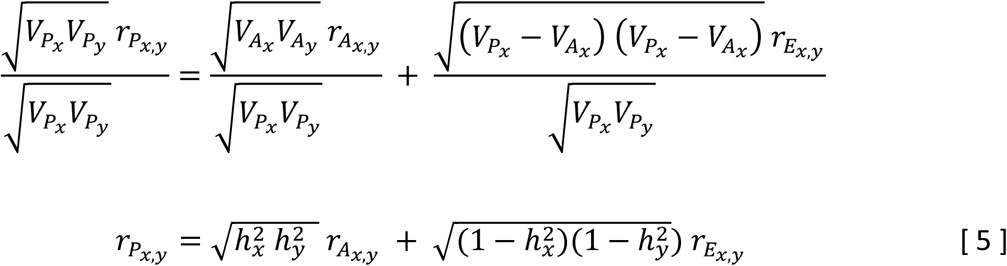

where 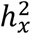 and 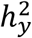 denote the narrow-sense heritabilities of the traits *x* and *y*. 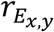 is the environmental correlation, and 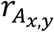 is the genetic correlation, defined as

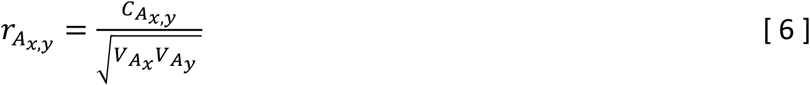

Equation [5], originally introduced by Hazel (1943, p. 480) in the context of selection indexes, models the phenotypic correlation as the sum of a weighed genetic correlation, namely the coheritability, which can be expressed as

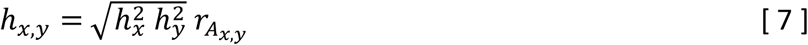

and a weighed environmental correlation, i.e., the coenvironmentability,

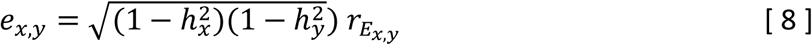

The expressions of coheritability given in Equations [4] and [7] are equivalent. Further equivalent can be done formulations using algebraic summations or by applying a path analytic method (or structural equation model) (Supplementary Information section 3.2). There are two more terms associated to the decomposition of the genetic covariance (Equation 1) and which involve terms of covariation of the breeding values a trait with the environmental values of the other (e.g. 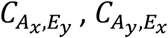). Some would opt to ignore these terms or consider them negligible, which is not different than including them in the environmental covariance term (Supplementary Information section 2.4-2.6). Their importance and influence deserve further study. From the relationships presented, it is clear that the phenotypic correlation’s magnitude and sign is directly influenced by the coheritability and coenvironmentability.

### The domains of coheritability, coenvironmentability and phenotypic correlation

The description of the phenotypic correlation as the sum of the coheritability and coenvironmentability imposes an intrinsic linear relationship between the three variables. Since the sum of *h*_*x,y*_ and *e*_*x,y*_ must yield 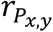, and the Cauchy-Schwarz inequality proves that the phenotypic correlation is bound to the domain [-1, +1]. This implies that if either of the variables *h*_*x,y*_ or *e*_*x,y*_ becomes zero, then the other variable would become equal to the phenotypic correlation. Therefore, *h*_*x,y*_ and *e*_*x,y*_ each must also have a domain within [-1, +1] with the added condition that the sum of their absolute values cannot exceed 1 (|*h*_*x,y*_| + |*e*_*x,y*_| ≤ 1). For instance, if one of the heritabilities is unity, or the 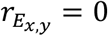, then *e*_*x,y*_ = 0, then equation [4] becomes 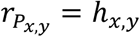. Similarly, if one the heritabilities or the genetic correlation is zero, then *h*_*x,y*_ = 0, and it results in 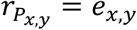. All this implies that both coheritability and coenvironmentability can be subject to the same inferential statistical methods as those designed for the assessment of correlations (Supplementary Information section 5.1).

The environmental correlation 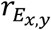, whose factors may remain unspecified, could be calculated as a residual derived from [4]:

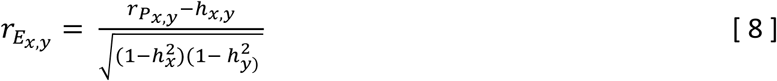

and then use it to determine the (broad-sense) coenvironmentability using equation [6]. Note that equation [7] will introduce dependency and collinearity if applied to a regression model involving the phenotypic correlation. A superior method would estimate the environmental correlation independent of the phenotypic correlation and coheritability, and relate to specified environmental factors. In this case, the coheritability and the narrow-sense coenvironmentability would not necessarily add up to 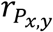 and should be an indication of that significant genotype x environment interaction terms are present.

### Visualization of 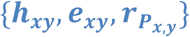

The three-dimensional coheritability-coenvironmentability-phenotypic correlation plane, 3DHER) plane is a Cartesian three-dimensional space employed to represent these three variables (Figure 1), where a single datum is defined by its coordinates 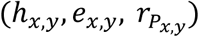. Spherical coordinates can also be used to describe the behavior of *h*_*x,y*_, *e*_*x,y*_, and 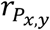 (Supplementary Information section 6.3) and serve as a complementary way to provide further insight. A first impression from observing this graph is that the data lies on a virtual slanted plane with zero volume. Mathematical theory tells that this is due to the existence of an intrinsic linear relationship among the three variables. Further corroboration that the possible values fall on a plane with zero volume is given by the fact that the determinant of a matrix involving any three distinct data points is zero (Supplementary Information section 6.4).

**Figure 1.**
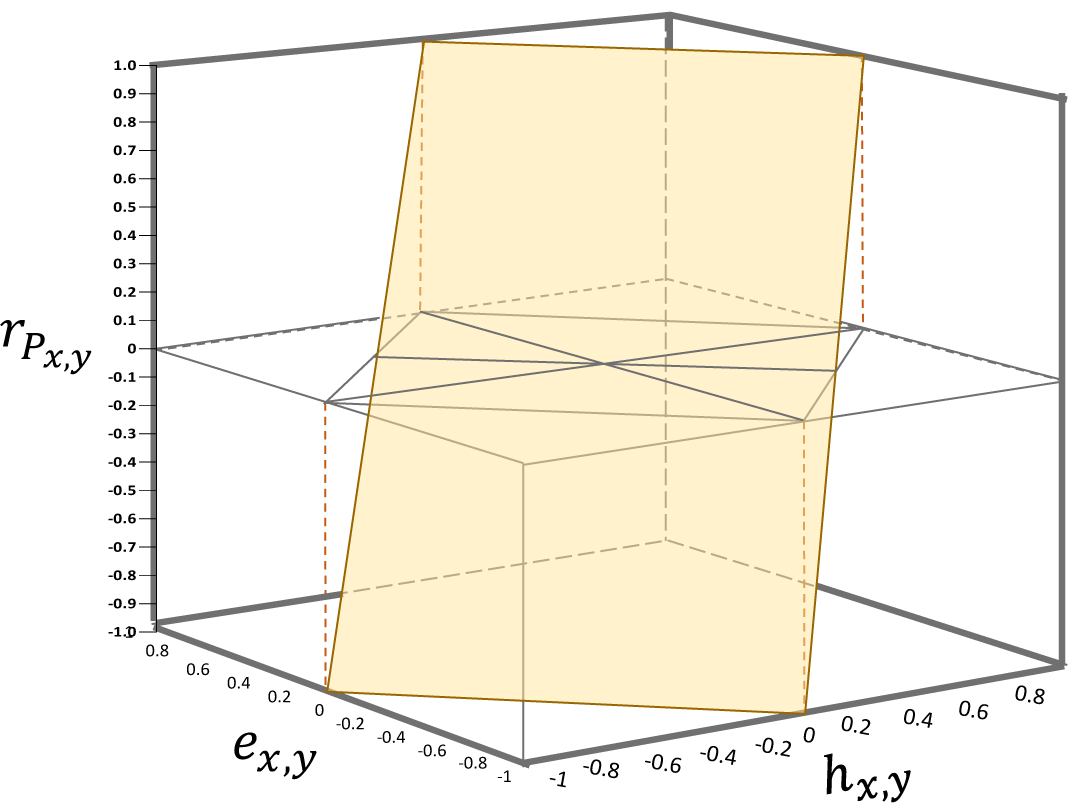
The three-dimensional 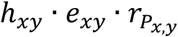-plane (3DHER-plane) presented in (A) the Cartesian coordinate system. The plane represents the area where the data of coheritability, coenvironmentability and phenotypic correlation satisfy the equation 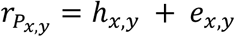, where 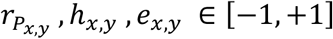.

The two-dimensional coheritability-coenvironmentability-phenotypic correlation (2DHER) field is the orthogonal projection of the 3Dplane onto the *h*_*x,y*_ · *e*_*x,y*_-surface when 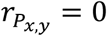 (see Figures 1), this graph retains information of the three variables (Figure 2, Supplementary Information section 6.4). This field represents an area bound by the relationship |*h*_*x,y*_| + |*e*_*x,y*_| = 1 (i.e. its borders are demarcated by the lines *h*_*x,y*_ + *e*_*x,y*_ = 1, *h*_*x,y*_ + *e*_*x,y*_ = −1, *h*_*x,y*_ - *e*_*x,y*_ = 1, and *h*_*x,y*_ - *e*_*x,y*_ = −1). The domain of the phenotypic correlation can be superimposed on it, knowing that the elements of a data point (*h*_*x,y*_, *e*_*x,y*_) must add to 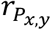. The field is a continuum domain of *h*_*x,y*_, *e*_*x,y*_ and 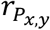, that accounts for all sign combinations among these three variables (excluding incongruous combinations consisting of *h*_*x,y*_ and *e*_*x,y*_ having the same sign, yet summing up to an 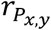 with a different sign). For the sake of clarity, the field can be divided by tracing the lines 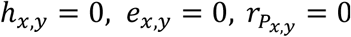. The six triangular partitions thus produced are labeled with the letter *S* followed by a subscript having a signed numeral, which is the sign of the coheritability (which in turn is conferred by the genetic correlation), and the numeral serves as a dummy indicator (Figure 1). The domain for variables *h*_*x,y*_, *e*_*x,y*_ and 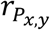 is specific for each partition (Figure 2B). Partitions that share a reciprocal angle become reciprocal partitions whose labels possess the same numeral by have different signs (i.e. there are three pairs of reciprocal partitions, namely *S*_+1_ and *S*_-1_; *S*_+2_ and *S*_-2_; *S*_+3_ and *S*_-3_). Notice that positive phenotypic correlations are found in partitions *S*_-3_, *S*_+1_, *S*_+2_, and the negative phenotypic correlations in *S*_+3_, *S*_-1_, *S*_-2_. Positive coheritabilities (and implicitly genetic correlations) are found in partitions *S*_+1_, *S*_+2_, *S*_+3_; and negative coheritabilities in partitions *S*_-1_, *S*_-2_, *S*_-3_. Lastly, partition, *S*_0_, contain all data with at least one of the three variables 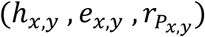 equal to zero and therefore lie on one of the diving lines. The *S*_+1_ and *S*_-1_ are reciprocal partitions where all the three variables 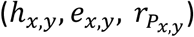 have the same sign, either positive (*S*_+1_) or negative (*S*_-1_), and each occupy an area equal to 0.5. Both are demarcated by the lines *h*_*x,y*_ = 0 and 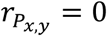, and line *h*_*x,y*_ + *e*_*x,y*_ = 1 in the case of *S*_+1_, and line *h*_*x,y*_ + *e*_*x,y*_ = −1 for *S*_-1_. The area is equally divided by the line *h*_*x,y*_ = *e*_*x,y*_ separating data that satisfy either *h*_*x,y*_ < *e*_*x,y*_ or *h*_*x,y*_ > *e*_*x,y*_.

**Figure 2.**
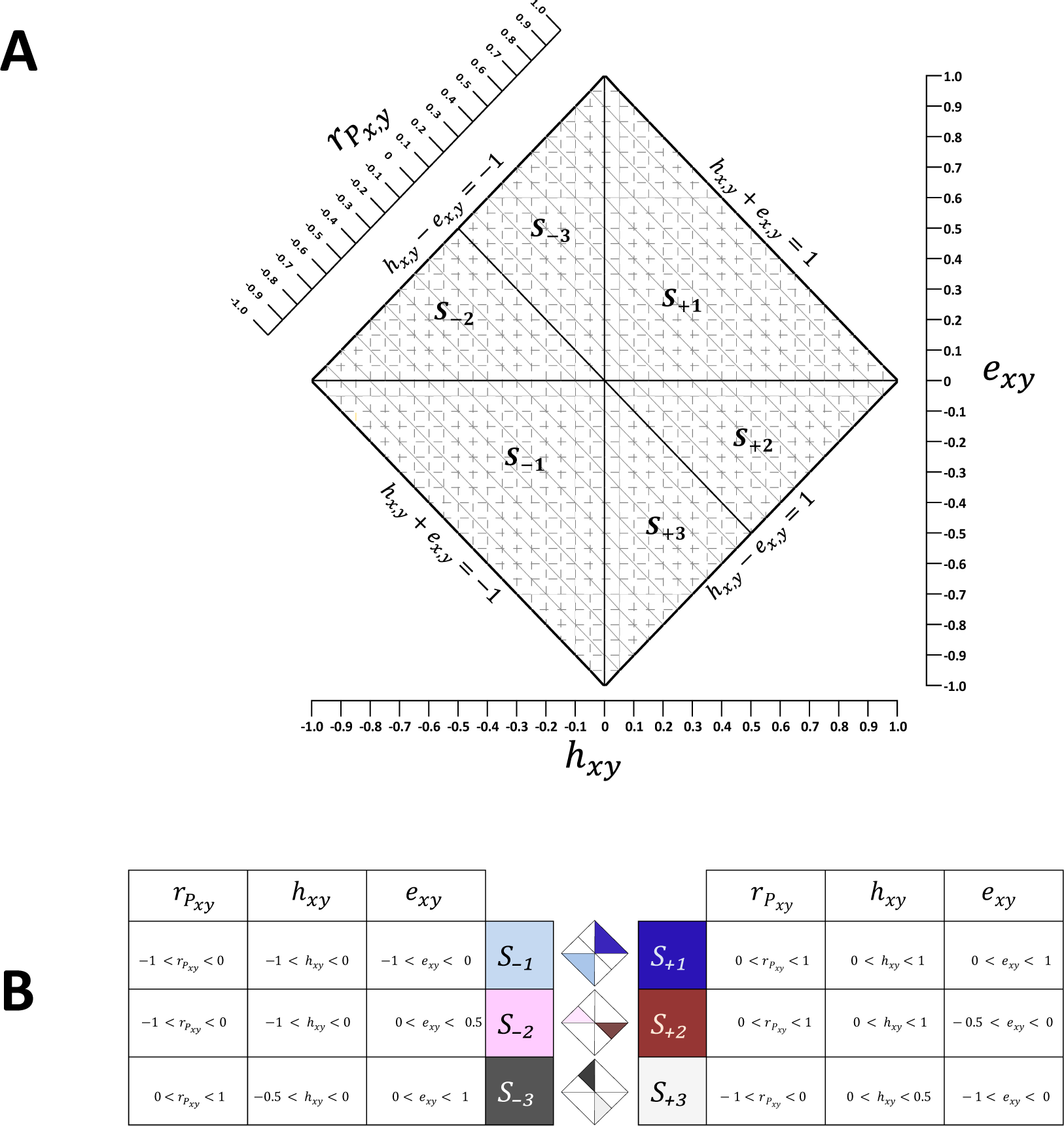
The two-dimensional 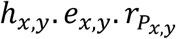-field (2DHER-field). This is the result of the projection of the 3D plane onto the *h*_*x,y*_ · *e*_*x,y*_ surface. (A) Partitions are demarcated by tracing the lines 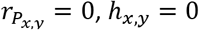, and *e*_*x,y*_ = 0. Partitions are labeled with a *S* symbol whose subscript positive (+) or negative (–) correspond to the sign of the genetic correlation followed by a numeral. Partitions denoted with the same numeral but with different sign are reciprocal. (B) The domain interval of *h*_*x,y*_, *e*_*x,y*_, and 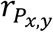 by partition. The equation |*h*_*x,y*_| + |*e*_*x,y*_| = 1 defines the boundary lines of the field, which are explicitly written at its sides. The total area of the field is 2. Partitions S_+1_ and S_-1_ occupy each an area of 0.5, while partitions *S*_+2_, *S*_-2_, *S*_+3_, and *S*_-3_ each cover an area equal to 0.25. The line *h*_*x,y*_ = *e*_*x,y*_ distinguishes all instances where *h*_*x,y*_ < *e*_*x,y*_ and *h*_*x,y*_ > *e*_*x,y*_.

The reciprocal partitions *S*_+2_ and *S*_-2_ are characterized by 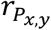 and *h*_*x,y*_ having the same sign, positive in the case of *S*_+2_, and negative in the case of *S*_-2_. In these partitions, the magnitude of coheritability is larger than coenvironmentability, and therefore shows the preponderant influence of a common genetic component upon the phenotypic correlation. These partitions occupy an area of 0.25 each and are demarcated by the lines 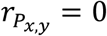 and *e*_*x,y*_ = 0, and either line *h*_*x,y*_ - *e*_*x,y*_ = 1 (for *S*_+2_) or *h*_*x,y*_ - *e*_*x,y*_ = −1 (for *S*_-2_). Note that the coenvironmentability is bound by the interval [0, +0.5] in *S*_-2_, and [−0.5, 0] for *S*_+2_. The reciprocal partitions *S*_+3_ and *S*_-3_ are characterized by 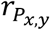 and *h*_*x,y*_ having different signs, due to the fact that the phenotypic and genetic correlation differ in direction. Each of these partitions cover an area of 0.25, and are delimited by 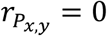 and *h*_*x,y*_ = 0, and line *h*_*x,y*_ - *e*_*x,y*_ = 1 (for *S*_+3_) or line *h*_*x,y*_ - *e*_*x,y*_ = −1 (for *S*_-3_). In these partitions, the magnitudes of *h*_*x,y*_ is smaller than the one for coenvironmentability, indicating that there is an overriding environmental effect on the phenotypic correlation. The line diving equally partitions *S*_+1_ and *S*_-1_ separates the instances where *h*_*x,y*_ < *e*_*x,y*_ (at the side of Partitions *S*_-2_ and *S*_-3_), and where *h*_*x,y*_ > *e*_*x,y*_ (at the side of *S*_+2_, *S*_+3_).

### Relationship between genetic correlation and heritabilities

The bivariate correlations and univariate heritabilities are independent, random variables, variation in one does not lead to a concomitant effect on the other. The weights expressed as functions of the heritabilities, however, are inversely related: as the geometric mean of the heritabilities 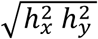 increases, there is a concomitant decrease of 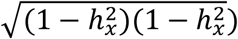. However, larger heritabilities do not necessarily imply larger influence of the genetic component. For instance, if the heritabilities of the traits are 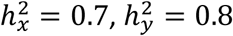, and the correlations 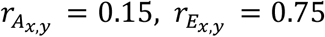, then the resulting coheritability *h*_*x,y*_ = 0.111, despite the large heritabilities of the traits, is smaller than the coenvironmentability *e*_*x,y*_ = 0.1836. On the other hand, a larger genetic correlation does not necessarily translate into a larger genetic influence either. For example, let us say that 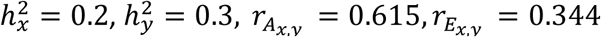 The coheritability becomes *h*_*x,y*_ = 0.15, which is smaller than the coenvironmentability *e*_*x,y*_ = 0.25, despite that the latter has a low environmental correlation. This fact may bring into reconsideration methods that attempt to map the degree of genetic influence on the phenotypic correlation based on the mere comparison of 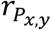 and 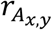. Rank between these statistics is not preserved when using the totality of the data.

If, on the other hand, one of the variable is set to a specified value, it would effectively limit the range of possible values on the other variables. Questions such as, given the value of a phenotypic correlation, what would be the possible values of the coheritability and coenvironmentability?, or, condition on specified values of coheritability and coenvironmentability, what are the possible values of the heritabilities of the traits and the genetic and the environmental correlations? These topics are duly elaborated in the supplementary information (SI section 7.1 and 7.2) in an empirical manner that yield possible, not probable, values. This is primarily meant to lay the groundwork for more rigorous development.

### The relationship of the magnitudes of the coheritability and of the genetic correlation

If the heritabilities of the traits are each unity, then the coenvironmentability becomes zero, the coheritability equates the genetic correlation, and only under this extreme condition,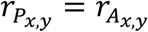. In the majority of the cases, however, the absolute value of the coheritability would always be less than the absolute value of the genetic correlation. The geometric mean of the heritabilities, 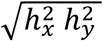, itself a decimal number between 0 and 1, multiplied to the genetic correlation results in the product (i.e. coheritability) with a smaller magnitude, such that the coheritability would move away from the 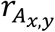 towards the origin. For instance, if 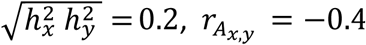 then *h*_*x,y*_ = –0.08. The latter is closer to zero than 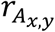. If 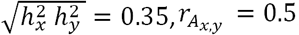, then *h*_*x,y*_ = 0.175, which also moves towards the zero direction Therefore, in terms of magnitude, the coheritability would be equal of less than the magnitude of the genetic correlation,

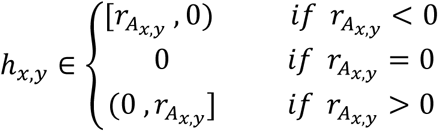

### Inference

Details on the characteristics of the base population and the sample, as well as aspects concerning hypothesis testing and confidence intervals determination for coheritability, coenvironmentability and phenotypic correlation are duly elaborated in the Supplementary Information sections 1 and 5. In brief, consider a large, genetically-structured population whose individuals possess heritable traits 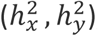, and with genetic 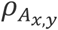, environmental 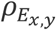, and phenotypic correlation 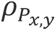 between the traits, the latter being the sum of the population-level coheritability *H*_*x,y*_ and coenvironmentability *E*_*x,y*_ parameters 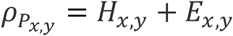. It is, therefore, feasible to make propositions about these population parameters using data obtained from sampling. If a sample of size *n* is obtained from such population, one could hypothesize the values of the parameters 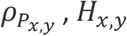, and *E*_*x,y*_, provided that they satisfy the relationship 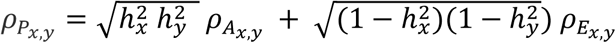. The distribution of the estimator of 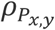 is the sample phenotypic correlation 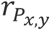 is construe to be the distribution of the correlation coefficients determined from *m* samples, all of size *n*, drawn from the original population (see Supplementary Information section 4 for derivation of sampling distribution of the correlation coefficient, Fisher 1915, Hotelling 1959). Naturally, sample size is an important consideration for estimation and hypothesis testing. Under a condition of moderate (70 < *n* ≤ 100) to large sample sizes (*n* > 100), asymptotic properties of the statistics can be assumed (Supplementary Information section 6). Other aspects of statistical inference on 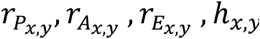, and *e*_*x,y*_ such as hypothesis testing, power, and confidence level of parameters, as well as aspects of experimental design regarding determination of an adequate sample size are presented in Supplementary Information section 5.

### Simulation of bivariate (***h***_***x***,***y***_, ***e***_***x***,***y***_) data

Simulation is fundamental to further an understanding on probabilities, estimation of the sampling distribution of statistics, calculation of coverage probability of confidence intervals, and to evaluate the robustness of statistical tests (Wicklin 2013). To facilitate these paths of inquiry, a method to generate bivariate (*h*_*x,y*_, *e*_*x,y*_) observations is presented (Supplementary Information section 10) based on a transformation of two random uniform random variables.

## Materials and Methods

### Data compilation and validation

An extensive search of data published in journal articles was carried out using bibliographic sources such as PubMed, Web of Science, GoogleScholar. The search was carried out using keywords: ‘coheritability’, ‘genetic parameters’, ‘phenotypic and genetic correlations’, ‘correlated response’, ‘age-age correlations’, and ‘early selection’, ‘fitness trade-offs’. To be included in the compilation (see Flow chart in Supplementary Material, File D1), the reported data had to involve continuous traits and minimally include trait heritabilities and at least two correlations (phenotypic, genetic, or environmental). Information was also gathered about the organism studied, standard errors of the parameter estimates (if any), and bibliographic citation. Coheritabilities and coenvironmentabilities were then calculated using equations [4] to [7]. The data was categorized into partitions according to the criteria expounded in Theoretical Background and summarized in Figure 2.

To ensure validation, three criteria were used for outlier detection, failure to satisfy at least one of them led to its exclusion from the data set: The first criterion checked whether the values of heritabilities and correlations were within their domains (*h*^2^ ∈ [0,1], *r* ∈ [–1,1]). A second criterion ascertained whether a given datum was within the boundaries of the 2DHER-field by holding the relationship |*h*_*x,y*_| + |*e*_*x,y*_| ≤ 1. Finally, the data had to satisfy the relationship 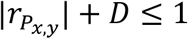, (*D*, disparity index, see below) to indicate that the values of 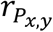 and 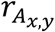 were coherent to the relationship expressed in equation [4]. Analyses of descriptive and inferential statistical analyses were carried out using SAS/STAT Software (SAS Institute, Cary, NC).

### Analysis of count data

It was of particular interest to see if the occupancy of the data in partitions (*S*_*0*_, *S*_+1_, *S*_+2_, *S*_+3_, *S*_–1_, *S*_–2_, *S*_–3_) came from independent trials. If each of n independent trials result in placing the data to one of the k partitions and the probability that a (collected) datum belongs to a given partition, is the same in every trial, then the count of the data in the partitions would follow a multinomial probability distribution. A goodness-of-fit test for multinomial distribution was therefore conducted to evaluate a null hypothesis proposing that the observed count of data points in each partition was the result of expected proportions assigned to each of the partitions.

### Empirical Distribution

Cognizant that the data came from disparate sources (organisms, populations, traits, experimental goals), it was not the intention to conduct a formal metaanalysis, but to simply visualize the general occupancy, tendency and dispersion of the phenotypic correlation, coheritability and coenvironmentability as scatter plot on the 3DHER-plane and 2DHER-field. Histograms were drawn to visualize the frequency and variability of the data.

### Modeling the phenotypic correlation

Though it was not the aim of this work to create a predictive statistical model, it was nevertheless of interest to observe how 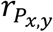, as a scalar, dependent variable, related to either (*h*_*x,y*_, *e*_*x,y*_) or 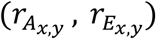 acting as regressors (or explanatory variables) in a multiple regression model. This exercise allowed also to check the relationship between regression parameters, and the consistency of regression equation using data as a whole and by partition. The models were:

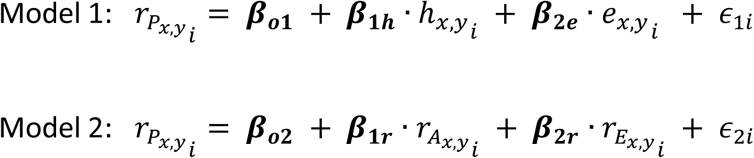

where the parameters ***β***_***0***_ is the intercept, ***β***_**1**_ is the slope of the regression of phenotypic correlation 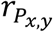 on the genetic factor, namely, *h*_*x,y*_ (model 1) or 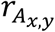 (model 2), and ***β***_**2**_ is the regression coefficient corresponding to the environmental factor, *e*_*x,y*_ (model 1), or 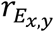 (model 2). Model 1 assumes that given the data set 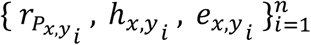 of *n* observations, a linear relationship exists between the dependent variable 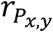 and the bivariate vector of regressors {*h*_*x,y*_, *e*_*x,y*_}. The ϵ term was assumed to have a normal distribution with zero mean and constant variance 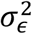 (a path analytical representation of model 1 is presented in Supplementary Information section 3.2.2). A similar rationale applies to model 2. The results of the regression analyses using Model 1 were assessed based upon the expectations derived from equation [3] (i.e. zero intercept, and each has a slope parameter equal to positive one). Model 2 was also expected to yield a zero intercept, and, if the genetic and environmental correlations are zero then the phenotypic correlations must also be zero. Otherwise, there are no theoretical relationship that relates 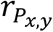 to 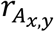 and 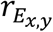 directly. The least-squares estimates of ***β***_**0**_, ***β***_**1**_ and ***β***_**2**_ were estimated from the data using the PROC REG procedure of SAS, applied to the data set as a whole as well as by partition. The statistics collected from the results were the intercept (*β*_*0*_), slope (*β*_1_), and the R-square of each model.

### Disparity Index Analyses

The disparity index (*D*) is defined as the absolute value of the difference between the phenotypic correlation and the genetic correlation, 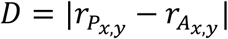 (Willis et al. 1991). This study used *D* for two purposes. One was as a means of validation (Supplementary Information Appendix 2). The second use was to evaluate the closeness of the magnitudes of 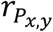 and 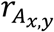. With the aim to further investigate the relationship between 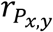 and 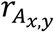, the disparity index was treated as a random variable, and for this purpose this work derived its probability density function, which used as a basis the transformation of the absolute difference of two uniform random variates, is formulated as *f*_*D*_(*D*) = 2 – 2*d* for 0 ≤ *D* ≤ 1 (Supplementary Information 7.5.2). A method to generate simulated disparity data is presented in Supplementary Information section 7.5.3.

### Illustrative Examples

Data from selected studies pertaining a range of topics of particular relevance to modern biology were used to help illustrate how the decomposition of 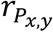 into *h*_*x,y*_ and *e*_*x,y*_ becomes an appropriate and pertinent approach to postulate hypotheses concerning the degree of influence that shared genetics and common environmental factors have on shaping an observable association between traits.

### Data availability

Supplementary Material is deposited and available in FigShare https://. It contains File D1 Data consisting of the raw numerical values obtained from the literature review and examples, File SI Supplementary Information including text explaining mathematical and inferential statistical aspects, and additional text and figures; and the File C1 Computing containing code and resources used in statistical and mathematical calculations (SAS, MS Excel).

## Results

This work analyzed more than 7700 observations comprising compiled data for the distribution, and examples, plus more than 16000 data points of the transcriptomics example. The data set compiled from journal articles amounted to *n* = 6287 observations, and involved 140 studies in the areas of human genetics, agronomy, forestry, fisheries, animal husbandry, ecology, and life history research. Data was collected from humans (*n* = 1069), animals (*n* = 3535, 39 genera, 40 species), plants (*n* = 1683, 26 genera, 33 species). Observations belonged to morphological (4288), physiological (1721), fitness-life history (20), and behavioral (258) traits. Around 60% of the studies were carried out under field conditions, and the rest in lab. The data collected consisted of heritabilities and correlations, and from them the coheritability and coenvironmentability corresponding to each observation, was derived including determination of the partition the datum belong to (see Theoretical Background section). The data set underwent an stringent validation procedure including detection and exclusion of outliers (Supplementary Information Appendix 2). Basic statistics for each variable (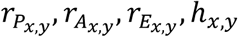 and *e*_*x,y*_) are presented in Supplementary Information Appendix 3A.

### Distribution of the *h*_*x,y*_, *e*_*x,y*_, *r*_*Px,y*_ data

Given the heterogeneity of the data in terms of objects of study, traits, sources, methods, precision, it was not the intention to perform a formal metanalysis, nor to combine data of the studies to create weighed averages. Rather, the objective here was to observed occurrence, tendency and variability of the data. Inspection of the coheritability and coenvironmentability data plotted on the 3DHER-plane (Figure 3A) or 2DHER-field (Figure 4A) shows it scattered on a two-dimensional object. The 3D plot of the phenotypic correlation as a function of the genetic and environmental correlations occupied a more variable volume (Figure 3B, 4B). The mean values of coheritability (0.075, SE 0.0039) and coenvironmentability (0.083, SE 0.0022) matched very well with the calculated overall mean phenotypic correlation (0.158, SE 0.0025). The variability, as measured by the standard deviation, was less for the coheritability (0.177) than for the coenvironmentability (0.198) and phenotypic correlation (0.309). The standard error of the estimates was the lowest for coheritability (mostly found below 0.16) (for formulas see Supplementary Information section 3.3). The dispersion was not uniform, all the variables presented abundance around the origin, but became more infrequent towards the borders (Figure 4A) particularly at the neighborhood of the lines *h*_*x,y*_ – *e*_*x,y*_ = 1 and *h*_*x,y*_ – *e*_*x,y*_ = –1. Overall, the tendency of the relationship of 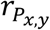 with 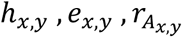 and 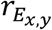 was positive.

**Figure 3.**
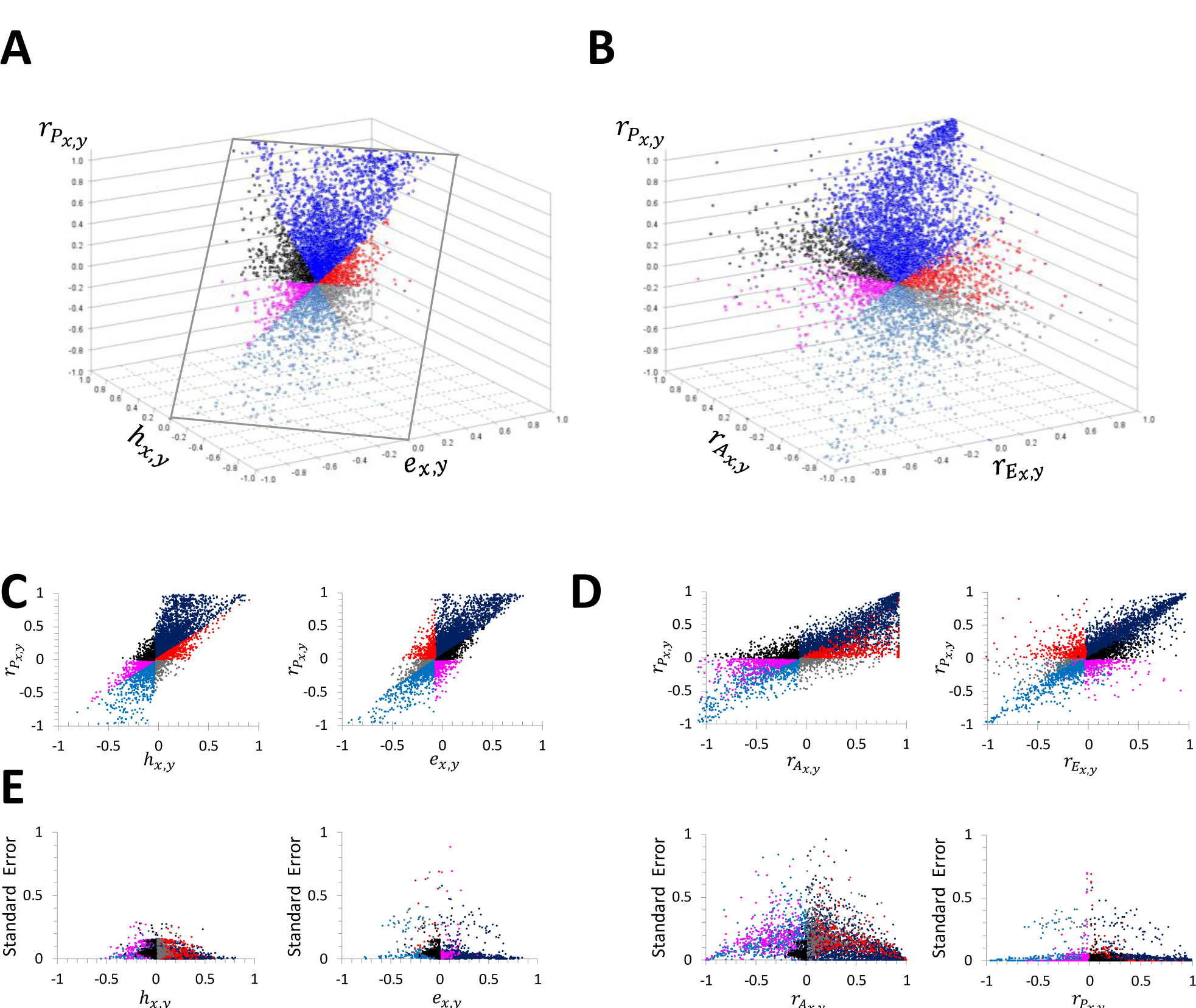
Three-dimensional scatter plot of the phenotypic correlation 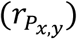 and the (A) coheritability *h*_*x,y*_, coenvironmentability *e*_*x,y*_ on the 3DHER-plane; and (B) the genetic correlation 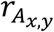 and the environmental correlation 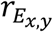. The bivariate plot of the phenotypic correlation against (C) the coheritability, and coenvironmentability, and against (D) the genetic and environmental correlations. (E) Standard errors of the parameters estimators. Total sample size *n* = 6287, color refers to each partition.

**Figure 4:**
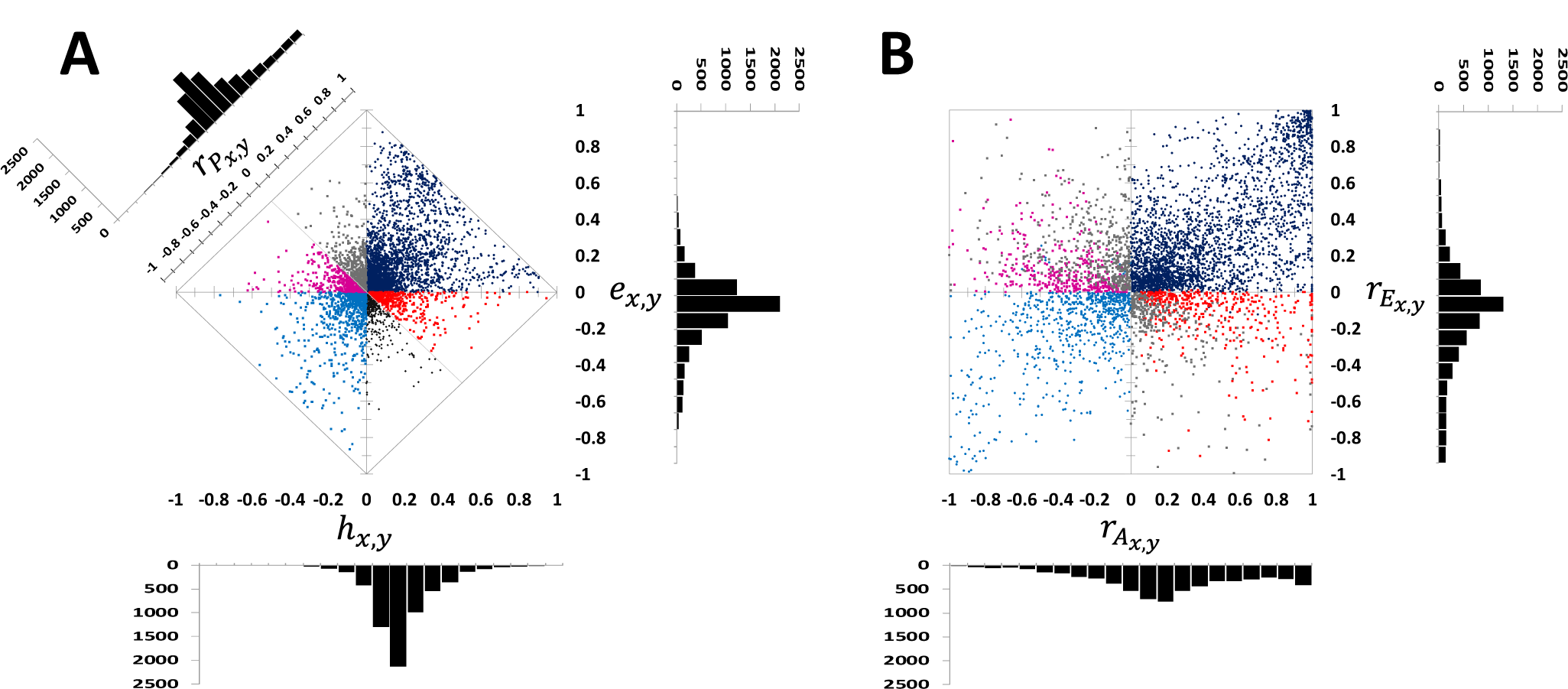
Two-dimensional scatter plot (*n* = 6287) of (A) Coheritability, coenvironmentability, and phenotypic correlations and histograms of the compiled data displayed on the 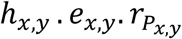 (2DHER)-field. (B) Plot of the genetic and environmental correlations, and histograms. All histograms exhibit a positively skew.

Inspection of the 2DHER-field showed, as expected, well demarcated division between the partitions (4A), a feature not apparent in the plot of the correlations which rather revealed a blurred distinction between partitions (4B). In the plot of the correlations, the data extends throughout its domain occupying most of the area. The histograms approximate a normal distribution, though the shape of all of them appeared somehow skewed towards the positive side.

## Multinomial Test

The abundance of the data varied significantly by partition. Partition *S*_+1_was the most populated (*n*_*S*+1_= 3312) amounting to 53% of the data, which was followed by *S*_-1_(*n*_*S*-1_= 880, or 14.2%). The occupancy in partitions *S*_+2_(*n*_*S*+2_= 617, 0.099%) and *S*_-3_ (*n*_*S*-3_ = 562, 0.09%) were approximately similar. The lowest count were found in partitions *S*_-2_ (*n*_*S*-2_ = 473, 0.076%) and *S*_+3_ (*n*_*S*+3_ = 367, 0.059%). The count in partition *S*_0_ was 67 and was not considered in further calculations. Both the chi-square and the chi-LRT gave qualitatively similar results so here I present only the results obtained using the Chi-Square test. A goodness-of-fit test for multinomial distribution under the null hypothesis 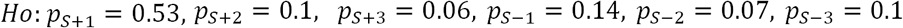, showed that the data fitted reasonably well the multinomial model 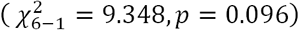 (Supplementary Information Appendix 6B).

### Multiple Regression Analysis

The purpose of these analyses was to evaluate the consistency of model equations and parameters, generated from the use of the whole data set and by partition. The phenotypic correlation was modeled as a function of coheritability-coenvironmentability (Model 1), and genetic-environmental correlations (Model 2). Figure 5 presents the regression parameter estimates corresponding to the genetic factor (*β*_1_), the environmental factor (*β*_2_), and the *R*-square (*R*^2^) for each model. Model 1 showed complete consistency and uniformity (*β*_1_ = *β*_2_ = 1; *R*^2^ = 1), as was clearly seen in producing the same linear regression equation regardless if the analysis used the whole data set or of each partitions. The ratio of slopes to in all cases maintained a 1-to-1 relationship. By Equation [3], the intercept was expected to be zero and the regression coefficients (slopes) of *h*_*x,y*_ and *e*_*x,y*_ each equaled to unity (Supplementary Information Appendix 3G), and the results from Model 1 satisfied these requirements. In Model 2, the relationship between the slopes of the genetic and environmental correlation varied widely (the test of heterogeneity of slopes shows that they are effectively different among each other, Supplementary Information Appendix 3G). The relationship 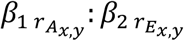, was 1:1.45 (overall) whereas by partition, it oscillated from 1:−0.17 to 1:3.64. Partition *S*_+2_and *S*_-2_ produced models with the lowest *R*-squares (0.299 for *S*_+2_, 0.235 for *S*_-2_). Analyses of residuals and the R-squares of Model 1 captured all of the variability of 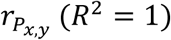, in contrast Model 2 exhibited variable degrees in explaining the variability of 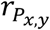, from low to high R-squares (0.23 to 0.94), and each partition yielded a different linear equation.

**Figure 5.**
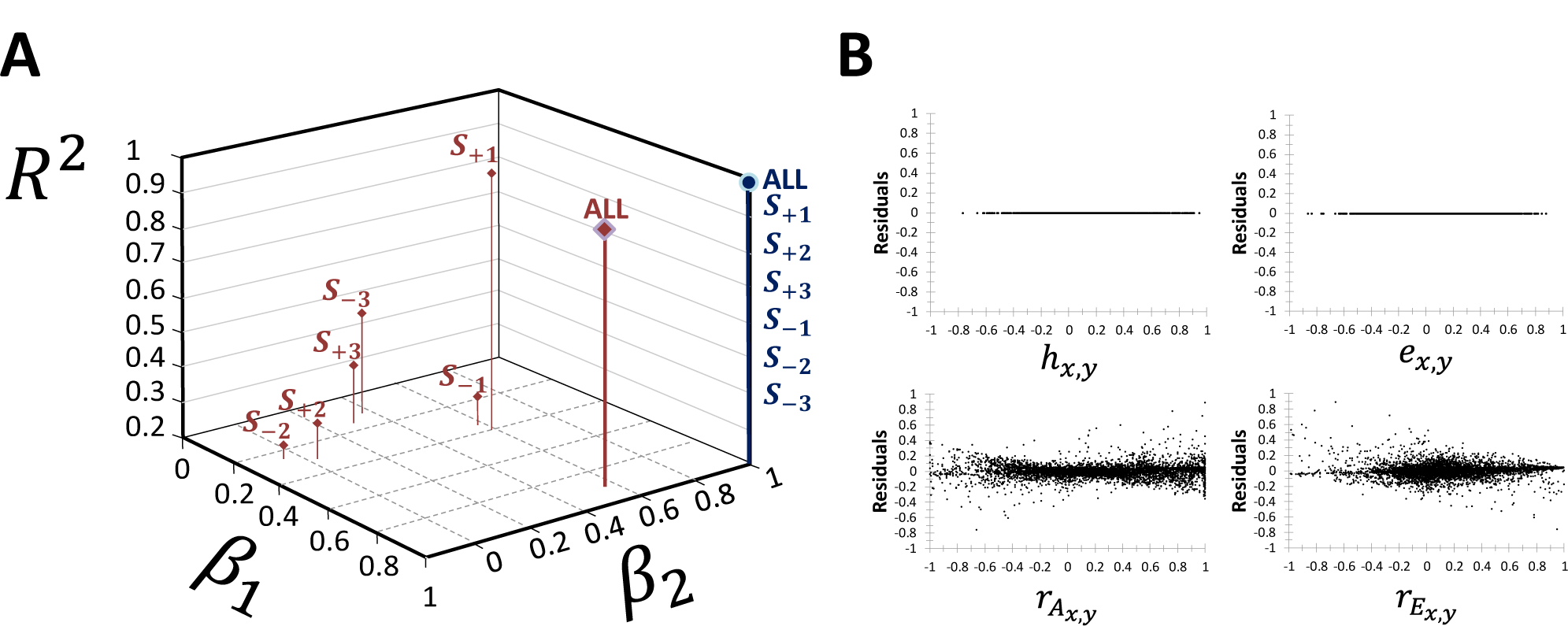
(A) Relationship of the multiple regression parameters (*β*_1_), slope (*β*_1_) for model 1 (blue needle with top circle) and Model 2 (brown needle with rhomboid top). Labels indicate the whole data set (ALL) or the partition subsets. (B) Plots of residual against coheritability and coenvironmentability (Model 1) display low variability dispersed around the horizontal axis indicating that the linear regression model is appropriate for the data. Compare this to the residual plots involving factors 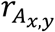 and 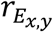 of Model 2.

The results point out to the dependability of the coheritability and coenvironmentability as the appropriate factors directly related to 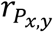, and the consistency of parameter estimates obtained by Model 1 whether using the overall or partition data.

## Analyses on 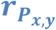 and 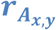

To address whether the phenotypic and genetic correlation exhibit some degree of similarity that would justify 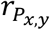 to be an appropriate proxy for 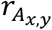, here I now compare the phenotypic and genetic correlation in terms of the Dissimilarity Index (*D*), namely, the degree of departure of their numerical values. A correlation analysis of the overall data set showed that these two variables maintained a high statistical association (*r* = 0.821, *p* < 0.0001, *n* = 6288). The regression of 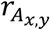 on 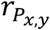 had a slope of 1.14 (standard error 0.01, R-square 0.67, Supplementary Information Appendix 3F). Approximately 2% of the data had a dissimilarity equal to zero, indicating that 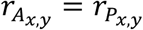, and if a *D* ≤ 0.03 represents an acceptable level of similarity between 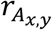 and 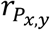, it occurred in 13% of all comparisons. However, there was a considerable departure of their values at the individual level. The mean of the absolute disparity *D* index was 0.181 (standard deviation 0.175, *n* = 6288) across all comparisons. In 15% of the data, 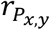 and 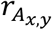 differened in sign (partitions *S*_-3_ and *S*_+3_). When both 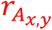 and 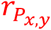 were positive 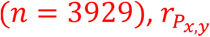 was, on average, 130% the magnitude of 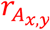. When both 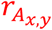 and 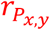 were negative 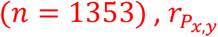 was 129% the magnitude of 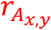. The relationship 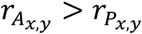 held in 58% (*n* = 3621) of all pairwise comparisons, 70% of the which were in partition *S*_+1_, and 100% in partitions *S*_+2_and *S*_+3_. Around 40% (*n* = 2526) of the overall comparisons involved 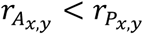, which was found in 67% of comparison in partition *S*_-1_, and 100% in partitions *S*_-2_ and *S*_-3_. If the disparity between 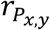 and 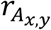 were largely due to measurement error of the genetic correlation, then it would be expected that the squared disparity (*D*^2^) would approach the sampling variance of 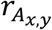. *D*^2^ and the sampling variance of 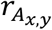 had a correlation not different than zero (*r* = 0.17, *p* = 0.001) indicating they were not strongly associated, and in 59% (*n* = 3353) of the cases that reported standard errors of the genetic correlation, *D*^2^ displayed a value larger than the estimated sampling variance of 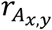. Under the null hypothesis Ho: the difference between the median of *D*^2^ and the median of 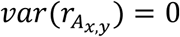, versus H1: median difference ≠ 0, the Wilcoxon Rank Sum Test had a critical value 3138479, test statistic 11.65, *p* < 0.0001, shows that there is enough evidence to reject *H*_0_ that the median of *D*^2^ and 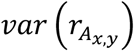 are quite similar. In addition, the Kruskal-Wallis test rejected the null hypothesis that there is no difference among partitions in mean rank of either *D*^2^ or the 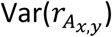. (Supplementary Information Appendix 3I). Therefore, differences between genetic and phenotypic correlations cannot be entirely explained by sampling error alone. The Disparity Index displayed its largest values when 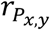 was around zero, and progressively decreased as the 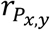 approached the limits of its domain (Figure 6A). The frequency distribution of the *D* values fitted a triangular model (Figure 6B, Supplementary Information 7.6). The relationship between 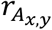 and 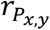 differed greatly by partition (Figure 6). With the goal to evaluate the capacity of the phenotypic correlation to predict the value of the genetic correlation, the simple regression of 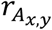 on 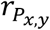 (Figure 6C) resulted in a model where 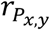 a low capacity to explain the variability of 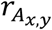 (R-square 0.674) and a slope equal to 1.141 (SD 0.01, p<0.0001, 95%CI [1.121,1.605], Supplementary Information Appendix 3F). Analyses by partition showed that the equations of the regression lines and the R-squares varied widely. In 4 out of the 6 partitions the R-squares were below 0.3. The largest R-square values were 0.73 (*S*_+1_) and 0.63 (*S*_-1_) which correspond to partitions where 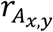 on 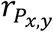 shared the same sign. The lowest R-square were 0.005 (*S*_+3_)and 0.001 (*S*_-3_), corresponding to partitions were both phenotypic and genetic correlations differed in sign.

**Figure 6.**
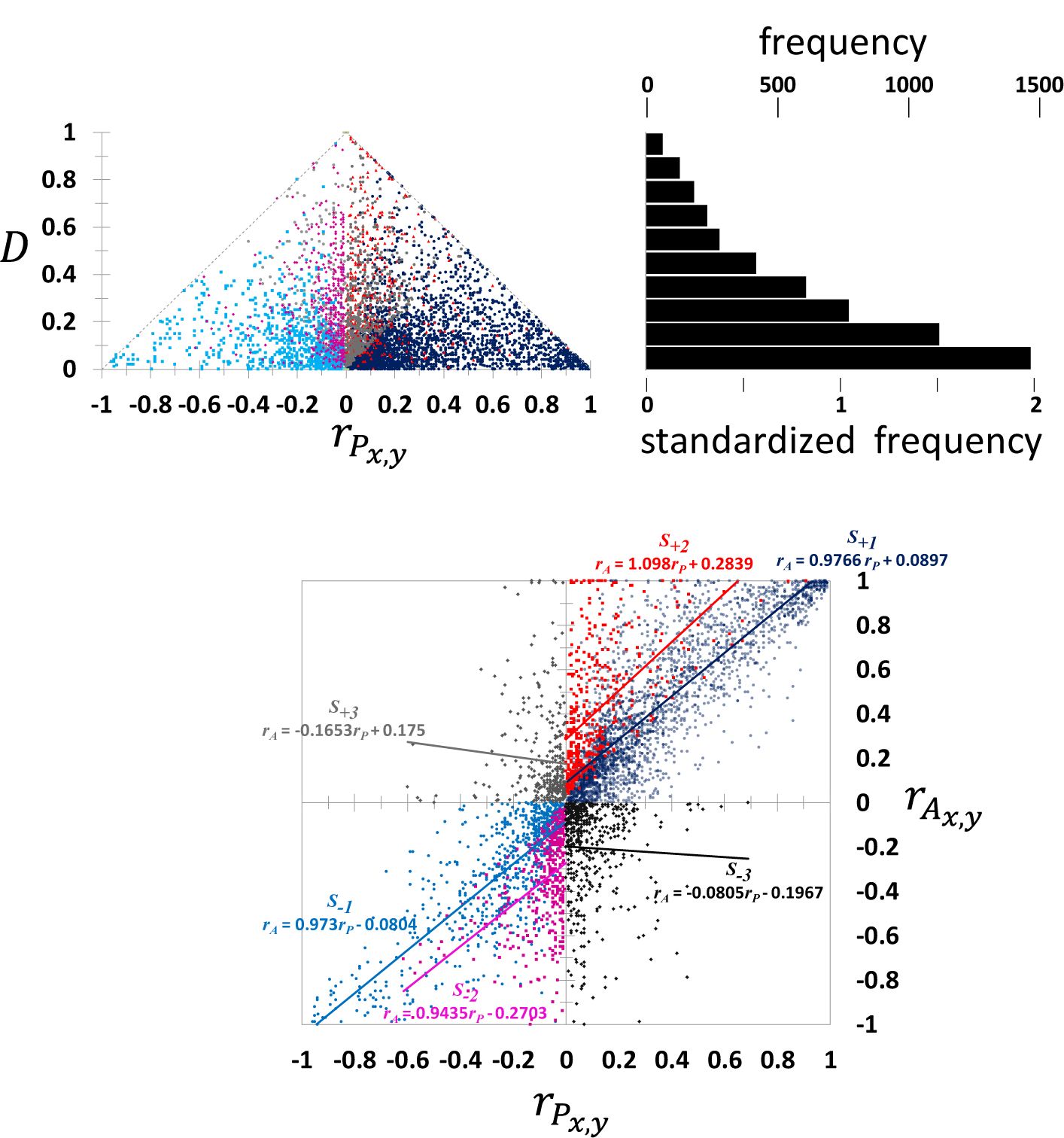
(A) Disparity Index *D* as a function of the phenotypic correlation value. Each dot represents the absolute value of the difference between 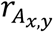 and 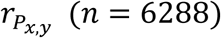. (B) Histograms reflecting the frequency of occurrences of comparisons with Disparity Index values within specified ranges. (C) Summary graph of the simple regression between 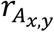 (scalar dependent variable) and 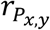 (explanatory variable) showing the model equation and the R-square.

### Illustrative Examples

To illustrate the application of the decomposition of the phenotypic correlation into the coheritability and coenvironmentability, data from published articles were used to gain insight into relevant topics of modern biology. Further examples can be found in the Supplementary Information Appendix 3.

#### Example 1. Changes due to selection over generations

Ramniwas et al. (2013) tested the hypothesis that abdominal melanisation in *Drosophila melanogaster* enhances desiccation resistance. They performed a series of upward and downward selections in the lab and reported the direct response in terms of abdominal coloration change (R_melanisation_) after five generations. In addition, correlated responses in several physiological traits related to water stress were also measured, including cuticular water loss (CR_cuticular_water_loss_). Figure 7 presents the results obtained from upward selection in female flies and the direct (R_melanization_) and correlated response (CR_cuticular_water_loss_) plotted against the selection differential (*S*_*C*_) for direct selection on melanisation. It can be seen that as the individuals were becoming darker, the cuticular water loss decreased concomitantly. Employing the regression slope of cumulative response against selection differential (*S*_*C*_) for darker individuals, the realized heritability for cuticular melanisation was 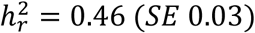. To determine the realized coheritability between melanisation (trait *x*) and cuticular water loss (trait *y*), a regression line was fitted having the correlated response of cuticular water loss as dependent variable against *S*_*C*_ as independent variable (Figure 7). The slope, *b* = −0.2673 (*SE* 0.012, *CI*_0.95_ [−0.27, −0.22]), represents half the coheritability of the traits 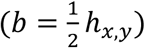 because only measurements obtained from female individuals were used. Therefore, the coheritability between melanisation and the rate of water loss was *h*_*x,y r*_ = −0.5346(*SE* 0.024). This negative coheritability is in agreement with melanisation-dessiccation hypothesis: darker individuals retain more water and are more resistant to desiccation.

**Figure 7.**
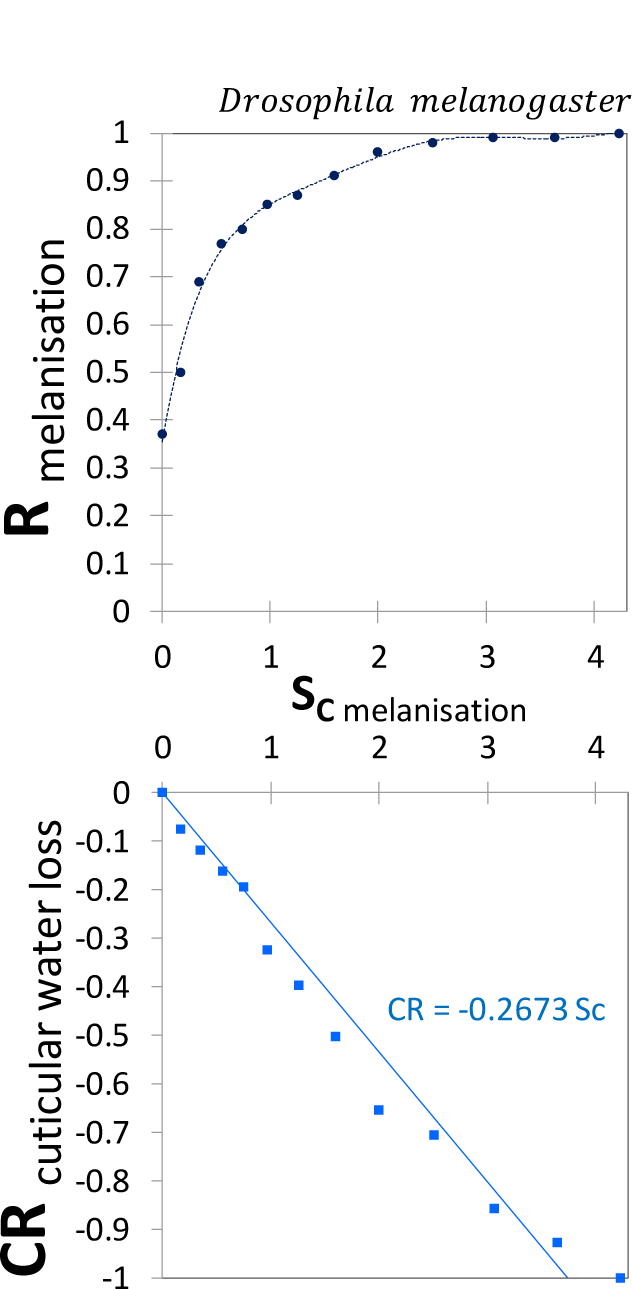
Direct response to melanization and the correlated response of the rate of cuticular water loss among female flies of *Drosophila melanogaster* (Source Ramniwas et al. 2013).

It is important to recognize, in a selection experiment, that each generation of selected parents constitutes a sample of the genetic correlation, and its value in successive generations of selection depend upon one another sequentially, therefore changes in the value of the genetic correlation will have some of the properties of a random walk (Gromko 1995), and manifest a large standard error.

#### Example 2. Changes through development

How do the contributions of shared genetics and shared environment affect the phenotypic correlation through development? Are the trajectories of the phenotypic correlation, coheritability and coenvironmentability informative about relationships between traits? Here I present the results derived from two field experiments. Diao et al. (2016) studied growth in Japanese larch (*Pinus kaempferi*) in order to evaluate the optimal age for early selection, an aspect particularly important for a long-term plantations. To this aim, the authors determined phenotypic and genetic parameters of absolute height measurements obtained in different ages and compared them to the one at age 16 (Figure 8A). All correlations (phenotypic, genetic and environmental) increased as the trees grew, the phenotypic correlation reached 10% of its maximum at age 11 and the genetic correlation around age 8. The coheritability and coenvironmentability also gradually increased and reached 10% from its maximum at age 6, which reveals that no significant gain from selection can be achieved after this age, setting an age earlier than the one based on the genetic correlation.

**Figure 8.**
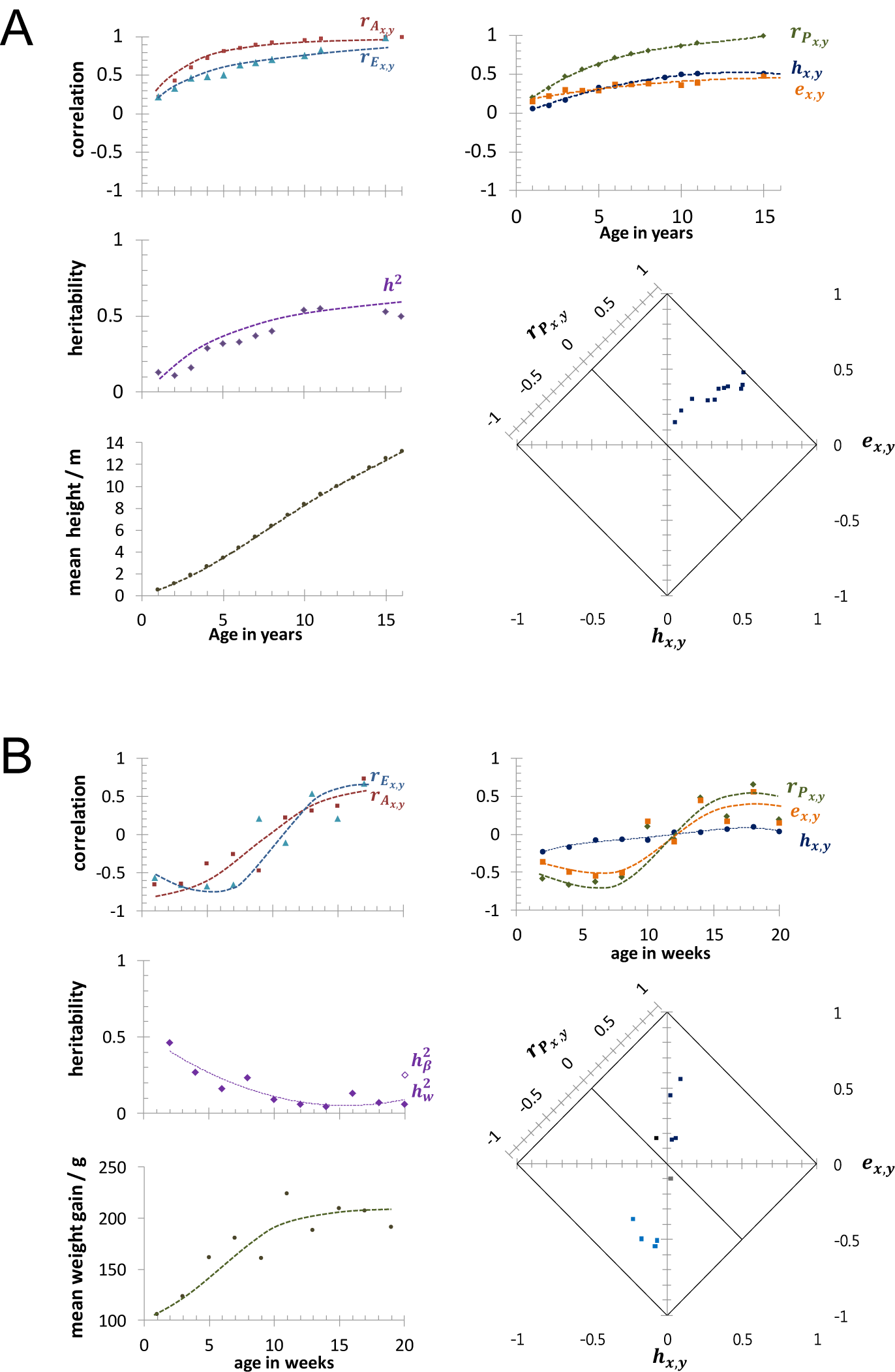
Age-dependent trend in genetic parameters in (A) height growth in japanese larch (*Larix kaempferi*) (Data source Diao et al. 2016), and (B) weight gain (increments) in Korean native chicken (*Gallus gallus domesticus*) (Data source: Manjula et al. 2017).

Another example involved incremental weight gain rather than absolute measurements of weight of Korean native chickens subject to a common diet (Manjula el al. 2017). The aim of the study was to find the best age to make selections that would predict growth at age 20. Measurements of weight were obtained from a number of individuals and subject to a logistic growth model, whose parameters were also treated as traits. Data presented in Figure 8B uses data of 2-week weight increments and the β parameter of the logistic model that captures the asymptotic mature weight gain. The heritability of weight increment decreased consistently as the animals aged. The genetic correlation between the weight increment and the β parameter from early life up to age 8 were negative, and from age 12 to late in life were positive. The phenotypic correlation followed a trend similar to the trajectory of the coenvironmentability indicating that the association between these traits is mainly influenced by common environmental factors. To elucidate what led to changes in the trait-trait relationship between ages 8 and 12, going from negative early in life to positive values late in life, it would necessitate to inquire from other lines of evidence in the metabolic, physiological, and developmental areas. The coheritability had minimal contribution to the phenotypic correlation, and varied in very narrow range of low values (−0.2 to 0.1). One can conclude that weight gain can be better achieved through poultry rearing practices (e.g. diet, feeding regime) than through genetics.

Analyses of longitudinal data present challenges because repeated measures from the same subject are often (auto)correlated, and cannot be assumed to be independent, such is the case of growth traits especially when measurements in certain point in time contain the measurement previously obtained. It would benefit to use suitable and informative covariates to help adjust values. The stochastic trends of observations across age/development/time, shows that any measurement at only a single time point, may over-or underestimate the genetic and environmental contributions to the 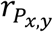.

#### Example 3. Modularity and Integration

The rationale of morphological integration states that functionally and developmentally related exhibit high levels of phenotypic correlation (Olson and Miller 1958), so that traits belonging to the same functional and/or developmental group are genetically more integrated than traits with different functions or developmental origins (Conner and Via 1993, Waitt and Levin 1998). Willmore et al. (2009) conducted a quantitative genetics study designed to compare patterns of mandibular integration between baboon and mouse. The traits were distances between homologous landmarks in the mandible of each species. Heritabilities of the traits as well as the correlations 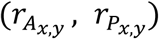 between them were determined in each species separately. Three cases are presented here that reveals the degree of insight provided by the decomposition of the phenotypic correlation. The first case deals with two traits whose genetic correlations are higher than the phenotypic correlation in both species. In the incisive alveolar module, the correlations 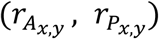 between the distances *sms-idp* and *sms-idi* were (0.911, 0.838) in baboon, and (0.862, 0.567) in mouse (displayed as a triangle in the 2DHER-field, Figure 9), which by a dimple examination of their magnitudes, it would suggest of a strong genetic influence. The use of the coheritability *and* coenvironmentability expand the inferential space, and point out that most of phenotypic correlation between these traits in baboon was due to the coenvironmentability (75%*r*_*p*_ or 0.624), whereas in the mouse due to the coheritability 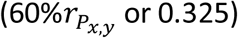. This aspect would otherwise have been overlooked by merely comparing the magnitudes of the correlations.

**Figure 9.**
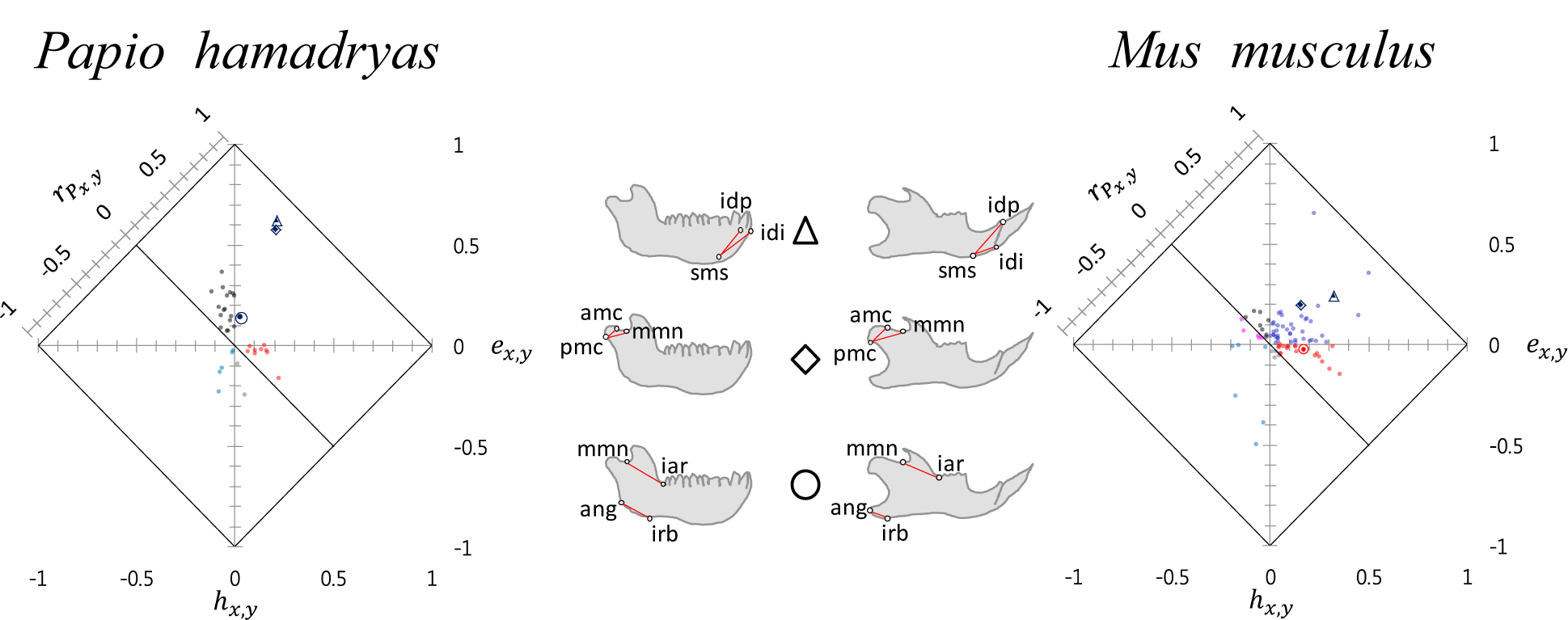
Modularity and integration. Comparison of mandibular integration between baboon (*Papio hamadryas*), and mouse (*Mus musculus*). At the center are depiction of the mandibles and the distance traits: triangle=*sms-idi* and *sms-idp*; rhomboid=*pmc-amc* and *pmc-mmn*; circle=*ang-irb* and *mmn-iar* (Data source: Willmore et al. 2009).

A second case involves trait-trait associations entirely due to either coheritability or coenvironmentability. In both species the distances *ang-irb* (angular process) and *mmn-iar* (coronoid process) are weakly correlated at the phenotypic level (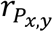: in baboon 0.213, in mouse 0.143; circle)(Figure 9A). However, in baboom the coheritability had a negligible contribution to 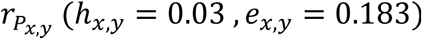, whereas in mouse the coheritability amounted to most of the phenotypic correlation (*h*_*x,y*_ = 0.170, *e*_*x,y*_ = −0.027).

Lastly, this case exemplifies that trends in one species cannot be extrapolated to related ones. The phenotypic correlation, coheritability and coenvironmentability in the alveolar *sms-idi* and *sms-idp* (Figure 9A, triangle) and in the condylar *pmc-amc* and condylar-coronoid *pmc-mmn*; (rhomboid) have similar values in baboon and they that plot almost together in the 2DHER-field. In mouse, however, these traits differ in terms phenotypic correlations and the relative amount contributed by the coheritability and coenvironmentability.

Without disregarding the high proportion of negative genetic correlations in the baboon (around 40%), the allometric data clearly indicates that there is a distinct pattern of modular development operating in the mandible of each species, which should be expected given their different ecological niche, feeding adaptations, mastication process, habits, and diet.

#### Example 4. Trade-offs of life history traits

A fundamental tenet in the study of fitness in natural populations is that life-history traits limit the individual to attain simultaneously maximal growth, reproduction, and survival (Lande 1982). A negative genetic correlations among life-history traits is often adduced to indicate a trade-off that constrain the correlated response to selection in natural populations (Stearns 1989, Zera and Harshman 2001). Figure 10 presents results of coheritability and coenvironmentability among life-history traits. Knowing that the sign of the coheritability is conferred by the genetic correlations, Figure 10 shows sufficient examples of life history traits relationships in both negative and positive directions, congruent with findings in other studies (Roff 1996, Kruuk 2003). Noteworthy is the study of Dutilleul et al. (2015) who analyzed fertility and survival of the nematode *Caenorhabditis elegans* subject to contaminated environments. They found that genotypes that achieve faster sexual maturity (early growth) were more fertile, but had a reduced life span. Their data shows that fertility (*x*) and early growth (*y*) were correlated at the phenotypic level 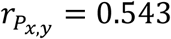, with approximately similar contributions by coheritability (*h*_*x,y*_ = 0.248) and coenvironmentability (*e*_*x,y*_ = 0.295). However, with late growth, fertility exhibited a negative phenotypic correlation 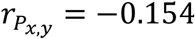, which was mostly due to the coheritability (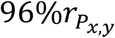 *or* - 0.148). The decrease of the phenotypic correlation between fertility from early to late growth could be explained by the antagonistic pleiotropy model which states that if the genes that promote fertility early in life are the same that cause deleterious effects late in life, then fitness is maximized for fertility when the organism is young, at the expense of detrimental performance as the individual age. It would be of great interest to explore whether the described trade-off results from a covariance of both traits to a third unmeasured trait, such as the individual’s body size. If individuals reach maturity early at a relatively small adult size (implying shorter growth period) then it would have the added benefit to produce progeny early to compensate for their short lifespan (Charmantier et al. 2006). Generally, traits associated to adaptive response resulting in enhanced fitness (e.g. fertility) will provide an advantage to the individuals that manifest such traits by favoring their reproduction, whose timing and plasticity would depend on the environment. Although selection pressure will always tend towards fitness increase (Fisher 1930, p. 35), it does not imply that fitness necessarily cause the competitive ability of a population to be superior with respect to other populations not interbreeding with it (Lerner and Dempster 1948).

**Figure 10.**
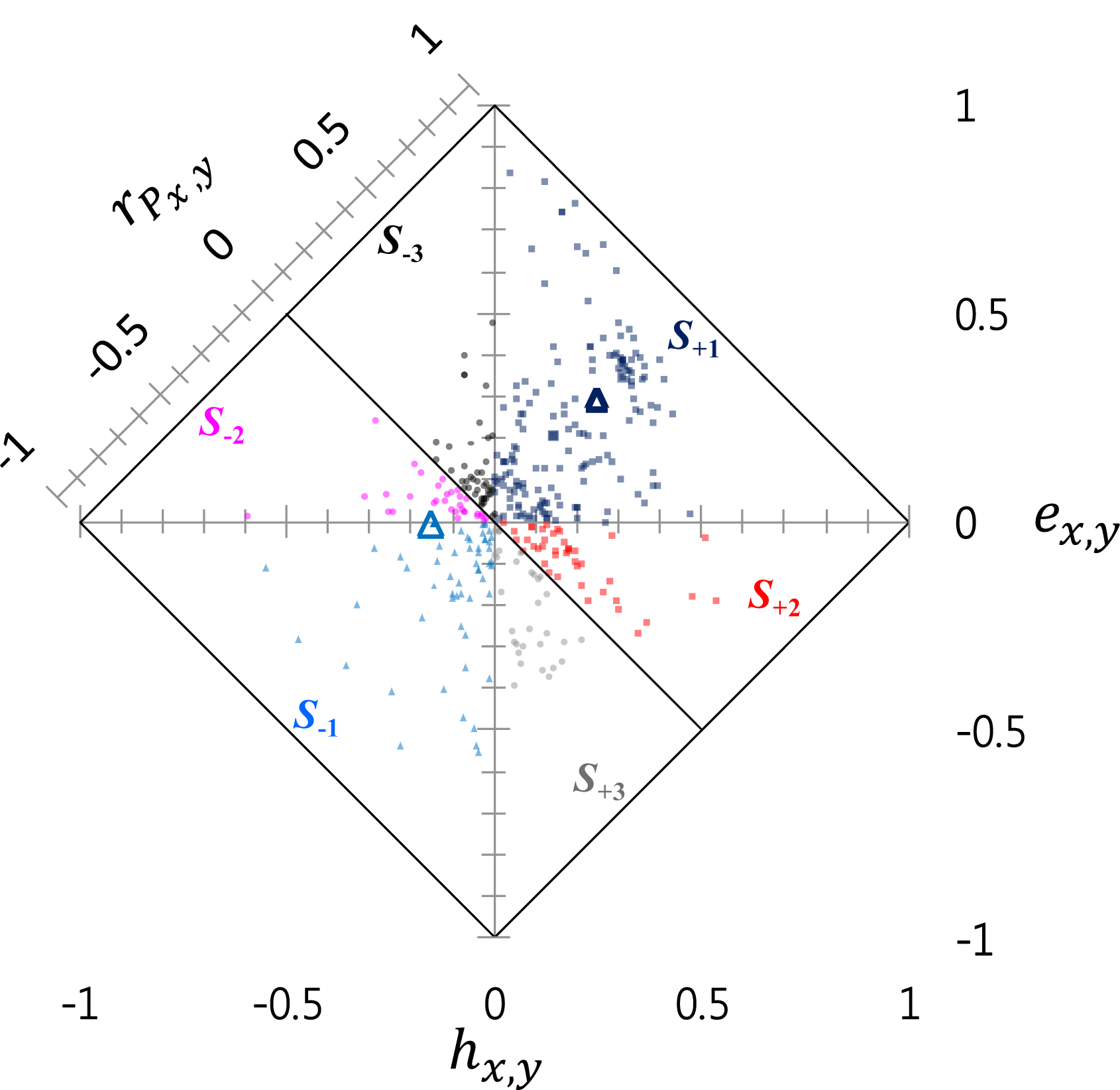
Trade-offs in life history traits in animals. Highlighted are the contrasting results of fertility and growth in *Caenorhabditis elegans* (Dutilleul et al. 2015). Potential trade-offs may be present between fertility and early growth (triangle solid). Fertility and late growth (triangle clear). (Data source: compiled from the literature).

The visualization of phenotypic, genetic, *and* environmental information in the 2DHER-field does facilitate the analysis in a single graph. There is an increasing recognition that the environment has a direct influence on the quantitation of genetic parameters underlying life-history traits, therefore it adds value to use the completeness of the data in these analyses to evaluate changes in environmental conditions can influence genetic interactions and covariances among life-history traits (Sgrò and Hoffman 2004).

#### Example 5. Gene co-expression

Gene expression is a major contributor to phenotypic co-variation in human complex trait (Gandal et al. 2018, Lee et al. 2012). Quantitative genetics of gene expression considers the abundance of a particular mRNA transcript as a “trait”. Gene expression can be influenced by a variety of biological factors, which are especially susceptible to interact with the environment. To gain insight into the mechanisms that control gene expression, Lukowski et al. (2017) investigated the proportion of co-expression between genes in whole blood samples. Figure 12 displays the results of thousands (*n*∼14000) of bivariate comparison in the 2DHER-field, and show that gene expression levels explore all sign configurations among 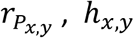, and *e*_*x,y*_. Most remarkable is the apparent lack of data points along the transect −0.1 < *h*_*x,y*_ < 0.1, which distinctly separates and defines two major patterns of coexpression based on the sign of the coheritability. Though classification is not the focus of this work, the visualization in the 2DHER-field nevertheless reveals a level of insight and could serve as a guide to further examine functional modes of co-expression.

**Figure 12.**
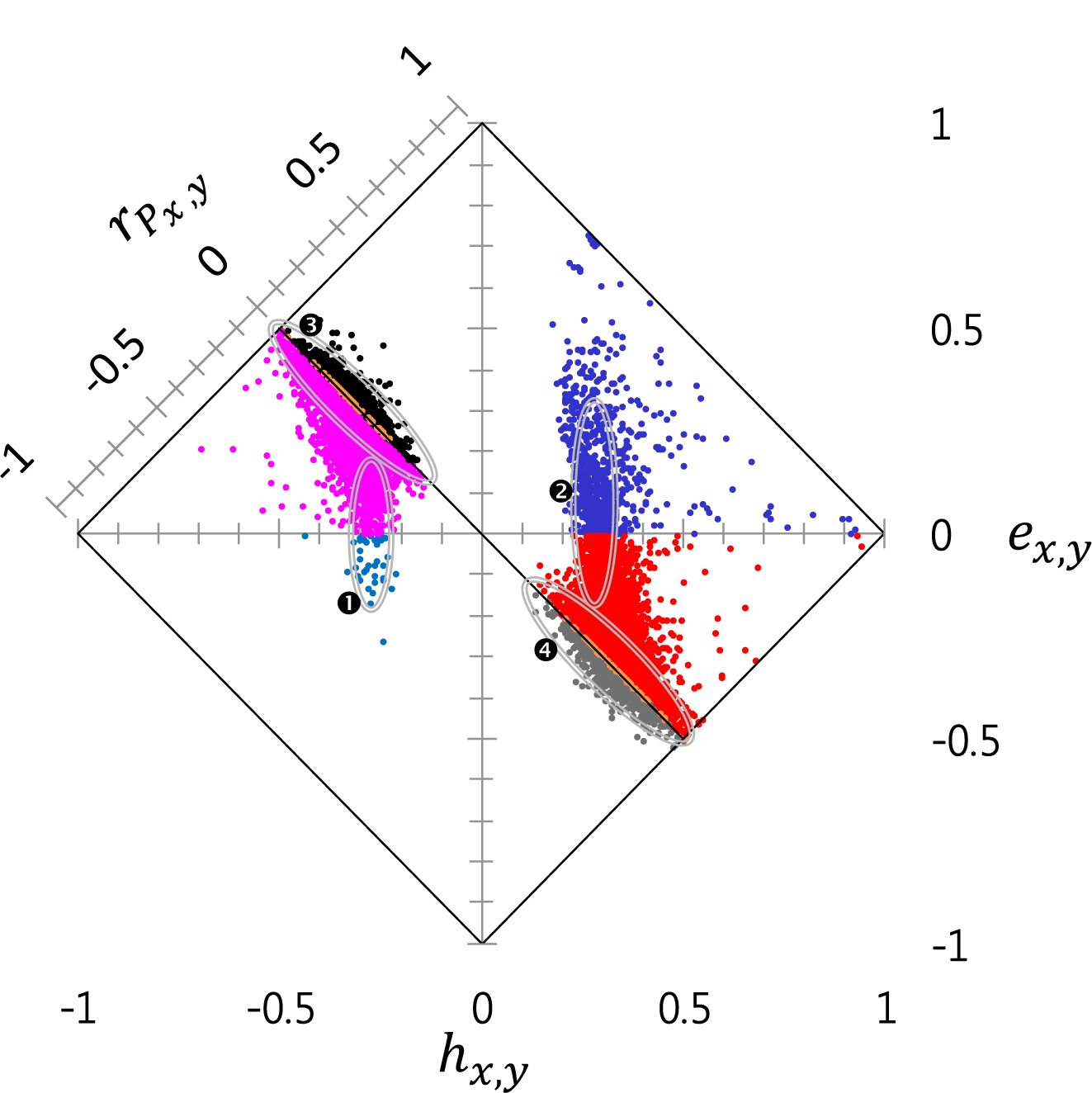
Shared genetic and shared environmental architecture of transcription in human peripheral blood. Significant genetically correlated probe pairs (study-wide false discovery rate *DR* < 0.05). Of these 14991 data pairs, 14020 (93%) are located on different chromosomes, 7886 have a positive coheritability and 7105 a negative coheritability (Data source: Lukowski et al. 2017).

Here special attention is placed on data points circumscribed by the ellipses in Figure 12. Groups 1 and 2 enclose observations with low coheritability, delimited to the interval −0.3 < *h*_*x,y*_ < −0.2 (Group 1), and 0.2 < *h*_*x,y*_ < 0.3 (Group 2). In these groups, much of the phenotypic correlation is under varying influence of the coenvironmentability. In Group 1 the phenotypic correlations are negative, mostly occupying partition *S*_-2_. The positive phenotypic correlations associated to Group 2, includes data with negative (partition *S*_+2_) and positive (partition *S*_+1_) coenvironmentabilities. Group 3 and 4 are found along a narrow interval around the null phenotypic correlations 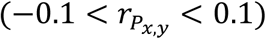. For the 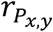 to be close to zero, the coheritability and coenvironmentability must have similar magnitudes but with different sign. Numerous instances where the phenotypic and genetic correlation differed in sign were found in partitions *S*_+3_ and *S*_-3_. The values of the genetic correlation averaged −0.019 and oscillated widely throughout all its domain [-1, 1] as can be seen by the large standard deviation (SD) 0.883, a fact that is corroborated by the large disparity values (mean *D* = 0.84, *SD* = 0.12). The coheritability allocated more or less equally to the positive and negative realms of its domain. All this indicated that the genetic correlation when the phenotypic correlation was close to or not different than zero differed greatly in magnitude and direction with respect to 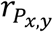. When sample sizes are large relative to the number of variables, the graphical method is likely to provide the most insight. Gene expression is subject to many factors including methodological bias (see Pereira et al. 2009), and is known to be responsive to environmental clues that alter mRNA abundances accordingly (Grishkevich and Yanai 2013). Therefore, it would be of great interest to conduct assessments of loci known to be up-or down regulated under different environmental conditions (Li and Burmeister 2005).

## Discussion

Biological studies rely on the phenotypic correlation between traits as an important measurement of association between the phenotypic values of two traits. There is an added interest to distinguish how much of the correlation can be attributed to additive genetics and how much to the environment. This paper presents a method that captures both the genetic and environmental contributions to the phenotypic correlation, based on quantitative genetics theory, and aims to formalize the decomposition of the phenotypic correlation into the coheritability and coenvironmentability, to further present its mathematical and statistical properties, and to provide a visualization tool capable to handle all the pertinent variables (i.e., 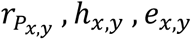) concurrently.

### Coheritability and genetic correlation

First, it is important to distinguish between coheritability and genetic correlation. Coheritability quantifies the contribution of additive genetic effects to the phenotypic correlation between two traits, and relates the genetic covariance to the bivariate phenotypic variability (i.e. geometric mean of the phenotypic variances, see Equation [4]). The genetic correlation measures the association between the breeding values of two traits and relates the genetic covariance to the bivariate *genetic* variability (i.e. geometric mean of the genetic variances, equation [6]). Thus, coheritability and genetic correlation do not share the same inferential space. Coheritability, together with the coenvironmentability have an additive relationship with the phenotypic correlation, and is therefore most proximal to the phenotypic correlation, an aspect not shared by the genetic and environmental correlation. In addition, the coheritability has predictive ability in regard to correlated response to selection. The formula of the coheritability (Equation 7) involves the modulation of the genetic correlation by the square root of the trait heritabilities, the latter being independent random variables, coheritability and genetic correlation, notwithstanding sharing the same sign, are not numerically similar nor follow necessarily the same trend nor rank.

### The biological and the biometrical concepts of coheritability

To further conceptualize the meaning of coheritability, it is useful to construe the term in a biological sense as well as in a biometrical sense. Biological coheritability is grounded on a gene-centric approach that rests on the degree of correspondence between specific phenotypic traits and the allelic content of identified, causal genes influencing them. This includes the effect of all loci that affect both traits (i.e. pleiotropic alleles), and the effects of non-randomly-associated, tightly linked loci acting on each trait singly (i.e. loci in linkage disequilibrium) (see Zhang et al. 2018, Tyler et al. 2009), as well as the loci that regulate their expression. Trait correlations caused by pleiotropic alleles are adaptive, different than trait correlations due to linkage disequilibrium, which could erode through recombination (Saltz et al. 2017).

The biometric concept of coheritability is based on population-level, statistical principles, and it recognizes that the observed association of traits is the net realization of a multiplicity of shared and not shared genetic factors influencing polygenic traits. It expresses the notion that two traits are inherited together (i.e. maintain the resemblance seen in the parental generation) due to the aggregate additive effect of genetic factors affecting the traits, including those acting on each trait individually, others on both traits. In this sense, the biometrical coheritability derived from multivariate analysis becomes a hypothesis on the degree of contribution of genetics to the phenotypic correlation of traits.

Biological coheritability deals with the genetic architecture of quantitative traits based upon identified causal gene variants, quantitative trait loci, additive and dominance effects, mapping, coding and noncoding sequences, and by understanding its variability due to the action of modifier genes adjusting its penetrance and expressivity, the occurrence of recombination events, or by the activity of regulatory elements in other parts of the genome (Short et al. 2018, Mackay et al. 2009). Discerning between pleiotropic and coincident linkage is an area of intensive experimental (Wagner and Zhang 2011, Solovieff et al. 2013, Gardner and Latta 2007) effort associated to a diversity statistical and computational (Hackinger and Zeggini 2017, Schaid et al. 2016, han and Pouget 2015, Yang et al. 2015, Carter et al. 2007) approaches involving sequencing, fine mapping, and functional characterization of the gene and gene product (Flint and MacKay 2009). The multifunctionality of a pleiotropic gene can be mediated by alternative splicing, by RNA editing, and tissue specific expressions. Pleiotropy can also exhibit multiple phenotypic consequences of a single molecular function, such as those acting on multiple pathways (Singh and Shaw 2012).

The biometrical coheritability invokes quantitative genetic concepts such as heritability, correlation (genetic, phenotypic), genetic factors. Its variability depends on population structure, the additive and phenotypic variances of each trait, and the genetic covariance between the traits (Carey 1988). The biometrical concept of coheritability does not carry the assumption that the genetic factors involved in the expression of the traits are related to genes with an additive mode of action, nor the existence of genetic covariance precludes the effects of genes with any degree of dominance or epistasis (cf. Huang and Mackay 2016).

Biometrical coheritability is not commensurate with relevance of importance of the trait-trait association. In a recent study on the relationship between amygdala (trait *x*) and emotion recognition (trait *y*), Knowles et al. (2015) determined heritabilities 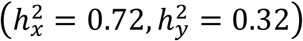 and genetic correlation 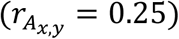 of these traits. It allows us to determine a coheritability *h*_*x,y*_ = 0.12, which appears to be of very low magnitude. Using bivariate linkage and association analyses, they identified the gene *PDE*5*A* (locus 4*q*26) whose gene product is the phosphodiesterase 5 (PDE5) enzyme expressed in various regions of the brain, and has been linked to deficits in memory recognition. As such the PDE5 enzyme has emerged as a potential drug target for treating cognitive deficits and neurological disease (Teich et al. 2016).

Whereas biological coheritability relates allelic variants and traits, biometrical coheritability focuses on trait-trait associations. At the interface is the extensive use of molecular markers (e.g. SNPs) in genome analyses which has occasionally presented loci harboring markers associated to several, sometimes disparate, traits. These cross-phenotype associations could have as underlying cause pleiotropy, linkage disequilibrium, or be artifactual nature (Solovieff et al. 2013). Besides the ‘missing’ heritability problem that arises in the estimation of heritability in GWAS (Yang et al. 2017), Gianola et al. (2015) demonstrated that correlation coefficients inferred using markers can provide a distorted picture of the actual genetic correlation between traits due to the lack of knowledge about linkage disequilibrium relationships between QTLs and markers. Thus, speculating about genetic correlations and even more about its causes (e.g. pleiotropy) using genomic data is conjectural (cf. Lee et al. 2012). Pleiotropy is, by definition, a property of a gene or locus, not of a marker. Therefore, caution must be exercised when interpreting (rethorically and conceptually) findings of marker-based genetic parameters, as to distinctly differentiate the causal (i.e., genes) component from the instrument (i.e., the marker). This distinction is critical.

### Coheritability under a null phenotypic correlation

A phenotypic correlation equal to or not significantly different than zero, does not necessarily imply that both coheritability and coenvironmentability are also zero or non-significant. The coheritability and coenvironmentability could be similar in magnitude but not in sign.

Therefore, an apparent lack of association between traits at the phenotypic level, cannot be dismissed as being of no interest at the genetic level. Disparity analyses revealed that 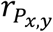 and 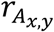 differed the most when the phenotypic correlation is around zero.

Under a null phenotypic correlation, the definition and metrics of the phenotypes must be reassessed. The ability to define meaningful traits in appropriate continuous or categorical scales would help avoid the confounding of distinct characters as a single phenotype (de Villemereuil 2017). This is of particular important in clinical research where a convergence of multiple traits, symptoms, signs in an individual may be a consequence of a cascade of disease causing defects in a complex network of interacting genes, proteins, and metabolites (Park et al. 2009, Emilsson et al. 2018). Yet given the heterogeneity of sources contributing to comorbidity, it is not obvious whether these traits are properties of a disease at the individual or at the population level. Among several options, two approaches can be used to address this problem. One approach is the use of subphenotype groups. A subphenotype is a subset of individuals drawn from a large cohort who feature a specific set of traits in common (Morris et al. 2010). The intent is to ‘concentrate’ individuals with similar trait-trait associations, generally using latent class analyses (Famous et al. 2017, Gårdlund et al. 2018), under the premise that, if the traits have a common genetic basis, their detection would be facilitated, otherwise by ascertaining all individuals simultaneously as having the same phenotype, the causal factors would be overlooked.

Another approach regards a complex phenotype as a composite of many traits, some of them more closely connected to the genetic cause of the complex phenotype. A trait with this characteristic is referred to as an endophenotype. Endophenotypes are widely employed in psychiatric genetics (Iacono 2018, Doyle et al 2005), livestock research (te Pas et al. 2017), immunology (Gregersen et al. 2015), and clinical research (Benyamin et al. 2007). Glahn (et al. 2012) used the absolute value of the coheritability as a criterion to select endophenotypes associated to major depression risk. Similarly, Hammer et al. (2006) proposed to dissect complex phenotypes into traits spanning different levels of biological organization within the individual in order to characterize the full suite of factors that contribute to quantitative variation across cellular, tissue, organ, organismal levels, as well as developmental stages (Cobb et al. 2013). Sun et al (2015) proposed a method to search for a combination of phenotypic characters of multivariate phenotypes and directly maximize the heritability of this combined trait. Though the aim of these models is to enhance the detection of trait-trait associations, it also brings ontogeny as a main factor shaping relationships among traits.

### Rules governing trait-trait associations

Inspection of the scatter plot of *h*_*x,y*_, *e*_*x,y*_ and *r*_*P*_ on the 3DHER-plane graphically showed that, despite the sizeable sample size, not all places in the plane appeared uniformly populated, suggesting that some areas more inaccessible than others such as close to the border where the relationship |*h*_*x,y*_ – *e*_*x,y*_| = 1 holds. At this area, data can only be generated by combining heritabilities and genetic correlations both at very high magnitudes. If the construction of traits entails optimization constraints, then heritabilities and genetic correlation reaching simultaneously extreme values in their domains will not be favored. The occupancy question should be better viewed as a consequences of how traits are constructed, and how ontogeny molds their relationship through life. The organized, sequential, and modulating events of development would manifest varying degrees of genetic and environmental control of two correlated traits (Badyaev and Martin 2000, Saltz et al. 2017). Under this condition, not all corners of the 2DHER-field would be accessible to the same degree.

The developmental process by which traits are constructed derive from genetic (Kellogg 2004) and epigenetic interactions (Peaston and Whitelaw 2006), as well as its interactions with the environment, all combined to shape the phenotypes as emergent products of such interactions. Certainly, phenotypic structures can be mathematical modeled using simple rules (e.g. egg shape by Stoddard et al. 2017, shell coiling by Raup 1966), or conform to empirical allometric scaling such as volume and surface area (Square-Cube Law; Haldane 1926), metabolic rate and body size (Kleiber’s Law; Niklas and Klutchera 2015, Hulbert 2014), speed and body mass (Meyer-Vernet and Rospans 2015). Though a constructionist perspective enhances our ability to account for the relationships among traits, these studies are not meant to simulate the actual biological mechanism of development. Developmental systems are often under strong stabilizing selection to maintain homeostasis, such that patterns of trait-trait covariation through ontogeny or within a particular ontogenic stage generally exhibit conservatism of developmental systems (Badyaev and Martin 2000). Genetic, phenotypic correlations and autocorrelations among traits would reduce independent variation of traits at different ages, limit the number of dimensions in growth trajectories, and overall present a powerful inducement for maintaining certain relationships between traits through ontogeny (Badyaev and Martin 2000).

### The biotic environment and its role in trait-trait association

The manner the coheritability undergoes changes under varying environmental circumstances reveal that the environment can potentially induce changes in the genetic architecture of complex traits (Sikkink et al. 2017), and inferences deduced from studies conducted in one environment cannot be generalized (Aastveit and Aastveit 1993). Stearns et al. (1991) proposed that environmentally induced changes in magnitude and sign of genetic covariances occur when traits are functionally uncoupled at the physiological or developmental levels.

Of particular importance is the need to clarify the extent, if any, of the existence of a covariance between the genetic effects of one trait and the environmental effects of the other. Generally, these terms are surmised to be zero. Nevertheless, instances of indirect genetic effects (IGEs, Bijma 2014) exerted by an individual on the trait values of another individual (e.g. maternal effects), admits that there may exist a partly heritable component in the social/biotic environment when IGEs occur (Ørsted et al. 2017). For instance, in isogentic lines of *Caenorhabditis elegans* nematodes, progeny traits such as fecundity and rate of development are due to maternal-dependent provisioning of vitellogenin protein to the embryos (Perez et al. 2017), parents’ influence juvenile body size by adjusting investment per offspring (Rollinson and Rowe 2015). Traits related to growth, nutrient assimilation, and nourishment of an organism, which are generally assumed to be an intrinsic property of an individual, is directly affected by the microbiome that colonizes alimentary tracts of animals (Org et al. 2017), or the rhizosphere (Nihorimbere et al. 2011, Wissuwa et al. 2009) and phyllosphere (Kembel et al. 2014, Rosado et al 2018) of plants. The microbial diversity present, in turn, is influenced by host traits (Li et al. 2018). Some researchers regard the commensal microbiome as a virtual organ of the body (Valdes et al 2018) whose genome brings forth emergent, novel capabilities (e.g. processing of drugs, resistance to toxins) to the host that would not otherwise exists (Kho and Lal 2018). Associations among traits manifested in an individual are not only the result from their own adaptations but also could derive from community level relationships. Grab et al. (2019) found that closely related pollinator bee species shared many behavioral and morphological traits including body size, plant fidelity, and visitation rate, but not flower-handling ability. On this basis, they were able to predict characteristics of fruit shape malformation from loss of diversity and abundance of pollinator bees in apple plants, and that phylogenetic diversity and species richness best predicted fruit weight and number of seeds per fruit.

### The coheritability and coenvironmentability are the most proximal factors affecting the phenotypic correlation

The coheritability and coenvironmentability are the most proximal factors influencing additively the value of the phenotypic correlation. There is a tendency to surmise that the genetic correlation *per se* will have a preponderant influence on the phenotypic correlation merely because its magnitude is sufficiently large. The theoretical and experimental examples presented in this paper make such proposition untenable. Equation [5] shows that the genetic correlation is embedded within the expression of the phenotypic correlation, where the magnitude of the genetic correlation is modulated by random variables, namely the trait heritabilities. Two traits may display high heritability values, yet be poorly correlated at the genetic level. Highly genetically correlated traits may each display low heritabilities (Drobniak and Cichon 2016). Therefore, the coheritability is not a linear transformation of the genetic correlation (i.e. it does not preserve rank). A given value of the coheritability may be the result of a numerous set of heritability and genetic correlation values, the coheritability is not a monotonic transformation of the genetic correlation because the variables do not possess a one-to-one relationship. This nuanced interpretation of Equation [5] must be contrasted with other views that treat heritabilities as scalars (cf. Cheverud 1988). Therefore the simple comparison of 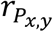 and 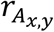 (or their matrices) cannot presume to convey information on the degree on which shared genetics influences 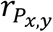. The argument presented here is persuasive for three reasons. First, the phenotypic correlation and coheritability share the same common denominator, a fact that distinguishes them from the genetic correlation, therefore comparison of 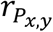 and *h*_*x,y*_ is made on the same basis. Second, the partition of the phenotypic correlation into coheritability and coenvironmentability directly follows, in a standardized form, from the partition of the phenotypic covariance into genetic and environmental covariance, maintaining additivity of the terms. That is a property that 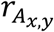 and 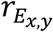 do not have with 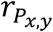 since their denominators involve different and independent random variables. Third, results of the regression analyses involving 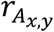 and 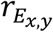 as regressors failed to produce a general relationship that can hold to any data set.

Differing from this approach that compares 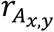 and 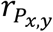 values in their own right, Cheverud (1988) postulated that phenotypic correlations could be used as a proxy for genetic correlations. The regression analyses presented in this study have shown that genetic correlation do not map directly to the phenotypic correlation, and overlooking genetic information (i.e. 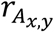) can qualitatively affect inferences (evolutionary, statistical, practical breeding outcomes) in ways a purely phenotypic approach cannot predict or explain (Rubin 2016, Kruuk et al. 2008). Therefore, correspondences between phenotypic-based predictions on genetic outcomes are not robust for all plausible assumptions regarding the underlying genetics of traits (Hadfield et al. 2007). By observing equation [5], to consider 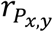 as a suitable predictor of 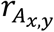 assumes that the phenotypic and genetic covariances possess the same sign, that the phenotypic covariance is entirely genetic in nature (i.e. no detectable environmental covariance is present), and that the heritabilities of both traits are unity, all becoming very demanding assumptions (Supplementary Information section 7.4). Thus, phenotypic data is not an adequate predictor of underlying genetics of natural or breeding populations. Since both genetic and environmental correlations combine together to give the phenotypic correlation, therefore, as Falconer and MacKay (1996, page 314) stated, ‘this dual nature makes it clear that the magnitude and even the sign of the genetic correlation cannot be determined from the phenotypic correlation alone’.

### Uses of the decomposition of the phenotypic correlations

Besides the applications presented in the illustrative examples of this work, there are further uses of the coheritability and coenvironmentability. These parameters could be utilized as objective means to construct common-cause classification of diseases that cluster together diseases with genetic and environmental similarities under the premise that shared genetics and environment would manifest similarities between diseases that have a common-cause nosology (Wang et al 2017). A promising application is enhancing the capability to recognize biomarkers and traits that are readily impacted by a disease challenge (Boulton et al. 2018, de Matos et al. 2018). Clinical diagnostics if construed as a type of indirect selection would benefit by helping detect measurable traits (i.e. phenotypic characters, biomarkers) that are both easily accessible, (positively or negatively) correlated with disease traits, and coheritable with it (see Yin et al. 2017, Mehr et al. 2018, Ngo et al, 2018; cf. Bastaranche et al. 2018).

Also, in the area of genomic prediction, knowledge on the degree genetic effects contribute to the phenotypic correlation can be utilized to simultaneously improve prediction accuracy of parameter estimates of breeding values of multiple traits (Crossa et al. 2017, Montesinos-Lopez et al. 2018). Another area where the decomposition of the phenotypic correlation would be particularly useful is in high-throughput phenotyping. It would exploit the data in their full capacity including the genetic interdependencies of traits (Xavier et al. 2017, Sun et al. 2017).

The coheritability of somatic, homologous traits common to both sexes would bring insight in the study of sexually dimorphic species especially in the manner males and females respond to selection in different directions given the tendency towards gender-optimal phenotypes (Mank 2009, Poissant et al. 2009, Chippindale et al. 2001). Application of the decomposition of the phenotypic correlation in morphometrics, a tool for the quantitative description and statistical analysis of morphology, could allow to model phenotypic changes between traits, emphasizing those trait-trait correlations that have strong, underlying genetic component, and decrease bias in certain direction of the morphospace (Polly 2008).

Climate change studies rely on models that attempt to translate meteorological data into a plausible biological response. Recent progress in modeling tree mortality under drought has led to the incorporation of plant hydraulic traits in the parameterization of simulation models in homogeneous forest communities (Choat et al. 2018, Martínez-Vilalta et al. 2009). The coinheritance of these traits with other morphological, physiological and life-history traits would add value to statements of climate change on trait diversity in natural habitats.

## Conclusions

The decomposition of the phenotypic correlation into coheritability and coenvironmentability is consistent with classical statistical genetics theory and therefore placed on a firm statistical footing. The decomposition provides interpretable results that employs the totality of the data (phenotypic, genetic, environmental), utilizes univariate (heritabilities) and bivariate (correlations) statistical measures, accounts for all combinations of magnitude and direction, standardizes covariance terms to a common denominator, all integrated into a simple, intuitive yet robust framework to simultaneously inform on portions of *r*_*P*_ contributed to joint genetic effects and joint environmental deviations. The graphical representations of *h*_*x,y*_, *e*_*x,y*_, and *r*_*P*_ in the 3DHER-plane or 2DHER-field help visualize all pertinent variables concurrently.

Improving our knowledge of underlying genetic basis of trait-trait associations helps to understand and predict species responses in a more variable environment. Although the coheritability and coenvironmentability, in a biometric sense, *per se* cannot capture subtleties of the causal factors underlying the genotypes or even their complete functional relationships between them, it nevertheless explore an important inferential space that point to relationships that deserve further scrutiny, refine searches, and in general, to supplement results obtained from other experimental lines of evidence.

## Supporting information

Coheritability and Coenvironmentability as Concepts for Partitioning the Phenotypic Correlation

