## Supplementary material for "Coheritability and Coenvironmentability as Concepts for Partitioning the Phenotypic Correlation": Coheritability and Coenvironmentability as Concepts for Partitioning the Phenotypic Correlation

Jorge Vasquez-Kool

4.1.1 Distribution of $r$ when $\rho$ is zero ... 37
4.1.2 Distribution of $r$ when $\rho$ is nonzero ... 38
4.2 Comments on the distribution of $r$ ... 50
4.2.1 General comments ... 50
4.2.2 Comments on the distribution of $r$ when the correlation coefficient parameter $\rho$ is zero ... 53
4.2.3 Comments on the distribution of $r$ when the correlation coefficient parameter $\rho$ is nonzero ... 54
4.3 Moments of the sample correlation coefficient distribution ... 56
4.4 The Fisher $r$-to-$Z$ transform of the correlation coefficient ... 57
5 Inferential statistics ... 60
5.1 Statistical justification ... 60
5.2 Hypothesis testing when null hypothesis sets the parameters to a zero value ... 62
5.3 Hypothesis testing for a parameter equal to a nonzero value ... 68
5.4 Power of the hypothesis test ... 74
5.5 Confidence interval of $rPxy$, $hxy$ and $exy$ ... 78
5.6 Sample size ... 83
6 Visualizing the $hxy \cdot exy \cdot rPxy$-relationship ... 87
6.1 The three-dimensional $h_{x,y} \cdot e_{x,y} \cdot r_{P_{x,y}}$ -plane (3DHER-plane) ... 87
6.2 The two-dimensional $h_{xy} \cdot e_{xy} \cdot r_{P_{xy}}$ -field (2DHER-field) ... 89
6.3 The $hxy \cdot exy \cdot rPxy$-plane in a three-dimensional spherical space ... 91
6.4 The determinant of the $hxy \cdot exy \cdot rPxy$-plane ... 95
7 Set of values and limiting cases ... 96
7.1 The range of values of the coheritability for a given value of the coenvironmentability ... 96
7.2 The set of values of coheritability and coenvironmentability for a given phenotypic correlation . 97
7.3 The set of possible values of heritabilities, and correlations (genetic, environmental) for a given (coheritability, coenvironmentability) datum. ... 99
7.4 The relationship of the heritabilities to yield a given geometric mean ... 102
7.5 Limiting cases ... 103
7.6 The assumption that $rPx,y = rAx,y$ ... 107
7.7 Disparity between the phenotypic and genetic correlations ... 108
7.7.1 The domain of the Disparity Index ... 108
7.7.2 Distribution of the Disparity Index ... 112

#### List of Symbols

(■ denotes subscript to indicate phenotypic P, (additive) genetic A, or environmental E)

| | |
|---|---|
| $P$ | phenotypic value of a trait |
| $A$ | breeding value of a trait |
| $E$ | environmental (residual) effect of a trait |
| $\rho_{■x,y}$ | correlation between traits $x$ and $y$ (population parameter) |
| $H_{x,y}$ | coheritability between traits $x$ and $y$ (population parameter) |
| $E_{x,y}$ | coenvironmentability between traits $x$ and $y$ (population parameter) |
| $r_{■x,y}$ | sample correlation between trait $x$ and trait $y$ |
| $h_x^2$ | narrow-sense heritability of trait $x$ (sample estimator) |
| $e_x^2$ | environmentability of trait $x$ $(1 - h_x^2)$ (sample estimator) |
| $h_{x,y}$ | coheritability of trait $x$ and trait $y$ (sample estimator) |
| $e_{x,y}$ | coenvironmentability of trait $x$ and trait $y$ (broad-sense) (sample estimator) |
| $\varepsilon_{x,y}$ | coenvironmentability of trait $x$ and trait $y$ (narrow-sense) (sample estimator) |
| $\epsilon$ | error term of regression model |
| $C_{■x,y}$ | covariance between trait $x$ and trait $y$ (population parameter) |
| $s_{■x,y}$ | covariance between trait $x$ and trait $y$ (sample estimator) |
| $V_{■x}$ | variance of trait $x$ (population parameter) |
| $s_{■x}^2$, $\sigma_{■x}^2$ | sample variance of trait $x$ |
| $\boldsymbol{x}$, $\boldsymbol{y}$ | character, trait, phenotype |

### 1 Characteristics of the population

#### 1.1 The base population

Consider a population consisting of a large number of individuals that have a known genetic structure. The phenotype of the individuals can be specified by a number of observable, measurable, variable phenotypic traits or characters, which are subject to genetic and environmental influences. The population is therefore assumed to be an infinite, multivariate population.

#### 1.2 Population parameters

The traits exhibit phenotypic variability. There is a fraction of the trait's phenotypic variability that can be attributed to all genetic contributions (additive, dominant, epistatic, maternal, paternal effects), and is measured by the population-level broad sense heritability. In addition, the additive component of the genetic variance is due to the average effects (additive effects) of the alleles. Since each parent transmits a single allele per locus to each offspring, the phenotypic resemblance among relatives is subject to the average effect of the single alleles. In this context, the additive genetic fraction of the phenotypic variance of the trait is the population-level narrow-sense-heritability $(\boldsymbol{h^2})$.

The strength of a linear association between two phenotypic characters measured in the individuals of the population can be quantified by the phenotypic correlation between traits $x$ and $y$ $(\rho_{P_{x,y}})$. The phenotypic correlation between two traits in the population is modelled as the sum of the underlying population-level coheritability $(H_{xy})$ and coenvironmentability $(E_{xy})$ coefficients.

From a bivariate normal distribution with standard deviations $\sigma_x = \sqrt{V_{P_x}}$ , $\sigma_y = \sqrt{V_{P_y}}$ , and the correlation coefficient $\rho_{P_{x,y}}$ , which can be expressed as

$$\rho_{P_{x,y}} \equiv \frac{C_{P_{x,y}}}{\sqrt{V_{P_x} \cdot V_{P_y}}} = \frac{C_{A\,x,y}}{\sqrt{V_{P_x} \cdot V_{P_y}}} + \frac{C_{E\,x,y}}{\sqrt{V_{P_x} \cdot V_{P_y}}}$$

Which is the sum of the population-level coheritability $H_{x,y}$,

$$H_{xy} = \frac{C_{A\,x,y}}{\sqrt{V_{P_x} \cdot V_{P_y}}}$$

And population-level coenvironmentability $E_{x,y}$

$$E_{x,y} = \frac{C_{E\,x,y}}{\sqrt{V_{P_x} \cdot V_{P_y}}}$$

We further assume that some of these traits, say a pair of $x$ and $y$, are under genetic control of alleles (in multiple loci) that affect (influence) both of them in varying (direct positive, negative, synergistic, antagonistic) degrees, causing a stable population-level genetic correlation between the traits $x$ and $y$ ( $\rho_{A_{x,y}}$ ).

$$\rho_A \equiv \frac{C_{A_{x,y}}}{\sqrt{V_{A_x} \cdot V_{A_y}}}$$

Here we are cognizant that the sign of the phenotypic and genetic correlation may not necessarily be the same as the sign of the genetic correlation. The population-level environmental correlation( $\rho_{E_{x,y}}$ ) is

$$\rho_E \equiv \frac{C_{E_{x,y}}}{\sqrt{V_{E_x} \cdot V_{E_{x,y}}}}$$

#### Phenotypic correlation, coheritability and coenvironmentability are population-based and population-dependent parameters.

### 2 Characteristics of the sample

Population parameters can only be calculated from the omniscient set of all observations of a population. However, estimator equations are used to estimate population parameters based on a collection of observations measured from a finite set of individuals drawn without observational bias from the whole population.

We draw a random sample of $n$ individuals from a population. Phenotypic characters are measured on these sampled individuals. Looking at two characters at the time, say $x$ and $y$, we consider a random bivariate sample $(x_i, y_i)$, $i = 1, \dots, n$ taken from a continuous bivariate population, whose joint distribution is denoted as a bivariate normal $f_{X,Y}(x, y)$, and the marginal distribution of the traits are $X \sim N(\mu_X, \sigma_X^2)$ and $Y \sim N(\mu_Y, \sigma_Y^2)$.

The purpose of this section is to briefly decompose the phenotypic statistics into its observational components of variance, namely the additive genetic, environmental and genetic x environment interaction terms.

#### 2.1 The phenotypic value

The phenotype is constituted by characteristics (e.g., characters, features, traits) observed in an organism. It is considered to be the realization of influences exerted by genotypic ($G$) and environmental ($E$) factors. The phenotypic value P is a random variable obtained by measuring an individual's metric character, and can be represented as the sum of the genotypic value $G$, and all environmental values $E$

$$P = G + E$$

The genotypic value can be further subdivided into a sum of random variable, namely additive genetic component (or the breeding value) $A$; and a non-additive ($NA$) component made of dominance deviations $D$ (interaction of alleles in a locus), and epistatic effects $I$ (interaction between alleles of different loci).

$$P = A + D + (AA + AD + DA + DD + AAA + \cdots) + (E_g + E_s) + (A \cdot E)$$

general specific

$$P = A + D + I + \epsilon$$

dominance genetic; epistatic interactions

$$P = A + NA + \epsilon$$

non-additive genetic

$$P = G + \epsilon$$

genetic; environmental

$$P = A + \boldsymbol{E}$$

additive genetic; environmental and non-additive genetic

Note: In regard to the classical partitioning of the genetic variance ($V_G$) of quantitative traits in terms of the additive ($V_A$), dominance ($V_D$), and interlocus epistatic interactions ($V_I$), sometimes, erroneously, gives an appearance that a relationship exist with the homonymous mode of gene action (i.e. additive gene action, dominant gene action, or epistatic gene action). This classical $V_G = V_A + V_D + V_I$ decomposition and the modes of gene action of genes do not possess a oneto-one relationship to each other, and the partition cannot be used to support any specific underlying genetic architecture (Huang and Mackay 2016).

2-1a The phenotypic value

$$P = A + E$$

2-1b The phenotypic mean

$$E(P_x) = \bar{P}_x = \frac{\sum_{i=1}^{n} P_{x_i}}{n}$$

2-1c Decomposition of the phenotypic mean

$$E(P_x) = \bar{P}_x = E(A_x + E_x) = \frac{\sum_{i=1}^{n}(A_{x\,i} + E_{x\,i})}{n} = \frac{\sum_{i=1}^{n} A_{x\,i}}{n} + \frac{\sum_{i=1}^{n} E_{x\,i}}{n} = \bar{A}_x + \bar{E}_x$$

#### 2.2 The sample phenotypic variance

The sample phenotypic variance of a specified trait $x$ is $s^2_{P_x}$, which becomes an estimator of the population parameter $V_{P_x}$. The variance is defined as the expected value of the square of deviations from the mean.

$$V_{P_x} = Var(P_x) = [\,E(P_x - E(P_x)\,]^2 = E\left(P_x^2\right) - [\,E(P_x)\,]^2$$

It is generally biased due to the finite sample count and when the population mean is unknown (which for this reason it is estimated by the sample mean). This introduces an underestimation bias. The Bessel's correction is used here by placing in the denominator $(n-1)$ rathern than $n$. The sample phenotypic variance

$$s^2_{P_x} = \frac{1}{n-1}\sum_{i=1}^{n}\left(P_{x_i} - \bar{P}_x\right)^2$$

$$= \frac{1}{n-1}\sum_{i=1}^{n}\left(P_{x_i}^2 - 2\bar{P}_x P_{x_i} + \bar{P}_x{}^2\right)$$

$$= \frac{1}{n-1}\sum_{i=1}^{n} P_{x_i}^2 - 2\frac{1}{n-1}\bar{P}_x\sum_{i=1}^{n} P_{x_i} + \frac{n}{n-1}\bar{P}_x^2$$

$$= \frac{1}{n-1}\sum_{i=1}^{n} P_{x_i}^2 - 2\frac{n}{n-1}\bar{P}_x\frac{\sum_{i=1}^{n} P_{x_i}}{n} + \frac{n}{n-1}\bar{P}_x^2$$

$$= \frac{1}{n-1}\sum_{i=1}^{n} P_{x_i}^2 - 2\frac{n}{n-1}\bar{P}_x{}^2 + \frac{n}{n-1}\bar{P}_x^2$$

$$= \frac{1}{n-1}\sum_{i=1}^{n} P_{x_i}^2 - \frac{n}{n-1}\bar{P}_x{}^2$$

$$= \frac{1}{n-1}\sum_{i=1}^{n}(A_x + E_x)^2 - \frac{n}{n-1}\left(\bar{A}_x + \bar{E}_x\right)^2$$

$$s^2_{P_x} = \frac{1}{n-1}\sum_{i=1}^{n} A_{x_i}^2 + \frac{2\sum_{i=1}^{n} A_{x_i}E_{x_i}}{n-1} + \frac{1}{n-1}\sum_{i=1}^{n} E_{x_i}^2 - \frac{n}{n-1}\left(\bar{A}_x{}^2 + 2A_xE_x + \bar{E}_x{}^2\right)$$

After some rearrangement

$$s^2_{P_x} = \left[\frac{1}{n-1}\sum_{i=1}^{n} A_{x_i}^2 - \frac{n\bar{A}_x^2}{(n-1)}\right] + \left[\frac{1}{n-1}\sum_{i=1}^{n} E_{x_i}^2 - \frac{n\,\bar{E}_x^2}{(n-1)}\right] + \left[\frac{2\sum_{i=1}^{n} A_{x_i}E_{x_i}}{n-1} - \frac{2\,n\,\bar{A}_x\bar{E}_x}{(n-1)}\right]$$

The sample phenotypic variance, therefore, is partitioned as follows:

**2-2** The sample phenotypic variance

$$s_{P_x}^2 = \frac{1}{n-1}\left[\sum_{i=1}^{n} \bar{A}_x^2 - n\,\bar{A}_x^2\right] + \frac{1}{n-1}\left[\sum_{i=1}^{n} \bar{E}_x^2 - n\,\bar{E}_x^2\right] + \frac{2}{n-1}\left[\sum\nolimits_{i=1}^{n} A_{x_i}E_{x_i} - n\bar{A}_x\,\bar{E}_x\right]$$

$$s_{P_x}^2 = s_{A_x}^2 + s_{E_x}^2 + 2\,s_{AE_x}^2$$

- $s_{A_x}^2$: sample additive genetic variance = sample variance of breeding values
- $s_{E_x}^2$: sample environmental variance
- $2\,s_{AE_x}^2$: sample variance of the interaction between breeding values and environmental values

#### 2.3 The sample phenotypic covariance

The sample phenotypic covariance between the phenotypic value of trait $x$ and the phenotypic value of trait $y$ , $s_{P_{x,y}}$, is the estimator of the population parameter, $C_{P_{x,y}}$ . The covariance is a measure of joint variability of two random variables, which are the phenotypic values for trait $x$, $P_x$ and trait $y$, $P_y$. The covariance is defined as the expected product of the deviations of the phenotypic values from their corresponding expected values.

$$C_{P_{x,y}} = Cov(P_x, P_y) = E[P_x - E(P_x)][P_y - E(P_y)] = [E(P_x P_y) - E(P_x)E(P_y)]$$

A sample of size $n$ is drawn from a population of the random variable pair $(P_x, P_y)$ such each observation $(P_{x_i}, P_{y_i})$ for $i = 1, \dots, n$, is drawn with equal probability $n^{-1}$ , and employing the Bessel's correction, sample phenotypic covariance is:

$$\begin{aligned}
s_{P_{x,y}} &= \frac{1}{n-1} \sum_{i=1}^{n} (P_{x_i} - \bar{P}_x)(P_{y_i} - \bar{P}_y) \\
&= \frac{1}{n-1} \sum_{i=1}^{n} (P_{x_i} P_{y_i} - P_{x_i} \bar{P}_y - \bar{P}_x P_{y_i} + \bar{P}_x \bar{P}_y) \\
&= \frac{1}{n-1} \sum_{i=1}^{n} (P_{x_i} P_{y_i} - P_{x_i} \bar{P}_y - \bar{P}_x P_{y_i} + \bar{P}_x \bar{P}_y) \\
&= \frac{1}{n-1} \left[ \sum_{i=1}^{n} P_{x_i} P_{y_i} - \sum_{i=1}^{n} P_{x_i} \bar{P}_y - \sum_{i=1}^{n} \bar{P}_x P_{y_i} + \sum_{i=1}^{n} \bar{P}_x \bar{P}_y \right] \\
&= \frac{1}{n-1} \left[ \sum_{i=1}^{n} P_{x_i} P_{y_i} - \frac{n}{n} \sum_{i=1}^{n} P_{x_i} \bar{P}_y - \frac{n}{n} \sum_{i=1}^{n} \bar{P}_x P_{y_i} + n\bar{P}_x \bar{P}_y \right] \\
&= \frac{1}{n-1} \left[ \sum_{i=1}^{n} P_{x_i} P_{y_i} - n\bar{P}_x \bar{P}_y - n\bar{P}_x \bar{P}_y + n\bar{P}_x \bar{P}_y \right]
\end{aligned}$$

**2-3** The sample phenotypic covariance

$$s_{P_{x,y}} = \frac{1}{n-1} \left[ \sum_{i=1}^{n} P_{x_i} P_{y_i} - n\bar{P}_x \bar{P}_y \right]$$

#### 2.4 Decomposition of the sample phenotypic covariance

The sample phenotypic covariance $C_{P_{xy}}$ between traits $x$ and $y$ can be thought to be the sum of an additive genetic covariance $C_{A_{xy}}$, an environmental, residual covariance term, $C_{E_{xy}}$, and interaction terms $C_{A_xE_y}$ and $C_{A_yE_x}$. Thus, $C_{P_{xy}} = C_{A_{xy}} + C_{E_{xy}}$. The genetic covariance $C_{A_{xy}}$ of two traits is a component of the sample phenotypic covariance that represents the joint measure of genetic association between two traits.

A quantitative genetic model of the value $P$ of trait is assumed to be the sum of an additive genetic value $A$ and an environmental, residual value $E$, thus $P = A + E$.

$$s_{P_{x,y}} = \frac{1}{n-1}\left[\sum_{i=1}^{n} P_{x_i}P_{y_i} - n\bar{P}_x\bar{P}_y\right]$$

Replacing the corresponding $P = A + E$ for each trait, we obtain

$$s_{P_{x,y}} = \frac{1}{n-1}\left[\sum_{i=1}^{n}(A_{x_i}+E_{x_i})(A_{y_i}+E_{y_i}) - n\frac{(\bar{A}_x+\bar{E}_x)}{n}\frac{(\bar{A}_y+\bar{E}_y)}{n}\right]$$

$$= \frac{1}{n-1}\left[\sum_{i=1}^{n}(A_{x_i}A_{y_i} + A_{x_i}E_{y_i} + A_{y_i}E_{x_i} + E_{x_i}E_{y_i}) - \frac{(\bar{A}_x+\bar{E}_x)(\bar{A}_y+\bar{E}_y)}{n}\right]$$

$$= \frac{1}{n-1}\left[\sum_{i=1}^{n}A_{x_i}A_{y_i} + \sum_{i=1}^{n}E_{x_i}E_{y_i} + \left(\sum_{i=1}^{n}A_{x_i}E_{y_i} + \sum_{i=1}^{n}A_{y_i}E_{x_i}\right) - n\bar{A}_{x_i}\bar{A}_{y_i} - n\bar{A}_x\bar{E}_y - n\bar{A}_y\bar{E}_x - n\bar{E}_x\bar{E}_y\right]$$

$$s_{P_{x,y}} = \frac{1}{n-1}\left[\sum_{i=1}^{n}(A_{x_i}A_{y_i} - n\bar{A}_x\bar{A}_y) + \sum_{i=1}^{n}(E_{x_i}E_{y_i} - n\bar{E}_x\bar{E}_y) + \left(\sum_{i=1}^{n}(A_{x_i}E_{y_i} - n\bar{A}_x\bar{E}_y) + \sum_{i=1}^{n}(A_{y_i}E_{x_i} - n\bar{A}_y\bar{E}_x)\right)\right]$$

**2-4** Decomposition of the sample phenotypic covariance between traits $x$ and $y$.

| Term | Symbol | Description | Group |
|---|---|---|---|
| $\frac{1}{n-1}\left[\sum_{i=1}^{n} P_{x_i}P_{y_i} - n\bar{P}_x\bar{P}_y\right]$ | $s_{P_{xy}}$ | phenotypic covariance | |
| $= \frac{1}{n-1}\sum_{i=1}^{n}(A_{x_i}A_{y_i} - n\bar{A}_x\bar{A}_y)$ | $s_{A_{xy}}$ | sample genetic covariance between the breeding values of traits $x$ and $y$ | $s_{A_{x,y}}$ additive genetic covariance |
| $+ \frac{1}{n-1}\sum_{i=1}^{n}(E_{x_i}E_{y_i} - n\bar{E}_x\bar{E}_y)$ | $s_{\epsilon_{xy}}$ | sample covariance between environmental deviations of traits $x$ and $y$ | $s_{E_{x,y}}$ environmental covariance |
| $+ \frac{1}{n-1}\sum_{i=1}^{n}\left(A_{x_i}E_{y_i} - n\bar{A}_x\bar{E}_y\right)$ | $s_{A_xE_y}$ | sample covariance of the GxE interaction of breeding values of trait $x$ and the environmental deviations of trait $y$ | |
| $+ \frac{1}{n-1}\sum_{i=1}^{n}\left(A_{y_i}E_{x_i} - n\bar{A}_y\bar{E}_x\right)$ | $s_{A_yE_x}$ | sample covariance of the GxE interaction of breeding values of trait $y$ and the environmental deviations of trait $x$ | |

#### 2.5 The sample phenotypic correlation

The sample phenotypic correlation between two traits $x$ and , $r_{P_{x,y}}$ , is the estimator of the population parameter $\rho_{x,y}$ also referred as the Pearson product-moment correlation coefficient, or Pearson's correlation coefficient, or simply "the correlation coefficient". It is obtained by dividing the covariance of the two variables by the product of their standard deviations.

The population correlation coefficient $\rho_{P_{x,y}}$ between two random variables, namely trait $x$ and trait $y$ with expected values $E(P_x)$ and $E(P_x)$ and variances $V_{P_x}$ and $V_{P_y}$ is defined as

$$\rho_{P_{x,y}} = Corr(P_x, P_y) = \frac{C_{P_{x,y}}}{\sqrt{V_{P_x} V_{P_y}}} = \frac{E[P_x - E(P_x)][P_y - E(P_y)]}{\sqrt{V_{P_x} V_{P_y}}}$$

for $-1 < \rho_{P_{x,y}} < 1$

If $\rho_{P_{x,y}} = \pm 1$, then the phenotypic characters $P_x$ and $P_y$ are linearly related.

If the phenotypic characters $P_x$ and $P_y$ are linearly related, then $\rho_{P_{x,y}} = \pm 1$.

Suppose $(P_x, P_y), (P_{x_i}, P_{y_i}), i = 1, \dots, n$ constitute a sample of $n$ independent and identically distributed (i.i.d) random vectors drawn from a bivariate normal distribution of phenotypic values of trait $x$ $(P_x)$ and trait $y$ $(P_y)$ whose population-level phenotypic correlation parameter is $\rho_{P_{x,y}}$. The trait means are $\bar{P}_x$ and $\bar{P}_y$, variances $s^2_{P_x}$ and $s^2_{P_y}$. The sample phenotypic correlation $r_{P_{x,y}}$ of traits $x$ and $y$ is defined as the sample phenotypic covariance $s_{P\,x,y}$ divided by the product of their standard deviations $s_{P_x} s_{P_y} = \sqrt{s^2_{P_x}\, s^2_{P_y}}$.

**2-5** The sample phenotypic correlation

$$r_{P_{x,y}} = \frac{s_{P\,x,y}}{\sqrt{s^2_{P_x} s^2_{P_y}}} = \frac{\frac{1}{n-1}\left[\sum_{i=1}^{n} P_{x_i} P_{y_i} - n\bar{P}_x \bar{P}_y\right]}{\sqrt{\frac{1}{n-1}\left[\sum_{i=1}^{n} P_{x_i} - \bar{P}_x\right]^2}\sqrt{\frac{1}{n-1}\left[\sum_{i=1}^{n} P_{y_i} - \bar{P}_y\right]^2}}$$

See section 3.2.1 for a path analytical treatment of the phenotypic correlation.

A discussion about the Fisher’s and Pearson’s formulations, and properties associated to them is elaborated by Plata (2006).

#### 2.6 Decomposition of the sample phenotypic correlation

The sample phenotypic correlation can be decomposed in an analogous as the sample phenotypic covariance.

$$r_{P_{x,y}} = \frac{\frac{1}{n-1}\sum_{i=1}^{n}(A_{x_i}A_{y_i} - n\bar{A}_x\bar{A}_y) + \frac{1}{n-1}\sum_{i=1}^{n}(E_{x_i}E_{y_i} - n\bar{E}_x\bar{E}_y) + \frac{1}{n-1}\sum_{i=1}^{n}\left(A_{x_i}E_{y_i} - n\bar{A}_x\bar{E}_y\right) + \frac{1}{n-1}\sum_{i=1}^{n}\left(A_{y_i}E_{x_i} - n\bar{A}_y\bar{E}_x\right)}{\sqrt{\frac{1}{n-1}\left[\sum_{i=1}^{n} P_{x_i} - \bar{P}_x\right]^2}\sqrt{\frac{1}{n-1}\left[\sum_{i=1}^{n} P_{y_i} - \bar{P}_y\right]^2}}$$

**2-6** Components of the sample phenotypic correlation

$r_{P_{x,y}} =$ Sample phenotypic correlation

$$\frac{\frac{1}{n-1}\sum_{i=1}^{n}(A_{x_i}A_{y_i} - n\bar{A}_x\bar{A}_y)}{\sqrt{\frac{1}{n-1}\left[\sum_{i=1}^{n} P_{x_i}{}^2 - \bar{P}_x^2\right]}\sqrt{\frac{1}{n-1}\left[\sum_{i=1}^{n} P_{y_i}{}^2 - \bar{P}_y^2\right]}}$$

Sample coheritability

$$h_{x,y} = \begin{cases} \dfrac{s_{A_{x,y}}}{\sqrt{s_{P_x}^2 s_{P_x}^2}} \\ \sqrt{h_y^2 h_x^2}\, r_{A_{x,y}} \end{cases}$$

$$+ \frac{\frac{1}{n-1}\sum_{i=1}^{n}(E_{x_i}E_{y_i} - n\bar{E}_x\bar{E}_y)}{\sqrt{\frac{1}{n-1}\left[\sum_{i=1}^{n} P_{x_i}{}^2 - \bar{P}_x^2\right]}\sqrt{\frac{1}{n-1}\left[\sum_{i=1}^{n} P_{y_i}{}^2 - \bar{P}_y^2\right]}}$$

Sample (narrow-sense) coenvironmentability

$$\varepsilon_{x,y} = \begin{cases} \dfrac{s_{E_{x,y}}}{\sqrt{s_{P_x}^2 s_{P_x}^2}} \\ \sqrt{(1-h_x^2)(1-h_y^2)}\, r_{E_{x,y}} \end{cases}$$

$$+ \frac{\frac{1}{n-1}\sum_{i=1}^{n}\left(A_{x_i}E_{y_i} - n\bar{A}_x\bar{E}_y\right)}{\sqrt{\frac{1}{n-1}\left[\sum_{i=1}^{n} P_{x_i}{}^2 - \bar{P}_x^2\right]}\sqrt{\frac{1}{n-1}\left[\sum_{i=1}^{n} P_{y_i}{}^2 - \bar{P}_y^2\right]}}$$

genotype x environment interaction component

$$a_x e_y = \begin{cases} \dfrac{s_{A_x E_y}}{\sqrt{s_{P_x}^2 s_{P_x}^2}} \\ \sqrt{h_x^2(1-h_y^2)}\, r_{A_x E_y} \end{cases}$$

$$+ \frac{\frac{1}{n-1}\sum_{i=1}^{n}\left(A_{y_i}E_{x_i} - n\bar{A}_y\bar{E}_x\right)}{\sqrt{\frac{1}{n-1}\left[\sum_{i=1}^{n} P_{x_i}{}^2 - \bar{P}_x^2\right]}\sqrt{\frac{1}{n-1}\left[\sum_{i=1}^{n} P_{y_i}{}^2 - \bar{P}_y^2\right]}}$$

genotype x environment interaction component

$$a_y e_x = \begin{cases} \dfrac{s_{A_y E_x}}{\sqrt{s_{P_x}^2 s_{P_x}^2}} \\ \sqrt{(1-h_x^2)h_y^2}\, r_{A_y E_x} \end{cases}$$

$e_{xy}$ coenvironmentability (broad-sense)

#### 2.7 The sample coheritability

The sample coheritability $h_{x,y}$ between two traits $x$ and $y$ is the genetic component of the sample phenotypic correlation $r_{P_{x,y}}$ , and is an estimator of the population coheritability ( $H_{x,y}$ ) which is defined as the ratio of the additive genetic covariance between the characters x and y divided by the geometric mean of the phenotypic variances of the traits.

The population coheritability is defined as

$$H_{x,y} = \frac{C_{A_{x,y}}}{\sqrt{V_{P_x} V_{P_y}}}$$

The sample coheritability is defined as the sample genetic covariance divided by the geometric mean of the sample phenotypic variances.

**2-7** sample coheritability

$$h_{x,y} = \frac{s_{A_{x,y}}}{\sqrt{s_{P_x}^2 s_{P_y}^2}} = \frac{\frac{1}{n-1}\left[\sum_{i=1}^{n} Ax_i Ay_i - \frac{\sum_{i=1}^{n} Ax_i \cdot \sum_{i=1}^{n} Ay_i}{n}\right]}{\sqrt{\frac{1}{n-1}\left[\sum_{i=1}^{n} P_{x_i} - \bar{P}_x\right]^2}\sqrt{\frac{1}{n-1}\left[\sum_{i=1}^{n} P_{y_i} - \bar{P}_y\right]^2}}$$

For $-1 < h_{x,y} < 1$

The following statements provide a notion on the meaning of the sample coheritability.

- The coheritability is the fraction of the phenotypic correlation of two traits that can be attributed to additive genetic effects.
- The coheritability is the proportion of the total phenotypic variability of two traits that is due to joint/common/shared genetic causes.
- The coheritability is the relative contribution of shared genetics in determining the phenotypic association of two traits.
- The coheritability measures the extent to which resemblance among relatives in regard to two traits in comparison to unrelated individuals of the same species, is due to common heredity.

#### 2.8 The sample coenvironmentability

The sample coenvironmentability $e_{x,y}$ between traits $x$ and $y$ is the residual, environmental component of the sample phenotypic correlation $r_{P_{x,y}}$ , and is an estimator of the population coheritability ( $E_{x,y}$ ).

The population coenvironmentability is defined as

$$E_{x,y} = \frac{C_{E\,x,y}}{\sqrt{V_{P_x}\,V_{P_y}}}$$

It is generally assumed that the genetic and environmental interaction effects between the breeding values of one trait and the environmental effects of another are negligible. These effects, if not explicitly formulated, will be added with the environmental covariance. Therefore, we can partition the numerator into a genetic covariance and a residual environmental covariance component, as follows:

**The narrow-sense coenvironmentability** is a measure of the shared effects of environmental deviations acting upon both traits.

$$\varepsilon_{x,y} = \sqrt{(1-h_x^2)(1-h_y^2)}\; r_{E_{X,y}}$$

**2-8a** The sample narrow-sense sample coenvironmentability

$$\varepsilon_{x,y} = \frac{s_{E\,x,y}}{\sqrt{s_{P_x}^2\,s_{P_x}^2}} = \frac{\frac{1}{n-1}\sum_{i=1}^{n}(E_{x_i}E_{y_i} - n\bar{E}_x\bar{E}_y)}{\sqrt{\frac{1}{n-1}\left[\sum_{i=1}^{n} P_{x_i} - \bar{P}_x\right]^2}\sqrt{\frac{1}{n-1}\left[\sum_{i=1}^{n} P_{y_i} - \bar{P}_y\right]^2}}$$

**The broad-sense coenvironmentability** is the combined effect of the narrow-sense coenvironmentability plus the interaction effects of the breeding values of a trait with the environmental effect of the other. Broad-sense coenvironmentability measures the degree to which the joint phenotypic variability of two traits is determined by all sources of variation excluding the additive genetic variation.

$$e_{x,y} = \sqrt{(1-h_x^2)(1-h_y^2)}\, r_{E_{X,y}} + \sqrt{h_x^2(1-h_y^2)}\, r_{A_x E_y} + \sqrt{(1-h_x^2)h_y^2}\, r_{A_y E_x}$$

**2-8b** The sample broad-sense sample coenvironmentability

$$e_{x,y} =$$

$$\frac{\frac{1}{n-1}\sum_{i=1}^{n}(E_{x_i}E_{y_i} - n\bar{E}_x\bar{E}_y)}{\sqrt{\frac{1}{n-1}\left[\sum_{i=1}^{n} P_{x_i} - \bar{P}_x\right]^2}\sqrt{\frac{1}{n-1}\left[\sum_{i=1}^{n} P_{y_i} - \bar{P}_y\right]^2}}$$

sample broad-sense coenvironmentability $e_{x,y}$

sample coenvironmentability (narrow-sense)

$$\varepsilon_{x,y} = \sqrt{(1-h_x^2)(1-h_y^2)}\, r_{E_{X,y}}$$

$$+ \frac{\frac{1}{n-1}\sum_{i=1}^{n}\left(A_{x_i}E_{y_i} - n\bar{A}_x\bar{E}_y\right)}{\sqrt{\frac{1}{n-1}\left[\sum_{i=1}^{n} P_{x_i} - \bar{P}_x\right]^2}\sqrt{\frac{1}{n-1}\left[\sum_{i=1}^{n} P_{y_i} - \bar{P}_y\right]^2}}$$

genotype x environment interaction component $A_xE_y$

$$a_x e_y = \sqrt{h_x^2(1-h_y^2)}\, r_{A_x E_y}$$

$$+ \frac{\frac{1}{n-1}\sum_{i=1}^{n}\left(A_{y_i}E_{x_i} - n\bar{A}_y\bar{E}_x\right)}{\sqrt{\frac{1}{n-1}\left[\sum_{i=1}^{n} P_{x_i} - \bar{P}_x\right]^2}\sqrt{\frac{1}{n-1}\left[\sum_{i=1}^{n} P_{y_i} - \bar{P}_y\right]^2}}$$

genotype x environment interaction component $A_yE_x$

$$a_y e_x = \sqrt{(1-h_x^2)h_y^2}\, r_{A_y E_x}$$

The broad-sense coenvironmentability included not only (narrow-sense) coenvironmentability, but in addition the interaction terms between genetic effects of one-trait and the environmental effects of the other. They are often assumed as being non-existent or negligible. Yet, statistically these components, if different than zero, may add “noise” to the estimators. In that case, it is expected that the interaction would inflate the (co-)environmental component. This addition of interaction effects on the coenvironmentability would cause increase in its magnitude and possibly surpassing the magnitude of the coheritability. Partitions (See section 6.2) where this may occur are $S_{+3}$ and $S_{-3}$. Thus though environmental effects affect all phenotypic correlations, they have a preponderant influence over the coheritability in these particular partitions.

### 3 Equivalent formulae of the sample coheritability

#### 3.1 Expressions of the sample coheritability

The value of an individual's phenotype (P) is determined by additive genetic (A) and environmental (E) effects, such that P = A + E. This simple decomposition holds also at the population level for variances ($s^2$), and covariances ($C$) between characters $x$ and $y$. We can therefore apply this decomposition to a bivariate dataset of measurements of two characters from a number of individuals sampled from a population and calculate an additive genetic covariance ($s_{A_{x,y}}$). All the rest constitutes the environmental covariance ($s_{E_{x,y}}$) which includes environmental effects together with non-additive genetic variation,

$$s_{P_{x,y}} = s_{A_{x,y}} + s_{E_{x,y}}$$

To standardize, both sides are divided by the square root of the product of phenotypic variances of each character, and using observational variance components symbols, we obtain

$$\frac{s_{P_{x,y}}}{\sqrt{s_{P_x}^2 \, s_{P_y}^2}} = \frac{s_{A_{x,y}}}{\sqrt{s_{P_x}^2 \, s_{P_y}^2}} + \frac{s_{E_{x,y}}}{\sqrt{s_{P_x}^2 \, s_{P_y}^2}}$$

**Expression I**

The ratio of the sample genetic covariance and the bivariate phenotypic variability (i.e., geometric mean of the sample phenotypic variances)

$$h_{xy} = \frac{s_{A_{x,y}}}{\sqrt{s_{P_x}^2 \, s_{P_y}^2}}$$

for $s_{P_x}^2 > 0$, $s_{P_y}^2 > 0$

Expression II

Rearranging Expression I by multiplying and diving by the geometric mean of the additive genetic variances we obtain:

$$h_{xy} = \frac{\sqrt{s_{A_x}^2 \, s_{A_y}^2}}{\sqrt{s_{A_x}^2 \, s_{A_y}^2}} \frac{s_{A_{x,y}}}{\sqrt{s_{P_x}^2 \, s_{P_y}^2}} = \sqrt{h_x^2 \, h_y^2} \, r_{A_{xy}}$$

for $s_{P_x}^2 > 0$ , $s_{P_y}^2 > 0$

Expression III

Multiplying the numerator and denominator of Expression I by the phenotypic covariance , we obtain:

$$h_{xy} = \frac{s_{A_{x,y}}}{\sqrt{s_{P_x}^2 \, s_{P_y}^2}} = \frac{s_{P_{x,y}}}{s_{P_{x,y}}} \frac{s_{A_{x,y}}}{\sqrt{s_{P_x}^2 \, s_{P_y}^2}} = \frac{s_{A_{x,y}}}{s_{P_{x,y}}} \frac{s_{P_{x,y}}}{\sqrt{s_{P_x}^2 \, s_{P_y}^2}}$$

$$h_{xy} = \frac{s_{A_{x,y}}}{s_{P_{x,y}}} \, r_{P_{x,y}}$$

where $s_{P_{x,y}} \neq 0$.

Notice that only under extreme and rare circumstance when the phenotypic correlation $r_{P_{x,y}} = 1$ will the ratio of genetic and phenotypic covariance become a valid expression of the coheritability, $h_{xy} = \frac{s_{A_{x,y}}}{s_{P_{x,y}}}$, otherwise its range is from $-\infty$ to $+\infty$. (a version of this formulation is presented by Yamada 1968).

Expressions IV and V involve the use of a regression parameter

Expression IV

In the model $P_y = \beta_o + \beta_{P_yP_x} P_x + \epsilon +, \beta_{P_xP_y},$

$\beta_{P_yP_x}$ represents the regression coefficient of independent phenotypic value $P_x$ on dependent phenotypic value $P_y$,

$$\beta_{P_yP_x} = \frac{s_{P_{x,y}}}{s^2_{P_x}}$$

which can be related to the phenotypic correlation using the following relationship

$$r_{P_{x,y}} = \beta_{P_yP_x} \frac{\sqrt{s^2_{P_x}}}{\sqrt{s^2_{P_y}}}$$

Then, replacing this form of the phenotypic correlation in Expression III, the coheritability is

$$h_{x,y} = \frac{s_{A_{x,y}}}{s_{P_{x,y}}} \beta_{P_yP_x} \frac{\sqrt{s^2_{P_x}}}{\sqrt{s^2_{P_y}}}$$

Expression V

In the regression model $P_x = \beta_o + \beta_{P_yP_x} P_y + \epsilon$,

$\beta_{P_xP_y}$ represents the regression coefficient of independent phenotypic value $P_y$ on dependent phenotypic value $P_x$, then

$$r_{P_{x,y}} = \beta_{P_xP_y} \frac{\sqrt{s^2_{P_y}}}{\sqrt{s^2_{P_x}}}$$

Then, replacing this form of the phenotypic correlation in expression III, the coheritability is

$$h_{x,y} = \frac{s_{A_{x,y}}}{s_{P_{x,y}}} \beta_{P_xP_y} \frac{\sqrt{s^2_{P_y}}}{\sqrt{s^2_{P_x}}}$$

**Equivalent Expressions of Coheritability Summary**

$$h_{x,y} = \begin{cases} \dfrac{s_{A_{x,y}}}{\sqrt{s_{P_x}^2 \; s_{P_y}^2}} & s_{P_x}^2 \neq 0 \,,\; s_{P_y}^2 \neq 0 \\ \sqrt{h_y^2 h_x^2}\; r_{A_{x,y}} & \\ \dfrac{s_{A_{x,y}}}{s_{P_{x,y}}}\; r_{P_{x,y}} & C_{P_{x,y}} \neq 0 \\ \dfrac{s_{A_{x,y}}}{s_{P_{x,y}}}\beta_{P_y P_x} \dfrac{\sqrt{s_{P_x}^2}}{\sqrt{s_{P_y}^2}} & s_{P_{x,y}} \neq 0, s_{P_y}^2 \neq 0 \\ \dfrac{s_{A_{x,y}}}{s_{P_{x,y}}}\beta_{P_x P_y} \dfrac{\sqrt{s_{P_y}^2}}{\sqrt{s_{P_x}^2}} & s_{P_x}^2 \neq 0, s_{P_{x,y}} \neq 0 \end{cases}$$

These expressions reduce to the heritability if the traits are the same $x = y$ .

Notice that in all forms, a coheritability value equal to zero implies that the genetic covariance is zero. The additive genetic variances, present in the denominator of the genetic correlation, cannot become zero since it would leave it undefined. Thus, the heritabilities cannot assume the value of zero. This shows that the coheritability estimator may, depending on the numbers involved, sometimes display numerical instability.

At this point the reader may wonder why we bother to spell out several mathematically equivalent formulations in a variety of variables and equations. The reasons for this are that (1) within a quantitative theoretical framework, it shows relationships among a variety of variables, thus providing conceptual insight among them. (2) The different expressions entail different implicit assumptions about what entities are related to each other. Third, some forms are more easily calculated than others, thus creates expressions that function more conceptually and others more operationally.

#### 3.2 Path Analysis

Path analysis is a mathematical method that attempts to model the relationship among correlated variables. The path analysis begins with a path diagram that considers two types of variables. Endogenous variables such as the phenotypic trait ($P$) are measurable and observed. They are represented enclosed in squares. Exogenous variables which genetic ($A$) , as well as the environmental ($E$) variables, cannot be directly observed or measured, and are enclosed in circles.

The path diagram involves arrows, either singled-headed or double-headed. A single-headed arrow indicates simply statistical predictability of a latent variable to the observed variable, with no commitment made about causality. In this sense $\textcircled{A_x} \rightarrow \boxed{P_x}$ denotes that the individual differences in exogenous variable $A$ may influence endogenous variable $P$, even when all other variables that predict $P$ are taken into account. Double-headed arrows indicate a correlation between exogenous variables, e.g. $\textcircled{A_x} \leftrightarrow \textcircled{A_y}$ However, a double-headed arrow can never be used between endogenous variables.

Path coefficients quantify the magnitude of a direct prediction. For example, the term $\sqrt{h_x^2}$ on the single-headed arrow between $\textcircled{A_x}$ and $\boxed{P_x}$ in figure 3.2.1 is a path coefficient that quantifies the additive genetic influences exerted by genotype $A_x$ on the phenotypic trait $P_x$. In figure , the additive genetic correlation measure the strength of a linear association between breeding values of trait x and trait y, thus $\textcircled{A_x} \leftrightarrow \textcircled{A_y}$ possess a quantifiable $r_{A_{xy}}$. The model provides no assumptions as to the source of the association.

##### 3.2.1 The phenotypic correlation

The phenotypic value of a trait is influenced by the (additive) genetic factors and the residual, environmental factors. The phenotypic correlation between traits $x$ and $y$, therefore, is the sum of the coheritability path and the (broad-sense) coenvironmentability path (cf. Sections 2.7 and 2.8)

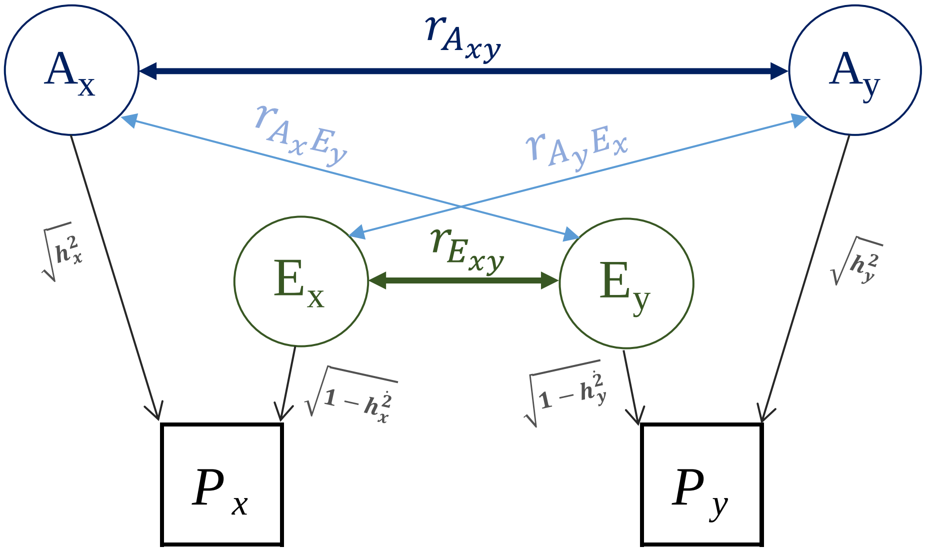

**Figure 3.2.1.** Path diagram relating two exogenous variables, namely, the phenotypic traits ($P_x, P_y$) to their corresponding additive genetic effects ($A_x, A_y$), and environmental influences ($E_x, E_y$), respectively. The path coefficient associated to a single headed arrow connecting a genotype A and its phenotypic trait $P$ is equal to the square root of heritability, $\sqrt{h^2}$ . Double headed arrows represent correlation coefficients. For the sake of completeness, genotype by environment covariance is considered.

Therefore, the phenotypic correlation becomes:

**3-2-1** Decomposition of the sample phenotypic correlation

$$r_{P_{x,y}} = \underbrace{\sqrt{h_x^2 h_y^2}\, r_{A_{x,y}}}_{h_{x,y}} + \underbrace{\sqrt{(1-h_x^2)(1-h_y^2)}\, r_{E_{x,y}}}_{\varepsilon_{x,y}} + \underbrace{\sqrt{h_x^2(1-h_y^2)}\, r_{A_x E_y}}_{a_x e_y} + \underbrace{\sqrt{(1-h_x^2)h_y^2}\, r_{A_y E_x}}_{a_y e_x}$$

$$r_{P_{x,y}} = h_{x,y} + \varepsilon_{x,y} + a_x e_y + a_y e_x$$

The last line invokes equation [ 2-6 ].

##### 3.2.2 Type B coheritability of one trait expressed in different environments

In field trials, breeders are interested in measuring the intensity of the interaction of the environment and the genotype. The correlation between the breeding values of a trait measured in two distinct localities can be considered as two distinct traits (Falconer 1952), whose genetic correlation is denoted as Type B-genetic genetic correlation ($r_{B\ x1,x2}$) (coined by Burdon 1977).

In this sense, environment can represent a locality (site 1, site2,…), sex (male, female), generation (parental, progeny). The same rationale applies if considering distinct ages or ontogenic stages (juvenile, adult), intrinsic conditions (normal, affected), etc.

An important consideration is not only to have $r_{B\ x1,x2}$, but also the phenotypic correlation of the phenotypic values of both characters expressed in the environments. This would allow to estimate the Type B coenvironmentability.

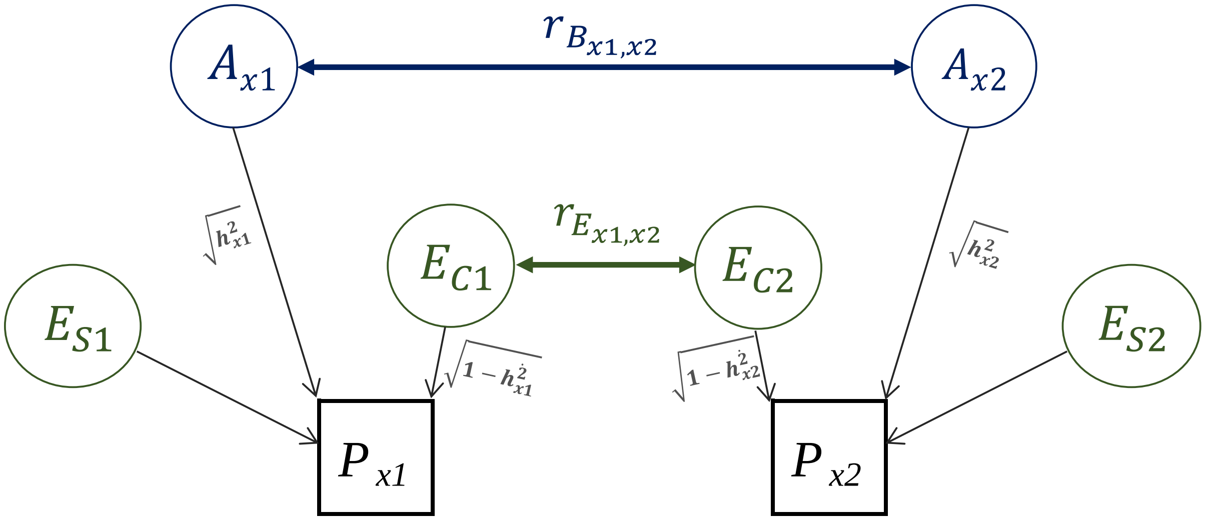

**Figure 3.2.2.** Type B genetic correlation between the breeding values of a trait measured in localities 1 and 2. $A_p$ is denotes by the genetic component determined in environment $p$ ($p = 1,2$), $E_c$ is the common environment, $E_s$ the specific environment. [Adapted from Falconer (1952)].

**3-2-2** Type B coheritability of the same trait x in two environments (named 1 and 2)

$$h_{B_{x1\,,\,x2}} = \sqrt{h^2_{x1} h^2_{x2}\, r_{B_{x1\,,\,x2}}}$$

##### 3.2.3 Multiple regression

The graph below represents a path diagram expressing a multiple linear regression equation with one outcome variable, $r_{P_{x,y}}$, and two predictors, namely the coheritability $h_{x,y}$ and the coenvironmentability $e_{x,y}$. The model is

$$r_{P_{x,y}} = \boldsymbol{\beta_o} + \boldsymbol{\beta_1} \cdot h_{x,y} + \boldsymbol{\beta_2} \cdot e_{x,y} + 1\epsilon$$

where $\epsilon$ is a residual term with a mean of zero. There are four single-headed arrows pointing into the phenotypic correlation $r_{P_{x,y}}$ , whose path coefficients are $\boldsymbol{\beta_o}$, $\boldsymbol{\beta_1}$, $\boldsymbol{\beta_2}$, and **1** corresponding to the four summands on the right hand side of regression equation. In a general linear model, the intercept $\boldsymbol{\beta_o}$ is estimated using a column of ones. For this reason, a constant is denoted by a triangle that maps onto that column of ones.

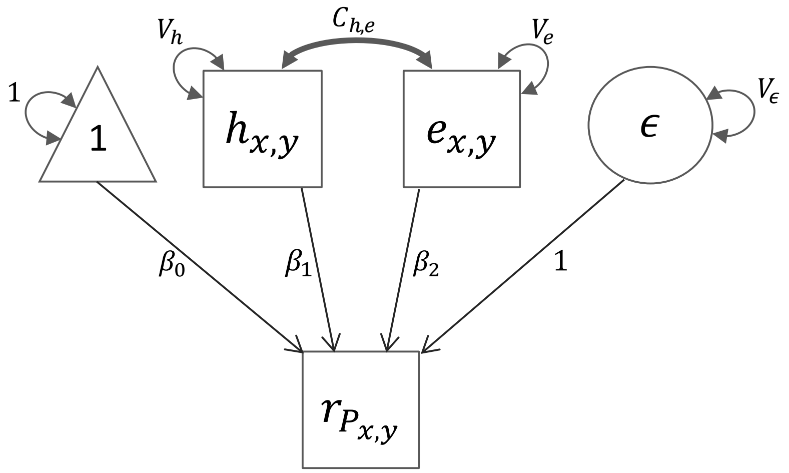

**Figure 3.2.3** Path diagram of a multiple regression where the phenotypic correlation is the dependent variable. The coheritability and coenvironmentability are the independent predictor variables.

For simplicity of presentation in Figure 3.2.3, the two predictor variables $h_{x,y}$ and $e_{x,y}$ have means of zero. If $h_{x,y}$ and $e_{x,y}$ had nonzero then single-headed arrows would be drawn from the triangle to $h_{x,y}$ and from the triangle to $e_{x,y}$ . There are double-headed variance arrows for each of the variables $h_{x,y}$ , $e_{x,y}$ , and $\epsilon$ on the right hand side of the equation, representing the variance of the predictor variables, $V_h$ , $V_e$ and residual variance $V_\epsilon$ respectively. The constant has, by convention, a nonzero variance term fixed at the value 1.0. While this double-headed arrow may seem counterintuitive since it is not formally a variance term, it is required in order to provide consistency to the path tracing rules described below. In addition, there is a double-headed arrow between $h_{x,y}$ and $e_{x,y}$ that specifies the potential covariance, $C_{h_{x,y} \cdot e_{x,y}}$ between the predictor variables.

#### 3.3 Sampling variance approximation

The Delta method involves a first order approximation of a Taylor series expansion of the function and then taking the expectation of according to Taylor's Theorem (see Appendix 6).

##### 3.3.1 Sampling variance of the phenotypic correlation

The sampling variance of the phenotypic correlation in terms of its components coheritability and coenvironmentability can be defined as:

$$Var\left(r_{P_{x,y}}\right) = Var\left(h_{x,y} + e_{x,y}\right) = Var\left(h_{x,y}\right) + Var\left(e_{x,y}\right) + 2Cov\left(h_{x,y}\,, e_{x,y}\right)$$

The last term can be calculated as:

$$Cov\left(h_{x,y}\,, e_{x,y}\right) = \tfrac{1}{2}\left[Var\left(r_{P_{x,y}}\right) - Var\left(h_{x,y}\right) - Var\left(e_{x,y}\right)\right].$$

The sample phenotypic correlation is defined in equation [ 2-5 ]

**3-3-1** Sampling variance of the phenotypic correlation $r_{P_{x,y}}$

$$Var\left(r_{P_{x,y}}\right) = \frac{1}{4} r_{P_{x,y}}^2 \left[4\frac{Var(s_{P_{x,y}})}{s_{P_{x,y}}} + \frac{Var(s_{P_x}^2)}{s_{P_x}^2} + \frac{Var(s_{P_y}^2)}{s_{P_y}^2} - 4\frac{Cov\left(s_{P_{x,y}}, s_{P_x}^2\right)}{s_{P_{x,y}} s_{P_x}^2} - 4\frac{Cov\left(s_{P_{x,y}}, s_{P_y}^2\right)}{s_{P_{x,y}} s_{P_y}^2} + 2\frac{Cov(s_{P_x}^2, s_{P_y}^2)}{s_{P_x}^2 s_{P_y}^2}\right]$$

For details, see Appendix 6.

##### 3.3.2 Sampling variance and standard error of the coheritability estimator

Based on Robertson's (1959) approximation of the variance of the genetic correlation, it is possible to derive an approximation to the sampling variance of the coheritability as a function of the genetic correlation, the sampling variance of the heritabilities, as follows

$$Var\left(r_{A_{x,y}}\right) \approx \left(1 - r_{A_{x,y}}^2\right)^2 \frac{\sqrt{Var(h_x^2) \cdot Var\left(h_y^2\right)}}{2\, h_x^2 h_y^2}$$

$$h_x^2 h_y^2 Var\left(r_{A_{x,y}}\right) \approx (1 - r_{A_{x,y}}^2)^2 \frac{\sqrt{Var(h_x^2) \cdot Var(h_y^2)}}{2}$$

the right-hand side of the equation becomes $h_x^2 h_y^2\, Var\left(r_{A_{x,y}}\right) = Var\left(\sqrt{h_x^2 h_y^2}\, r_{A_{x,y}}\right) = Var(h_{x,y})$.

Therefore,

**3-3-2a** Sampling variance of the coheritability estimator $h_{x,y}$ (as an extension of the Robertson's approximation of the sampling variance of the genetic correlation)

$$Var\left(h_{x,y}\right) \approx \left(1 - r_{A_{x,y}}^2\right)^2 \frac{\sqrt{Var(h_x^2) \cdot Var\left(h_y^2\right)}}{2} = \left(1 - r_{A_{x,y}}^2\right)^2 \frac{SE(h_x^2) \cdot SE\left(h_y^2\right)}{2}$$

**3-3-2b** Standard error (SE) of the coheritability estimator $h_{x,y}$ (as an extension of the Robertson's approximation of the sampling variance of the genetic correlation)

$$SE\left(h_{x,y}\right) = \sqrt{Var\left(h_{x,y}\right)} = (1 - r_{A_{x,y}}^2) \sqrt{\frac{SE(h_x^2) \cdot SE(h_y^2)}{2}}$$

Another way to approximate the sampling variance of the coheritability is to use the Delta method

**3-3-2c** Sampling variance of the coheritability estimator $h_{x,y}$ (Delta Method, see Appendix 6 )

$$Var\left(h_{x,y}\right) \approx \frac{1}{4}\left(h_{x,y}\right)^2 \left[ \frac{Var\,[h_x^2]}{(h_x^2)^2} + \frac{Var\,[h_y^2]}{\left(h_y^2\right)^2} + \frac{4\,Var\,\left[r_{A_{x,y}}\right]}{r_{A_{x,y}}^2} + \frac{2\,Cov\,[h_x^2, h_y^2]}{h_x^2\; h_y^2} + \frac{4\,Cov\,\left[h_x^2\,,\; r_{A_{x,y}}\right]}{h_x^2\;\; r_{A_{x,y}}} + \frac{4\,Cov\,\left[h_y^2\,,\; r_{A_{x,y}}\right]}{h_y^2\;\; r_{A_{x,y}}} \right]$$

##### 3.3.3 Sampling variance of the coenvironmentability estimator

**3-3-3** Sampling variance of the coenvironmentability estimator $e_{x,y}$ (Delta Method)

$$Var\left(e_{x,y}\right) \approx \frac{1}{4}\left(e_{x,y}\right)^2\left[\frac{Var\left[h_x^2\right]}{\left(1-h_x^2\right)^2}+\frac{Var\left[h_y^2\right]}{\left(1-h_y^2\right)^2}+4\frac{Var\left(r_{E_{x,y}}\right)}{\left(r_{E_{x,y}}\right)^2} + 2\frac{Cov\left[h_x^2, h_y^2\right]}{\left(1-h_x^2\right)\left(1-h_y^2\right)}-4\frac{Cov\left[h_x^2, r_{E_{x,y}}\right]}{\left(1-h_x^2\right)r_{E_{x,y}}}-4\frac{Cov\left[h_y^2, r_{E_{x,y}}\right]}{\left(1-h_y^2\right)r_{E_{x,y}}}\right]$$

For details, see Appendix 6.

##### 3.3.4 Sampling variance of the heritability estimator

**3-3-4** Sampling variance of the heritability estimator $h^2 = \frac{V_A}{V_P}$ (Delta Method)

$$Var\ (\ h^2) \approx \left(h^2\right)^2 \left[ \frac{Var\ [V_A]}{(V_A)^2} + \frac{Var\ [V_P]}{(V_P)^2} + 2\frac{Cov\ [V_A, V_P]}{V_A\ V_P} \right]$$

For details, see Appendix 6.

##### 3.3.5 Sampling variance and standard error of the genetic correlation estimator

The relationships presented in this section correspond to the formulae of Robertson (1959). For extensive review on the topic of the sampling variance of the genetic correlation, see Visscher (1998) and Koots and Gibson (1996).

**3-3-5a** Sampling variance of the genetic correlation estimator (from Robertson 1959 approximation)

$$Var\left(r_{A_{x,y}}\right) \approx \left(1 - r_{A_{x,y}}^2\right)^2 \frac{\sqrt{Var(h_x^2) \cdot Var\left(h_y^2\right)}}{2\ h_x^2 h_y^2} = \left(1 - r_{A_{x,y}}^2\right)^2 \frac{SE(h_x^2) \cdot SE\left(h_y^2\right)}{2\ h_x^2 h_y^2}$$

**3-3-5b** Standard error of the genetic correlation estimator (from Robertson 1959 aproximation)

$$SE(r_{A_{x,y}}) = \sqrt{Var\left(r_{A_{x,y}}\right)} \approx \sqrt{\left(1 - r_{A_{x,y}}^2\right)^2 \frac{SE(h_x^2) \cdot SE\left(h_y^2\right)}{2\ h_x^2 h_y^2}} = \left(1 - r_{A_{x,y}}^2\right) \sqrt{\frac{SE(h_x^2) \cdot SE\left(h_y^2\right)}{2\ h_x^2 h_y^2}}$$

### 4 Distribution of the sample correlation coefficient

#### 4.1 Derivation of the distribution of the sample correlation coefficient

This section succinctly elaborates on the distribution of the sample phenotypic correlation in the case when the correlation parameter $\rho$ is zero, and nonzero.

##### 4.1.1 Distribution of $r$ when $\rho$ is zero

When $\rho = 0$, the three variates $r$ , $s_x^2$ and $s_y^2$ are a completely independent set, since $s_x^2$ and $s_y^2$ are functions of separate sets of independent variates and are therefore mutually independent. Further details can be found elsewhere ( Fisher 1928, Kym 1968, Chance 1986).

**4-1-1 Distribution of the sample correlation coefficient when $\rho = 0$**

$$f_r(r|\rho = 0\,, n) = \frac{\Gamma\left(\frac{n-1}{2}\right)}{\pi^{\frac{1}{2}}\,\Gamma\left(\frac{n-2}{2}\right)}(1-r)^{\frac{\eta-4}{2}}\,dr \qquad -1 \leq r \leq +1$$

##### 4.1.2 Distribution of $r$ when $\rho$ is nonzero

The exact density function of the sample correlation coefficient $r$ was originally obtained by Fisher (1915, 1928), following a geometric argument For ease of exposition, I present an exegesis of the derivation of the distribution the work by Hotelling (1950) (whose equation numbers are kept in this section for easy reference) .For purposes of this explanation we will use $\eta = (n-1)$, where $n$ is the sample size.

The sample variances follow a chi-square distribution

**Equation 10 Distribution of the sample variances**

| sample variance | chi-square distribution of the sample variance | |
|---|---|---|
| $s_x^2 = \frac{\Sigma(x_i-\bar{x})^2}{\eta}$ , | $\frac{\eta s_x^2}{\sigma_x^2} \sim \chi_\eta^2$ | $f_{s_x}\left(\frac{\eta s_x^2}{2}\right) = \frac{\left(\frac{\eta s_x^2}{2}\right)^{\frac{\eta}{2}-1} e^{-\frac{\eta s_x^2}{2}}}{\Gamma\left(\frac{\eta}{2}\right)} \, d\left(\frac{\eta s_x^2}{2}\right)$ |
| $s_y^2 = \frac{\Sigma(y_i-\bar{y})^2}{\eta}$ , | $\frac{\eta s_y^2}{\sigma_y^2} \sim \chi_\eta^2$ | $f_{s_y}\left(\frac{\eta s_y^2}{2}\right) = \frac{\left(\frac{\eta s_y^2}{2}\right)^{\frac{\eta}{2}-1} e^{-\frac{\eta s_y^2}{2}}}{\Gamma\left(\frac{\eta}{2}\right)} \, d\left(\frac{\eta s_y^2}{2}\right)$ |

Multiplying together the independent probability density functions of the correlation coefficient (when $\rho = 0$), the sample variance for trait $x$, and the sample variance for trait $y$.

Equation 2 Equation 10

$$\underbrace{f_r(r|\rho=0\,,\eta)} \quad \cdot \quad \underbrace{f_{s_x}\left(\frac{\eta s_x^2}{2}\right)} \quad \cdot \quad \underbrace{f_{s_y}\left(\frac{\eta s_y^2}{2}\right)}$$

$$g(r, s_x, s_y|\rho=0)\, dr\, ds_x\, ds_y = \left[\frac{\Gamma\left(\frac{\eta}{2}\right)}{\pi^{\frac{1}{2}}\,\Gamma\left(\frac{\eta-1}{2}\right)}(1-r^2)^{\frac{\eta-3}{2}}\, dr\right]\left[\frac{\left(\frac{\eta s_x^2}{2}\right)^{\frac{\eta}{2}-1} e^{-\frac{\eta s_x^2}{2}}}{\Gamma\left(\frac{\eta}{2}\right)}\, d\left(\frac{\eta s_x^2}{2}\right)\right]\left[\frac{\left(\frac{\eta s_y^2}{2}\right)^{\frac{\eta}{2}-1} e^{-\frac{\eta s_y^2}{2}}}{\Gamma\left(\frac{\eta}{2}\right)}\, d\left(\frac{\eta s_y^2}{2}\right)\right]$$

$$= \frac{1}{\pi^{\frac{1}{2}}\,\Gamma\left(\frac{\eta-1}{2}\right)\,\Gamma\left(\frac{\eta}{2}\right)}(1-r^2)^{\frac{\eta-3}{2}}\, dr\,.\ \frac{\eta^{\frac{\eta-2}{2}} {s_x}^{2\left(\frac{\eta-2}{2}\right)} e^{-\frac{\eta s_x^2}{2}}}{2^{\frac{\eta-2}{2}}}\left(\frac{2\eta s_x}{2} ds_x\right)\ \frac{\eta^{\frac{\eta-2}{2}} {s_y}^{2\left(\frac{\eta-2}{2}\right)} e^{-\frac{\eta s_y^2}{2}}}{2^{\frac{\eta-2}{2}}}\left(\frac{2\eta s_y}{2} ds_y\right)$$

$$= \frac{(1-r^2)^{\frac{\eta-3}{2}}\, dr}{\pi^{\frac{1}{2}}\,\Gamma\left(\frac{\eta-1}{2}\right)\,\Gamma\left(\frac{\eta}{2}\right)} \cdot \frac{\eta^{\frac{\eta-2}{2}+\frac{\eta-2}{2}+1+1}\ {s_x}^{\eta-2+1}\ {s_y}^{\eta-2+1}\ e^{-\frac{\eta}{2}(s_x^2+s_x^2)}}{2^{\frac{\eta-2}{2}+\frac{\eta-2}{2}}}\, ds_x\, ds_y$$

After some rearrangement, we obtain:

**Equation 11 The joint distribution of $r\ , s_x\ , s_y$**

$$g(r, s_x, s_y|\rho = 0)\ dr\ ds_x\ ds_y = \frac{\eta^\eta\ (1-r^2)^{\frac{\eta-3}{2}}}{\pi^{\frac{1}{2}}\ 2^{\eta-2}\ \Gamma\left(\frac{\eta-1}{2}\right)\Gamma\left(\frac{\eta}{2}\right)}\ .\ e^{-\frac{\eta}{2}(s_x^2+s_x^2)}(s_x s_y)^{\eta-1}\ ds_x\ ds_y\ dr$$

$$= \frac{(1-r^2)^{\frac{\eta-3}{2}}\ dr}{\pi^{\frac{1}{2}}\Gamma\left(\frac{\eta-1}{2}\right)\ \Gamma\left(\frac{\eta}{2}\right)}\ .\frac{\eta^\eta\ \ e^{-\frac{\eta}{2}(s_x^2+s_x^2)}(s_x s_y)^{\eta-1}}{2^{\eta-2}}\ ds_x\ ds_y\ dr$$

**Equation 12** Proof that the denominator $\pi^{\frac{1}{2}}\Gamma\left(\frac{\eta-1}{2}\right)\ \Gamma\left(\frac{\eta}{2}\right)2^{\eta-2} = \pi(\eta-2)!$

Let $(\eta - 1) = m$

$$\eta = m + 1$$

Then $\pi^{\frac{1}{2}}\Gamma\left(\frac{m}{2}\right)\ \Gamma\left(\frac{m+1}{2}\right)\ 2^{m-1}$

The product of the gamma functions

$$\Gamma\left(\frac{m}{2}\right)\Gamma\left(\frac{m+1}{2}\right) = \frac{\pi^{\frac{1}{2}}\Gamma(m)}{2^{m-1}}$$

$$\pi^{\frac{1}{2}}\ \frac{\pi^{\frac{1}{2}}\ \Gamma(m)}{2^{m-1}}\ 2^{m-1} = \pi^{\frac{1}{2}}\ \pi^{\frac{1}{2}}\ \Gamma(m) = \pi(m-1)!\ = \pi\ (\eta-2)!$$

Consider two random variables $X{\sim}Normal(0,1)$ and $Y{\sim}Normal(0,1)$

We assume that each $(x, y)$ datum has a bivariate normal probability density

$$f_{X,Y}(x,y) = \frac{1}{2\pi(1-\rho^2)^{\frac{1}{2}}}\ e^{\left\{-\frac{1}{2(1-\rho^2)}\ (x^2-2\rho xy+y^2)\right\}}dx\ dy$$

Multiplying the independent probability density functions assumed for each bivariate datum, it yields the joint distribution of the $2\eta$ observations which is written with the help of equations 10 and 11.

$$\varphi(r,s_x,s_y,\rho) = \prod_{i=1}^{\eta} f_{XY_i}(x,y) = \prod_{i=1}^{\eta} \frac{\eta}{2\pi^{\eta}(1-\rho^2)^{\frac{1}{2}\eta}}\, e^{\left\{-\frac{1}{2(1-\rho^2)}\,(x_i^2 - 2\rho x_i y_i + y_i^2)\right\}} dx_i dy_i$$

$$= \frac{1}{2\pi^{\eta}(1-\rho^2)^{\frac{1}{2}\eta}}\, e^{-\frac{\eta}{2(1-\rho^2)}(\sum x_i^2 - 2\rho \sum x_i y_i + \sum y_i^2)} dx_i dy_i$$

Since we know that $\bar{x} = \bar{y} = 0$, the following relationships can be use to replace terms of $\sum x_i^2$, $\sum x_i\, y_i$, and $\sum y_i^2$.

$$s_x^2 = \frac{\sum(x_i-\bar{x})^2}{\eta} \quad \rightarrow \quad \sum(x_i-\bar{x})^2 = \eta s_x^2 \quad \rightarrow \sum x_i^2 = \eta s_x^2$$

$$r = \frac{\sum(x_i-\bar{x})(y_i-\bar{y})}{\eta\, s_y\, s_x} \rightarrow \quad \sum(x_i-\bar{x})(y_i-\bar{y}) = \eta\, s_y\, s_x\, r \quad \rightarrow \sum x_i\, y_i = \eta\, s_y\, s_x\, r$$

$$s_y^2 = \frac{\sum(y_i-\bar{y})^2}{\eta} \quad \rightarrow \quad \sum(y_i-\bar{y})^2 = \eta s_y^2 \quad \rightarrow \sum y_i^2 = \eta s_y^2$$

We obtain

**Equation 14**

$$\varphi(r,s_x,s_y,\rho) dx_1 dy_1\, dx_2 dy_{2\ldots}\, dx_{\eta} dy_{\eta} = \prod_{i=1}^{\eta} f_{XY_i}(x,y)$$

which becomes

**Equation 15** **Joint distribution of $r$, $s_x$, $s_y$, $\rho$**

$$\varphi(r,s_x,s_y,\rho) = \prod_{i=1}^{n} f_{XY_i}(x,y) = = \frac{1}{2\pi^{\eta}(1-\rho^2)^{\frac{\eta}{2}}}\, e^{-\frac{\eta}{2(1-\rho^2)}(s_x^2 - 2\rho r s_x s_y + s_y^2)} dx_1 dy_1 \ldots dx_n dy_n$$

When $\rho$ varies, the joint probability elements of $r, s_x$, $s_y$ varies in proportion to the density of equation 15.

The derivation of the joint probability distribution of $r, s_x$, $s_y$, $\rho$ for $\rho = 0$

$$f(r\,, s_x\,, s_y) = \frac{[\,Equation\ 15\,]}{[\,Equation\ 15\ when\ \rho = 0\,]}[\,equation\ 11\,]$$

$$= \frac{\varphi(r, s_x, s_y, \rho) dx_1 dy_1 \dots dx_n dy_n}{\varphi(r, s_x, s_y, 0) dx_1 dy_1 \dots dx_n dy_n}\, g(r, s_x, s_y)\, ds_x\, ds_y\, dr$$

$$= \frac{\dfrac{1}{2\pi^{\eta}(1-\rho^2)^{\frac{\eta}{2}}}\, e^{-\frac{\eta}{2(1-\rho^2)}(s_x^2 - 2\rho r s_x s_y + s_y^2)}}{\dfrac{1}{2\pi^{\eta}}\, e^{-\frac{\eta}{2}(s_x^2 + s_y^2)}} \frac{(1-r^2)^{\frac{\eta-3}{2}}}{\pi^{\frac{1}{2}}\, \Gamma\left(\frac{\eta-1}{2}\right)\, \Gamma\left(\frac{\eta}{2}\right)} \cdot \frac{\eta^{\eta}\, e^{-\frac{\eta}{2}(s_x^2 + s_y^2)} (s_x s_y)^{\eta-1}}{2^{\eta-2}} ds_x\, ds_y dr$$

**Equation 16** **The joint probability distribution of $r,\ s_x,\ s_y, \rho$ for $\rho = 0$**

$$f(r\,, s_x\,, s_y) = \frac{\eta^{\eta}}{\pi\,(\eta-2)!}(1-\rho^2)^{-\frac{\eta}{2}}(1-r^2)^{\frac{\eta-3}{2}}\, e^{\left\{-\frac{\eta}{2(1-\rho^2)}(s_x^2 - 2\rho r s_x s_y + s_y^2)\right\}}(s_x s_y)^{\eta-1}\, ds_x\, ds_y\, dr$$

The distribution of $r$ is found by integrating (16) with respect to $s_x$ and $s_y$ from 0 to positive infinity. For this purpose the following transformation is useful.

Put

$$s_x = \alpha^{\frac{1}{2}}\, e^{-\frac{1}{2}\beta}$$

$$s_y = \alpha^{\frac{1}{2}}\, e^{\frac{1}{2}\beta}$$

Note that $s_x s_y = \alpha$

The Jacobian is

$$J = \frac{\delta(s_x, s_y)}{\delta(\alpha, \beta)} = \begin{bmatrix} \dfrac{\delta s_x}{\delta\, \alpha} & \dfrac{\delta s_x}{\delta\, \beta} \\ \dfrac{\delta s_y}{\delta\, \alpha} & \dfrac{\delta s_y}{\delta\, \beta} \end{bmatrix} = \begin{bmatrix} \dfrac{\delta}{\delta\, \alpha}\left(\alpha^{\frac{1}{2}}\, e^{-\frac{1}{2}\beta}\right) & \dfrac{\delta}{\delta\, \beta}\left(\alpha^{\frac{1}{2}}\, e^{-\frac{1}{2}\beta}\right) \\ \dfrac{\delta}{\delta\, \alpha}\left(\alpha^{\frac{1}{2}}\, e^{\frac{1}{2}\beta}\right) & \dfrac{\delta}{\delta\, \beta}\left(\alpha^{\frac{1}{2}}\, e^{\frac{1}{2}\beta}\right) \end{bmatrix}$$

$$= \begin{bmatrix} \dfrac{1}{2}\alpha^{\frac{1}{2}}\, e^{-\frac{1}{2}\beta} & -\dfrac{1}{2}\alpha^{\frac{1}{2}}\, e^{-\frac{1}{2}\beta} \\ \dfrac{1}{2}\alpha^{\frac{1}{2}}\, e^{\frac{1}{2}\beta} & \dfrac{1}{2}\alpha^{\frac{1}{2}}\, e^{\frac{1}{2}\beta} \end{bmatrix}$$

Now working in the exponential part of equation 16

$$f(s_x, s_y, r) = \frac{\eta^\eta}{\pi\,(\eta-2)!}(1-\rho^2)^{-\frac{\eta}{2}}(1-r^2)^{\frac{\eta-3}{2}}\ e^{\left\{-\frac{\eta}{2(1-\rho^2)}\left(s_x^2-2\rho r s_x s_y+s_y^2\right)\right\}}(s_x s_y)^{\eta-1} ds_x\, ds_y\, dr$$

$$\left\{-\frac{\eta}{2(1-\rho^2)}\left(s_x^2 - 2\rho r s_x s_y + s_y^2\right)\right\}$$

$$= \left\{-\frac{\eta}{2(1-\rho^2)}\left(\left(\alpha^{\frac{1}{2}} e^{-\frac{1}{2}\beta}\right)^2 - 2\rho r\left(\alpha^{\frac{1}{2}} e^{-\frac{1}{2}\beta}\right)\left(\alpha^{\frac{1}{2}} e^{\frac{1}{2}\beta}\right) + \left(\alpha^{\frac{1}{2}} e^{\frac{1}{2}\beta}\right)^2\right)\right\}$$

$$= -\frac{\eta}{2(1-\rho^2)}\left(\alpha e^{-\beta} - 2\,\rho r\alpha + \alpha e^{\beta}\right)$$

$$= -\frac{\eta\,\alpha}{(1-\rho^2)}\left(\frac{e^{-\beta}+e^{\beta}}{2} - \frac{2\,\rho r}{2}\right)$$

$$= -\frac{\eta\,\alpha}{(1-\rho^2)}\left(\cosh\beta - \rho r\right)$$

Working on the differentials $ds_x\, ds_y$ of equation 16.

$$f(s_x, s_y, r) = \frac{\eta^\eta}{\pi\,(\eta-2)!}(1-\rho^2)^{-\frac{\eta}{2}}(1-r^2)^{\frac{\eta-3}{2}}\ e^{\left\{-\frac{\eta}{2(1-\rho^2)}\left(s_x^2-2\rho r s_x s_y+s_y^2\right)\right\}}\left(s_x s_y\right)^{\eta-1} ds_x\, ds_y\, dr$$

We note that

$$\frac{\delta\,(s_x s_y)}{\delta\, s_x} = \frac{\delta\,\alpha}{\delta\, s_x} = s_y \quad \rightarrow \quad d\alpha = ds_x \cdot s_y$$

and

$$\frac{s_x}{s_y} = \frac{\alpha^{\frac{1}{2}} e^{\frac{1}{2}\beta}}{\alpha^{\frac{1}{2}} e^{-\frac{1}{2}\beta}} = e^{\beta} \quad \rightarrow \quad ln\left(\frac{s_x}{s_y}\right) = ln\left(e^{\beta}\right) = \beta$$

$$\frac{d\,\beta}{d\,s_y} = \frac{d\left(\ln\frac{s_x}{s_y}\right)}{d\,s_y} = \frac{1}{s_y} \quad \rightarrow \quad d\beta = \frac{1}{s_y} ds_y$$

Therefore, the product

$$d\alpha\, d\beta = ds_x \cdot s_y \,\frac{1}{s_y} ds_y = ds_x\, ds_y$$

We mark the factors involving $s_x$ and $s_y$ in Equation 16 and express them in terms of $\alpha$ and $\beta$.

$$e^{\left\{-\frac{\eta}{2(1-\rho^2)}\left(s_x^2 - 2\rho r s_x s_y + s_y^2\right)\right\}}(s_x s_y)^{\eta-1} ds_x\, ds_y \;=\; \left|\frac{1}{2}\right| \alpha^{\eta-1}\, e^{-\eta\frac{\alpha}{1-\rho^2}[\cosh\beta - \rho r]} d\alpha\, d\beta$$

Integration with respect to $\alpha$ (with integration limits $0 < \alpha < +\infty$)

$$\frac{1}{2}\int_0^{+\infty} e^{-\eta\frac{\alpha}{1-\rho^2}[\cosh\beta - \rho r]}\; \alpha^{\eta-1}\, d\alpha\, d\beta$$

$$\alpha = a$$

$$(1-\rho^2) = y$$

$$\cosh\beta - \rho r = x$$

$$\frac{1}{2}\int_0^{\infty} e^{-\eta\frac{a}{y}x}\; a^{\eta-1}\, da\, d\beta$$

Steps to integrate with respect to $a$ (Equation 16b)

$$\frac{1}{2}\int_0^{\infty} e^{-\eta\,\frac{a}{y}x}\ \ a^{\eta-1}\,da\,d\beta$$

① substitute

$$u = \frac{a}{y}\ \ \rightarrow\ \ a = uy \qquad and \qquad \frac{\delta a}{\delta u} = \frac{\delta(uy)}{\delta u} = y\ \ \rightarrow\ \ \delta a = y\,\delta u$$

Use $a^{\eta-1} = y^{\eta-1}u^{\eta-1}$

$$u = \frac{a}{y}$$

$$\frac{1}{2}\int_0^{\infty} y^{\eta-1}\,u^{\eta-1}e^{-\eta\,ux}\ y\,du\,d\beta$$

$$\frac{1}{2}y^{\eta}\int_0^{\infty} u^{\eta-1}e^{-\eta\,xu}\ du\,d\beta$$

② substitute in $\frac{1}{2}y^{\eta}\int_0^{\infty} u^{\eta-1}e^{-\eta\,ux}\ du\,d\beta$

$b = \eta - 1 \quad \rightarrow \quad b + 1 = \eta$

$m = x(b+1) \quad \rightarrow \quad m = x\eta$

$$\frac{1}{2}y^{\eta}\int_0^{\infty} u^{b}e^{-mu}\ du\,d\beta = \frac{1}{2}y^{\eta}\frac{\Gamma(b+1)}{m^{b+1}}d\beta$$

Undoing of variable substitution above

③ Undo substitution

$$\frac{1}{2}y^{\eta}\frac{\Gamma(b+1)}{m^{b+1}}d\beta \quad \rightarrow \quad \frac{1}{2}y^{\eta}\frac{\Gamma(\eta)}{x^{\eta}\,\eta^{\eta}}d\beta\ = \frac{1}{2}(1-\rho^2)^{\eta}\ \frac{\Gamma(\eta)}{\eta^{\eta}}\ \frac{1}{(cosh\beta - \ \rho r)^2}d\beta$$

④ Undo variable change. Finally we obtain

$$\frac{1}{2}\int_0^{\infty} e^{-\eta\,\frac{a}{y}x}\ \ a^{\eta-1}\,da\,d\beta = \frac{1}{2}\frac{(1-\rho^2)^{\eta}\,\Gamma(\eta)}{\eta^{\eta}}\frac{1}{(\cosh\beta - \rho r)^{\eta}}d\beta$$

$$= \frac{1}{2}\Gamma(\eta)\left(\frac{1-\rho^2}{\eta}\right)^{\eta}\frac{1}{(\cosh\beta - \rho r)^{\eta}}d\beta$$

Now we can reformulate the integrant as follows

$$\frac{1}{2}\int_0^{\infty} e^{-\frac{\eta\alpha}{1-\rho^2}[\cosh\beta-\rho r]}\ \alpha^{\eta-1}d\alpha\, d\beta$$

Finally, integrating with respect to $\alpha$,

$$\frac{1}{2}\int_{\alpha=0}^{\alpha\to+\infty} \alpha^{\eta-1}\, e^{-\frac{\eta\alpha}{1-\rho^2}[\cosh\beta-\rho r]}\ d\alpha\, d\beta\ =\ \frac{1}{2}(\eta-1)!\left(\frac{1-\rho^2}{\eta}\right)^{\eta}\frac{1}{(\cosh\beta-\rho r)^{\eta}}d\beta$$

Equation 16c

Integration with respect to $\beta$ (integration limits $-\infty<\beta<+\infty$), but since we work from $0$ to $+\infty$, we multiply by 2.

$$\frac{1}{2}(\eta-1)!\left(\frac{1-\rho^2}{\eta}\right)^{\eta}\ 2\int_0^{+\infty}\frac{1}{(\cosh\beta-\rho r)^{\eta}}d\beta$$

Up to now, we have arrived to the following expression

$$f(r,\rho)=\frac{\eta^{\eta}}{\pi\,(\eta-2)!}(1-\rho^2)^{-\frac{\eta}{2}}\,(1-r^2)^{\frac{1}{2}(\eta-3)}\,(\eta-1)!\left(\frac{1-\rho^2}{\eta}\right)^{\eta}\left[\int_0^{\infty}\frac{1}{(\cosh\beta-\rho r)^{\eta}}d\beta\right]dr$$

After some simplification and rearrangement, the general distribution of $r$ is

$$f_n(r,\rho)dr=\frac{\eta-1}{\pi}(1-\rho^2)^{\frac{\eta}{2}}\ (1-r^2)^{\frac{1}{2}(\eta-3)}\left[\int_0^{\infty}\frac{1}{(\cosh\beta-\rho r)^{\eta}}d\beta\right]dr$$

Introducing the notation

**Equation 17**

$$I_n(\rho r)=\int_0^{\infty}\frac{1}{(\cosh\beta-\rho r)^{\eta}}d\beta \qquad |p|<1$$

We replace the integral for equation 17

$$f_\eta(r,\rho)dr = \frac{\eta-1}{\pi}(1-\rho^2)^{\frac{\eta}{2}}\ (1-r^2)^{\frac{1}{2}(\eta-3)}I_n(\rho r)dr$$

The difficulties with the form presented lies in the integral factor, which will be denoted as $I_n(\rho r)$.
We can re-express

**Equation 18**

$$f_\eta(r,\rho)dr = \frac{\eta-1}{\pi}(1-\rho^2)^{\frac{1}{2}\eta}\ (1-r^2)^{\frac{1}{2}(\eta-3)}I_n(\rho r)dr$$

Introducing the notation $p = \rho r$

$$I_\eta(p) = \int_0^\infty \frac{1}{(\cosh\beta - p)^n}d\beta$$

A fundamental improvement is the use of the following substitution:

**Equation 21**

$$\cosh\beta = \frac{1-pz}{1-z}$$

Therefore

$$\cosh\beta - p = \left(\frac{1-pz}{1-z} - p\right) = \frac{1-p}{1-z}$$

$$\beta = arcosh\left(\frac{1-pz}{1-z}\right)$$

Taking the derivative of $\beta$ with respect to $z$,

$$\frac{d\beta}{dz} = \frac{1-p}{(z-1)^2\sqrt{\frac{(1-pz)^2}{(1-z)^2}-1}}$$

$$= \frac{1-p}{(z-1)^2\sqrt{\frac{1-pz}{1-z}-1}\sqrt{\frac{1-pz}{1-z}+1}}$$

$$= \frac{1-p}{(z-1)^2\sqrt{\frac{z(1-p)}{1-z}}\sqrt{\frac{2-z-pz}{1-z}}}$$

$$= \frac{(1-p)(1-z)}{(z-1)^2\sqrt{z}\sqrt{1-p}\sqrt{2-z-pz}}$$

Finally, we note that the term $(z-1)^2 = (1-z)^2$ and after simplification

$$d\beta = \frac{(1-p)^{\frac{1}{2}}}{(1-z)\sqrt{z}\sqrt{2-z-pz}}dz$$

Thus

$$I_\eta(p) = \int_0^\infty \frac{1}{(\cosh\beta - p)^\eta}d\beta = \int_0^\infty \frac{(1-z)^\eta}{(1-p)^\eta}\frac{(1-p)}{(1-z)\sqrt{z}\sqrt{2-z-pz)}}dz$$

After some simplification and rearrangement we obtain

$$I_\eta(p) = \int_0^\infty (1-p)^{-\eta+\frac{1}{2}}\, z^{-\frac{1}{2}}\,(1-z)^{\eta-1}\frac{1}{\sqrt{2-z-pz}}dz$$

Multiplying the integrand by $\frac{\sqrt{2}}{\sqrt{2}}$

$$I_\eta(p) = \int_0^\infty (1-p)^{-\eta+\frac{1}{2}}\, z^{-\frac{1}{2}}\,(1-z)^{\eta-1}\frac{1}{\sqrt{2-z-pz}}\frac{\sqrt{2}}{\sqrt{2}}dz$$

**Equation 22**

$$I_\eta(p) = \frac{1}{\sqrt{2}}\,(1-p)^{-\eta+\frac{1}{2}} \int_0^\infty z^{-\frac{1}{2}}\,(1-z)^{\eta-1}\left(1-\frac{(1+p)}{2}z\right)^{-\frac{1}{2}} dz$$

This integral has the kernel of a hypergeometric integral

$${}_2F_1(a_1, a_2\,;\, b_1; x) = \frac{\Gamma(b_1)}{\Gamma(a_2)\Gamma(b_1-a_2)} \int_0^1 z^{a_2}(1-z)^{b_1-a_2}(1-kz)^{-a_1} dz$$

$$\frac{\Gamma(a_2)\Gamma(b_1-a_2)}{\Gamma(b_1)} \cdot {}_2F_1(a_1, a_2\,;\, b_1; x) = \int_0^1 z^{a_2}(1-z)^{b_1-a_2}(1-kz)^{-a_1} dz$$

where $a_1 = \frac{1}{2}$, $a_2 = \frac{1}{2}$, $b_1 = \eta + \frac{1}{2}$, $k = \frac{1+p}{2}$

Replacing this variables in equation we obtain

$$I_n(p) = \frac{1}{\sqrt{2}}\,(1-p)^{-\eta+\frac{1}{2}} \int_0^\infty z^{-\frac{1}{2}}\,(1-z)^{\eta-1}\left(1-\frac{(1+p)}{2}z\right)^{-\frac{1}{2}} dz$$

$$I_n(p) = \frac{1}{\sqrt{2}}\,(1-p)^{-\eta+\frac{1}{2}}\,\frac{\Gamma\left(\frac{1}{2}\right)\Gamma(\eta)}{\Gamma(\eta+\frac{1}{2})} \cdot {}_2F_1\left(\frac{1}{2}, \frac{1}{2}\,;\, \eta+\frac{1}{2}; \frac{1+p}{2}\right)$$

Since $\Gamma\left(\frac{1}{2}\right) = \pi^{\frac{1}{2}}$

$$I_n(p) = \frac{1}{\sqrt{2}}\,(1-p)^{-\eta+\frac{1}{2}}\,\frac{\pi^{\frac{1}{2}}\,\Gamma(\eta)}{\Gamma(\eta+\frac{1}{2})} \cdot {}_2F_1\left(\frac{1}{2}, \frac{1}{2}\,;\, \eta+\frac{1}{2}; \frac{1+p}{2}\right)$$

**Equation 24**

$$I_n(p) = \frac{\pi^{\frac{1}{2}}\,\Gamma(\eta)}{\sqrt{2}\,\Gamma(\eta+\frac{1}{2})}(1-p)^{-\eta+\frac{1}{2}}\;{}_2F_1\left(\frac{1}{2}, \frac{1}{2}\,;\, \eta+\frac{1}{2}; \frac{1+p}{2}\right)$$

Replacing $p = \rho r$, we obtain

$$I_\eta(\rho r) = \frac{\pi^{\frac{1}{2}}\,\Gamma(\eta)}{\sqrt{2}\,\Gamma(\eta+\frac{1}{2})}(1-\rho r)^{-\eta+\frac{1}{2}}\ {}_2F_1\left(\frac{1}{2},\frac{1}{2};\eta+\frac{1}{2};\frac{1+\rho r}{2}\right)$$

$$f_\eta(r|\rho,\eta)dr = \frac{\eta-1}{\pi}(1-\rho^2)^{\frac{\eta}{2}}\ (1-r^2)^{\frac{1}{2}(\eta-3)} I_\eta(\rho r)dr$$

Replacing this $I_\eta(\rho r)$ term in Equation 18, we obtain

**Equation 25**

$$f_\eta(r|\rho,\eta) = \frac{\eta-1}{\pi}(1-\rho^2)^{\frac{\eta}{2}}\ (1-r^2)^{\frac{1}{2}(\eta-3)} \frac{\pi^{\frac{1}{2}}\,\Gamma(\eta)}{\sqrt{2}\,\Gamma(\eta+\frac{1}{2})}(1-\rho r)^{-\eta+\frac{1}{2}}\ {}_2F_1\left(\frac{1}{2},\frac{1}{2};\eta+\frac{1}{2};\frac{1+\rho r}{2}\right)$$

which converges for $-1 < \rho r < +1$.

Finally, we exchange $\eta = n-1$, and obtain the sampling distribution of the correlation coefficient in terms of the sample size $n$.

Probability density function of the distribution of the sample correlation coefficient in terms of $\rho, n$

$$\boldsymbol{f_{\mathrm{r}}(r|\rho,n) = \frac{n-2}{\sqrt{2\pi}}\frac{\Gamma(n-1)}{\Gamma\left(n-\frac{1}{2}\right)}\ (1-\ \rho^2)^{\frac{n-1}{2}}\ (1-r^2)^{\frac{n-4}{2}}\ (1-\ \rho r)^{-n+\frac{3}{2}} \cdot {}_2F_1\left(\frac{1}{2},\frac{1}{2};n-\frac{1}{2};\frac{1+\rho r}{2}\right)}$$

The hypergeometric function is expanded in Appendix 5. By using the first three terms one can obtain a very good approximation of the probability density function of $r$.

$$\boldsymbol{f_{\mathrm{r}}(r|\,\rho,n) \approx \frac{n-2}{\sqrt{2\pi}}\frac{\Gamma(n-1)}{\Gamma\left(n-\frac{1}{2}\right)}(1-\ \rho^2)^{\frac{n-1}{2}}(1-r^2)^{\frac{n-4}{2}}(1-\ \rho r)^{-n+\frac{3}{2}}\left[1+\frac{1}{4}\frac{(1-\rho r)}{(2n-1)}+\frac{9}{32}\frac{(1+\rho r)^2}{(2n-1)(2n+1)}+\cdots\right]}$$

#### 4.2 Comments on the distribution of $r$

##### 4.2.1 General comments

The shape of the sample correlation coefficient when $\rho = 0$ is symmetric and/or when the sample size is very large. The distribution has a zero skew value indicating that the left and right hand sides of the distribution are roughly equally balanced around the mean.

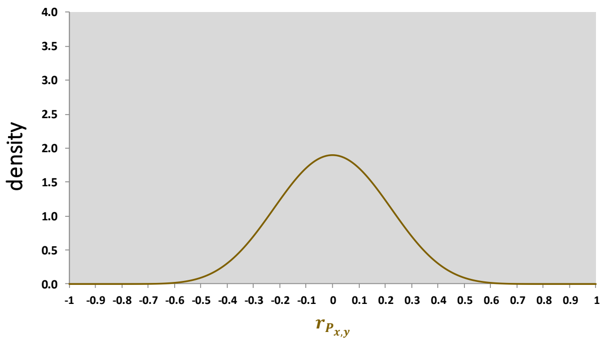

However, the shape of the sampling distribution of the correlation coefficient is asymmetric when it differs from $\rho = 0$. This means that the tail on one side is longer or fatter than the other side. The reason for this asymmetry of the probability distribution is that the correlation coefficient cannot take values less than -1 or greater than +1. The skew of the distribution is more noticeable when the sample size is small ($n < 25$).

When $\rho < 0$ the distribution is positively skewed.

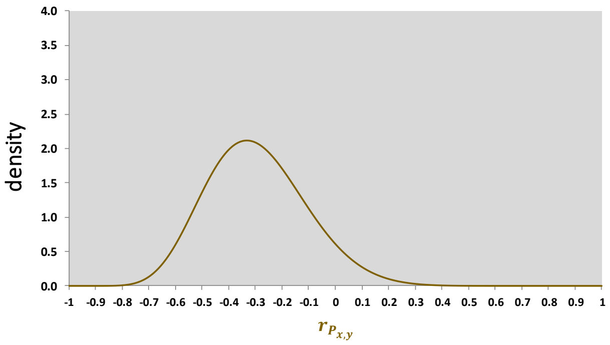

When $r > 0$, the distribution is negatively skewed.

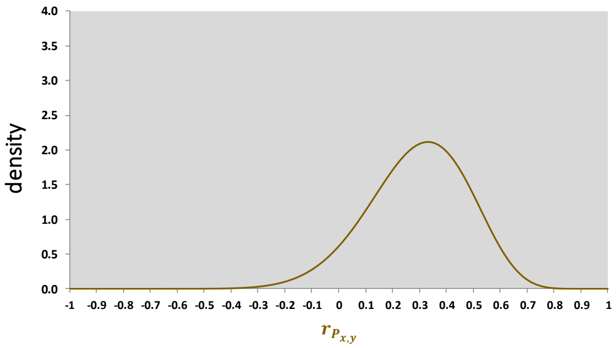

As the sample size $n$ increases
the skew is less pronounced,
the distribution becomes narrower
(variance decreases),
and density ($y$ axis) reaches larger values

$n =$ 25

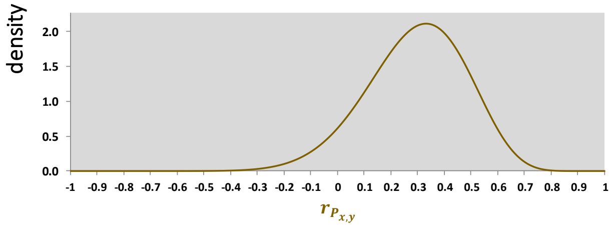

$n =$ 50

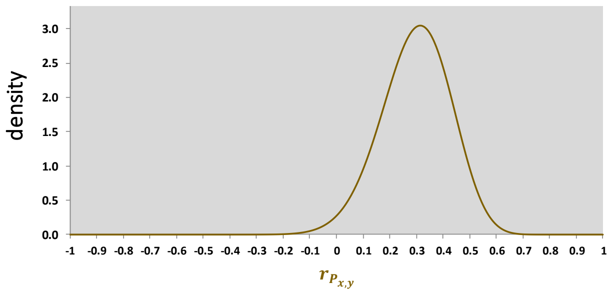

$n =$ 75

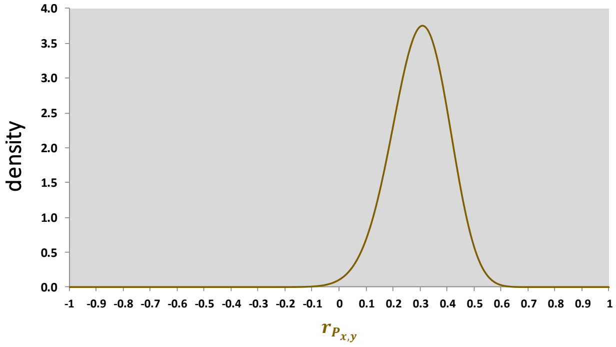

$n =$ 100

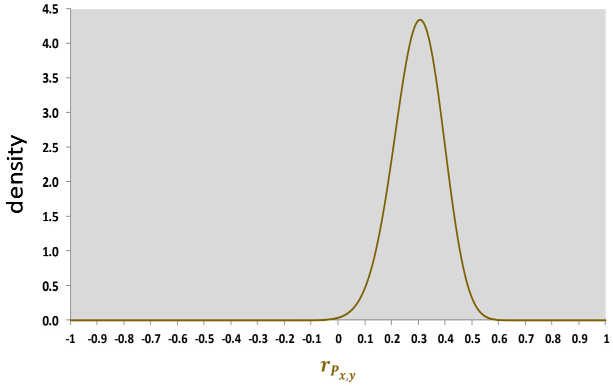

As $\rho$ approaches $+1$ or $-1$, the sampling variance decreases, so that when $\rho$ is either at $+1$ or $-1$, all sample values equal the parameter and the sampling variance is zero.

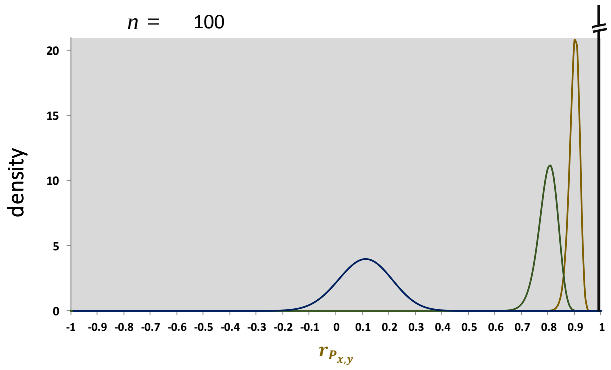

Note that as |ρ| approaches 1, the sampling variance approaches zero.

The shape of the sampling distribution depends on the sample size $n$. The shape becomes increasingly normal with large values of $n$, and becomes increasingly skewed with increasing $|\rho|$.

##### 4.2.2 Comments on the distribution of $r$ when the correlation coefficient parameter $\rho$ is zero

Independence implies zero correlation, but not the converse.

If the population phenotypic correlation is zero, then two possible cases exist:

(a) both the population coheritability $H_{x,y}$ and the population coenvironmentability $E_{x,y}$ are zero,

or

(b) both the population coheritability $H_{x,y}$ and the population coenvironmentability $E_{x,y}$ are nonzero, have the same magnitude, but differ in sign.

In the latter case, one must be mindful that a zero phenotypic correlation does not necessarily indicate that the coheritability and coenvironmentability are zero. From a statistical point of view, their values may be significantly different than zero, yet possess a different sign.

Examples of distribution of $r_{P_{x,y}}$, $h_{x,y}$, and $e_{x,y}$ when the parameter values of the population-level coheritability and coenvironmentability sum up to zero.

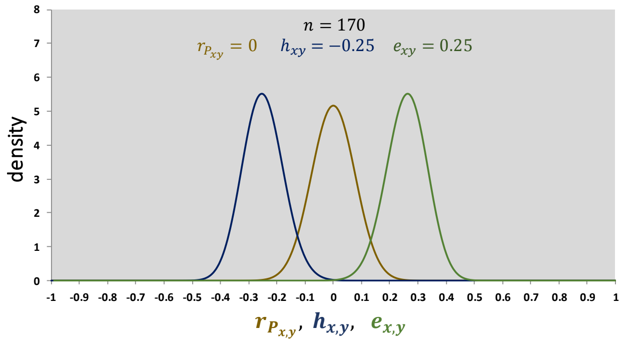

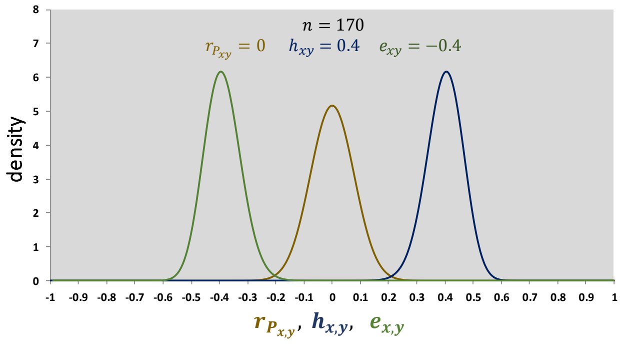

##### 4.2.3 Comments on the distribution of $r$ when the correlation coefficient parameter $\rho$ is nonzero

The closed-form representation of the probability density function of the sample correlation coefficient $r$ is dependent on the population correlation coefficient $\rho$ and the sample size $n$ (see section 4.1.2). Figure 4.2.3-1 presents a hypothetical but realistic case of data plotted on the 2DHER-field where the phenotypic correlation is $r_{P_{x,y}} = -0.5$, the coheritability $h_{x,y} = -0.35$, and the coenvironmentability $e_{x,y} = -0.15$. Further examples are presented in Figure 4.2.3-2 where, for ease of comparison, the density curves are overlaid in the same graph.

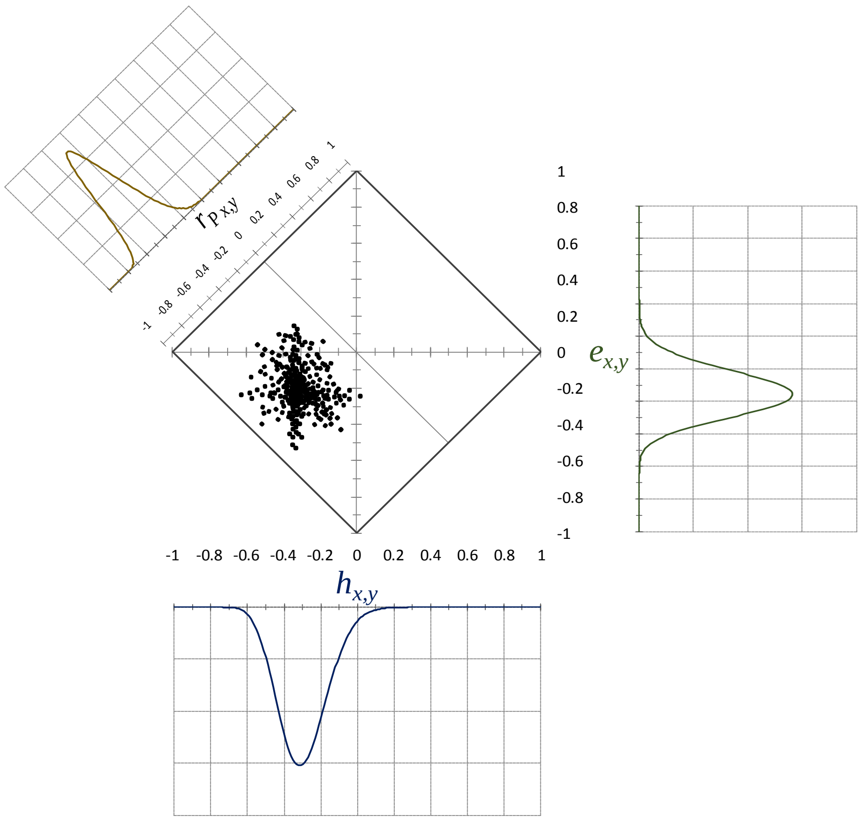

**Figure 4.2.3-1** Sampling distribution of phenotypic correlation, coheritability, and coenvironmnetability. Data is plotted on the 2DHER-field showing the phenotypic correlation between two traits obtained from $n = 150$ samples drawn from a population with $\rho_{P_{x,y}}$ =-0.51, decomposed into its coheritability, coenvironmentability components

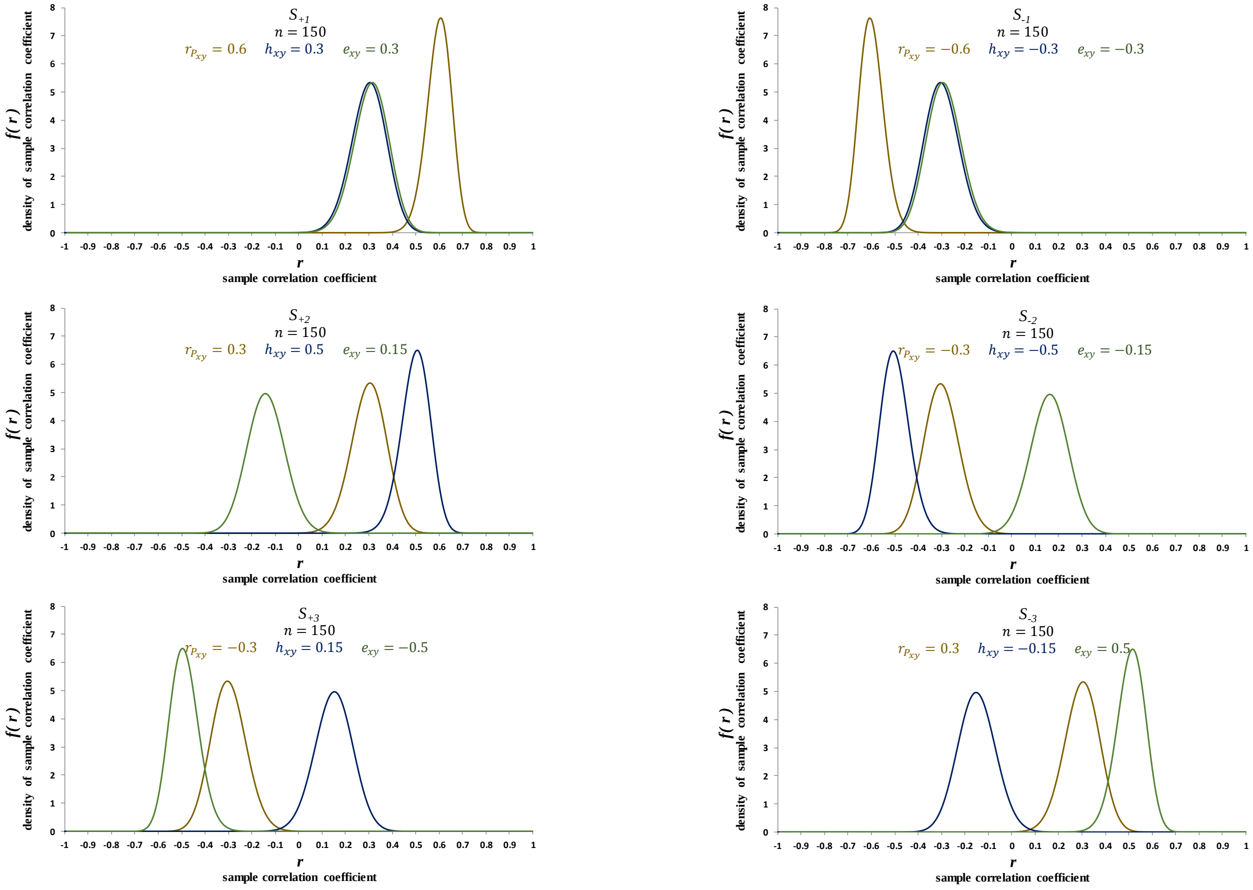

**Figure 4.2.3-2** Example of distributions of the phenotypic correlation, coheritability and coenvironmentability. Density curves are overlaid to facilitate comparison.

Pham-Gia and Choulakian (2014) proposed an original expression for the density of the correlation matrix, with sample variances as parameters for the case of the multivariate normal population with non-identity population correlation.

#### 4.3 Moments of the sample correlation coefficient distribution

Hotelling (1953, page 212) presents the moments of the sample correlation coefficient $r$ around its mean. Rady et al. (2005) introduced a method to easily determine the moments of $r$ around its mean, which are the same as those found by Gosh (1966). Romero-Padilla (2016) presents the moments of $r$ around the origin using hypergeometric functions.

Mean sample correlation

**4-3a** Mean of the sample correlation coefficient around its mean

$$E(r) = \bar{r} = \rho\left\{1 - \frac{1}{2(n-1)}(1-\rho^2) - \frac{3}{8(n-1)^2}(1-\rho^2)(1+3\rho^2)\right\}$$

**4-3b** Variance of the sample correlation coefficient

$$Var(r) = \sigma_r^2 = \frac{(1-\rho^2)^2}{n-1}\left\{1 + \frac{5.5\,\rho^2}{n-1} + \frac{-24\rho^2+75\rho^4}{2(n-1)^2} +. . .\right\}$$

**4-3c** Skewness of the sample correlation coefficient

$$skewness = \frac{6\rho}{\sqrt{n-1}}\, 1 + \frac{-30+77\rho^2}{12\,(n-1)} +. . .$$

**4-3c** Kurtosis of the sample correlation coefficient

$$kurtosis = \frac{6(12\rho^2-1)}{\sqrt{n-1}} +. . .$$

#### 4.4 The Fisher $r$-to-$Z$ transform of the correlation coefficient

The Pearson product moment correlation coefficient $r$ based on $n$ independent units of observation has an asymptotic normal distribution. However, its distribution can be highly skewed at small-to-moderate sample sizes, and its sampling variance varies as a function of the population correlation coefficient $\rho$. Noting these limitations, Fisher (1921) proposed using the inverse hyperbolic tangent function $Z_r$ as a normalizing and variance-stabilizing transformation:

**4-4a** Fisher r-to-Z transformation of $Z_r$

$$Z_r = atanh(r) = \frac{1}{2} \ln\left(\frac{1+r}{1-r}\right)$$

note the equivalent ways to denote the inverse hyperbolic tangent:

$$atanh(r) = artanh(r) = arctanh(r) = tanh^{-1}(r)$$

For the transformed $z$, the approximate variance $\sigma^2_{Z_r} = 1/(n-3)$ is independent of the correlation. Fouladi and Steiger (2008) provide a thorough discussion on the consequence of the use of this transformation and its implications in inferential statistics.

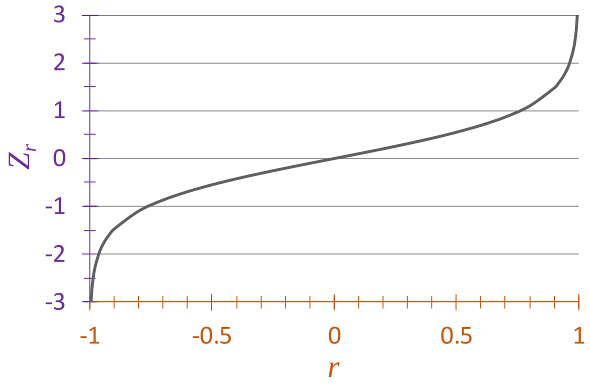

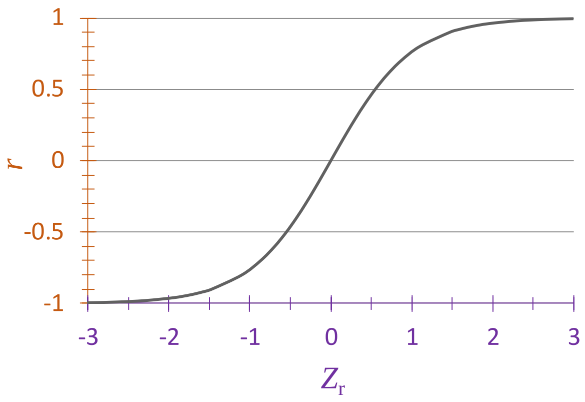

4-4b Mean of the Fisher transform of $Z_r$

$$\bar{Z}_r = E\left(Z_r - Z_\rho\right)$$

$$= \frac{\rho}{2(n-1)} + \frac{5\rho + \rho^3}{8(n-1)^2} + \frac{11\rho + 2\rho^3 + 3\rho^5}{16(n-1)^3} + \frac{83\rho + 13\rho^3 - 27\rho^5 + 75\rho^7}{128(n-1)^4} + O(n^{-5})$$

4-4c Variance of the Fisher transform of $Z_r$

$$\sigma^2_{Z_r - Z_\rho} = E(Z_r - Z_\rho - \bar{r})^2$$

$$= \frac{1}{n-1} + \frac{8 - \rho^2}{4(n-1)^2} + \frac{88 - 9\rho^2 - 9\rho^4}{24(n-1)^3} + \frac{384 - 19\rho^2 + 2\rho^4 - 75\rho^6}{64(n-1)^4} + O(n^{-5})$$

**Figure 4.4**. Distribution of the sample correlation coefficient $r$ (orange line), and its corresponding Fisher transform $Z_r$ (purple line).

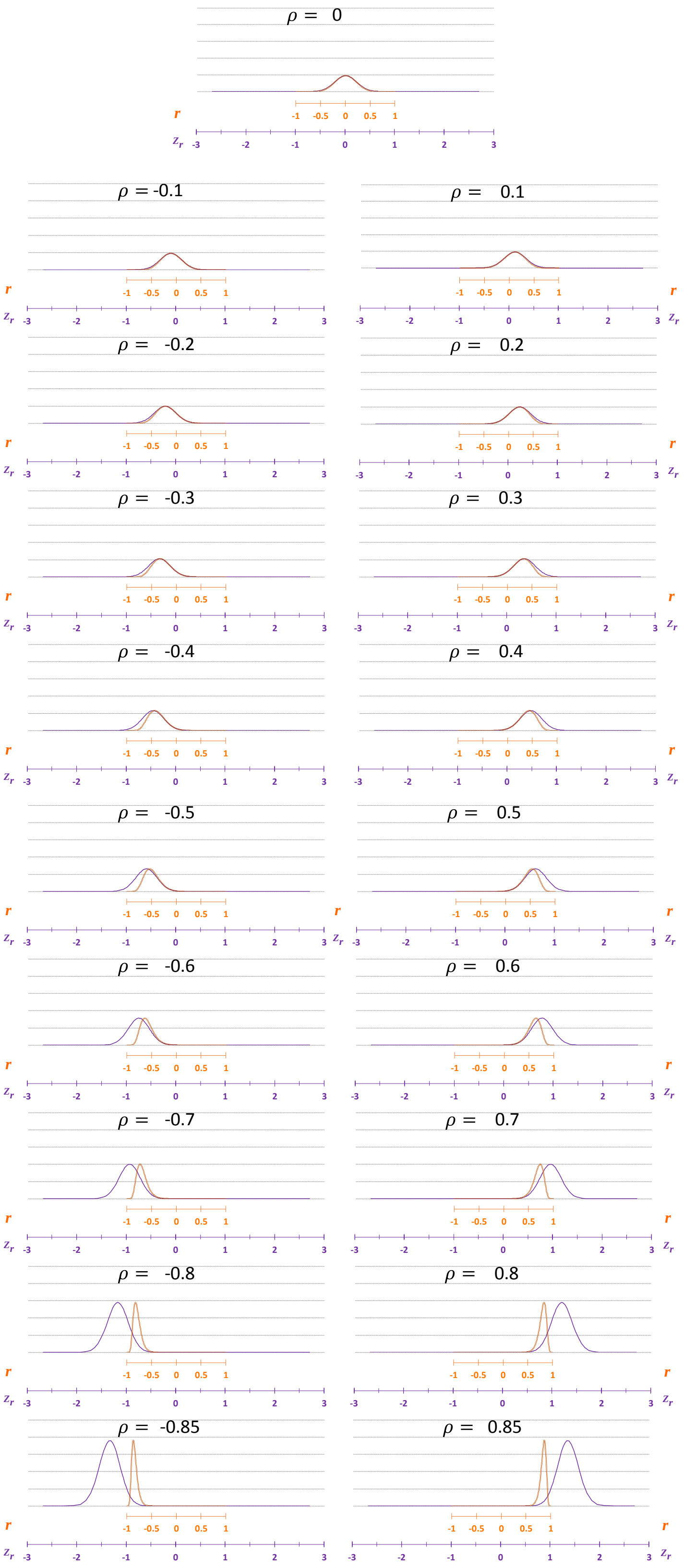

### 5 Inferential statistics

Statistical inference is often conducted through significance testing and confidence interval construction. Although closely related, significance testing focuses on a single *a priori* hypothesis, usually a null value (e.g. $\rho = 0$). In contrast, a confidence interval can avoid such problems by providing a range of plausible parameter values. The confidence interval reveals both the magnitude and the precision of the estimated effect, whereas the p-value obtained from significance testing confounds these two aspects of the data (Lee 2016).

#### 5.1 Statistical justification

In this section I apply well-established statistical tools for the analysis of correlations and extend them to the analysis of the coheritability coefficient. The heritabilities are treated here as scalars, which may be considered a potential caveat of the method.

The distribution of a random variable multiplied by a scalar creates a scale family distribution, with well-known statistical properties. Thus, the coheritability $h_{xy}$ has a distribution similar to the one of $r_{A_{x,y}}$, but rescaled by $\sqrt{h_x^2\, h_y^2}$ .

(1) It has been demonstrated that at limiting cases (Section 7.4) the phenotypic correlation can equate either the coheritability (if both heritabilities are unity) or the coenvironmentability (if at least one heritability is zero). This fact indicates that the same inferential statistical tools to test correlations can also be used to test the coheritability and the coenvironmentability.

(2) The coheritability is composed of additive genetic correlation multiplied by the factor $\sqrt{h_x^2\, h_y^2}$ which is the geometric mean of the heritabilities. Although, the heritabilities are random variables, here they are going to be regarded as constants (i.e. scalars). Thus, the coheritability coefficient $h_{xy} = \sqrt{h_x^2\, h_y^2}\, r_{A_{x,y}}$ can be thought of as the weighed genetic correlation, with a distribution that is a scale family of the original correlation distribution.

The $\sqrt{h_x^2\, h_x^2}$ factor is expressed as a strictly positive decimal numeral bound by the interval (0 , +1]. This causes the values of $h_{x,y}$ to compare to $r_{A_{x,y}}$ as follows:

Multiplying $\sqrt{h_x^2\, h_x^2}$ to the genetic correlation $r_{A_{x,y}}$ will cause (a) the modulus (i.e. absolute value) of the product to decrease, but still maintaining the same sign.

(a) if $-1 \leq r_{A_{x,y}} < 0$ then the value of the coheritability would oscillate in the range $r_{A_{x,y}} \leq h_{x,y} < 0$, because multiplying a decimal numeral to a negative $r_{A_{x,y}}$ will cause its magnitude to move towards zero, making $h_{x,y}$ comparatively higher than$r_{A_{x,y}}$, yet keeping its sign.

(b) If $0 < r_{A_{xy}} \leq 1$ then $0 < h_{xy} \leq r_{A_{xy}}$, because multiplying a positive decimal numeral to a strictly positive $r_{A_{xy}}$ will cause its magnitude to move towards zero thus decreasing it, and maintain its sign.

(3) The form of the $r_{A_{xy}}$ sampling distribution does not change with the factoring $\sqrt{h_x^2 \, h_y^2}$. It simply becomes a scale family distribution of $r_{A_{xy}}$.

(4) The $r$-to-$Z$ Fisher transformed of the phenotypic correlation cannot be decomposed into a coheritability term and a coenvironmentability term. However, a naïve and heuristic approach could be assumed in which the proportion of the coheritability or coenvironmentability with respect to the phenotypic correlation in the $r$-space does not change with the transformation to the $Z$-space.

#### 5.2 Hypothesis testing when null hypothesis sets the parameters to a zero value

Although the fundamental results and associated use of the sample correlation are described in most introductory textbooks of statistics. The rationale used for the testing of the correlation coefficients (phenotypic, genetic, environmental) $\rho = 0$, is extended, in a heuristic way, to the hypothesis test of the coheritability $H_{x,y} = 0$, and the coenvironmentability $E_{x,y} = 0$. Here I used the term parameter to refer to $\theta = \{\rho_{P_{x,y}}, \rho_{A_{x,y}}, \rho_{E_{x,y}}, H_{x,y}, E_{x,y}\}$.

The sample correlation coefficient $r$ is a point estimator of $\rho$. The distribution of r when $\rho = 0$ is presented in Section 4.1.1. To test whether or not a linear relationship exists between two traits $x$ and $y$, a following test statistic is used

$$t_{\alpha,(n-2)} = \frac{r}{\sqrt{\frac{1-r^2}{n-2}}}$$

Which can also be rearranged as

$$r = \frac{t_{\alpha,(n-2)}}{\sqrt{n-2+t^2}}$$

(see Samiuddin 1970 for an exact test statistic).

Which is the basis of the critical value method, used in this work.

$$r_{critical} = \frac{t_{\alpha,(n-2)_{critical}}}{\sqrt{n-2+t^2}}$$

There is extensive criticism in the literature of testing of parameters hypothesized to be equal to zero (Beaulieu-Prévost 2006, Lee 2016). Simply reporting that a correlation is significant at a given $\alpha$ level just because the p-value for the test $Ho\colon \rho = 0$ is less than $\alpha$ is generally not sufficient (Looney 2008). The argument that $Ho\colon \rho = 0$ is the appropriate null hypothesis to test and that using sample sizes that actually yield a desirable level of power (say 80%) for the test can result in confidence intervals that are so wide that they provide very little useful information about the magnitude of the population correlation coefficient.

A better approach is to test null values other than $\rho_o = 0$ (Section 5.3), and then determine the sample size (Section 5.6) so as to achieve a certain level of precision of the estimate of $\rho$ as measured by the width of the resulting confidence interval.

### Hypothesis test of population phenotypic correlation

hypotheses

$H_o\!: \ \rho_{P_{x,y}} = 0$

$H_1: \ \rho_{P_{x,y}} < 0$

significance level

$\alpha =$ 0.05

critical value

$r_{P_{crit}} = \dfrac{t_{crit}}{\sqrt{n-2+t_{crit}^2}} =$ -0.1646

$n =$ 101 $\quad \alpha =$ 0.05

$t_{crit} = -t_{\alpha,n-2} =$ -1.6604

test statistic

$r_{P_{x,y}} =$ 0.45

test criteria

$if \begin{cases} r_{P\,x,y} \le r_{P_{crit}} & \text{then reject the } H_o \\ otherwise & \text{do not reject } H_o \end{cases}$

test result

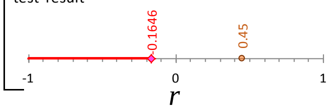

test conclusion

Because $r_{P_{x,y}} < r_{P_{crit}}$

The test rejects the $H_o$ at an
α = 0.05 significance level

hypotheses

$H_o\!: \ \rho_{P_{x,y}} = 0$

$H_1: \ \rho_{P_{x,y}} \neq 0$

significance level

$\alpha =$ 0.05

critical value

$r_{P_{crit}} = \dfrac{t_{crit}}{\sqrt{n-2+t_{crit}^2}} = \pm$ 0.1956

$n =$ 101 $\quad \alpha =$ 0.05

$|t_{crit}| \to \pm t_{1-\frac{\alpha}{2},n-2} = \pm$1.9842

test statistic

$r_{P_{x,y}} =$ 0.45

test criteria

$if \begin{cases} |r_{P\,x,y}| \ge r_{P_{crit}} & \text{then reject the } H_o \\ otherwise & \text{do not reject } H_o \end{cases}$

test result

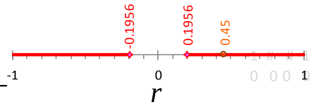

test conclusion

Because $|r_{P_{x,y}}| > r_{P_{crit}}$

The test rejects the $H_o$ at an
α = 0.05 significance level

hypotheses

$H_o\!: \ \rho_{P_{x,y}} = 0$

$H_1: \ \rho_{P_{x,y}} > 0$

significance level

$\alpha =$ 0.05

critical value

$r_{P_{crit}} = \dfrac{t_{crit}}{\sqrt{n-2+t_{crit}^2}} =$ 0.1646

$n =$ 101 $\quad \alpha =$ 0.05

$t_{crit} = t_{\alpha,n-2} =$ 1.6604

test statistic

$r_{P_{x,y}} =$ 0.45

test criteria

$if \begin{cases} r_{P\,x,y} \ge r_{P_{crit}} & \text{then reject the } H_o \\ otherwise & \text{do not reject } H_o \end{cases}$

test result

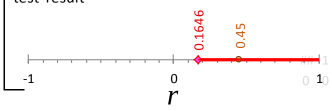

test conclusion

Because $r_{P_{x,y}} < r_{P_{crit}}$

The test does not rejects the $H_o$ at an
α = 0.05 significance level

### Hypothesis test of population genetic correlation

**hypotheses**

$H_o\text{: } \rho_{A_{x,y}} = 0$

$H_1 : \rho_{A_{x,y}} < 0$

**significance level**

$\alpha =$ 0.05

**critical value**

$r_{A_{crit}} = \dfrac{t_{crit}}{\sqrt{n-2+t_{crit}^2}} =$ -0.1646

$n =$ 101 $\alpha =$ 0.05

$t_{crit} = -t_{\alpha,n-2} =$ -1.6604

**test statistic**

$r_{A_{x,y}} =$ -0.8165

**test criteria**

$if \begin{cases} r_{A_{x,y}} \le r_{A_{crit}} \\ otherwise \end{cases}$

then reject the $H_o$

do not reject $H_o$

**test result**

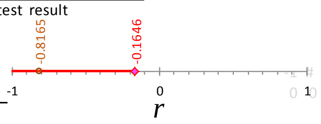

**test conclusion**

Because $r_{A_{x,y}} < r_{A_{crit}}$

The test rejects the $H_o$ at an
α = 0.05 significance level

**hypotheses**

$H_o\text{: } \rho_{A_{x,y}} = 0$

$H_1 : \rho_{A_{x,y}} \ne 0$

**significance level**

$\alpha =$ 0.05

**critical value**

$r_{A_{crit}} = \dfrac{t_{crit}}{\sqrt{n-2+t_{crit}^2}} = \pm$ 0.1956

$n =$ 101 $\alpha =$ 0.05

$|t_{crit}| \rightarrow \pm t_{1-\frac{\alpha}{2},n-2} = \pm$ 1.9842

**test statistic**

$r_{A_{x,y}} =$ -0.8165

**test criteria**

$if \begin{cases} |r_{A_{x,y}}| \ge r_{A_{crit}} \\ otherwise \end{cases}$

then reject the $H_o$

do not reject $H_o$

**test result**

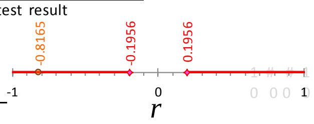

**test conclusion**

Because $|r_{A_{x,y}}| > r_{A_{crit}}$

The test rejects the $H_o$ at an
α = 0.05 significance level

**hypotheses**

$H_o\text{: } \rho_{A_{x,y}} = 0$

$H_1 : \rho_{A_{x,y}} > 0$

**significance level**

$\alpha =$ 0.05

**critical value**

$r_{A_{crit}} = \dfrac{t_{crit}}{\sqrt{n-2+t_{crit}^2}} =$ 0.1646

$n =$ 101 $\alpha =$ 0.05

$t_{crit} = t_{\alpha,n-2} =$ 1.6604

**test statistic**

$r_{A_{x,y}} =$ -0.8165

**test criteria**

$if \begin{cases} r_{A_{x,y}} \ge r_{A_{crit}} \\ otherwise \end{cases}$

then reject the $H_o$

do not reject $H_o$

**test result**

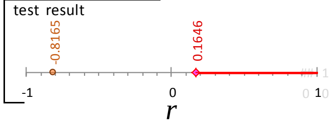

**test conclusion**

Because $r_{A_{x,y}} < r_{A_{crit}}$

The test does not rejects the $H_o$ at an
α = 0.05 significance level

#### Hypothesis test of population environmental correlation

**hypotheses**

$H_o\colon\ \rho_{E_{x,y}} = 0$

$H_1:\ \rho_{E_{x,y}} < 0$

**significance level**

$\alpha =$ 0.05

**critical value**

$r_{E_{crit}} = \dfrac{t_{crit}}{\sqrt{n-2+t_{crit}^2}} =$ -0.1646

$n =$ 101 $\alpha =$ 0.05

$t_{crit} = -t_{\alpha,n-2} =$ -1.6604

**test statistic**

$r_{E_{x,y}} =$ 0.8686

**test criteria**

$if \begin{cases} r_{E_{x,y}} \le r_{E_{crit}} & \text{then reject the } H_o \\ otherwise & \text{do not reject } H_o \end{cases}$

**test result**

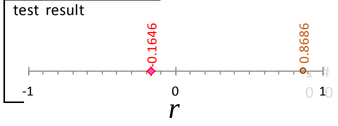

**test conclusion**

Because $r_{E_{x,y}} < r_{E_{crit}}$

The test does not reject the $H_o$ at an

α = 0.05 significance level

---

**hypotheses**

$H_o\colon\ \rho_{E_{x,y}} = 0$

$H_1:\ \rho_{E_{x,y}} \ne 0$

**significance level**

$\alpha =$ 0.05

**critical value**

$r_{E_{crit}} = \dfrac{t_{crit}}{\sqrt{n-2+t_{crit}^2}} = \pm$ 0.1956

$n =$ 101 $\alpha =$ 0.05

$|t_{crit}| \to \pm t_{1-\frac{\alpha}{2},n-2} = \pm$ 1.9842

**test statistic**

$r_{E_{x,y}} =$ 0.8686

**test criteria**

$if \begin{cases} |r_{E_{x,y}}| \ge r_{E_{crit}} & \text{then reject the } H_o \\ otherwise & \text{do not reject } H_o \end{cases}$

**test result**

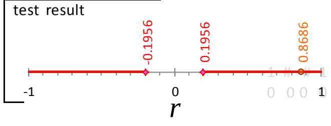

**test conclusion**

Because $|r_{E_{x,y}}| > r_{E_{crit}}$

The test rejects the $H_o$ at an

α = 0.05 significance level

---

**hypotheses**

$H_o\colon\ \rho_{E_{x,y}} = 0$

$H_1:\ \rho_{E_{x,y}} > 0$

**significance level**

$\alpha =$ 0.05

**critical value**

$r_{E_{crit}} = \dfrac{t_{crit}}{\sqrt{n-2+t_{crit}^2}} =$ 0.1646

$n =$ 101 $\alpha =$ 0.05

$t_{crit} = t_{\alpha,n-2} =$ 1.6604

**test statistic**

$r_{E_{x,y}} =$ 0.8686

**test criteria**

$if \begin{cases} r_{E_{x,y}} \ge r_{E_{crit}} & \text{then reject the } H_o \\ otherwise & \text{do not reject } H_o \end{cases}$

**test result**

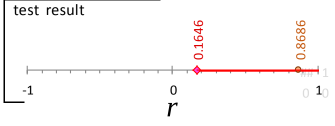

**test conclusion**

Because $r_{E_{x,y}} > r_{E_{crit}}$

The test rejects the $H_o$ at an

α = 0.05 significance level

### Hypothesis test of population coheritability

hypotheses

$H_o$: $H_{x,y} = 0$

$H_1$: $H_{x,y} < 0$

significance level

$\alpha =$ 0.05

critical value

$h_{x,y_{crit}} = \sqrt{h_x^2\, h_y^2}\; r_{A_{x,y_{crit}}} = \dfrac{\sqrt{h_x^2\, h_y^2}\; t_{crit}}{\sqrt{n-2+t_{crit}^2}} =$ -0.0403

$n =$ 101 $\alpha =$ 0.05

$t_{crit} = -t_{\alpha, n-2} =$ -1.6604

test statistic

$h_{x,y} =$ -0.2

test criteria

$if \begin{cases} h_{x,y} \le h_{x,y_{crit}} & \text{then reject the } H_0 \\ otherwise & \text{do not reject } H_0 \end{cases}$

test result

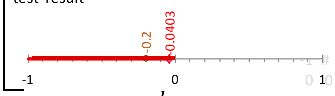

$h_{x,y}$

test conclusion

Because $h_{x,y} < h_{x,y_{crit}}$

The test rejects the $H_0$ at an

α = 0.05 significance level

hypotheses

$H_o$: $H_{x,y} = 0$

$H_1$: $H_{x,y} \neq 0$

significance level

$\alpha =$ 0.05

critical value

$h_{x,y_{crit}} = \sqrt{h_x^2\, h_y^2}\; r_{A_{x,y_{crit}}} = \dfrac{\sqrt{h_x^2\, h_y^2}\; t_{crit}}{\sqrt{n-2+t_{crit}^2}} = \pm$ 0.0479

$n =$ 101 $\alpha =$ 0.05

$|t_{crit}| \to \pm t_{1-\frac{\alpha}{2}, n-2} = \pm$ 1.9842

test statistic

$h_{x,y} =$ -0.2

test criteria

$if \begin{cases} |h_{x,y}| \ge h_{x,y_{crit}} & \text{then reject the } H_0 \\ otherwise & \text{do not reject } H_0 \end{cases}$

test result

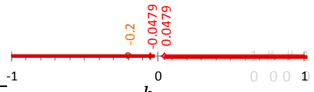

$h_{x,y}$

test conclusion

Because $|h_{x,y}| > h_{x,y_{crit}}$

The test rejects the $H_0$ at an

α = 0.05 significance level

hypotheses

$H_o$: $H_{x,y} = 0$

$H_1$: $H_{x,y} > 0$

significance level

$\alpha =$ 0.05

critical value

$h_{x,y_{crit}} = \sqrt{h_x^2\, h_y^2}\; r_{A_{x,y_{crit}}} = \dfrac{\sqrt{h_x^2\, h_y^2}\; t_{crit}}{\sqrt{n-2+t_{crit}^2}} =$ 0.0403

$n =$ 101 $\alpha =$ 0.05

$t_{crit} = t_{\alpha, n-2} =$ 1.6604

test statistic

$h_{x,y} =$ -0.2

test criteria

$if \begin{cases} h_{x,y} \ge h_{x,y_{crit}} & \text{then reject the } H_0 \\ otherwise & \text{do not reject } H_0 \end{cases}$

test result

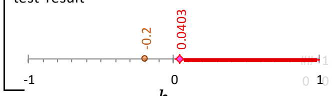

$h_{x,y}$

test conclusion

Because $h_{x,y} < h_{x,y_{crit}}$

The test does not rejects the $H_0$ at an

α = 0.05 significance level

### Hypothesis test of population coenvironmentability

hypotheses

$H_o\!: \; E_{x,y} = 0$

$H_1: \; E_{x,y} < 0$

significance level

$\alpha =$ 0.05

critical value

$$e_{x,y_{crit}} = \sqrt{(1-h_x^2)(1-h_y^2)}\; r_{E_{x,y_{crit}}} = \frac{\sqrt{(1-h_x^2)(1-h_y^2)}\; t_{crit}}{\sqrt{n-2+t_{crit}^2}}$$

$=$ -0.1232

$n =$ 101 $\alpha =$ 0.05

$t_{crit} = -t_{\alpha,n-2} =$ -1.6604

test statistic

$e_{x,y} =$ 0.65

test criteria

$if \begin{cases} e_{x,y} \le e_{x,y_{crit}} & \text{then reject the } H_o \\ otherwise & \text{do not reject } H_o \end{cases}$

test result

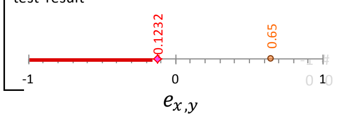

test conclusion

Because $r_{E_{x,y}} < r_{E_{crit}}$

The test does not reject the $H_o$ at an $\alpha =$ 0.05 significance level

hypotheses

$H_o\!: \; E_{x,y} = 0$

$H_1: \; E_{x,y} \neq 0$

significance level

$\alpha =$ 0.05

critical value

$$e_{x,y_{crit}} = \sqrt{(1-h_x^2)(1-h_y^2)}\; r_{E_{x,y_{crit}}} = \frac{\sqrt{(1-h_x^2)(1-h_y^2)}\; t_{crit}}{\sqrt{n-2+t_{crit}^2}}$$

$= \pm$ 0.1464

$n =$ 101 $\alpha =$ 0.05

$|t_{crit}| \rightarrow \pm t_{1-\frac{\alpha}{2},n-2} = \pm$ 1.9842

test statistic

$e_{x,y} =$ 0.65

test criteria

$if \begin{cases} |e_{x,y}| \ge e_{x,y_{crit}} & \text{then reject the } H_o \\ otherwise & \text{do not reject } H_o \end{cases}$

test result

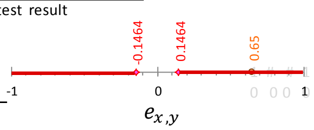

test conclusion

Because $|r_{E_{x,y}}| > r_{E_{crit}}$

The test rejects the $H_o$ at an $\alpha =$ 0.05 significance level

hypotheses

$H_o\!: \; E_{x,y} = 0$

$H_1: \; E_{x,y} > 0$

significance level

$\alpha =$ 0.05

critical value

$$e_{x,y_{crit}} = \sqrt{(1-h_x^2)(1-h_y^2)}\; r_{E_{x,y_{crit}}} = \frac{\sqrt{(1-h_x^2)(1-h_y^2)}\; t_{crit}}{\sqrt{n-2+t_{crit}^2}}$$

$=$ 0.1232

$n =$ 101 $\alpha =$ 0.05

$t_{crit} = t_{\alpha,n-2} =$ 1.6604

test statistic

$e_{x,y} =$ 0.65

test criteria

$if \begin{cases} e_{x,y} \ge e_{x,y_{crit}} & \text{then reject the } H_o \\ otherwise & \text{do not reject } H_o \end{cases}$

test result

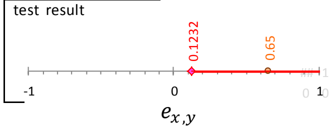

test conclusion

Because $r_{E_{x,y}} > r_{E_{crit}}$

The test rejects the $H_o$ at an $\alpha =$ 0.05 significance level

#### 5.3 Hypothesis testing for a parameter equal to a nonzero value

To test the significance of point estimators of the population parameters $\theta = \{\rho_{P_{x,y}}, \rho_{A_{x,y}}, \rho_{E_{x,y}}, H_{x,y}, E_{x,y}\}$, the hypothesized values used in $H_o$ must satisfy the following relationship:

$$\rho_{P_{x,y}} = \underbrace{\sqrt{h_x^2\, h_y^2}\ \rho_{A_{x,y}}}_{H_{x,y}} + \underbrace{\sqrt{(1-h_x^2)(1-\ h_y^2)}\ \rho_{E_{x,y}}}_{E_{x,y}}$$

Since the sampling distribution of the Pearson's $r$ is not normally distributed, a common practice is to use the Fisher's $r$-to-$Z$ transformation (Section 4.4).

### Hypothesis test of population phenotypic correlation

hypotheses

$H_o$: $\rho_{P_{x,y}} =$ 0.6

$H_1$: $\rho_{P_{x,y}} <$ 0.6

significance level

$\alpha =$ 0.05

Fisher $r$-to-$Z$ transformation

$$Z_{r_P} = \frac{1}{2} ln\left[\frac{1+(\ 0.45\ )}{1-(\ 0.45\ )}\right] = 0.4847$$

$$Z_{\rho_P} = \frac{1}{2} ln\left[\frac{1+(\ 0.6\ )}{1-(\ 0.6\ )}\right] = 0.6931$$

$$\sigma_{Z_{r_P}} \approx \sqrt{\frac{1}{n-3}} = 0.1010$$

critical value

$z_{crit} = z_\alpha =$ -1.645

test statistic

$$z = \frac{z_{r_P} - z_{\rho_P}}{\sigma_{Z_{r_P}}} = \frac{(0.4847) - (0.69315)}{0.1010} = -2.06$$

test criteria

$if \begin{cases} z \le z_{P_{crit}} & \text{then reject the H}_o \\ otherwise & \text{do not reject H}_o \end{cases}$

test result

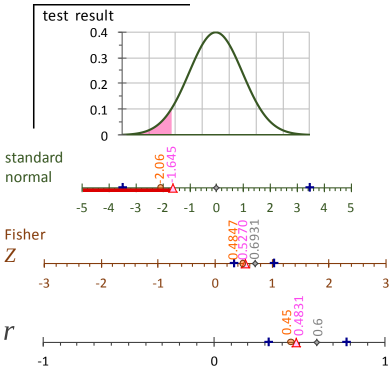

test conclusion

Because $Z < z_{P_{crit}}$

The test rejects the $H_o$ at an

$\alpha =$ 0.05 significance level

hypotheses

$H_o$: $\rho_{P_{x,y}} =$ 0.6

$H_1$: $\rho_{P_{x,y}} \neq$ 0.6

significance level

$\alpha =$ 0.05

Fisher $r$-to-$Z$ transformation

$$Z_{r_P} = \frac{1}{2} ln\left[\frac{1+(\ 0.45\ )}{1-(\ 0.45\ )}\right] = 0.4847$$

$$Z_{\rho_P} = \frac{1}{2} ln\left[\frac{1+(\ 0.6\ )}{1-(\ 0.6\ )}\right] = 0.6931$$

$$\sigma_{Z_{r_P}} \approx \sqrt{\frac{1}{n-3}} = 0.1010$$

critical value

$z_{crit} = \pm z_{1-\frac{\alpha}{2}} = \pm 1.96$

test statistic

$$z = \frac{z_{r_P} - z_{\rho_P}}{\sigma_{Z_{r_P}}} = \frac{(0.4847) - (0.69315)}{0.1010} = -2.06$$

test criteria

$if \begin{cases} |z| \ge z_{P_{crit}} & \text{then reject the H}_o \\ |z| < z_{crit} & \text{do not reject H}_o \end{cases}$

test result

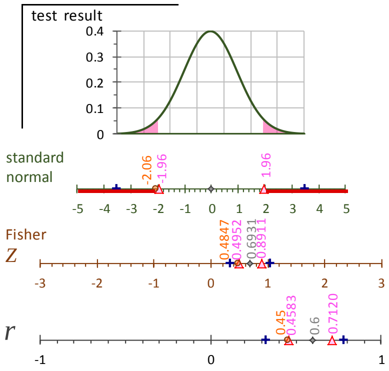

test conclusion

Because $Z < -z_{P_{crit}}$

The test rejects the $H_o$ at an

$\alpha =$ 0.05 significance level

hypotheses

$H_o$: $\rho_{P_{x,y}} =$ 0.6

$H_1$: $\rho_{P_{x,y}} >$ 0.6

significance level

$\alpha =$ 0.05

Fisher $r$-to-$Z$ transformation

$$Z_{r_P} = \frac{1}{2} ln\left[\frac{1+(\ 0.45\ )}{1-(\ 0.45\ )}\right] = 0.4847$$

$$Z_{\rho_P} = \frac{1}{2} ln\left[\frac{1+(\ 0.6\ )}{1-(\ 0.6\ )}\right] = 0.6931$$

$$\sigma_{Z_{r_P}} \approx \sqrt{\frac{1}{n-3}} = 0.1010$$

critical value

$z_{P_{crit}} = z_{1-\frac{\alpha}{2}} = 1.645$

test statistic

$$z = \frac{z_{r_P} - z_{\rho_P}}{\sigma_{Z_{r_P}}} = \frac{(0.4847) - (0.69315)}{0.1010} = -2.06$$

test criteria

$if \begin{cases} z \ge z_{P_{crit}} & \text{then reject the H}_o \\ otherwise & \text{do not reject H}_o \end{cases}$

test result

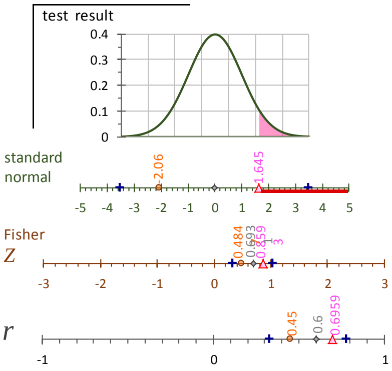

test conclusion

Because $Z < z_{P_{crit}}$

The test does not reject the $H_o$ at an

$\alpha =$ 0.05 significance level

### Hypothesis test of population genetic correlation

hypotheses

$H_o$: $\rho_{A_{x,y}} =$ -0.7071

$H_1$: $\rho_{A_{x,y}} <$ -0.7071

significance level

$\alpha = 0.05$

Fisher $r$-to-$Z$ transformation

$$Z_{r_A} = \frac{1}{2} ln\left[\frac{1+(\text{-0.8165})}{1-(\text{-0.8165})}\right] \quad \text{-1.1462}$$

$$Z_{\rho_A} = \frac{1}{2} ln\left[\frac{1+(\text{-0.7071})}{1-(\text{-0.7071})}\right] \quad \text{-0.8814}$$

$$\sigma_{Z_{r_A}} \approx \sqrt{\frac{1}{n-3}} = 0.1010$$

critical value

$$z_{A_{crit}} = z_\alpha = \text{-1.645}$$

test statistic

$$z = \frac{z_{r_A} - z_{\rho_A}}{\sigma_{Z_{r_A}}} = \frac{(\text{-1.1462}) - (\text{-0.8814})}{0.1010} = \text{-2.62}$$

test criteria

$if \begin{cases} z \le z_{A_{crit}} & \text{then reject the } H_0 \\ otherwise & \text{do not reject } H_0 \end{cases}$

test result

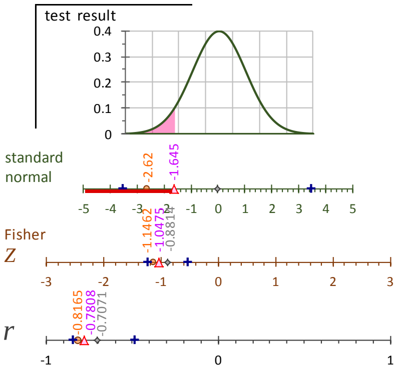

test conclusion

Because $z < z_{A_{crit}}$

The test rejects the $H_0$ at an α = 0.05 significance level

hypotheses

$H_o$: $\rho_{A_{x,y}} =$ -0.7071

$H_1$: $\rho_{A_{x,y}} \neq$ -0.7071

significance level

$\alpha = 0.05$

Fisher $r$-to-$Z$ transformation

$$Z_{r_A} = \frac{1}{2} ln\left[\frac{1+(\text{-0.8165})}{1-(\text{-0.8165})}\right] \quad \text{-1.1462}$$

$$Z_{\rho_A} = \frac{1}{2} ln\left[\frac{1+(\text{-0.7071})}{1-(\text{-0.7071})}\right] \quad \text{-0.8814}$$

$$\sigma_{Z_{r_A}} \approx \sqrt{\frac{1}{n-3}} = 0.1010$$

critical value

$$z_{A_{crit}} = \pm z_{1-\frac{\alpha}{2}} = \pm 1.96$$

test statistic

$$z = \frac{z_{r_A} - z_{\rho_A}}{\sigma_{Z_{r_A}}} = \frac{(\text{-1.1462}) - (\text{-0.8814})}{0.1010} = \text{-2.62}$$

test criteria

$if \begin{cases} |z| \ge z_{A_{crit}} & \text{then reject the } H_0 \\ |z| < z_{crit} & \text{do not reject } H_0 \end{cases}$

test result

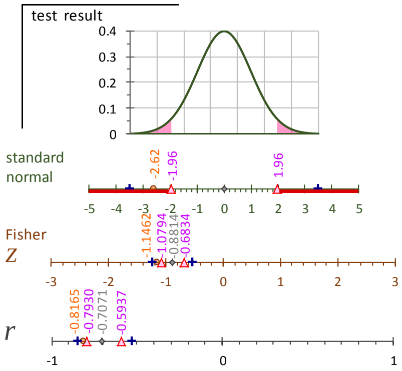

test conclusion

Because $z < -z_{A_{crit}}$

The test rejects the $H_0$ at an α = 0.05 significance level

hypotheses

$H_o$: $\rho_{A_{x,y}} =$ -0.7071

$H_1$: $\rho_{A_{x,y}} >$ -0.7071

significance level

$\alpha = 0.05$

Fisher $r$-to-$Z$ transformation

$$Z_{r_A} = \frac{1}{2} ln\left[\frac{1+(\text{-0.8165})}{1-(\text{-0.8165})}\right] \quad \text{-1.1462}$$

$$Z_{\rho_A} = \frac{1}{2} ln\left[\frac{1+(\text{-0.7071})}{1-(\text{-0.7071})}\right] \quad \text{-0.8814}$$

$$\sigma_{Z_{r_A}} \approx \sqrt{\frac{1}{n-3}} = 0.1010$$

critical value

$$z_{A_{crit}} = z_{1-\frac{\alpha}{2}} = 1.645$$

test statistic

$$z = \frac{z_{r_A} - z_{\rho_A}}{\sigma_{Z_{r_A}}} = \frac{(\text{-1.1462}) - (\text{-0.8814})}{0.1010} = \text{-2.62}$$

test criteria

$if \begin{cases} z \ge z_{A_{crit}} & \text{then reject the } H_0 \\ otherwise & \text{do not reject } H_0 \end{cases}$

test result

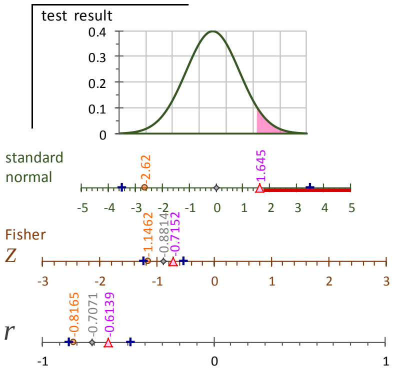

test conclusion

Because $z < z_{A_{crit}}$

The test does not reject the $H_0$ at an α = 0.05 significance level

### Hypothesis test of population environmental correlation

hypotheses $H_o\colon\ \rho_{A_{x,y}} =$ 0.8250

$H_1\colon\ \rho_{A_{x,y}} <$ 0.8250

significance level

$\alpha = 0.05$

Fisher $r$-to-$Z$ transformation

$$Z_{r_E} = \frac{1}{2} ln\left[\frac{1 + (\ 0.8686\ )}{1 - (0.8686\ )}\right] = 1.3273$$

$$Z_{\rho_E} = \frac{1}{2} ln\left[\frac{1 + (\ 0.8250\ )}{1 - (0.8250\ )}\right] = 1.1721$$

$$\sigma_{Z_{r_E}} \approx \sqrt{\frac{1}{n-3}} = 0.1010$$

critical value

$z_{crit} = z_\alpha =$ -1.645

test statistic

$$z = \frac{z_{r_E} - z_{\rho_E}}{\sigma_{Z_{r_E}}} = \frac{(\text{-}1.1462) - (\text{-}0.8814)}{0.1010} = 1.54$$

test criteria

$if \begin{cases} z \le z_{E_{crit}} \\ otherwise \end{cases}$ then reject the H₀ / do not reject H₀

test result

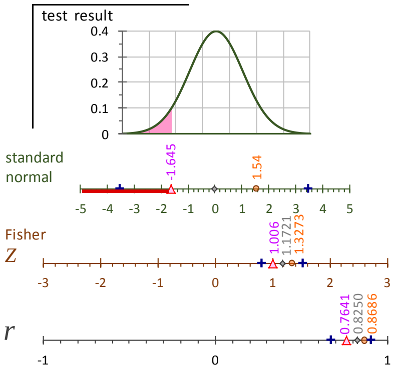

test conclusion

Because $z\ >\ z_{r_E\,crit}$

The test does not reject the H₀ at an α = 0.05 significance level

hypotheses $H_o\colon\ \rho_{A_{x,y}} =$ 0.8250

$H_1\colon\ \rho_{A_{x,y}} \ne$ 0.8250

significance level

$\alpha = 0.05$

Fisher $r$-to-$Z$ transformation

$$Z_{r_E} = \frac{1}{2} ln\left[\frac{1 + (\ 0.8686\ )}{1 - (\ 0.8686\ )}\right] = 1.3273$$

$$Z_{\rho_E} = \frac{1}{2} ln\left[\frac{1 + (\ 0.8250\ )}{1 - (\ 0.8250\ )}\right] = 1.1721$$

$$\sigma_{Z_{r_E}} \approx \sqrt{\frac{1}{n-3}} = 0.1010$$

critical value

$z_{crit} = \pm z_{1-\frac{\alpha}{2}} = \pm 1.96$

test statistic

$$z = \frac{z_{r_E} - z_{\rho_E}}{\sigma_{Z_{r_E}}} = \frac{(\text{-}1.1462) - (\text{-}0.8814)}{0.1010} = 1.54$$

test criteria

$if \begin{cases} |z| \ge z_{E_{crit}} \\ |z| < z_{crit} \end{cases}$ then reject the H₀ / do not reject H₀

test result

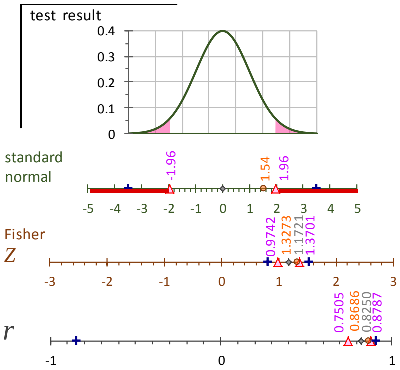

test conclusion

Because $-z_{r_E\,crit} < z < +z_{r_E\,crit}$

The test does not reject the H₀ at an α = 0.05 significance level

hypotheses $H_o\colon\ \rho_{A_{x,y}} =$ 0.8250

$H_1\colon\ \rho_{A_{x,y}} >$ 0.8250

significance level

$\alpha = 0.05$

Fisher $r$-to-$Z$ transformation

$$Z_{r_E} = \frac{1}{2} ln\left[\frac{1 + (\ 0.8686\ )}{1 - (\ 0.8686\ )}\right] = 1.3273$$

$$Z_{\rho_E} = \frac{1}{2} ln\left[\frac{1 + (\ 0.8250\ )}{1 - (\ 0.8250\ )}\right] = 1.1721$$

$$\sigma_{Z_{r_E}} \approx \sqrt{\frac{1}{n-3}} = 0.1010$$

critical value

$z_{crit} = z_{1-\frac{\alpha}{2}} = 1.645$

test statistic

$$z = \frac{z_{r_E} - z_{\rho_E}}{\sigma_{Z_{r_E}}} = \frac{(\ 1.3273\ ) - (\ 1.17214)}{0.1010} = 1.54$$

test criteria

$if \begin{cases} z \ge z_{E_{crit}} \\ otherwise \end{cases}$ then reject the H₀ / do not reject H₀

test result

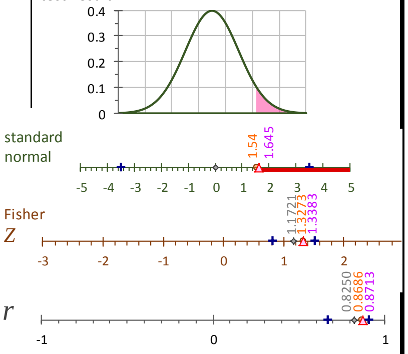

test conclusion

Because $z\ >\ z_{r_E\,crit}$

The test does not reject the H₀ at an α = 0.05 significance level

### Hypothesis test of population environmental correlation

hypotheses $H_o\!: \rho_{A_{x,y}} = 0.8250$

$H_1: \rho_{A_{x,y}} < 0.8250$

significance level

$\alpha = 0.05$

Fisher $r$-to-$Z$ transformation

$$Z_{r_E} = \frac{1}{2} ln\left[\frac{1+(0.8686)}{1-(0.8686)}\right] = 1.3273$$

$$Z_{\rho_E} = \frac{1}{2} ln\left[\frac{1+(0.8250)}{1-(0.8250)}\right] = 1.1721$$

$$\sigma_{Z_{r_E}} \approx \sqrt{\frac{1}{n-3}} = 0.1010$$

critical value

$z_{crit} = z_\alpha = -1.645$

test statistic

$$z = \frac{z_{r_E} - z_{\rho_E}}{\sigma_{Z_{r_E}}} = \frac{(-1.1462) - (-0.8814)}{0.1010} = 1.54$$

test criteria

$if \begin{cases} z \le z_{E_{crit}} \\ otherwise \end{cases}$ then reject the H₀ / do not reject H₀

test result

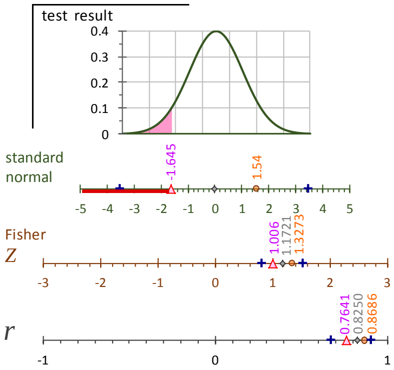

test conclusion

Because $z > z_{r_E\,crit}$

The test does not reject the H₀ at an α = 0.05 significance level

---

hypotheses $H_o\!: \rho_{A_{x,y}} = 0.8250$

$H_1: \rho_{A_{x,y}} \neq 0.8250$

significance level

$\alpha = 0.05$

Fisher $r$-to-$Z$ transformation

$$Z_{r_E} = \frac{1}{2} ln\left[\frac{1+(0.8686)}{1-(0.8686)}\right] = 1.3273$$

$$Z_{\rho_E} = \frac{1}{2} ln\left[\frac{1+(0.8250)}{1-(0.8250)}\right] = 1.1721$$

$$\sigma_{Z_{r_E}} \approx \sqrt{\frac{1}{n-3}} = 0.1010$$

critical value

$z_{crit} = \pm z_{1-\frac{\alpha}{2}} = \pm 1.96$

test statistic

$$z = \frac{z_{r_E} - z_{\rho_E}}{\sigma_{Z_{r_E}}} = \frac{(-1.1462) - (-0.8814)}{0.1010} = 1.54$$

test criteria

$if \begin{cases} |z| \ge z_{E_{crit}} \\ |z| < z_{crit} \end{cases}$ then reject the H₀ / do not reject H₀

test result

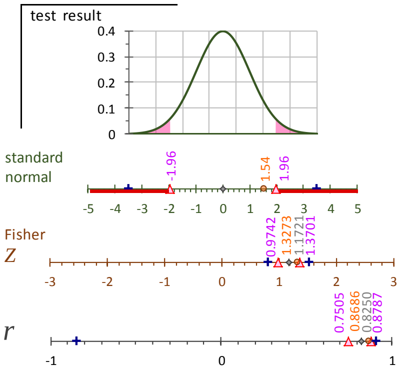

test conclusion

Because $-z_{r_E\,crit} < z < +z_{r_E\,crit}$

The test does not reject the H₀ at an α = 0.05 significance level

---

hypotheses $H_o\!: \rho_{A_{x,y}} = 0.8250$

$H_1: \rho_{A_{x,y}} > 0.8250$

significance level

$\alpha = 0.05$

Fisher $r$-to-$Z$ transformation

$$Z_{r_E} = \frac{1}{2} ln\left[\frac{1+(0.8686)}{1-(0.8686)}\right] = 1.3273$$

$$Z_{\rho_E} = \frac{1}{2} ln\left[\frac{1+(0.8250)}{1-(0.8250)}\right] = 1.1721$$

$$\sigma_{Z_{r_E}} \approx \sqrt{\frac{1}{n-3}} = 0.1010$$

critical value

$z_{crit} = z_{1-\frac{\alpha}{2}} = 1.645$

test statistic

$$z = \frac{z_{r_E} - z_{\rho_E}}{\sigma_{Z_{r_E}}} = \frac{(1.3273) - (1.17214)}{0.1010} = 1.54$$

test criteria

$if \begin{cases} z \ge z_{E_{crit}} \\ otherwise \end{cases}$ then reject the H₀ / do not reject H₀

test result

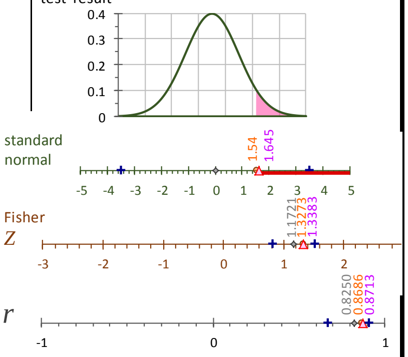

test conclusion

Because $z > z_{r_E\,crit}$

The test does not reject the H₀ at an α = 0.05 significance level

#### Hypothesis test of population coenvironmentability

**hypotheses**

$H_o$: $E_{x,y} =$ 0.7

$H_1$: $E_{x,y} <$ 0.7

**significance level**

$\alpha = 0.05$

**Fisher r-to-Z transformation**

$\widehat{Z_{e_{x,y}}} = Z_{r_P} - \widehat{Z_{h_{x,y}}} = (0.4847) - (0.2154) = 0.7001$

$Z_{E_{x,y}} = \frac{1}{2} ln\left[\frac{1 + (0.7)}{1 - (0.7)}\right] = 0.8673$

$\sigma_{\widehat{Z_{e_{x,y}}}} \approx \sqrt{\frac{1}{n-3}} = 0.1010$

**critical value**

$z_{e\,crit} = z_\alpha = -1.645$

**test statistic**

$z = \frac{\widehat{Z_{e_{x,y}}} - Z_{E_{x,y}}}{\sigma_{\widehat{Z_{e_{x,y}}}}} = \frac{(0.7001) - (0.8673)}{0.1010} = -1.65$

**test criteria**

if $z \le z_{e\,crit}$ then reject the $H_0$

*otherwise* do not reject $H_0$

**test result**

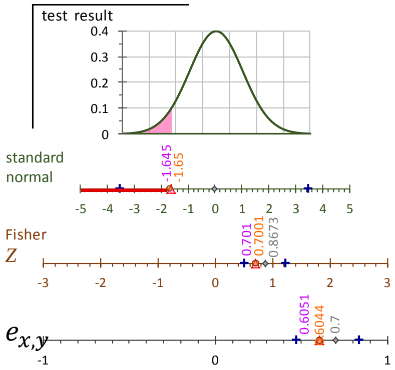

**test conclusion**

Because $z > z_{e\,crit}$

The test does not reject the $H_0$ at an α = 0.05 significance level

---

**hypotheses**

$H_o$: $E_{x,y} =$ 0.7

$H_1$: $E_{x,y} \neq$ 0.7

**significance level**

$\alpha = 0.05$

**Fisher r-to-Z transformation**

$\widehat{Z_{e_{x,y}}} = Z_{r_P} - \widehat{Z_{h_{x,y}}} = (0.4847) - (0.2154) = 0.7001$

$Z_{E_{x,y}} = \frac{1}{2} ln\left[\frac{1 + (0.7)}{1 - (0.7)}\right] = 0.8673$

$\sigma_{\widehat{Z_{e_{x,y}}}} \approx \sqrt{\frac{1}{n-3}} = 0.1010$

**critical value**

$z_{e\,crit} = \pm z_{1-\frac{\alpha}{2}} = \pm 1.96$

**test statistic**

$z = \frac{\widehat{Z_{e_{x,y}}} - Z_{E_{x,y}}}{\sigma_{\widehat{Z_{e_{x,y}}}}} = \frac{(0.7001) - (0.8673)}{0.1010} = -1.65$

**test criteria**

if $|z| \ge z_{e_{crit}}$ then reject the $H_0$

$|z| < z_{h_{crit}}$ do not reject $H_0$

**test result**

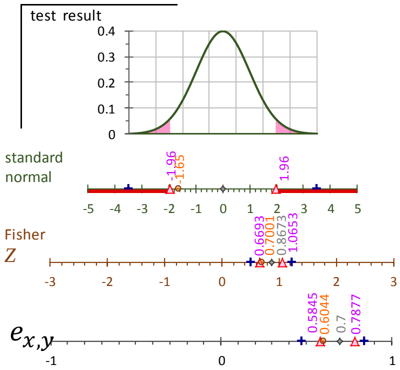

**test conclusion**

Because $-z_{e\,crit} < z < +z_{e\,crit}$

The test does not reject the $H_0$ at an α = 0.05 significance level

---

**hypotheses**

$H_o$: $E_{x,y} =$ 0.7

$H_1$: $E_{x,y} >$ 0.7

**significance level**

$\alpha = 0.05$

**Fisher r-to-Z transformation**

$\widehat{Z_{e_{x,y}}} = Z_{r_P} - \widehat{Z_{h_{x,y}}} = (0.4847) - (0.2154) = 0.7001$

$Z_{H_{x,y}} = \frac{1}{2} ln\left[\frac{1 + (0.7)}{1 - (0.7)}\right] = 0.8673$

$\sigma_{\widehat{Z_{e_{x,y}}}} \approx \sqrt{\frac{1}{n-3}} = 0.1010$

**critical value**

$z_{e\,crit} = z_{1-\frac{\alpha}{2}} = 1.645$

**test statistic**

$z = \frac{\widehat{Z_{e_{x,y}}} - Z_{E_{x,y}}}{\sigma_{\widehat{Z_{e_{x,y}}}}} = \frac{(0.7001) - (0.8673)}{0.1010} = -1.65$

**test criteria**

if $z \ge z_{e_{crit}}$ then reject the $H_0$

*otherwise* do not reject $H_0$

**test result**

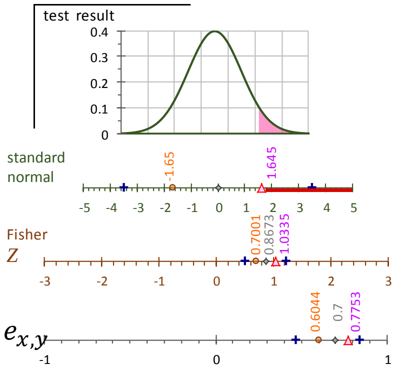

**test conclusion**

Because $z > z_{e\,crit}$

The test does not reject the $H_0$ at an α = 0.05 significance level

#### 5.4 Power of the hypothesis test

Power (sensitivity) is defined as the chance of appropriately rejecting the $H_0$ if the data are drawn from the alternative hypothesis $H_1$. It is calculated from the area of $H_1$ in the points set under the Ho distribution as the critical values.

Here the classical approach for determining the power of the test is presented for the case where the null hypothesis states that the parameter is has a nonzero value.

The method to calculate the power of the test for the coheritability is based on the Fisher's Z transformation. Since there is no way to analytically determine the Z transform of the heritability and coenvironmentability given the value of the phenotypic correlation, the approach taken here was to determine the proportion of the coheritability in reference to the phenotypic correlation (i.e. $\frac{h_{x,y}}{r_{P_{x,y}}}$). The assumption is that the proportion is retained even when the phenotypic correlation has been transformed to $Z_{r_P}$ (which is a very demanding assumption). Multiplying the calculated proportion times the $Z_{r_P}$ will yield a product " $Z_{h_{x,y}}$" which is assumed to be the coheritability in the Z-space.

This method is certainly not robust enough, and rather indicates that further work is needed in this topic.

### Hypothesis test of population phenotypic correlation

Hypotheses

$H_0: \rho_P = -0.5$

$H_1: \rho_P < -0.5$

This is a single left-tailed test

Significance level

α = 0.05

Fisher r-to-Z transformation

$$Z_{r_P} = \ln\left[\frac{1+(-0.3183)}{1-(-0.3183)}\right] = -0.330$$

$$Z_{\rho_P} = \ln\left[\frac{1+(-0.5)}{1-(-0.5)}\right] = -0.549$$

Standard deviation $\sigma_{Zr_P}$ = 0.101

Critical Value

P( $z < z_{CRIT}$ ) = α = 0.05 $z_{CRIT}$ = -1.645

Test Statistic

$$z = \frac{Z_{r_P} - Z_{\rho_P}}{\sigma_{Zr_P}} = \frac{(-0.330) - (-0.549)}{0.101} = 2.173$$

Test Criterion

if $z \le z_{crit}$ then reject the $H_0$

if $z > z_{crit}$ do not reject $H_0$

Test Result

Standard normal score $z$

Fisher Z

Sample Correlation coefficient r

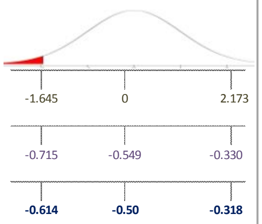

Test Conclusion

Because $z > z_{CRIT}$

The test does not reject the $H_0$ at an

α = 0.05 significance level

Hypotheses

$H_0: \rho_P = -0.5$

$H_1: \rho_P \neq -0.5$

Significance level

α = 0.05

Fisher r-to-Z transformation

$$Z_{r_P} = \ln\left[\frac{1+(-0.3183)}{1-(-0.3183)}\right] \quad -0.330$$

$$Z_{\rho_P} = \ln\left[\frac{1+(-0.5)}{1-(-0.5)}\right] \quad -0.549$$

Standard deviation $\sigma_{Zr_P}$ 0.101

Critical Value

P( $|z| > z_{CRIT}$ ) = α = 0.05 $z_{CRIT}$ = ± 1.96

Test Statistic

$$z = \frac{Z_{r_P} - Z_{\rho_P}}{\sigma_{Zr_P}} = \frac{(-0.330) - (-0.549)}{0.101} = 2.173$$

Test Criterion

if $|z| \ge z_{crit}$ then reject the $H_0$

if $|z| < z_{crit}$ do not reject $H_0$

Test Result

Standard normal score $z$

Fisher Z

Sample Correlation correlation r

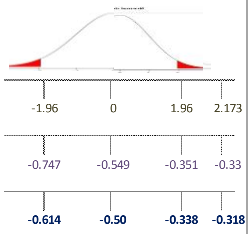

Test Conclusion

Because $z > z_{CRIT}$

The test rejects the $H_0$ at an

α = 0.05 significance level

Hypotheses

$H_0: \rho_P = -0.5$

$H_1: \rho_P > -0.5$

Significance level

α = 0.05

Fisher r-to-Z transformation

$$Z_{r_P} = \ln\left[\frac{1+(-0.3183)}{1-(-0.3183)}\right] = -0.330$$

$$Z_{\rho_P} = \ln\left[\frac{1+(-0.5)}{1-(-0.5)}\right] = -0.549$$

Standard deviation $\sigma_{Zr_P}$ = 0.101

Critical Value

P( $z < z_{CRIT}$ ) = α = 0.05 $z_{CRIT}$ = 1.645

Test Statistic

$$z = \frac{Z_{r_P} - Z_{\rho_P}}{\sigma_{Zr_P}} = \frac{(-0.330) - (-0.549)}{0.101} = 2.173$$

Test Criterion

if $z \ge z_{crit}$ then reject the $H_0$

if $z < z_{crit}$ do not reject $H_0$

Test Result

Standard normal score $z$

Fisher Z

Sample Correlation coefficient r

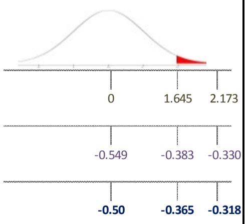

Test Conclusion

Because $z > z_{CRIT}$

The test rejects the $H_0$ at an

α = 0.05 significance level

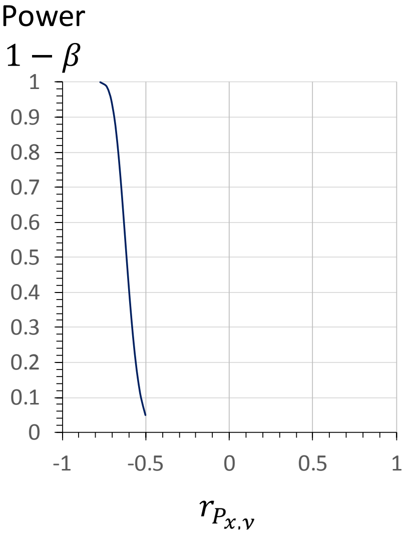

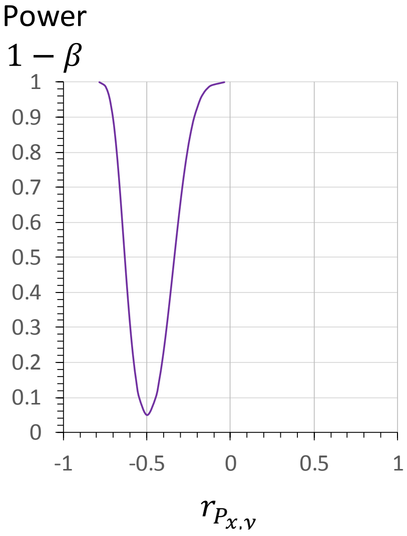

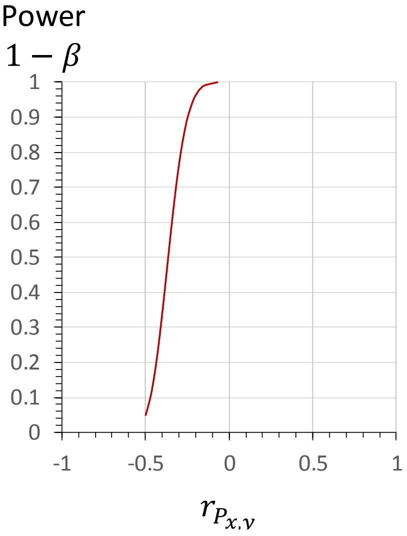

### Hypothesis test of population genetic correlation

**Hypotheses**

$H_0: \rho_A = 0.6$

$H_1: \rho_A < 0.6$

This is a single left-tailed test

Significance level

$\alpha = 0.05$

Fisher r-to-Z transformation

$$Z_{r_A} = \ln\left[\frac{1+(0.4)}{1-(0.4)}\right] = 0.424$$

$$Z_{\rho_A} = \ln\left[\frac{1+(0.6)}{1-(0.6)}\right] = 0.693$$

Standard deviation $\sigma_{Zr_P} = 0.101$

Critical Value

$P(z < z_{CRIT}) = \alpha = 0.05$ $\quad z_{CRIT} = -1.645$

Test Statistic

$$\boldsymbol{z} = \frac{Z_{r_A} - Z_{\rho_A}}{\sigma_{Zr_A}} = \frac{(-0.330) - (-0.549)}{0.101} = -2.668$$

Test Criterion

if $\boldsymbol{z} \le \boldsymbol{z}_{crit}$ then reject the $H_0$

$\boldsymbol{z} > \boldsymbol{z}_{crit}$ do not reject $H_0$

Test Result

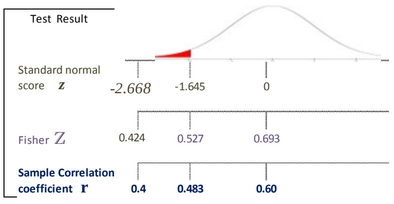

Test Conclusion

Because $z < z_{CRIT}$

The test reject the $H_0$ at an $\alpha = 0.05$ significance level

**Hypotheses**

$H_0: \rho_A = 0.6$

$H_1: \rho_A \ne 0.6$

Significance level

$\alpha = 0.05$

Fisher r-to-Z transformation

$$Z_{r_A} = \ln\left[\frac{1+(0.4)}{1-(0.4)}\right] \quad 0.424$$

$$Z_{\rho_A} = \ln\left[\frac{1+(0.6)}{1-(0.6)}\right] \quad 0.693$$

Standard deviation $\sigma_{Zr_P} \quad 0.101$

Critical Value

$P(|z| > z_{CRIT}) = \alpha = 0.05$ $\quad z_{CRIT} = \pm 1.96$

Test Statistic

$$\boldsymbol{z} = \frac{Z_{r_A} - Z_{\rho_A}}{\sigma_{Zr_A}} = \frac{(-0.330) - (-0.549)}{0.101} = -2.668$$

Test Criterion

if $|\boldsymbol{z}| \ge \boldsymbol{z}_{crit}$ then reject the $H_0$

$|\boldsymbol{z}| < \boldsymbol{z}_{crit}$ do not reject $H_0$

Test Result

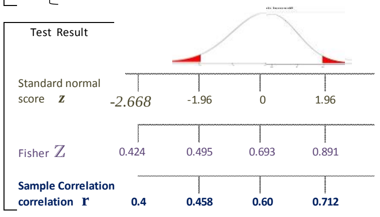

Test Conclusion

Because $z < z_{CRIT}$

The test rejects the $H_0$ at an $\alpha = 0.05$ significance level

**Hypotheses**

$H_0: \rho_A = 0.6$

$H_1: \rho_A > 0.6$

Significance level

$\alpha = 0.05$

Fisher r-to-Z transformation

$$Z_{r_A} = \ln\left[\frac{1+(0.4)}{1-(0.4)}\right] = 0.424$$

$$Z_{\rho_A} = \ln\left[\frac{1+(0.6)}{1-(0.6)}\right] = 0.693$$

Standard deviation $\sigma_{Zr_P} = 0.101$

Critical Value

$P(z < z_{CRIT}) = \alpha = 0.05$ $\quad z_{CRIT} = 1.645$

Test Statistic

$$\boldsymbol{z} = \frac{Z_{r_A} - Z_{\rho_A}}{\sigma_{Zr_A}} = \frac{(-0.330) - (-0.549)}{0.101} = -2.668$$

Test Criterion

if $\boldsymbol{z} \ge \boldsymbol{z}_{crit}$ then reject the $H_0$

$\boldsymbol{z} < \boldsymbol{z}_{crit}$ do not reject $H_0$

Test Result

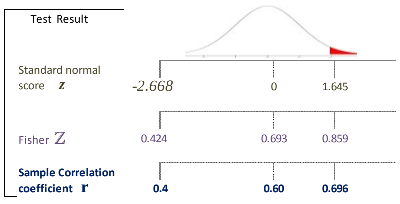

Test Conclusion

Because $z < z_{CRIT}$

The test does not reject the $H_0$ at an $\alpha = 0.05$ significance level

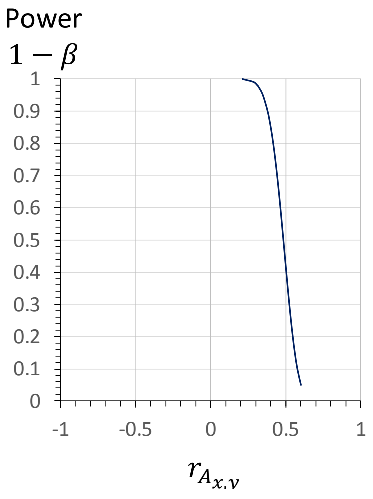

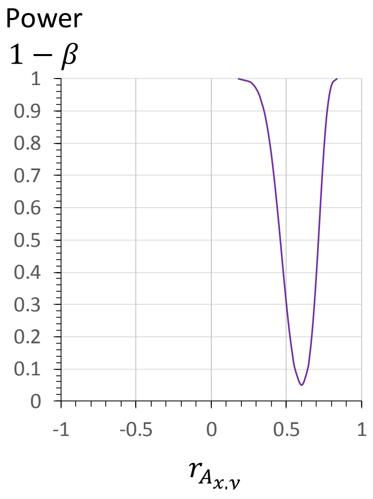

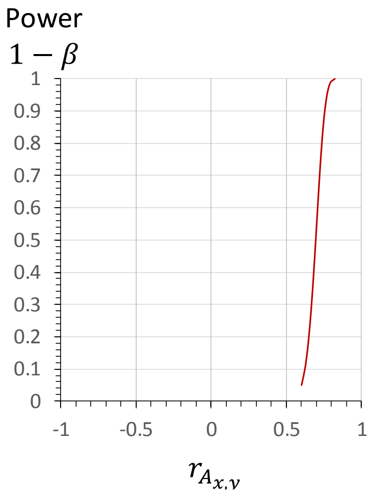

#### Hypothesis test of population coheritability

Hypotheses

$H_0: H_{xy} = 0.25$

$H_1: H_{xy} < 0.25$

This is a single left-tailed test

Significance level

$\alpha = 0.05$

Fisher r-to-Z transformation

$Z_{hxy} = \frac{hxy}{rP}\text{arctanh}(rP) = 0.160$

$Z_{Hxy} = \ln\left[\frac{1+(0.25)}{1-(0.25)}\right] = 0.255$

Standard deviation $\sigma_{Zhxy} = 0.101$

Critical Value

$P(z < z_{CRIT}) = \alpha = 0.05$ $\quad z_{CRIT} = -1.645$

Test Statistic

$$z = \frac{Z_{hxy} - Z_{Hxy}}{\sigma_{Zhxy}} = \frac{(0.160) - (0.255)}{0.101} = -0.940$$

Test Criterion

if $z \le z_{crit}$ then reject the $H_0$; $z > z_{crit}$ do not reject $H_0$

Test Result

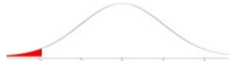

Standard normal score $z$: -1.645, -0.940, 0

Fisher Z: 0.089, 0.16, 0.255

**Sample Correlation coefficient r**: **0.089**, **0.159**, **0.25**

Test Conclusion

Because $z < z_{CRIT}$

The test reject the $H_0$ at an $\alpha = 0.05$ significance level

Hypotheses

$H_0: H_{xy} = 0.25$

$H_1: H_{xy} \neq 0.25$

Significance level

$\alpha = 0.05$

Fisher r-to-Z transformation

$Z_{hxy} = \frac{hxy}{rP}\text{arctanh}(rP) = 0.160$

$Z_{Hxy} = \ln\left[\frac{1+(0.25)}{1-(0.25)}\right] = 0.255$

Standard deviation $\sigma_{hxy}$ 0.101

Critical Value

$P(|z| > z_{CRIT}) = \alpha = 0.05$ $\quad z_{CRIT} = \pm 1.96$

Test Statistic

$$z = \frac{Z_{rP} - Z_{\rho P}}{\sigma_{ZrP}} = \frac{(0.160) - (0.255)}{0.101} = -0.940$$

Test Criterion

if $|z| \ge z_{crit}$ then reject the $H_0$; $|z| < z_{crit}$ do not reject $H_0$

Test Result

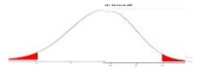

Standard normal score $z$: -1.96, -0.940, 0, 1.96

Fisher Z: 0.057, 0.16, 0.255, 0.453

**Sample Correlation correlation r**: **0.057**, **0.159**, **0.25**, **0.425**

Test Conclusion

Because $z < z_{CRIT}$

The test rejects the $H_0$ at an $\alpha = 0.05$ significance level

Hypotheses

$H_0: H_{xy} = 0.25$

$H_1: H_{xy} > 0.25$

Significance level

$\alpha = 0.05$

Fisher r-to-Z transformation

$Z_{hxy} = \frac{hxy}{rP}\text{arctanh}(rP) = 0.160$

$Z_{Hxy} = \ln\left[\frac{1+(0.25)}{1-(0.25)}\right] = 0.255$

Standard deviation $\sigma_{hxy} = 0.101$

Critical Value

$P(z < z_{CRIT}) = \alpha = 0.05$ $\quad z_{CRIT} = 1.645$

Test Statistic

$$z = \frac{Z_{rP} - Z_{\rho P}}{\sigma_{ZrP}} = \frac{(0.160) - (0.255)}{0.101} = -0.940$$

Test Criterion

if $z \ge z_{crit}$ then reject the $H_0$; $z < z_{crit}$ do not reject $H_0$

Test Result

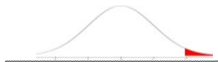

Standard normal score $z$: -0.940, 0, 1.645

Fisher Z: 0.16, 0.255, 0.422

**Sample Correlation coefficient r**: **0.159**, **0.25**, **0.398**

Test Conclusion

Because $z < z_{CRIT}$

The test does not reject the $H_0$ at an $\alpha = 0.05$ significance level

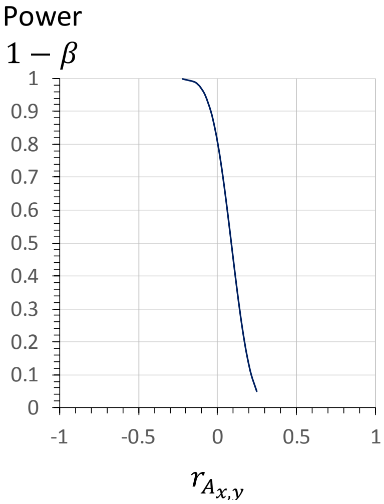

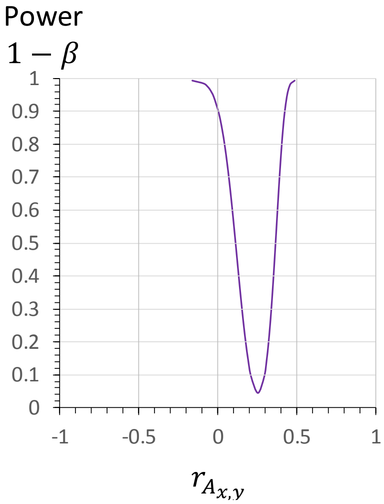

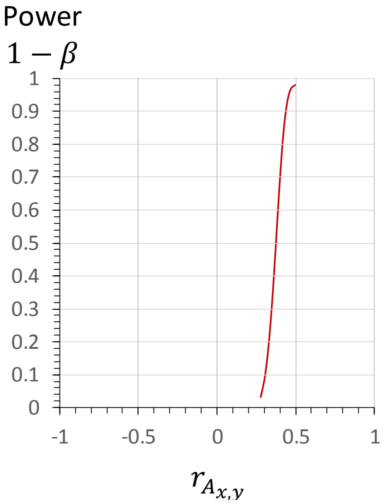

#### 5.5 Confidence interval of $r_{P_{xy}}$, $h_{xy}$ and $e_{xy}$

The confidence interval for a single correlation ( $\rho_\blacksquare$: phenotypic $\rho_P$, genetic $\rho_A$, environmental $\rho_E$) is often obtained using Fisher's $r$ to $Z$ transformation because the sampling distribution of $r$ displays a positive (for $r < 0$) and negative ($r > 0$) skew. To do this, one first forms confidence limits for $Z_{\rho_\blacksquare} = \operatorname{atanh}(r) = \frac{1}{2}\ln\left[\frac{1+\rho_\blacksquare}{1-\rho_\blacksquare}\right]$ (Section 4.4) and then back transform the resultant limits $r = \tanh(Z_r) = \frac{e^{2Z_r}-1}{e^{2Z_r}+1}$ to obtain a confidence interval for $\rho_\blacksquare$ .

The $Z_r$ is the Fisher transform of $\rho$, whose confidence interval are calculated as follows:

the lower bound $Z_L = Z_r - z_{1-\frac{\alpha}{2}}\sqrt{\sigma^2_{Z_r}}$ , and

the upper bound $Z_U = Z_r + z_{1-\frac{\alpha}{2}}\sqrt{\sigma^2_{Z_r}}$

Where $\alpha$ is the significance level, $z_{1-\frac{\alpha}{2}}$ is the $1-\frac{\alpha}{2}$ quantile of the standard normal distribution

$$z_{1-\frac{\alpha}{2}} = \begin{cases} 1.645 & for\ \ \alpha = 90\%\ CI \\ 1.96 & for\ \ \alpha = 95\%\ CI \\ 2.57 & for\ \ \alpha = 99\%\ CI \end{cases}$$

and $\sigma^2_{Z_r} = \frac{1}{n-3}$ is the (approximate) variance of the $Z_r$ distribution, which depends only in the sample size $n$.

It is assumed that the variance of the Fisher transform can be adequately approximated by this value. Fouladi and Steiger (2008) show that although a better approximation to the variance of the Fisher transform $(n-3)^{-1}$ can be obtained, the use of these values or even exact value for the variance of the Fisher transform does not directly translate into statistics with improved size or coverage probabilities with respect to tests or confidence intervals on single correlations.

A worked example follows accompanied by Figure 5.5-1. Another example, in a more succinct presentation is given and represented graphically in Figure 5.5-2.

#### STEP 0. Data

The following be parameter estimators must be obtained from the sample

| | |
|---|---|
| Sample size | $\boldsymbol{n} =$ |
| Phenotypic correlation of traits x and y | $\boldsymbol{r}_{\boldsymbol{P}_{x,y}}$ |
| Additive genetic correlation of traits $x$ and $y$ | $\boldsymbol{r}_{\boldsymbol{A}_{x,y}} =$ |
| Environmental correlation of traits $x$ and $y$ | $\boldsymbol{r}_{\boldsymbol{E}_{x,y}} =$ |
| Narrow-sense heritability of trait $x$ | $\boldsymbol{h}_x^2 =$ |
| Narrow-sense heritability of trait $y$ | $\boldsymbol{h}_y^2 =$ |

EXAMPLE

Let the following data obtained from a quantitative genetics experiment

| | | |
|---|---|---|
| $\boldsymbol{n} =$ | 100 | |
| $\boldsymbol{r}_{\boldsymbol{P}_{x,y}}$ | 0.34 | |
| $\boldsymbol{r}_{\boldsymbol{A}_{x,y}} =$ | -0.4131 | |
| $\boldsymbol{r}_{\boldsymbol{E}_{x,y}} =$ | 0.8451 | |
| $\boldsymbol{h}_x^2 =$ | 0.5 | $\boldsymbol{h}_{x,y} =$ -0.16 |
| $\boldsymbol{h}_y^2 =$ | 0.3 | $\boldsymbol{e}_{x,y} =$ 0.50 |

#### STEP 1. Fisher r-to-Z transformation

Perform the Fisher r-to-Z transformation of the phenotypic $r_{P_{x,y}}$ and genetic $r_{A_{x,y}}$ correlations.

*Phenotypic*

$$Z_{r_P} = 0.5\ ln\left(\frac{1 + r_{P_{xy}}}{1 - r_{P_{xy}}}\right)$$

*Genetic*

$$Z_{r_A} = 0.5\ ln\left(\frac{1 + r_{A_{xy}}}{1 - r_{A_{xy}}}\right)$$

*Phenotypic*

$$Z_{r_P} = 0.5\ ln\left(\frac{1+0.34}{1-0.34}\right) = 0.3541$$

*Genetic*

$$Z_{r_A} = 0.5\ ln\left(\frac{1+(-0.4131)}{1-(-0.4131)}\right) = -0.4393$$

#### STEP 2. Determination of upper and lower bounds for a given α level

for a two-tailed confidence interval the the standard normal $z$ score is given for a fiven significance level α.

$$\underbrace{\left[Z_r - \left(z_{1-\frac{\alpha}{2}}\right)\sqrt{\frac{1}{N-3}}\right]}_{\substack{Z_{r\_L} \\ lower\ \ bound}} < Z_r < \underbrace{\left[Z_r - \left(z_{1-\frac{\alpha}{2}}\right)\sqrt{\frac{1}{N-3}}\right]}_{\substack{Z_{r\_U} \\ upper\ \ bound}}$$

$$z_{1-\frac{\alpha}{2}} = \begin{cases} 1.960 & for\ \alpha = 0.05,\ z_{1-\frac{0.95}{2}} \\ 2.576 & for\ \alpha = 0.01,\ z_{1-\frac{0.99}{2}} \end{cases}$$

for a two-tailed 95% confidence interval we obtained the following bounds:

*Phenotypic*

$$0.3541 - 1.96\sqrt{\frac{1}{100-3}} < Z_{r_P} < 0.3541 + 1.96\sqrt{\frac{1}{100-3}}$$

$$\underbrace{0.1551}_{Z_{r_{P\_L}}} < Z_{r_P} < \underbrace{0.5531}_{Z_{r_{P\_U}}}$$

*Genetic*

$$-0.4393 - 1.96\sqrt{\frac{1}{100-3}} < Z_{r_A} < -0.4393 + 1.96\sqrt{\frac{1}{100-3}}$$

$$\underbrace{-0.6384}_{Z_{r_{A\_L}}} < Z_{r_A} < \underbrace{-0.2403}_{Z_{r_{A\_U}}}$$

#### STEP 3. Back-transformation of $Z_r$ in $r$

Use the following relationship to transform $Z_r$ back into correlations in $r$

$$\underbrace{\frac{e^{2\,Z_{r_{P\_L}}}-1}{e^{2\,Z_{r\_L}}+1}}_{r_{■\_L}} < \rho < \underbrace{\frac{e^{2\,Z_{r\_U}}-1}{e^{2\,Z_{r\_U}}+1}}_{r_{■\_U}}$$

*Phenotypic*

$$\frac{e^{2\,(0.1551)}-1}{e^{2\,(0.1551)}+1} < \rho_P < \frac{e^{2\,(0.5531)}-1}{e^{2\,(0.5531)}+1}$$

$$\underbrace{0.1539}_{r_{P\_L}} < \rho_P < \underbrace{0.5028}_{r_{P\_U}}$$

*Genetic*

$$\frac{e^{2\,(-0.6384)}-1}{e^{2\,(-0.6384)}+1} < \rho_A < \frac{e^{2\,(-0.2403)}-1}{e^{2\,(-0.2403)}+1}$$

$$\underbrace{-0.5638}_{r_{A\_L}} < \rho_A < \underbrace{-0.2358}_{r_{A\_U}}$$

#### STEP 4. Confidence interval of the coheritability

Multiply the confidence interval of the genetic correlation by the product of the square root of the heritabilities

$$\underbrace{\sqrt{h_x^2 . h_y^2}\, r_{A\_L}}_{h_{x,y\_L}} < \underbrace{\sqrt{h_x^2 . h_y^2}\, \rho_A}_{H_{x,y}} < \underbrace{\sqrt{h_x^2 . h_y^2}\, r_{A\_U}}_{h_{x,y\_U}}$$

$$\sqrt{0.5x0.3}(-0.5638) < \sqrt{h_x^2 . h_y^2}\, \rho_A < \sqrt{0.5x0.3}(-0.2358)$$

$$\underbrace{-0.2183}_{h_{x,y\_L}} < H_{x,y} < \underbrace{-0.0913}_{h_{x,y\_U}}$$

#### STEP 5. Confidence level of the coenvironmentability

The lower bound $r_{E_L}$ equals the lower bound of phenotypic correlation minus the lower bound of the coheritability.

The upper bound $r_{E_U}$ equals the upper bound of the phenotypic correlation minus the upper bound of the coheritability.

$$\underbrace{(r_{P_L} - h_{x,y\_L})}_{r_{E_L}} < E_{x,y} < \underbrace{(r_{P_U} - h_{x,y\_U})}_{r_{E_U}}$$

$$0.1539 - (-0.2138) < E_{x,y} < 0.5028 - (-0.0913)$$

$$0.4059 < E_{x,y} < 0.5941$$

To summarize, the confidence interval for the parameters (sample size n=100) are

$$\underbrace{0.1539}_{r_{P\_L}} < \rho_P < \underbrace{0.5028}_{r_{P\_U}}$$

$$\underbrace{-0.2183}_{h_{x,y\_L}} < H_{x,y} < \underbrace{-0.0913}_{h_{x,y\_U}}$$

$$0.4059 < E_{x,y} < 0.5941$$

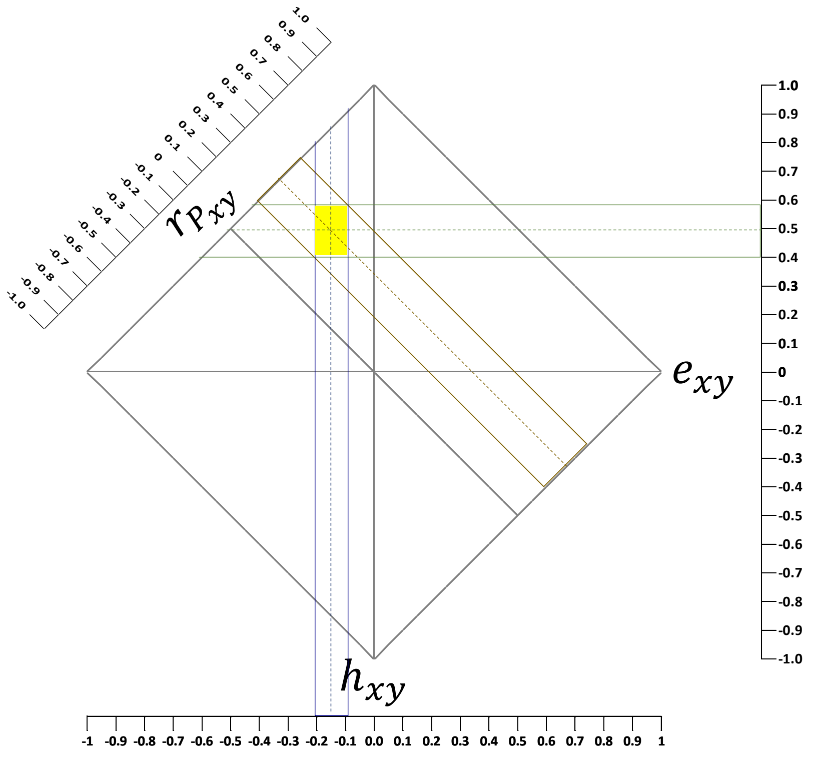

**Figure 5.5**-1 Confidence intervals of the phenotypic correlation, coheritability and coenvironmentability (yellow rectangle) .

| $n = 628$<br>$h_x^2 = 0.3$<br>$h_y^2 = 0.4$ | | Fisher's $r$-to-$Z$ transformation<br>$Z_{r} = 0.5\, ln\left(\frac{1 + r_{x,y}}{1 - r_{x,y}}\right)$ | Confidence Interval of $Z_r$<br>$\left[Z_r - (z_{1-\frac{\alpha}{2}})\sqrt{\frac{1}{N-3}}\right] < Z_r < \left[Z_r - (z_{1-\frac{\alpha}{2}})\sqrt{\frac{1}{N-3}}\right]$ | back transformation from $Z_r$ to $r$<br>$\underbrace{\frac{e^{2 Z_{r_{P\_L}}} - 1}{e^{2 Z_{r\_L}} + 1}}_{r_{\_L}} < \rho < \underbrace{\frac{e^{2 Z_{r\_U}} - 1}{e^{2 Z_{r\_U}} + 1}}_{r_{\_U}}$ | confidence interval coheritability<br>$\underbrace{\sqrt{h_x^2 . h_y^2}\, r_{A\_L}}_{h_{x,y\_L}} < \underbrace{\sqrt{h_x^2 . h_y^2}\, \rho_A}_{H_{x,y}} < \underbrace{\sqrt{h_x^2 . h_y^2}\, r_{A\_U}}_{h_{x,y\_U}}$ | confidence interval coenvironmentability<br>$\underbrace{(r_{P_L} - h_{x,y\_L})}_{r_{E_L}} < E_{x,y} < \underbrace{(r_{P_U} - h_{x,y\_U})}_{r_{E_U}}$ |
|---|---|---|---|---|---|---|
| $r_{P_{x,y}}$ | 0.24 | 0.549 | $0.549 - 1.96\sqrt{\frac{1}{628-3}} < Z_{r_P} < 0.549 + 1.96\sqrt{\frac{1}{628-3}}$<br>$\underbrace{0.4706}_{Z_{r_{P\_L}}} < Z_{r_P} < \underbrace{0.6274}_{Z_{r_{P\_U}}}$ | $\frac{e^{2(0.4706)} - 1}{e^{2(0.4706)} + 1} < \rho_P < \frac{e^{2(0.6274)} - 1}{e^{2(0.6274)} + 1}$<br>$\underbrace{\mathbf{0.165}}_{\mathbf{r_{P\_L}}} < \boldsymbol{\rho_P} < \underbrace{\mathbf{0.312}}_{\mathbf{r_{P\_U}}}$ | | |
| $r_{A_{x,y}}$ | -0.58 | -0.662 | $-0.662 - 1.96\sqrt{\frac{1}{628-3}} < Z_{r_A} < -0.662 + 1.96\sqrt{\frac{1}{628-3}}$<br>$\underbrace{-0.7404}_{Z_{r_{A\_L}}} < Z_{r_A} < \underbrace{-0.5836}_{Z_{r_{A\_U}}}$ | $\frac{e^{2(-0.7404)} - 1}{e^{2(-0.7404)} + 1} < \rho_A < \frac{e^{2(-0.5836)} - 1}{e^{2(-0.5836)} + 1}$<br>$\underbrace{-0.629}_{r_{A\_L}} < \rho_A < \underbrace{-0.526}_{r_{A\_U}}$ | | |
| $h_{x,y} = \sqrt{h_x^2\, h_y^2}\, r_{A_{x,y}}$ | -0.2 | | | | $\sqrt{0.3\ x\ 0.4}(-0.629) < \sqrt{h_x^2 . h_y^2}\, \rho_A < \sqrt{0.3\ x\ 0.4}(-0.526)$<br>$\underbrace{\mathbf{-0.218}}_{\mathbf{h_{x,y\_L}}} < \mathbf{H_{x,y}} < \underbrace{\mathbf{-0.182}}_{\mathbf{h_{x,y\_U}}}$ | |
| $e_{x,y} = r_{P_{x,y}} - h_{x,y}$ | 0.348 | | | | | $0.165 - (-0.218) < E_{x,y} < 0.312 - (-0.182)$<br>$\mathbf{0.383} < \mathbf{E_{x,y}} < \mathbf{0.494}$ |

#### 5.6 Sample size

An important aspect in planning a study that will require inference on the phenotypic correlation and its components is the determination of the appropriate sample size $n$ in order to maximize the probability of credible results. Sometimes, the sample size is determined by practical issues regarding resources, time, and money. In this case the number of individuals to be measure is known, yet it is nevertheless important to determine the inferential extent the data can provide when used for hypothesis testing. A usual procedure of choosing $n$ aims to achieve sufficient power of the test and level of significance to determine whether a parameter is zero ( $Ho: \theta = 0$). The following formula can be used to that effect:

$$n = \left( \frac{z_{1-\frac{\alpha}{2}} + z_{1-\beta}}{\frac{1}{2} ln \left[ \frac{1+\theta}{1-\theta} \right]} \right)^2 + 3$$

where $z_{1-\frac{\alpha}{2}}$ and $z_{1-\beta}$ are the standard normal $z$ scores corresponding to the $1 - \frac{\alpha}{2}$ and $1 - \beta$ quantiles of the standard normal distribution (e.g. for $\alpha = 0.05$ , $z_{1-\frac{\alpha}{2}} = 1.96$; $\beta = 0.2$, $z_{1-\beta} = 0.842$), and the denominator within the brackets is the Fisher's transformation of $\theta$ (where $\theta$ is the parameter to be assessed, e.g., $r_{P_{x,y}}$ , $r_{A_{x,y}}$ , $h_{x,y}$)

A better procedure for the determination of the optimum sample size for the study of joint inheritance of two traits uses confidence intervals. This procedure aims to find a sample size that guarantees confidence intervals with a given width to detect a given effect size. Here we intend to specify a null hypothesis in which the parameter is equal to a nonzero value. The attention, therefore, is on confidence level estimation of the population parameter instead of the test of a parameter hypothesized null value. The following equation can be used to determine the sample size required to obtain a 95% CI for the parameter $\theta$ with a desired width (Moinester and Gottfried 2014).

$$n = \frac{3.84\,(1-\theta^2)^2}{w^2} + 6\theta^2 + 1$$

And the corresponding equation for the half-width of the confidence interval is

$$w = 1.96\frac{(1-\theta^2)}{\sqrt{n-1-6\theta^2}}$$

where $\theta$ is the effect size, that is the value of the parameter of interest, e.g., $r_{P_{x,y}}$ , $r_{A_{x,y}}$ , $h_{x,y}$; and $w$ is half-width of the desired confidence interval. If the sample size is already fixed, one can calculate the corresponding half-width w for a given expected effect size $\theta$.

Effect size $\theta$ is the expected value of the parameter that one suspects or guesses.

**A**

half-width $w$

| effect size $\lvert\theta\rvert$ | 0.03 | 0.035 | 0.04 | 0.05 | 0.07 | 0.1 |
|---|---|---|---|---|---|---|
| 0.05 | 4255 | 3127 | 2394 | 1533 | 782 | 384 |
| 0.10 | 4192 | 3080 | 2358 | 1510 | 771 | 378 |
| 0.15 | 4086 | 3003 | 2299 | 1472 | 751 | 369 |
| 0.20 | 3942 | 2896 | 2218 | 1420 | 725 | 356 |
| 0.25 | 3759 | 2762 | 2115 | 1354 | 692 | 340 |
| 0.30 | 3542 | 2603 | 1993 | 1276 | 652 | 320 |
| 0.35 | 3294 | 2420 | 1854 | 1187 | 606 | 298 |
| 0.40 | 3019 | 2218 | 1699 | 1088 | 556 | 273 |
| 0.45 | 2721 | 2000 | 1532 | 981 | 502 | 247 |
| 0.50 | 2408 | 1769 | 1355 | 868 | 444 | 219 |
| 0.55 | 2083 | 1531 | 1173 | 752 | 385 | 190 |
| 0.60 | 1754 | 1290 | 988 | 634 | 325 | 161 |
| 0.65 | 1429 | 1051 | 806 | 517 | 265 | 132 |
| 0.70 | 1116 | 821 | 629 | 404 | 208 | 104 |
| 0.75 | 823 | 606 | 465 | 299 | 155 | 78 |
| 0.80 | 559 | 412 | 317 | 204 | 107 | 55 |
| 0.85 | 335 | 247 | 191 | 124 | 66 | 35 |
| 0.90 | 160 | 119 | 93 | 61 | 34 | 20 |
| 0.95 | 47 | 36 | 29 | 21 | 14 | 10 |

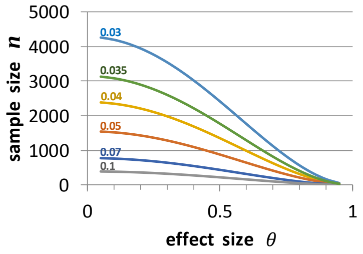

**B**

| test statistic $\lvert\theta\rvert$ | $\alpha = 0.05$, $\beta = 0.20$ |
|---|---|
| 0.05 | 3138 |
| 0.10 | 783 |
| 0.15 | 347 |
| 0.20 | 194 |
| 0.25 | 123 |
| 0.30 | 85 |
| 0.35 | 62 |
| 0.40 | 47 |
| 0.45 | 36 |
| 0.50 | 29 |
| 0.55 | 24 |
| 0.60 | 19 |
| 0.65 | 16 |
| 0.70 | 13 |
| 0.75 | 11 |
| 0.80 | 10 |
| 0.85 | 8 |
| 0.90 | 7 |
| 0.95 | 5 |

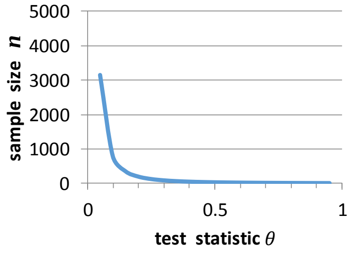

**Figure 5.6**. (A) Sample size required to obtain a desired 95% confidence interval half-width for different values of the correlations, or coheritability) according to the formula of Moinester and Gottfried (2014). (B) Sample size required to test a null hypothesis that the parameter is zero with a significance level $\alpha = 0.05$, and power $1-\beta = 0.8$.

Since in absolute value the coheritability is generally smaller than the genetic correlation, it has implications on lowering the effect size, and it is expected that the sample size for testing the coheritability will generally be larger than that for the genetic correlation. For example, if a test is to be carried out having a test statistic $r_{A_{x,y}} = 0.6$ and set the probability to reject a true null hypothesis to $\alpha = 0.05$ (Type I error), and a probability for failing to reject a false null hypothesis to $\beta = 0.2$ (Type II error) then the sample size required to determine whether a genetic correlation differs from zero would be $n = 19$. In contrast, given the same genetic correlation and $h_x^2 = 0.28$, $h_y^2 = 0.405$, the coheritability becomes $h_{x,y} = (\sqrt{0.28 \cdot 0.405} \cdot 0.6) = 0.202$, and the sample size would be $n = 190$ to establish a 80% power of the hypothesis test.

A more meaningful approach, however, is to determine a sample size that guarantees a narrow confidence interval for the coheritability. For instance, using the data from Figure 5.5.2, if the genetic correlation is expected to be $r_{A_{x,y}} = -0.58$ (this is the effect size), and would like to measure it within a confidence interval $[-0.62\ , -0.54]$, which is $-0.58 \pm 0.04$, then using the equation of Moinester and Gottfried refereed above,

$$n = \frac{3.84\ (1 - (0.58)^2)^2}{(0.04)^2} + 6\ (0.58)^2 + 1 = \mathbf{1596}$$

If the coheritability is expected to be $h_{x,y} = -0.2$, and also would like to measure around a $\pm w = \pm 0.04$, that is $[-0.24, -0.16]$, then

$$n = \frac{3.84\ (1 - (0.2)^2)^2}{(0.04)^2} + 6\ (0.2)^2 + 1 = \mathbf{2305}$$

which results in a larger sample size for the coheritability because its effect size $(0.2)$ is smaller than the one for the genetic correlation $(0.58)$.

On the other hand, if we have already a fixed number of individuals to measure, say a sample of size $n = 981$ and would like to know what is the confidence level that can be obtained if the coheritability is $h_{x,y} = -0.2$. Again, we use the equation from Moinester and Gottfried,

$$w = 1.96\frac{(1-(0.2)^2)}{\sqrt{981-1-6(0.2)^2}} = 0.06$$

Therefore, with a fixed sample size $n = 981$, the coheritability $h_{x,y} = -0.2$ would be calculated within a confidence interval $[(-0.2-0.06),(-0.2+0.06)] = [-0.26\,,-0.14]$. Note that the interval is a little broader because of the reduced sample size.

### 6 Visualizing the $\boldsymbol{h_{xy}} \cdot \boldsymbol{e_{xy}} \cdot \boldsymbol{r_{P_{xy}}}$-relationship

#### 6.1 The three-dimensional $h_{x,y}$▪$e_{x,y}$▪$r_{P_{x,y}}$ -plane (3DHER-plane)

The relationship $r_{P_{x,y}} = (h_{x,y} + e_{x,y})$ allow us to infer the domains of the phenotypic correlation, coheritability and coenvironmentability, as follows

$$-1 \leq r_{P_{x,y}} \leq 1$$

$$-1 \leq h_{x,y} \leq 1$$

$$-1 \leq e_{x,y} \leq 1$$

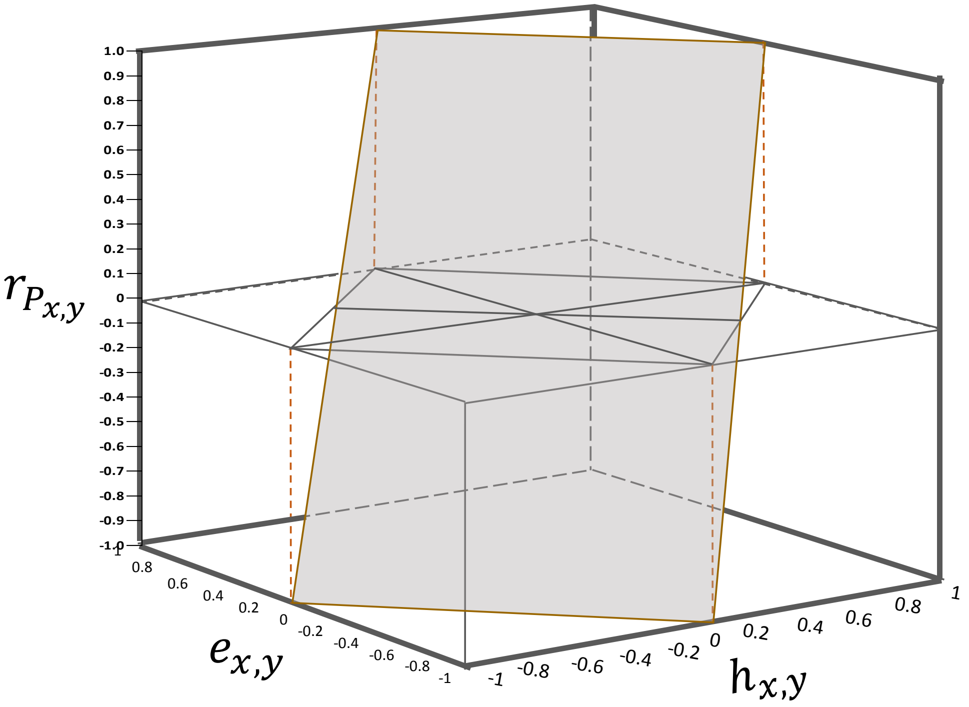

**Figure 6.1** 3DHER-plane

If each variable (coheritability, coenvironmentability, phenotypic correlation) are assigned to an ordered triplet of axis lines, which are numerical, intersect at the origin, are pair-wise perpendicular, and all have a single unit of length, it is possible to uniquely specify, in this Cartesian system, the values of a single point by signed distances from the origin, i.e. three numbers in a chosen order $(h_{x,y}, e_{x,y}, r_{P_{x,y}})$. This is illustrated in Figure 6.1 and is called the three-dimensional coheritability-coenvironmentability-phenotypic correlation plane, 3DHER-plane.

The data align on a plane (i.e. it has zero volume) due to the interdependencies established by the intrinsic linear dependency of the axes, namely, $r_{P_{x,y}} = \left(h_{x,y} + e_{x,y}\right)$ (see Section 6.4).

#### 6.2 The two-dimensional $h_{xy}$▪$e_{xy}$▪$r_{P_{xy}}$ -field (2DHER-field)

The orthogonal projection of the 3D $h_{x,y} \cdot e_{x,y} \cdot r_{P_{x,y}}$ –plane on the surface defined by $h_{xy}$ x $e_{xy} | \, r_{P_{xy}} = 0$, results in the two-dimensional $h_{x,y} \cdot e_{x,y} \cdot r_{P_{x,y}}$ –field. This field is circumscribed by the relationship $|h_{x,y}| + |e_{x,y}| = 1$, which in a more explicit way, is demarcated by the lines $h_{x,y} + e_{x,y} = 1$ , $h_{x,y} + e_{x,y} = -1$, $h_{x,y} - e_{x,y} = 1$, and $h_{x,y} - e_{x,y} = -1$ (Figure 6.2)

The field $h_{xy}$▪$e_{xy}$▪$r_P$ can be partitioned according to the sign of the phenotypic correlation, coheritability and coenvironmentability. Note here that the coheritability sign is conferred by the additive genetic correlation. The sign of the cohenvironmentability is given by the environmental correlation. By tracing the lines $r_{P_{xy}} = 0$, $h_{xy} = 0$, and $e_{xy} = 0$, sectors are demarcated within the field forming isosceles triangles. These partitions are characterized by (1) a particular sign combination among components, (2) the range of values the components can have, and (3) and the relationship between the sign of the phenotypic correlation and the coheritability and coenvironmentability.

The partitions are denoted by the letter $S$ followed by a subscripted numeral having the sign of the coheritability of that sector. For instance, partitions $S_{+1}$ , $S_{+2}$ , $S_{+3}$ have a positive coheritability, and therefore a positive genetic correlation. The sign of the phenotypic correlation does not follow necessarily the sign of the coheritability or genetic correlation.

The subscripts with the same numeral but with opposite signs indicate sectors that are reciprocal to each other. For example, partition $S_{+3}$ has $h_{xy} > 0$, and $e_{xy} < 0$, while partition $S_{-3}$ has $h_{xy} < 0$, and $e_{xy} > 0$. In both $S_{+3}$ and $S_{-3}$ the phenotypic correlation $r_{P_{xy}}$ has the sign of $e_{xy}$ , indicating the preponderant effect of the coenvironmentability over the coheritability component. Notice that it implies that $r_{P_{xy}}$ and $r_{A_{xy}}$ have opposite sign.

Partitions $S_{+1}$ and $S_{-1}$ are reciprocal sectors where the coheritability and coenvironmentability have the same sign, and contribute to the magnitude of the phenotypic correlation in the same direction (sign). Both $h_{xy}$ and $e_{xy}$ act in the same direction (same sign).

Partitions $S_{+2}$ and $S_{-2}$ are reciprocal sectors characterized by the preponderant effect of the coheritability on the phenotypic correlation, at the expense of the coenvironmentability. This is clearly seen by $r_{P_{xy}}$ having the same sign as the coheritability (and $r_{A_{xy}}$). Both $h_{xy}$ and $e_{xy}$ display an antagonistic relationship, they act in opposite directions (different signs). This may also suggest the reduced effect of the GxE interaction.

Partitions $S_{+3}$ and $S_{-3}$ are reciprocal sectors characterized by the phenotypic correlation and the coheritability having opposite signs, and the coenvironmentability having a preponderant effect on the magnitude and sign of the phenotypic correlation. Both $h_{xy}$ and $e_{xy}$ display an antagonistic relationship, they act in opposite directions (different signs). This may be an indicator that a probable GxE interaction may be present.

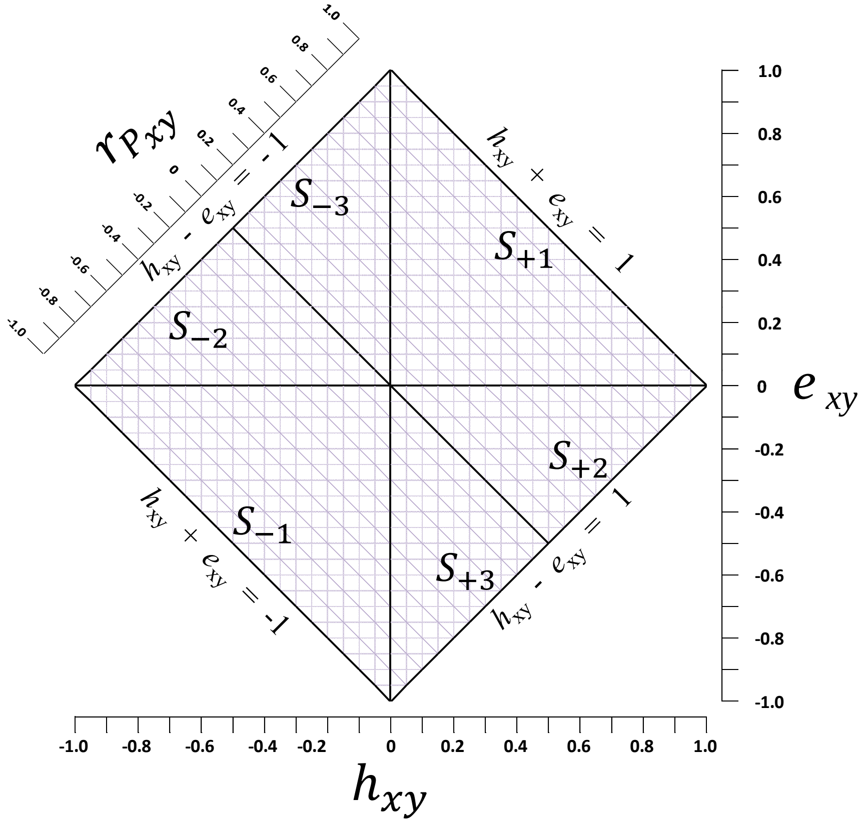

| $r_{P_{xy}}$ | $h_{xy}$ | $e_{xy}$ | |
|---|---|---|---|
| [ -1 , 0 ) | [ -1 , 0 ) | [ -1 , 0 ) | $S_{-1}$ |
| [ -1 , 0 ) | [ -1 , 0 ) | ( 0 , 0.5 ] | $S_{-2}$ |
| ( 0 , 1 ] | [ -0.5 , 0 ) | ( 0 , 1 ] | $S_{-3}$ |

| | $r_{P_{xy}}$ | $h_{xy}$ | $e_{xy}$ |
|---|---|---|---|
| $S_{+1}$ | ( 0 , 1 ] | ( 0 , 1 ] | ( 0 , 1 ] |
| $S_{+2}$ | ( 0 , 1 ] | ( 0 , 1 ] | [ -0.5 , 0 ) |
| $S_{+3}$ | [ -1 , 0 ) | ( 0 , 0.5 ] | [ -1 , 0 ) |

**Figure 6.2** (A) The two-dimensional coheritability-coenvironmentability-phenotypic correlation-field (2DHER). This field is demarcated by the relationship $|h_{x,y}| + |e_{x,y}| = 1$. Lines at zero for each variable define partitions, labeled by an S followed by the sign of the coheritability and an indicator numeral. (B) Range of values for the variables for each partition.

#### 6.3 The $\boldsymbol{h_{xy}} \cdot \boldsymbol{e_{xy}} \cdot \boldsymbol{r_{P_{xy}}}$-plane in a three-dimensional spherical space

The coheritability, coenvironmentability and the phenotypic correlation, being three numerical coordinates that define a point  in a 3-dimensional Cartesian system, could easily be represented into a spherical coordinate system where the position of a point in space is specified by spherical coordinates $(\mathrm{r}\,,\ \theta\,,\ \varphi\,)$, (symbols as often used in mathematics)(Figure 6.3-1).

$\mathrm{r}$ ,   the radius (radial coordinate, $\mathrm{r}\ \geq 0$ ) defined as the Euclidean distance from the origin O to the point P.

$\theta$,   the azimuthal angle (azimuth coordinate, $0 < \theta < 2\pi$) is the signed angle measured from the azimuth reference direction (positive $x$-axis) to the orthogonal projection of the line segment OP on the reference plane ($x$-$y$ plane). The azimuth coordinate corresponds to the coheritability.

$\varphi$,   the polar angle (zenith angle coordinate, $0 < \varphi < \pi$) is the angle between the zenith direction ($z$-axis) to the line segment OP that connects the origin O to the point P. The zenith coordinate corresponds to the phenotypic correlation.

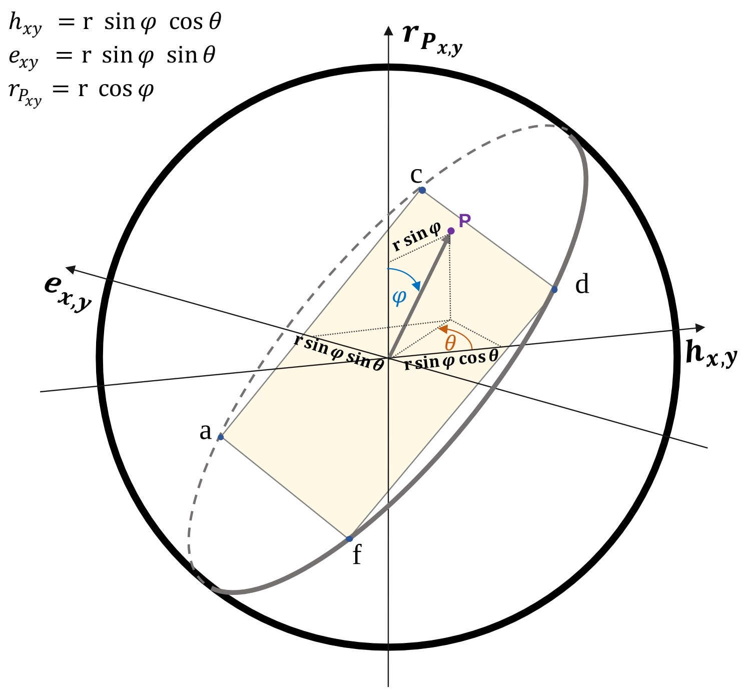

**Figure 6.3-1** The spherical coordinate system corresponding to the variables coheritability, coenvironmentability and phenotypic correlation.  The point can be uniquely specified by $(\mathrm{r}\,, \theta\,, \varphi\,)$.   The plane is inclined 35.26438968° in reference to the $r_{P_{x,y}}.\,e_{x,y}$-plane and rotated 45°. The radius of the sphere is equal to $\sqrt{2}$.

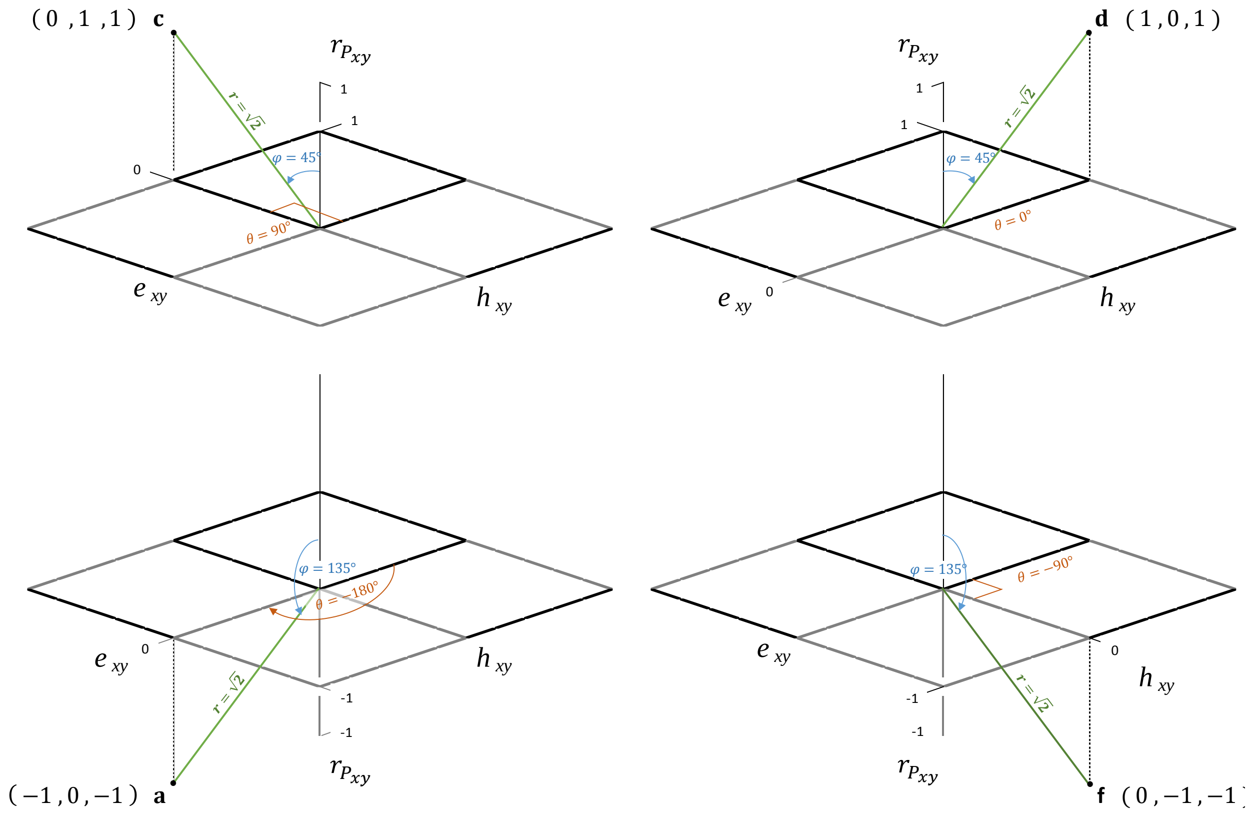

| | | Cartesian coordinates | | | Spherical coordiantes | | | | | | | | | |
|---|---|---|---|---|---|---|---|---|---|---|---|---|---|---|
| | | ***h*** *xy* | ***e*** *xy* | ***r*** *p* | **azimuth Θ** | **polar φ** | **radius r** | **sinΘ** | **cosΘ** | **sinφ** | **cosφ** | ***h*** *x,y* **ρ sin φ cos Θ** | ***e*** *x,y* **ρ sin ϕ sin Θ** | ***r*** *Px,y* **ρ cos ϕ** |
| point | **a** | 0 | 1 | 1 | 90 | 45 | $\sqrt{2}$ | 1 | 0 | $\frac{\sqrt{2}}{2}$ | $\frac{\sqrt{2}}{2}$ | **0** | **1** | **1** |
| | **b** | 1 | 0 | 1 | 0 | 45 | $\sqrt{2}$ | 0 | 1 | $\frac{\sqrt{2}}{2}$ | $\frac{\sqrt{2}}{2}$ | **1** | **0** | **1** |
| | **c** | 0 | -1 | -1 | -90 | 135 | $\sqrt{2}$ | -1 | 0 | $\frac{\sqrt{2}}{2}$ | $-\frac{\sqrt{2}}{2}$ | **0** | **-1** | **-1** |
| | **d** | -1 | 0 | -1 | 180 | 135 | $\sqrt{2}$ | 0 | -1 | $\frac{\sqrt{2}}{2}$ | $-\frac{\sqrt{2}}{2}$ | **-1** | **0** | **-1** |

**Figure 6.3-2** Examples of determination of specified positions in the spherical coordinate system corresponding to the coheritability, coenvironmentability and phenotypic correlation.

The spherical coordinates $(\mathrm{r}\,,\ \theta\,,\ \varphi\,)$ are related to the Cartesian coordinates $(h_{xy}\,,\ e_{xy}\,,\ r_{P_{xy}})$ by the formulae

$$\mathrm{r} = \sqrt{h_{xy}^2 + e_{xy}^2 + r_{P_{xy}}^2}\,, \qquad \mathrm{r}\,\epsilon\,\left[0\,,\sqrt{2}\right]$$

$$\theta = \arctan\left(\frac{e_{xy}}{h_{xy}}\right) \qquad , \qquad \theta\ \epsilon \begin{cases} [0\,,2\pi) \\ [0°\,,360°) \end{cases}$$

$$\varphi = \arccos\left(\frac{r_{P_{xy}}}{\mathrm{r}}\right) \qquad , \qquad \varphi\ \epsilon \begin{cases} [0\,,\ \pi) \\ [0°\,,180°) \end{cases}$$

The Cartesian coordinates are related to the spherical coordinates as follows

$$h_{xy} = x = \mathrm{r}\ \sin\varphi\ \cos\theta$$

$$e_{xy} = y = \mathrm{r}\ \sin\varphi\ \sin\theta$$

$$r_{P_{xy}} = z = \mathrm{r}\ \cos\varphi$$

The following relationship must be satisfied

$$\overbrace{\mathrm{r}\cos(\varphi)}^{r_{P_{x,y}}} = \overbrace{\sin(\varphi)cos\theta}^{h_{x,y}} + \overbrace{\sin(\varphi)\sin(\theta)}^{e_{x,y}}$$

Relationship between the azimuthal $\theta$ and polar $\varphi$ angles (Figure 6.3-3) is derived as follow

$$r_{P_{x,y}} = h_{x,y} + e_{x,y}$$

$$\mathrm{r}\cos\varphi = \mathrm{r}\sin\varphi\ \cos\theta + \mathrm{r}\sin\varphi\ \sin\theta$$

$$\cos\varphi = \sin\varphi\,[\sin\theta + \cos\theta]$$

$$\frac{\cos\varphi}{\sin\varphi} = \sin\theta + \cos\theta$$

$$\left(\frac{1}{\tan\varphi}\right)^2 = (\sin\theta + \cos\theta)^2 = (\ sin^2\ \theta + cos^2\ \theta + 2\sin\theta\cos\theta\ ) = 1 + 2\sin\theta\cos\theta$$

$$\frac{1}{\varphi} = \sqrt{1 \quad + 2\sin\theta \quad \cos\theta} \quad = \sqrt{1 + \sin(2\theta)}$$

$$\tan\varphi = \frac{1}{\sqrt{1 + \sin(2\theta)}}$$

Finally we obtain

$$\varphi = arctan\left[\frac{1}{\sqrt{1 + \sin(2\theta)}}\right]$$

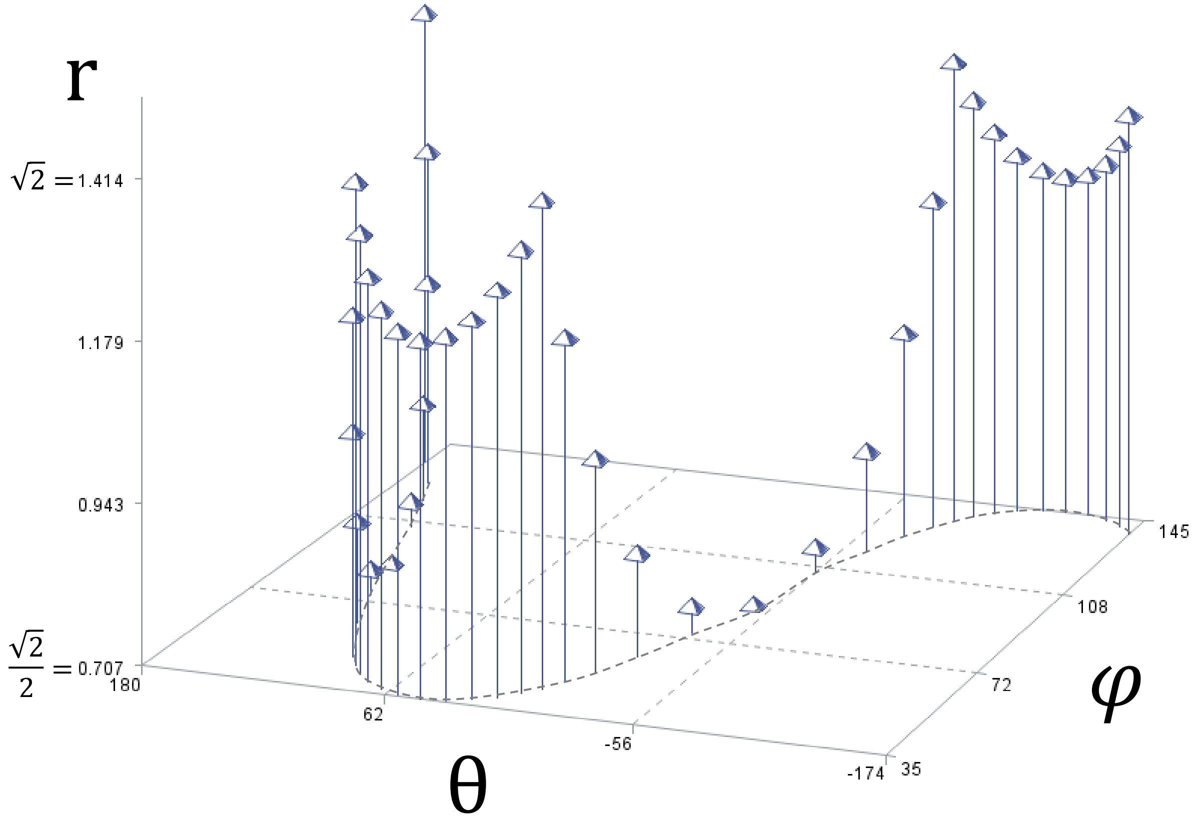

**Figure 6.3-3** Relationship between the radius r, the azimuthal angle $\theta$, and the polar angle $\varphi$.

#### 6.4 The determinant of the $\boldsymbol{h_{xy} \cdot e_{xy} \cdot r_{P_{xy}}}$-plane

The equation $r_{P_{xy}} = h_{xy} + e_{xy}$ establishes a *linear* dependency between the variables that constraints their values. Therefore, a 3x3 matrix in which a column corresponds to a single datum, of the form $\begin{vmatrix} h_{xy} \\ e_{xy} \\ r_{P_{xy}} \end{vmatrix}$ will have a determinant equal to zero, which means that all the data is "squeezed" to a flat plane of zero volume.

Using three of the points a, c, d, f which touch an imaginary sphere of radius $\sqrt{2}$

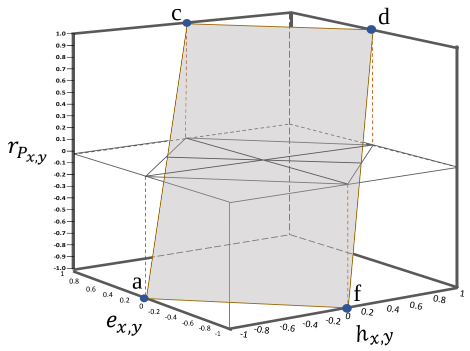

$$\mathbf{a} = \begin{vmatrix} -1 \\ 0 \\ -1 \end{vmatrix}, \quad \mathbf{c} = \begin{vmatrix} 0 \\ 1 \\ 1 \end{vmatrix}, \quad \mathbf{d} = \begin{vmatrix} 1 \\ 0 \\ 1 \end{vmatrix}, \quad \mathbf{f} = \begin{vmatrix} 0 \\ -1 \\ -1 \end{vmatrix}$$

In Cartesian Coordinates

$$det\,[\,\mathbf{a\,c\,d}\,] \quad = \quad det\,[\,\mathbf{a\,c\,f}\,] \quad = \quad det\,[\,\mathbf{a\,d\,f}\,] \quad = \quad det\,[\mathbf{c\,d\,f}\,]$$

$$det \begin{vmatrix} -1 & 0 & 1 \\ 0 & 1 & 0 \\ -1 & 1 & 1 \end{vmatrix} = det \begin{vmatrix} -1 & 0 & 0 \\ 0 & 1 & -1 \\ -1 & 1 & -1 \end{vmatrix} = det \begin{vmatrix} -1 & 1 & 0 \\ 0 & 0 & -1 \\ -1 & 1 & -1 \end{vmatrix} = det \begin{vmatrix} 0 & 1 & 0 \\ 1 & 0 & -1 \\ 1 & 1 & -1 \end{vmatrix} = 0$$

Using the values in of Figure 6.3-2 above, one can find the determinant for three points, say points c, d, f, the determinant has the following format:

$$det \begin{vmatrix} (\mathrm{r}\ \sin\varphi\ \cos\theta)_c & (\mathrm{r}\ \sin\varphi\ \cos\theta)_d & (\mathrm{r}\ \sin\varphi\ \cos\theta)_f \\ (\mathrm{r}\ \sin\varphi\ \sin\theta)_c & (\mathrm{r}\ \sin\varphi\ \sin\theta)_d & (\mathrm{r}\ \sin\varphi\ \sin\theta)_f \\ (\quad \mathrm{r}\cos\varphi \quad)_c & (\quad \mathrm{r}\cos\varphi \quad)_d & (\quad \mathrm{r}\cos\varphi \quad)_f \end{vmatrix} = 0$$

### 7 Set of values and limiting cases

#### 7.1 The range of values of the coheritability for a given value of the coenvironmentability

Given the interdependencies, it is possible to relate the range of values of the coheritability based on a given value of the coenvironmentability, as follows:

$$-(1-|e_{x,y}|) < h_{x,y} < (1-|e_{x,y}|) \qquad -1 < e_{x,y} \leq 1$$

which can be written more explicitly as

$$-(1-e_{x,y}) < h_{x,y} < (1-e_{x,y}) \qquad 0 < e_{x,y} \leq 1$$

$$-1 < h_{x,y} < 1 \qquad e_{x,y} = 0$$

$$-(1+e_{x,y}) < h_{x,y} < (1+e_{x,y}) \qquad -1 \leq e_{x,y} < 0$$

To illustrate: for $e_{x,y} = 0.8$ $\quad -(1-0.8) < h_{x,y} < (1-0.8) \quad \rightarrow \quad -0.2 < h_{x,y} < 0.2$

$e_{x,y} = 0$ $\quad -1 < h_{x,y} < 1 \quad \rightarrow \quad -1 < h_{x,y} < 1$

$e_{x,y} = 0.8$ $\quad -(1+(-0.35)) < h_{x,y} < (1+(-0.35)) \quad \rightarrow \quad -0.65 < h_{x,y} < 0.65$

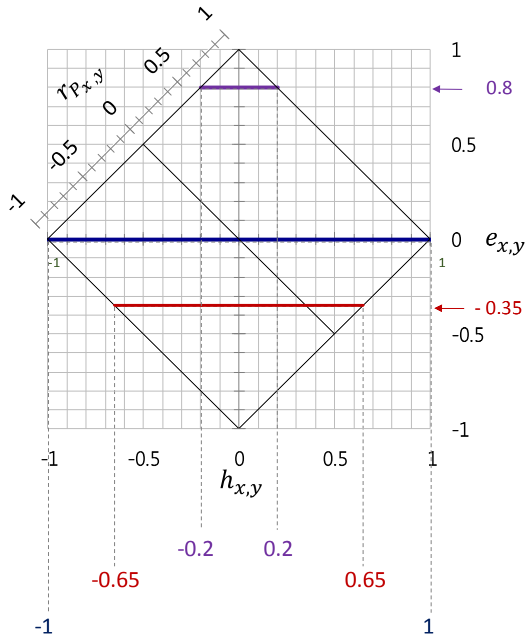

#### 7.2 The set of values of coheritability and coenvironmentability for a given phenotypic correlation

Provided a value of phenotypic correlation, what is the range values allowed for its coheritability and coenvironmentability components? The problem can be expressed as follows:

$$r_{P_{x,y}} \rightarrow (h_{x,y}\,, e_{x,y} |\, r_{P_{x,y}})$$

(a) The set of $(h_{xy}\,, e_{xy})$ values valid for a given $r_{P_{x,y}}$, is a collection of bivariate data whose boundaries are:

$$\left(-0.5(1-\ r_{P_{x,y}})\ ,\ 0.5(1+r_{P_{x,y}})\right) < \left(h_{x,y}\,, e_{x,y}|r_{P_{x,y}}\right) < \left(\ 0.5(1+r_{P_{x,y}})\ ,\ -0.5(1-0.5\ r_{P_{x,y}})\right)$$

and (b) the elements of the datum sum up to the phenotypic correlation

$$h_{x,y} + e_{x,y} = r_{P_{xy}}$$

and, (c) where the linear equation relating the coenvironmentability as a function of the coheritability is

$$e_{x,y} = r_{P_{xy}} -\ h_{x,y}$$

a line whose intercept is the value of the phenotypic correlation, and the slopeof the coheritability factor is -1.

For example, if the phenotypic correlation $r_{P_{xy}}\ = -0.35$, then the $(\ h_{x,y}\ ,\ e_{x,y})$ values follow:

(a) The $(\ h_{x,y}\ ,\ e_{x,y})$ must be within the boundaries

$$(-0.675\,, 0.325) \leq\ (h_{x,y}\,, e_{x,y}|\ r_{P_{x,y}}\ = -0.35) \leq\ (\ 0.325\ \ , -0.675\ )$$

(b) The elements of the datum must necessarily sum up to the phenotypic correlation

$$h_{x,y} + e_{x,y} = -0.35$$

using the boundaries as an example:

$$-0.675 + 0.325\ =\ -0.35, \text{ and }\ 0.325 - 0.675\ =\ -0.35$$

(c) and the linear relationship is that relates the coenvironmentability and coheritability is

$$e_{x,y} = -0.35 - h_{x,y}\ , \qquad (-0.675 < h_{x,y} < 0.325)$$

This line allows to determine the coenvironmentability value for any value of coheritability within the set boundaries. Figure 7.2 illustrates this example.

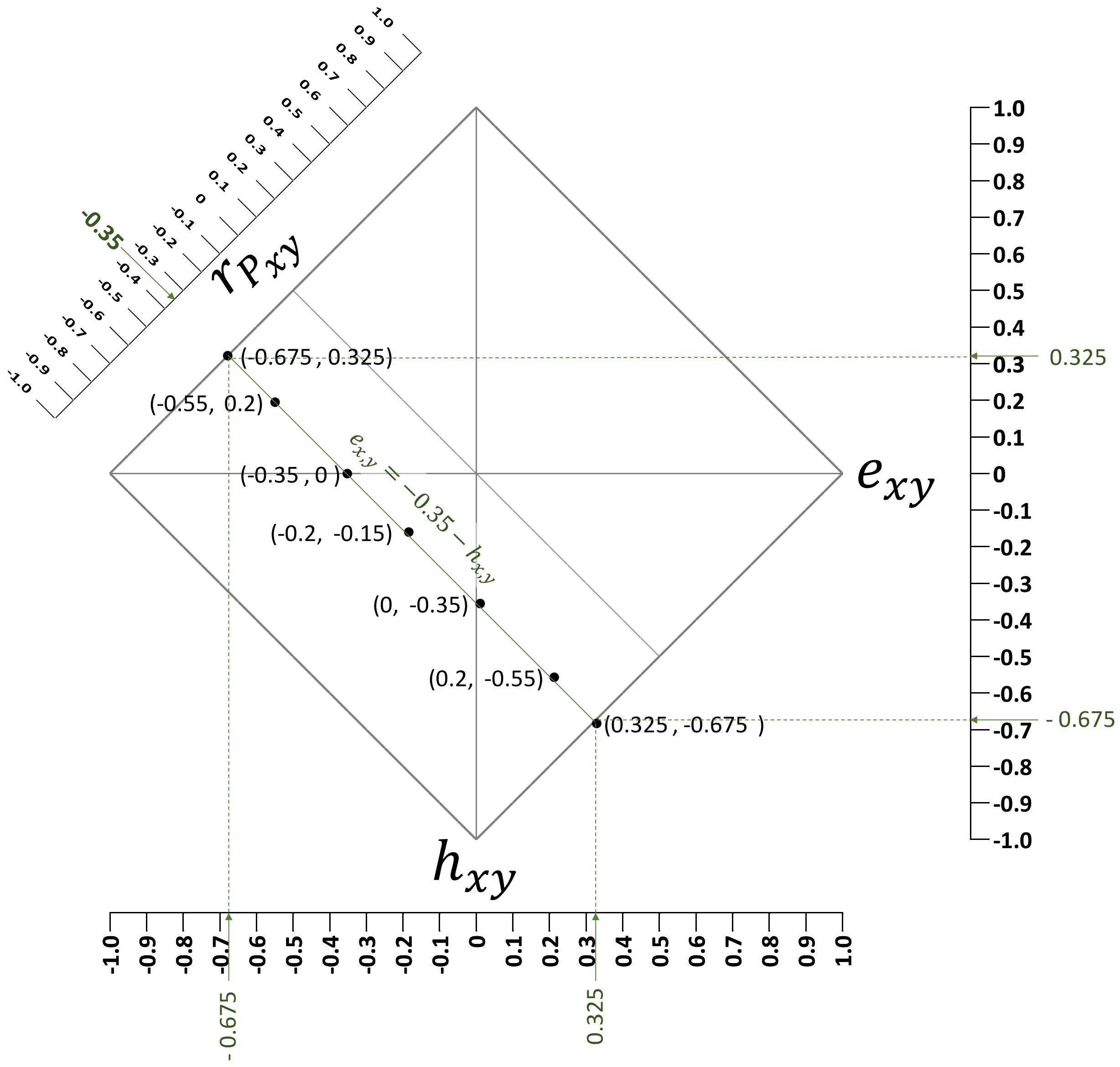

Figure 7.2. Example of some of the possible values of the bivariate datum (coheritability, coenvironmentability) for the given phenotypic correlation of -0.35, presented in the 2DHER-field. Notice that the boundaries of coheritability and coenvironmentability are the numerically the same yet antiparallel. Note that the elements constituting each datum sum up to the phenotypic correlation. The line equation has as intercept the value of the phenotypic correlation, and a slope of -1.

#### 7.3 The set of possible values of heritabilities, and correlations (genetic, environmental) for a given (coheritability, coenvironmentability) datum.

Provided a value of a (coheritability, coenvironmentability) datum, what are the possible values of the heritabilities of the traits and the genetic and the environmental correlations? In other words, knowing a datum involving coheritability,coenvironmentability corresponding to a phenotypic correlation $(h_{x,y}\ , e_{x,y} \mid r_{P_{x\ ,y}})$ what would be the collection of possible values of the heritability of trait $x$ and of trait $y$, the genetic correlation, and the environmental correlation, $(h_x^2\ , h_y^2\ , r_{A_{x,y}}\ , r_{E_{x,y}})$? The problem can be expressed as follows:

$$(h_{x,y}\ , e_{x,y} \mid r_{P_{x\ ,y}}) \rightarrow \left\{ h_x^2\ , \ h_y^2\ , \ r_{A_{x,y}}\ , \ r_{E_{x,y}} \right\}$$

In principle, the values of the phenotypic, genetic and environmental correlations are independent and random variables. A high phenotypic correlation does not imply a comparable value of the genetic correlation. A high genetic correlation between two traits, do not necessarily indicates that the heritabilities of the traits are high also. Traits that have high heritabilities may display a weak genetic correlation, and visceversa. In this sense the heritabilities and genetic correlations are independent. In addition, the effect of the heritabilities may result in changes in the rank of the genetic correlation compared to the rank observed in the coheritabilities.

However, as explained in the previous section, by imposing a constraint (i.e. fixing the value of a variable) such as equating the phenotypic correlation to a specific value would affect the set of bivariate coheritability-coenvironmentability combinations.

To address the problem of finding heritabilities and correlations, here I use an empirical, graphical method where the set of values $r_{A_{x,y}}$, $r_{E_{x,y}}$ are plotted against $\sqrt{h_x^2 h_y^2}$. The algorithm to can be sumnmarized as follows:

(1) list a set of heritabilities combinations, ranked by the value of their geometric mean $\sqrt{h_x^2 h_y^2}$.

| $\sqrt{h_x^2\, h_y^2}$ | 0.1 | 0.14142 | 0.17321 | 0.2 | 0.22361 | 0.24495 | 0.26458 | 0.28284 | 0.3 | 0.31623 | 0.34641 | 0.37417 | 0.3873 | 0.4 | 0.42426 | 0.44721 | 0.45826 |
|---|---|---|---|---|---|---|---|---|---|---|---|---|---|---|---|---|---|
| $h^2_x$ | 0.1 | 0.2 | 0.3 | 0.2 | 0.5 | 0.3 | 0.7 | 0.4 | 0.9 | 0.5 | 0.4 | 0.7 | 0.5 | 0.4 | 0.9 | 0.5 | 0.7 |
| $h^2_y$ | 0.1 | 0.1 | 0.1 | 0.2 | 0.1 | 0.2 | 0.1 | 0.2 | 0.1 | 0.2 | 0.3 | 0.2 | 0.3 | 0.4 | 0.2 | 0.4 | 0.3 |
| $\sqrt{(1-h_x^2)(1-h_y^2)}$ | 0.9 | 0.8485 | 0.7937 | 0.8 | 0.6708 | 0.7483 | 0.5196 | 0.6928 | 0.3 | 0.6325 | 0.6481 | 0.4899 | 0.5916 | 0.6 | 0.2828 | 0.5477 | 0.4583 |

| $\sqrt{h_x^2\, h_y^2}$ | 0.4899 | 0.5 | 0.51962 | 0.52915 | 0.54772 | 0.56569 | 0.59161 | 0.6 | 0.63246 | 0.64807 | 0.69282 | 0.7 | 0.73485 | 0.79373 | 0.8 | 0.84853 | 0.9 |
|---|---|---|---|---|---|---|---|---|---|---|---|---|---|---|---|---|---|
| $h^2_x$ | 0.6 | 0.5 | 0.9 | 0.7 | 0.6 | 0.8 | 0.7 | 0.9 | 0.8 | 0.7 | 0.8 | 0.7 | 0.9 | 0.9 | 0.8 | 0.9 | 0.9 |
| $h^2_y$ | 0.4 | 0.5 | 0.3 | 0.4 | 0.5 | 0.4 | 0.5 | 0.4 | 0.5 | 0.6 | 0.6 | 0.7 | 0.6 | 0.7 | 0.8 | 0.8 | 0.9 |
| $\sqrt{(1-h_x^2)(1-h_y^2)}$ | 0.4899 | 0.5 | 0.2646 | 0.4243 | 0.4472 | 0.3464 | 0.3873 | 0.2449 | 0.3162 | 0.3464 | 0.2828 | 0.3 | 0.2 | 0.1732 | 0.2 | 0.1414 | 0.1 |

in this case, the values are within the interval [0.1 , 0.9] in order to avoid numerical instability.

(2) Second, set the boundaries of the heritabilities. In which the magnitude of the correlation becomes 1. This is determined using the following relationship:

$h_x^2 = h_y^2 = \sqrt{h_x^2 h_y^2} = |\, h_{x,y} \,|$, it results in $r_{A_{x,y}} = \frac{|\, h_{x,y} \,|}{h_{x,y}}$, with value either -1 or +1

and

$h_x^2 = h_y^2 = \sqrt{h_x^2 h_y^2} = 1 - |\, e_{x,y} \,|$, it results in $r_{E_{x,y}} = \frac{|\, e_{x,y} \,|}{e_{x,y}}$, with value either -1 or +1

To summarize, when both heritabilities are either $|\, h_{x,y} \,|$ or $(1 - |\, e_{x,y} \,|)$, it establish the boundaries of the heritability values. Note that even within this interval the correlations may be greater than +1 or less than -1.

(3) The correlations that are within the heritability boundaries and contained in [-1 , +1 ] are the possible valid values to be selected.

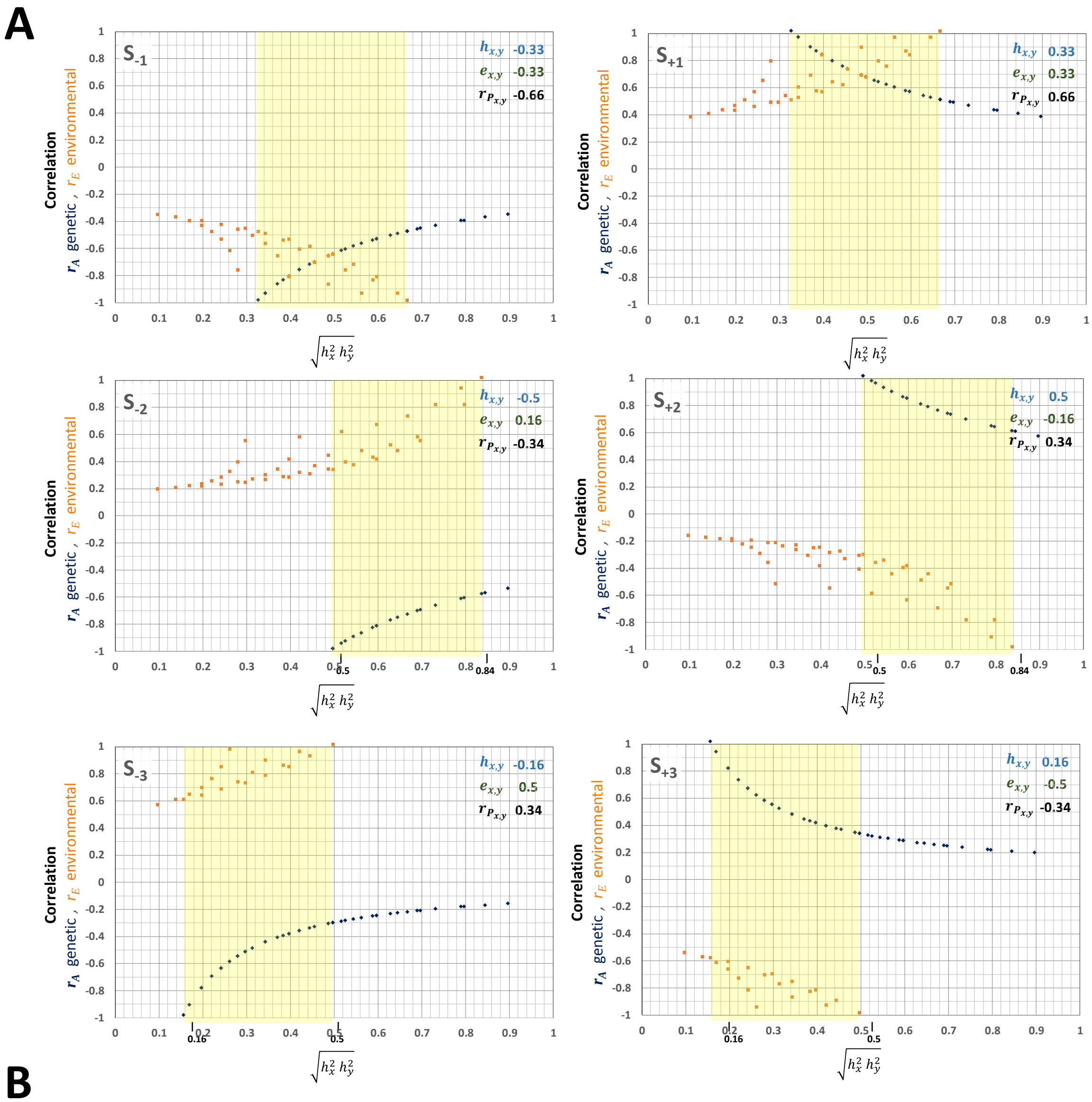

**B**

| $\sqrt{h_x^2\,h_y^2}$ | 0.1 | 0.14142 | 0.17321 | 0.2 | 0.22361 | 0.24495 | 0.26458 | 0.28284 | 0.3 | 0.31623 | 0.34641 | 0.37417 | 0.3873 | 0.4 | 0.42426 | 0.44721 | 0.45826 | 0.4899 |
|---|---|---|---|---|---|---|---|---|---|---|---|---|---|---|---|---|---|---|
| $h^2_x$ | 0.1 | 0.2 | 0.3 | 0.2 | 0.5 | 0.3 | 0.7 | 0.4 | 0.9 | 0.5 | 0.4 | 0.7 | 0.5 | 0.4 | 0.9 | 0.5 | 0.7 | 0.6 |
| $h^2_y$ | 0.1 | 0.1 | 0.1 | 0.2 | 0.1 | 0.2 | 0.1 | 0.2 | 0.1 | 0.2 | 0.3 | 0.2 | 0.3 | 0.4 | 0.2 | 0.4 | 0.3 | 0.4 |
| $\sqrt{(1-h_x^2)(1-h_y^2)}$ | 0.9 | 0.8485 | 0.7937 | 0.8 | 0.6708 | 0.7483 | 0.5196 | 0.6928 | 0.3 | 0.6325 | 0.6481 | 0.4899 | 0.5916 | 0.6 | 0.2828 | 0.5477 | 0.4583 | 0.4899 |

| $\sqrt{h_x^2\,h_y^2}$ | 0.5 | 0.51962 | 0.52915 | 0.54772 | 0.56569 | 0.59161 | 0.6 | 0.63246 | 0.64807 | 0.69282 | 0.7 | 0.73485 | 0.79373 | 0.8 | 0.84853 | 0.9 |
|---|---|---|---|---|---|---|---|---|---|---|---|---|---|---|---|---|
| $h^2_x$ | 0.5 | 0.9 | 0.7 | 0.6 | 0.8 | 0.7 | 0.9 | 0.8 | 0.7 | 0.8 | 0.7 | 0.9 | 0.9 | 0.8 | 0.9 | 0.9 |
| $h^2_y$ | 0.5 | 0.3 | 0.4 | 0.5 | 0.4 | 0.5 | 0.4 | 0.5 | 0.6 | 0.6 | 0.7 | 0.6 | 0.7 | 0.8 | 0.8 | 0.9 |
| $\sqrt{(1-h_x^2)(1-h_y^2)}$ | 0.5 | 0.2646 | 0.4243 | 0.4472 | 0.3464 | 0.3873 | 0.2449 | 0.3162 | 0.3464 | 0.2828 | 0.3 | 0.2 | 0.1732 | 0.2 | 0.1414 | 0.1 |

**Figure 7.3** Example of values of heritabilities and correlations (genetic, environmental) for given values of coheritability and coenvironmentability $(h_{x,y}\,, e_{x,y})\; r_{P_{x,y}}$. (A) Values of genetic (blue) and environmental (orange) correlations as a function of the square root of the product of heritabilities. These graphs are dependent on (B) the list of combinations of heritabilities of the traits ranked by their geometric mean.

#### 7.4 The relationship of the heritabilities to yield a given geometric mean

In the previous section, the algorithm requires to list an ordered set of geometric means of pair of heritabilities. The major caveat of such a method is that there is not a 1-to-1 relationship between a geometric mean and a pair of heritabilities. In fact, a given geometric mean can be obtained by a number of alternative pairs.

For instance, following example consists of heritability pairs whose product is 0.18, and its geometric mean is 0.424264.

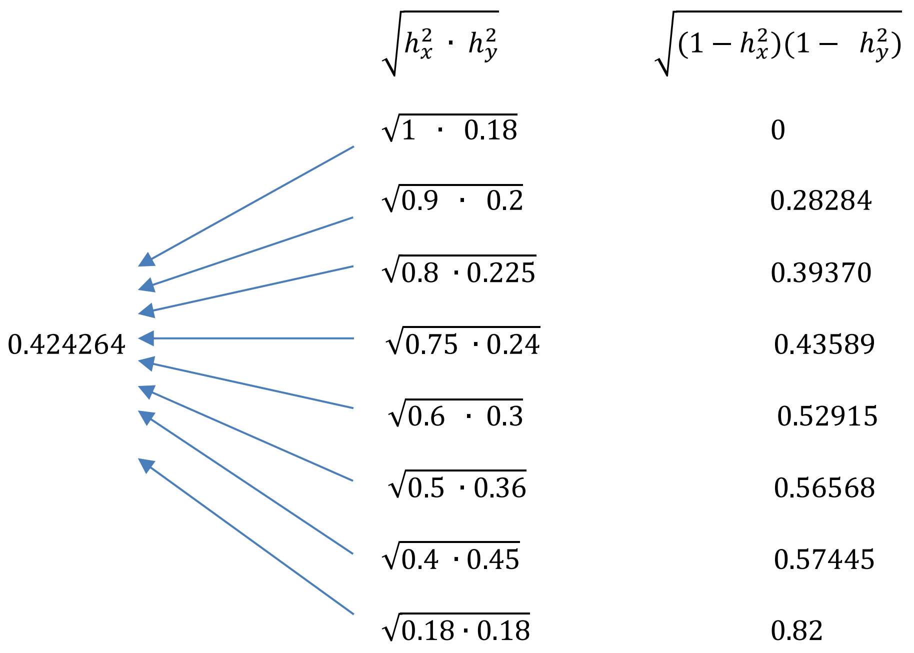

#### 7.5 Limiting cases

Limiting cases provide insight into the nature of the coheritability and coenvironmentability in relation to the phenotypic correlation. For instance, if one of the heritabilities approaches 1, results in vanishing coenvironmentability, and $r_{P_{x,y}} = h_{xy}$. If conversely one of the heritabilities approaches zero, it causes the coheritability to vanish, leading to $r_{P_{x,y}} = e_{xy}$. Clearly this shows that at extreme values of the heritability, the phenotypic correlation can become either the coheritability or coenvironmentability. Therefore the statistical inferences designed to test $r_{P_{x,y}}$ can be used for $h_{xy}$ and $e_{xy}$ as well.

Note that

if $r_{A_{x,y}} < 0$, the multiplying by $\sqrt{h_x^2\, h_y^2}$, will result in coheritability with larger value than $r_{A_{x,y}}$.

If $r_{A_{x,y}} > 0$ then the product of $\sqrt{h_x^2\, h_y^2}$ and $r_{A_{x,y}}$ results in a coheritability value smaller than $r_{A_{x,y}}$.

In both cases, the coheritability will move in a direction towards zero.

Additionally, if $h_x^2 + h_y^2 = 1$, then $\sqrt{h_x^2\, h_y^2} = \sqrt{(1-h_x^2)(1-h_x^2)}$.

The phenotypic correlation would be zero if the coheritability and coenvironmentability are of the same magnitude but of different sign. Another case exists when both the coheritability and coenvironmentability are zero; this would happen if one of the heritabilities is unity and the other zero, or if both genetic and environmental correlations is zero.

The sample correlation coefficient, as well as the coheritability estimator, depending on the sample size involved, can display numerical instability, especially at the proximity$\pm 1$. Both the sample correlation and the coheritability are not sufficiently robust, so its values can be misleading if outliers are present. Cognizant of this fact, we can make some deductions in this regard:

(A) A heritability equal to zero can only occur if the additive genetic variance (in the numerator) is zero. Under this condition, however, such zero genetic variance enters into the formula of the genetic correlation and causes it to be mathematically undefined (i.e. division by zero). Therefore, zero heritability cannot coexist with non null genetic correlation.

(B) A similar rationale can also be applied to the environmental correlation. A heritability equal to 1 implies that the additive genetic variance and the phenotypic variance are the same, and that the environmental variance is zero. Since the zero environmental variance is part of the denominator of the environmental correlation, it will cause to become undefined. Therefore, a heritability equal to 1 and a non null environmental correlation cannot exist.

Table 7.5 presents the algebraic formula for coheritability, coenvironmentability and phenotypic correlations, as affected by the heritabilities and the correlations.

### Table 7.5 Special Cases

| | $0 < h_x^2 < 1$ and $0 < h_y^2 < 1$ | | $h_x^2 = 0.5$ and $h_y^2 = 0.5$ | | $h_x^2 = h_y^2$ | | $h_x^2 + h_y^2 = 1$ | |
|---|---|---|---|---|---|---|---|---|
| $-1 < r_{A_{x,y}} < 1$<br>$-1 < r_{E_{x,y}} < 1$ | [1] $r_{P_{x,y}} = \sqrt{h_x^2 h_y^2}\, r_{A_{x,y}} + \sqrt{(1-h_x^2)(1-h_y^2)}\, r_{E_{x,y}}$ | | [10] $r_{P_{x,y}} = 0.5\left[r_{A_{x,y}} + r_{E_{x,y}}\right]$ | | [19] $r_{P_{x,y}} = h_x^2\,(r_{A_{x,y}} + r_{E_{x,y}}) + r_{E_{x,y}}$ | | [28] $r_{P_{x,y}} = \sqrt{h_x^2(1-h_x^2)}\left[r_{A_{x,y}} + r_{E_{x,y}}\right]$ | |
| | $h_{x,y} = \sqrt{h_x^2 h_y^2}\, r_{A_{x,y}}$ | $e_{x,y} = \sqrt{(1-h_x^2)(1-h_y^2)}\, r_{E_{x,y}}$ | $h_{x,y} = 0.5\, r_{A_{x,y}}$ | $e_{x,y} = 0.5\, r_{E_{x,y}}$ | $h_{x,y} = h_x^2\, r_{A_{x,y}}$ | $e_{x,y} = (1-h_x^2)\, r_{E_{x,y}}$ | $h_{x,y} = \sqrt{h_x^2 h_y^2}\, r_{A_{x,y}}$ | $e_{x,y} = \sqrt{h_x^2(1-h_x^2)}\, r_{E_{x,y}}$ |
| $-1 < r_{A_{x,y}} < 1$<br>$r_{E_{x,y}} = 0$ | [2] $r_{P_{x,y}} = \sqrt{h_x^2 h_y^2}\, r_{A_{x,y}}$ | | [11] $r_{P_{x,y}} = 0.5\, r_{A_{x,y}}$ | | [20] $r_{P_{x,y}} = h_x^2\, r_{A_{x,y}}$ | | [29] $r_{P_{x,y}} = \sqrt{h_x^2(1-h_x^2)}\, r_{A_{x,y}}$ | |
| | $h_{x,y} = \sqrt{h_x^2 h_y^2}\, r_{A_{x,y}}$ | $e_{x,y} = 0$ | $h_{x,y} = 0.5\, r_{A_{x,y}}$ | $e_{x,y} = 0$ | $h_{x,y} = h_x^2\, r_{A_{x,y}}$ | $e_{x,y} = 0$ | $h_{x,y} = \sqrt{h_x^2(1-h_x^2)}\, r_{A_{x,y}}$ | $e_{x,y} = 0$ |
| $r_{A_{x,y}} = 0$<br>$-1 < r_{E_{x,y}} < 1$ | [3] $r_{P_{x,y}} = \sqrt{(1-h_x^2)(1-h_y^2)}\, r_{E_{x,y}}$ | | [12] $r_{P_{x,y}} = 0.5\, r_{E_{x,y}}$ | | [21] $r_{P_{x,y}} = r_{A_{x,y}} + (1-h_x^2)$ | | [30] $r_{P_{x,y}} = \sqrt{h_x^2(1-h_x^2)}\, r_{E_{x,y}}$ | |
| | $h_{x,y} = 0$ | $e_{x,y} = \sqrt{(1-h_x^2)(1-h_y^2)}\, r_{E_{x,y}}$ | $h_{x,y} = 0$ | $e_{x,y} = 0.5\, r_{E_{x,y}}$ | $h_{x,y} = 0$ | $e_{x,y} = (1-h_x^2)\, r_{E_{x,y}}$ | $h_{x,y} = 0$ | $e_{x,y} = \sqrt{h_x^2(1-h_x^2)}\, r_{E_{x,y}}$ |
| $-1 < r_{A_{x,y}} < 1$<br>$r_{E_{x,y}} = 1$ | [4] $r_{P_{x,y}} = \sqrt{h_x^2 h_y^2}\, r_{A_{x,y}} + \sqrt{(1-h_x^2)(1-h_y^2)}$ | | [13] $r_{P_{x,y}} = 0.5\left[r_{A_{x,y}} + 1\right]$ | | [22] $r_{P_{x,y}} = h_x^2\, r_{A_{x,y}} + (1-h_x^2)$ | | [31] $r_{P_{x,y}} = \sqrt{h_x^2(1-h_x^2)}\left[r_{A_{x,y}} + 1\right]$ | |
| | $h_{x,y} = \sqrt{h_x^2 h_y^2}\, r_{A_{x,y}}$ | $e_{x,y} = \sqrt{(1-h_x^2)(1-h_y^2)}$ | $h_{x,y} = 0.5\, r_{A_{x,y}}$ | $e_{x,y} = 0.5$ | $h_{x,y} = h_x^2\, r_{A_{x,y}}$ | $e_{x,y} = (1-h_x^2)$ | $h_{x,y} = \sqrt{h_x^2(1-h_x^2)}\, r_{A_{x,y}}$ | $e_{x,y} =$ |
| $r_{A_{x,y}} = 1$<br>$-1 < r_{E_{x,y}} < 1$ | [5] $r_{P_{x,y}} = \sqrt{h_x^2 h_y^2} + \sqrt{(1-h_x^2)(1-h_y^2)}\, r_{E_{x,y}}$ | | [14] $r_{P_{x,y}} = 0.5\left[1 + r_{E_{x,y}}\right]$ | | [23] $r_{P_{x,y}} = h_x^2 + (1-h_x^2)\, r_{E_{x,y}}$ | | [53] $r_{P_{x,y}} = \sqrt{h_x^2(1-h_x^2)}\left[r_{E_{x,y}} + 1\right]$ | |
| | $h_{x,y} = \sqrt{h_x^2 h_y^2}$ | $e_{x,y} = \sqrt{(1-h_x^2)(1-h_y^2)}\, r_{E_{x,y}}$ | $h_{x,y} = 0.5$ | $e_{x,y} = 0.5\, r_{E_{x,y}}$ | $h_{x,y} = h_x^2$ | $e_{x,y} = (1-h_x^2)\, r_{E_{x,y}}$ | $h_{x,y} = \sqrt{h_x^2(1-h_x^2)}$ | $e_{x,y} = \sqrt{h_x^2(1-h_x^2)}$ |
| $r_{A_{x,y}} = r_{E_{x,y}}$ | [6] $r_{P_{x,y}} = \left[\sqrt{h_x^2 h_y^2} + \sqrt{(1-h_x^2)(1-h_y^2)}\right] r_{A_{x,y}}$ | | [15] $r_{P_{x,y}} = r_{A_{x,y}} = r_{E_{x,y}}$ | | [24] $r_{P_{x,y}} = 1$ | | [32] $r_{P_{x,y}} = 2\sqrt{h_x^2(1-h_x^2)}\, r_{A_{x,y}}$ | |
| | $h_{x,y} = \sqrt{h_x^2 h_y^2}\, r_{A_{x,y}}$ | $e_{x,y} = \sqrt{(1-h_x^2)(1-h_y^2)}$ | $h_{x,y} = 0.5\, r_{A_{x,y}}$ | $e_{x,y} = 0.5\, r_{E_{x,y}}$ | $h_{x,y} = h_x^2\, r_{A_{x,y}}$ | $e_{x,y} = (1-h_x^2)\, r_{A_{x,y}}$ | $h_{x,y} = \sqrt{h_x^2(1-h_x^2)}\, r_{A_{x,y}}$ | $e_{x,y} = \sqrt{h_x^2(1-h_x^2)}\, r_{A_{x,y}}$ |
| $r_{A_{x,y}} = -\, r_{E_{x,y}}$ | [7] $r_{P_{x,y}} = \left[\sqrt{h_x^2 h_y^2} - \sqrt{(1-h_x^2)(1-h_y^2)}\right] r_{A_{x,y}}$ | | [16] $r_{P_{x,y}} = 0$ | | [25] $r_{P_{x,y}} = -1 + 2\, h_x^2\, r_{A_{x,y}}$ | | [33] $r_{P_{x,y}} = 0$ | |
| | $h_{x,y} = \sqrt{h_x^2 h_y^2}\, r_{A_{x,y}}$ | $e_{x,y} = -\sqrt{(1-h_x^2)(1-h_y^2)}$ | $h_{x,y} = 0.5\, r_{A_{x,y}}$ | $e_{x,y} = -0.5\, r_{A_{x,y}}$ | $h_{x,y} = h_x^2\, r_{A_{x,y}}$ | $e_{x,y} = -(1-h_x^2)\, r_{A_{x,y}}$ | $h_{x,y} = \sqrt{h_x^2(1-h_x^2)}\, r_{A_{x,y}}$ | $e_{x,y} = -\sqrt{h_x^2(1-h_x^2)}\, r_{A_{x,y}}$ |
| $r_{A_{x,y}} + r_{E_{x,y}} = 1$ | [8] $r_{P_{x,y}} = \sqrt{h_x^2 h_y^2}\, r_{A_{x,y}} + \sqrt{(1-h_x^2)(1-h_y^2)}\, r_{E_{x,y}}$ | | [17] $r_{P_{x,y}} = 0.5$ | | [26] $r_{P_{x,y}} = 1 + 2\, h_x^2\, r_{A_{x,y}} - h_x^2 - r_{A_{x,y}}$ | | [34] $r_{P_{x,y}} = \sqrt{h_x^2(1-h_x^2)}$ | |
| | $h_{x,y} = \sqrt{h_x^2 h_y^2}\, r_{A_{x,y}}$ | $e_{x,y} = \sqrt{(1-h_x^2)(1-h_y^2)}\, r_{E_{x,y}}$ | $h_{x,y} = 0.5\, r_{A_{x,y}}$ | $e_{x,y} = 0.5\,(1 - r_{A_{x,y}})$ | $h_{x,y} = h_x^2\, r_{A_{x,y}}$ | $e_{x,y} = (1-h_x^2)(1 - r_{A_{x,y}})$ | $h_{x,y} = \sqrt{h_x^2(1-h_x^2)}\, r_{A_{x,y}}$ | $e_{x,y} = \sqrt{h_x^2(1-h_x^2)} - h_{x,y}$ |
| $r_{A_{x,y}} + r_{E_{x,y}} = 1$ | [9] $r_{P_{x,y}} = \sqrt{h_x^2 h_y^2} + \sqrt{(1-h_x^2)(1-h_y^2)}$ | | [18] $r_{P_{x,y}} = 1$ | | [27] $r_{P_{x,y}} = 1$ | | [35] $r_{P_{x,y}} = 2\sqrt{h_x^2(1-h_x^2)}$ | |
| | $h_{x,y} = \sqrt{h_x^2 h_y^2}$ | $e_{x,y} = \sqrt{(1-h_x^2)(1-h_y^2)}$ | $h_{x,y} = 0.5$ | $e_{x,y} = 0.5$ | $h_{x,y} = h_x^2$ | $e_{x,y} = (1-h_x^2)$ | $h_{x,y} = \sqrt{h_x^2(1-h_x^2)}$ | $e_{x,y} = \sqrt{h_x^2(1-h_x^2)}$ |

Finally, the value that of the phenotypic correlation "approaches" as the covariance input "approaches" zero, shows that at the limit the phenotypic correlation can become one of its components, the coheritability or the coenvironmentability.

$$\lim_{C_{E_{x,y}} \to 0} \left( r_{P_{x,y}} \right) = \lim_{C_{E_{x,y}} \to 0} \left( \frac{C_{P_{x,y}}}{\sqrt{V_{P_x} V_{P_y}}} \right) = \lim_{C_{E_{x,y}} \to 0} \left( \frac{C_{A_{x,y}} + C_{E_{x,y}}}{\sqrt{V_{P_x} V_{P_y}}} \right) = h_{x,y}$$

and

$$\lim_{C_{A_{x,y}} \to 0} \left( r_{P_{x,y}} \right) = \lim_{C_{A_{x,y}} \to 0} \left( \frac{C_{P_{x,y}}}{\sqrt{V_{P_x} V_{P_y}}} \right) = \lim_{C_{A_{x,y}} \to 0} \left( \frac{C_{A_{x,y}} + C_{E_{x,y}}}{\sqrt{V_{P_x} V_{P_y}}} \right) = e_{x,y}$$

#### 7.6 The assumption that $\boldsymbol{r_{P_{x,y}}} = \boldsymbol{r_{A_{x,y}}}$

To assume that the phenotypic correlation is a suitable proxy for the genetic correlation. At the extreme case when $r_{P_{x,y}} = r_{A_{x,y}}$ , some implications can be deduced:

(A) The phenotypic variance of each trait is entirely composed of additive genetic variance and null environmental variance.
(B) This implies that the heritability of each trait is unity.
(C) The environmental correlation becomes undefined because its denominator would include zero.
(D) Thus,

$$r_{P_{x,y}} = \sqrt{h_x^2\, h_y^2}\, r_{A_{x,y}} + \sqrt{(1 - h_x^2)(1 - h_y^2)}\; r_{E_{x,y}}$$

$$r_{P_{x,y}} = \sqrt{1 \cdot 1}\, r_{A_{x,y}} = r_{A_{x,y}}$$

These are problematic assumptions, especially if applied in studies in the wild, and even in field tests of plant and animal breeding, where the environmental effects are generally strong and of importance.

#### 7.7 Disparity between the phenotypic and genetic correlations

Disparity D denotes the absolute difference of the phenotypic and genetic correlations (Willis et al. 1991).

$$D = |r_{P_{x,y}} - r_{A_{x,y}}|$$

This is a measure to assess the closeness of the values of the phenotypic and genetic correlations to one another.

The disparity index D is equivalent to the following relationship,

$$\left| \underbrace{\sqrt{h_x^2\, h_y^2}\, r_{A_{x,y}} + \sqrt{(1-h_x^2)(1-\ h_y^2)}\ r_{E_{x,y}}}_{r_{P_{x,y}}} - r_{A_{x,y}} \right|$$

which reduces to

$$D = \left| \left( \sqrt{h_x^2\, h_y^2}\ - 1 \right) r_{A_{x,y}}\ + \sqrt{(1-h_x^2)(1-\ h_y^2)}\ r_{E_{x,y}} \right|$$

##### 7.7.1 The domain of the Disparity Index

To properly address issues regarding the domain of the Disparity Index, it is useful to review the relationship maintained by the heritabilities and correlations, as presentred in table 7.7.1.

If $r_{P_{x,y}} = 1$, it means that either the coheritability or the coenvironmentability equals 1. Let's analyze each case.

**Case 1: If** $\boldsymbol{r_{P_{x,y}} = 1,\ and\ h_{x,y} = 1\ (\ e_{x,y} = 0\ )}$. If the coheritability equals 1 then the only combination of values allowed is that all $h_x^2 = 1, h_y^2 = 1,\ r_{A_{x,y}}\ = 1$ (causing the coenvironmentability to become zero, note if the heritabilities are 1, then $\sqrt{(1-h_x^2)(1-\ h_y^2)} = 0$). The disparity is $D = |1-1| = 0$.

**Case 2: If $\boldsymbol{r_{P_{x,y}} = 1,\ and\ h_{x,y} = 0\ (\ e_{x,y} = 1\ )}$.** In the case that the coheritability is zero, it is useful to analyze it by decomposing it into its constituents.

$$h_{x,y} = \sqrt{h_x^2\, h_y^2} \cdot r_{A_{x,y}} \qquad \text{for } 0 < h_{\blacksquare}^2 < 1\,, -1 < r_{A_{x,y}} < 1$$

$$= \frac{\sigma_{A_x}}{\sigma_{P_x}} \frac{\sigma_{A_y}}{\sigma_{P_y}} \cdot \frac{\sigma_{A_{x,y}}}{\sigma_{A_x}\sigma_{A_y}} \qquad \text{for } \sigma_{A_x} > 0\,,\ \sigma_{A_x} > 0\,, \sigma_{P_x} > 0,\ \sigma_{P_y} > 0$$

One notices that the numerator of the square root of the product of the heritabilities is also the denominator of the genetic correlation. If a heritability equals zero, it means that the genetic variance (the numerator) is zero, is also part of the denominator of the genetic correlation (see Section 7.5) Additional cases are presented in Table 7.7.1 .

An intuitive way to express this finding is presented below.

$$D = |r_{P_{x,y}} - r_{A_{x,y}}|$$

$$\left| r_{P_{x,y}} - r_{A_{x,y}} \right| < 1$$

$$-1 < r_{P_{x,y}} - r_{A_{x,y}} \leq 1$$

$$-1 + r_{A_{x,y}} < r_{P_{x,y}} < 1 + r_{A_{x,y}}$$

Note at this point that if $r_{A_{x,y}} = 0$, then $-1 < r_{P_{x,y}} < 1$

Let's analyze each bound separately.

When $r_{P_{x,y}} > 0$, the upper bound, $r_{P_{x,y}} < 1 + r_{A_{x,y}}$ cannot exceed unity, therefore to satisfy this condition $r_{A_{x,y}}$ must have negative values.

When $r_{P_{x,y}} < 0$, the lower bound, $r_{P_{x,y}} > -1 + r_{A_{x,y}}$, cannot be lower than -1, therefore $r_{A_{x,y}}$ should have positive values.

To summarize, the bounds the bounds of the Disparity Index is $-1 < D < 1$, or more explicitly

$$D = \begin{cases} r_{P_{x,y}} - r_{A_{x,y}} < 1 & for\ r_{A_{x,y}} < 0\,,\ r_{P_{x,y}} > 0 \\ r_{P_{x,y}} & for\ r_{A_{x,y}} = 0 \\ r_{P_{x,y}} - r_{A_{x,y}} > -1 & for\ r_{A_{x,y}} > 0, r_{P_{x,y}} < 0 \end{cases}$$

| $r_{P_{x,y}}$ | $\sqrt{h_x^2}$ | $\sqrt{h_y^2}$ | $r_{A_{x,y}}$ | $h_{x,y}$ | $\sqrt{(1-h_x^2)}$ | $\sqrt{(1-h_y^2)}$ | $r_{E_{x,y}}$ | $e_{x,y}$ | $D = \lvert r_{P_{x,y}} - r_{A_{x,y}} \rvert$ |
|---|---|---|---|---|---|---|---|---|---|
| **1** | $\sqrt{\mathbf{1}}$ | $\sqrt{\mathbf{1}}$ | **1** | $h_{x,y} = 1$ | $\sqrt{1-\mathbf{1}}$ | $\sqrt{1-\mathbf{1}}$ | $r_{E_{x,y}}$ | $e_{x,y} = 0$ | **0** = \|**1** - **1**\| |
| **1** | $\sqrt{\mathbf{1}}$ | $\sqrt{\mathbf{1}}$ | $r_{A_{x,y}}$ | $h_{x,y} = r_{A_{x,y}}$ | $\sqrt{1-\mathbf{1}}$ | $\sqrt{1-\mathbf{1}}$ | $r_{E_{x,y}}$ | $e_{x,y} = 0$ | $\lvert 1 - r_{A_{x,y}} \rvert$ |
| **1** | $\sqrt{\mathbf{0.5}}$ | $\sqrt{\mathbf{0.5}}$ | **1** | $h_{x,y} = 0.5$ | $\sqrt{1-\mathbf{0.5}}$ | $\sqrt{1-\mathbf{0.5}}$ | 1 | $e_{x,y} = 0.5$ | **0** = \|**1** - **1**\| |
| **-1** | $\sqrt{\mathbf{1}}$ | $\sqrt{\mathbf{1}}$ | **-1** | $h_{x,y} = -1$ | $\sqrt{1-\mathbf{1}}$ | $\sqrt{1-\mathbf{1}}$ | $r_{E_{x,y}}$ | $e_{x,y} = 0$ | **0** = \|**-1** - (**-1)**\| |
| **-1** | $\sqrt{\mathbf{0.5}}$ | $\sqrt{\mathbf{0.5}}$ | **-1** | $h_{x,y} = -0.5$ | $\sqrt{1-\mathbf{0.5}}$ | $\sqrt{1-\mathbf{0.5}}$ | -1 | $e_{x,y} = -0.5$ | **0** = \|**-1** -(- **1)**\| |
| **0** | $\sqrt{\mathbf{1}}$ | $\sqrt{h_y^2}$ | **0** | $h_{x,y} = 0$ | $\sqrt{1-\mathbf{1}}$ | $\sqrt{1-h_y^2}$ | $r_{E_{x,y}}$ | $e_{x,y} = 0$ | **0** = \|**0** - **0**\| |
| **0** | $\sqrt{h_x^2}$ | $\sqrt{h_y^2}$ | **0** | $h_{x,y} = 0$ | $\sqrt{(1-h_y^2)}$ | $\sqrt{1-h_y^2}$ | 0 | $e_{x,y} = 0$ | **0** = \|**0** - **0**\| |
| **0** | $\sqrt{\mathbf{0.5}}$ | $\sqrt{\mathbf{0.5}}$ | **1** | $h_{x,y} = 0.5$ | $\sqrt{1-\mathbf{0.5}}$ | $\sqrt{1-\mathbf{0.5}}$ | -1 | $e_{x,y} = -0.5$ | **1** = \|**0** - **1**\| |
| **0** | $\sqrt{\mathbf{0.5}}$ | $\sqrt{\mathbf{0.5}}$ | **-1** | $h_{x,y} = -0.5$ | $\sqrt{1-\mathbf{0.5}}$ | $\sqrt{1-\mathbf{0.5}}$ | 1 | $e_{x,y} = 0.5$ | **1** = \|**0** - (**-1)**\| |
| **0** | $\sqrt{\mathbf{0.5}}$ | $\sqrt{\mathbf{0.5}}$ | $r_{A_{x,y}}$ | $h_{x,y} = 0.5\, r_{A_{x,y}}$ | $\sqrt{1-\mathbf{0.5}}$ | $\sqrt{1-\mathbf{0.5}}$ | $r_{E_{x,y}} = -r_{A_{x,y}}$ | $e_{x,y} = -0.5\, r_{A_{x,y}}$ | $\lvert \boldsymbol{r_{A_{x,y}}} \rvert = \lvert 0 - r_{A_{x,y}} \rvert$ |
| 1 | $\sqrt{\mathbf{0}}$ | $\sqrt{\mathbf{0}}$ | $r_{A_{x,y}}$ | $h_{x,y} = 0$ | $\sqrt{1-\mathbf{0}}$ | $\sqrt{1-\mathbf{0}}$ | 1 | $e_{x,y} = 1$ | $\lvert \mathbf{1} - r_{A_{x,y}} \rvert$ |
| -1 | $\sqrt{\mathbf{0}}$ | $\sqrt{\mathbf{0}}$ | $r_{A_{x,y}}$ | $h_{x,y} = 0$ | $\sqrt{1-\mathbf{0}}$ | $\sqrt{1-\mathbf{0}}$ | -1 | $e_{x,y} = -1$ | $\lvert -\mathbf{1} - r_{A_{x,y}} \rvert$ |
| 1 | $\sqrt{\mathbf{0}}$ | $\sqrt{\mathbf{0}}$ | **-1** | $h_{x,y} = 0$ | $\sqrt{1-\mathbf{0}}$ | $\sqrt{1-\mathbf{0}}$ | 1 | $e_{x,y} = 1$ | 2 = \| 1 - (-1) \| |
| -1 | $\sqrt{\mathbf{0}}$ | $\sqrt{\mathbf{0}}$ | **1** | $h_{x,v} = 0$ | $\sqrt{1-\mathbf{0}}$ | $\sqrt{1-\mathbf{0}}$ | -1 | $e_{x,y} = -1$ | 2 = \| -1 - 1 \| |

**Table 7.7.1** Disparity index measures under varying values of phenotypic and genetic correlations. Note the last four rows, which show incongruence relationship between the genetic and the environmental correlations, produce invalid disparity indices.

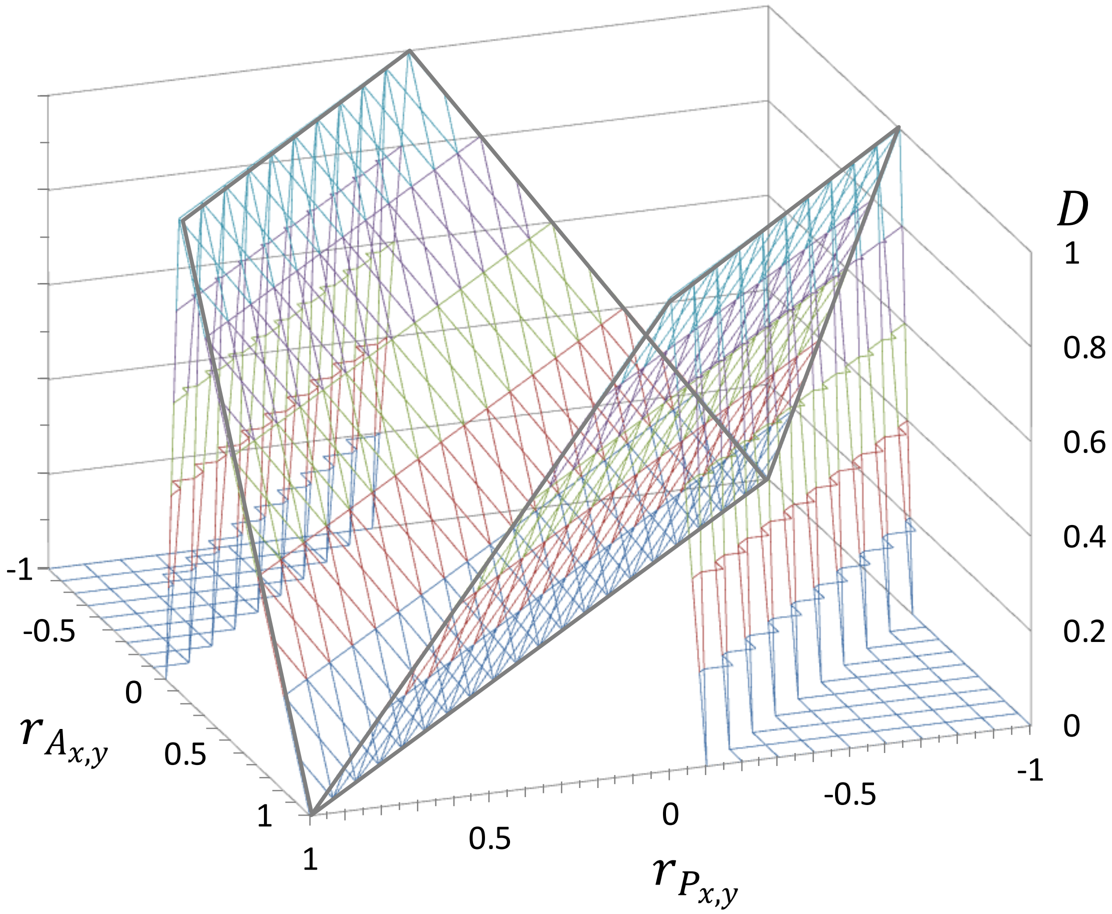

Figure 7.7.1 Surface graph depicting the Disparity Index $D$ as a function of the phenotypic correlation $r_{P_{x,y}}$ and the genetic correlation $r_{A_{x,y}}$.

##### 7.7.2 Distribution of the Disparity Index

The distribution of D can be modelled as the absolute difference of two uniform variables, which results in the triangular distribution. Our purpose now is to find the cumulative density function (CDF) and the probability density function (PDF) of $D$.

Let $P$ and $A$ be random variables distributed identically and independently as standard uniform, such that

$$P \sim U(0,1)\,, \quad f_P(p) = 1\,, \quad 0 < p < 1\,,$$

$$A \sim U(0,1)\,, \quad f_A(a) = 1\,, \quad 0 < a < 1\,,$$

Define the new variable $D = |\,P - A\,|$.

The subset within the unit square that is between the two lines represents the area where the event $\{\,|P - A| \leq d\,\}$ for $0 \leq d \leq 1$ occurs. The area of the unit square is 1. The dimension of the subset (a hexagon) is obtained by subtracting the area of the two triangles, each encompassing $\frac{(1-d)^2}{2}$.

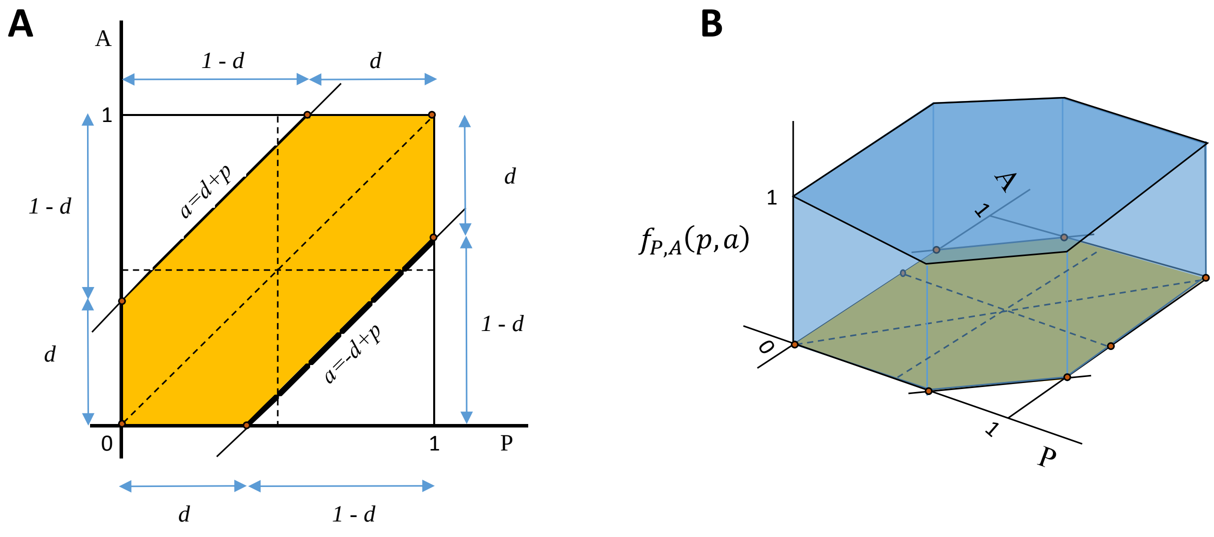

**Figure 7.7.2-1** Event $\{\,|P - A| \leq d\,\}$. (A) Area representing the occurrence of the event. (B) Volume representing the probability of the event.

Within this region, the probability is the volume within the two lines (yellow region in Figure 7.7.2-1A). The overall volume is a cube with dimension 1x1x1=1 is obtained by multiplying by 1 since that is the uniform pdf in each point $(P, A)$ (Figure 7.7.2-2).

The cumulative density function of $D$

Is obtained by determining the probability of $D$ taking values less than $d$

$$F_D(d) = Pr\{D \le d\} = Pr\{\{||P - A|\}| \le d\}$$

$$= 1 - 2\frac{(1-d)^2}{2} = 2d - d^2$$

$$= d(2-d) \qquad \text{for } 0 \le d \le 1$$

$$F_D(d) = \begin{cases} 0 & d < 1 \\ d(2-d) & 0 \le d \le 1 \\ 0 & d > 1 \end{cases}$$

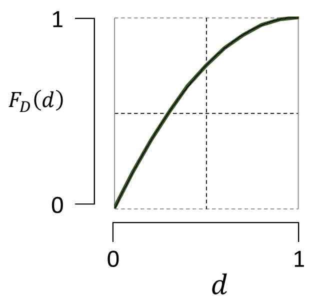

The probability density function of $D$

Is obtained by differentiating the cumulative function $F_D$ with respect to $d$.

$$f_D(d) = \begin{cases} 0 & d < 0 \\ \frac{\mathrm{d}(F_D)}{\mathrm{d}d} = 2 - 2d & 0 \le d \le 1 \\ 0 & d > 1 \end{cases}$$

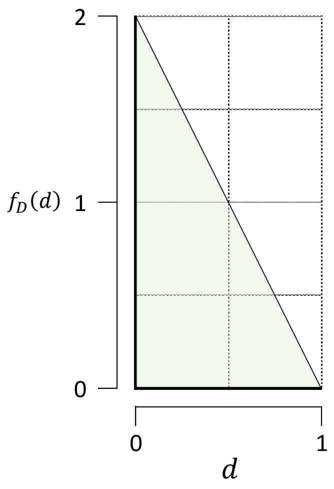

##### 7.7.3 A simulation model to generate disparity data

The disparity between the phenotypic and genetic correlation, as we have seen, can only obtain values between zero and one (Section 7.7.1).

To generate simulated data of the disparity index, a model can be devised as the distribution of a simple difference of two standard uniform random variables. Here a convolution method is employed.

Let $X_1$ and $X_2$ be random variables distributed identically and independently as standard uniform, such that

$$X_1 \sim U(0,1)\,, \quad f_{X_1}(x_1) = 1\,, \quad 0 < x_1 < 1$$

$$X_2 \sim U(0,1)\,, \quad f_{X_2}(x_2) = 1\,, \quad 0 < x_2 < 1$$

The joint density of $(X_1, X_2)$ is

$$f_{X_1,X_2}(x_1, x_2) = f_{X_1}(x_1) \cdot f_{X_2}(x_2) = \begin{cases} 1 & for\ \ 0 \le x_1 \le 1\ ,\ \ 0 \le x_2 \le 1 \\ 0 & elsewhere \end{cases}$$

Let's define $= X_1 - X_2$ , such that the value of $x_1 = z + x_2$

$$0 \le x_1 \le 1$$

$$0 \le z + x_2 \le 1$$

$$-x_2 \le \ z \ \le 1 - x_2$$

which can be replaced in the joint density pdf, as follows.

$$f_{X_1,X_2}(x_1, x_2) = \ f_{X_1,X_2}(\,z + x_2\,,\ x_2) = \begin{cases} 1 & for\ \ 0 \le x_2 \le 1\ ,\ \ z \ge -x_2\ ,\ \ z + x_2 \le 1 \\ 0 & elsewhere \end{cases}$$

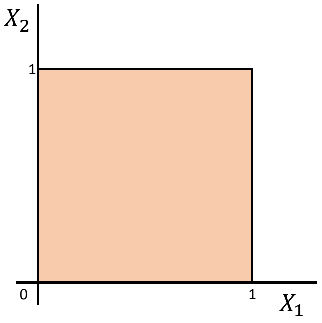

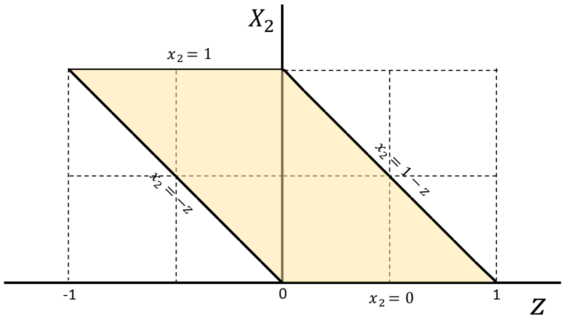

| $X_1$ | $X_2$ |
|---|---|
| 0 | 0 |
| 0 | 1 |
| 1 | 0 |
| 1 | 1 |

| $Z$ $X_1 - X_2$ | $X_2$ |
|---|---|
| 0 | 0 |
| -1 | 1 |
| 1 | 0 |
| 0 | 1 |

The tables above help corroborate that this is a 1-to-1 transformation.

The probability density function of $z$ is obtained by solving the convolution integrals

$$f_Z(z) = \int_{-\infty}^{+\infty} f_{X_1,X_2}(\, z + x_2 \,,\, x_2)\, dx_2 = \int_{-\infty}^{+\infty} f_{X_1}(\, z + x_2\,) \cdot f_{X_2}(x_2)\, dx_2 = \int_{-\infty}^{+\infty} f_{X_1}(\, z + x_2 \,)\, dx_2$$

$$f_Z(z) = \begin{cases} 0 & z < -1 \\ \int_{-z}^{1} 1\, dx_2 = \ [1 - (-z)] = 1 + z & -1 \le z < 0 \\ 1 & z = 0 \\ \int_{0}^{1-z} 1\, dx_2 = [1 - z - 0] = 1 - z & 0 < z \le 1 \\ 0 & z > 0 \end{cases}$$

And the cumulative density function of $z$ is

$$F_Z(z) = \ Pr\{\, Z \le z \,\} \ = Pr\{\, X_1 - X_2 \le z \,\}$$

$$F_Z(z) = \begin{cases} 0 & z < -1 \\ \int_{x_1=0}^{z} \int_{x_2=x_1-z}^{1} 1 \, dx_2 \, dx_1 = \frac{z^2}{2} + z + \frac{1}{2} & -1 \le z < 0 \\ 0.5 & z = 0 \\ 1 - \int_{x_1=z}^{1} \int_{x_2=0}^{x_1-z} 1 \, dx_2 \, dx_1 = -\frac{z^2}{2} + z + \frac{1}{2} & 0 < z \le 1 \\ 0 & z > 0 \end{cases}$$

The cumulative density function of $z$

$$F_Z(z) = \begin{cases} 0 & z < -1 \\ \frac{z^2}{2} + z + \frac{1}{2} & -1 \le z < 0 \\ 0.5 & z = 0 \\ -\frac{z^2}{2} + z + \frac{1}{2} & 0 < z \le 1 \\ 0 & z > 0 \end{cases}$$

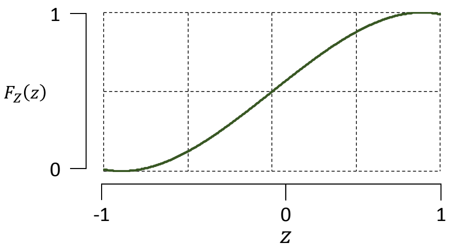

The probability density function of $z$

$$f_Z(z) = \begin{cases} 0 & z < -1 \\ 1 + z & -1 \le z < 0 \\ 1 & z = 0 \\ 1 - z & 0 < z \le 1 \\ 0 & z > 0 \end{cases}$$

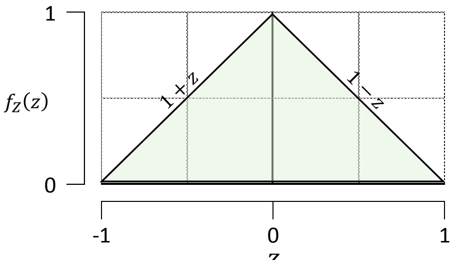

##### 7.7.4 The ratio of the genetic correlation over the phenotypic correlation

The ratio $k$ between the genetic correlation and the phenotypic correlation can be deduced if the phenotypic correlation and the disparity index are known. This ratio is another parameter that reveal the degree of difference between the genetic and phenotypic correlations.

Figure 7.7.4 illustrates how the ratio behave as a function of the phenotypic correlation and the disparity index.

(A) When the genetic and phenotypic correlation have the same sign, the ratio oscillates between zero and is 1 (or 100%). The latter is equivalent that $r_{A_x} = r_{P_x}$, and its line correspond to the base of the triangle.

(B) When the genetic and phenotypic correlation have different sign, the ratio goes from zero to extreme values (< -900%). A ratio equal to -200% indicates that the henetic correlation value is the double of the phenotypic correlation, $|r_{A_x}| = 2|r_{P_x}|$, the sign indicating the genetic correlation has a different sign than the phenotypic correlation.

(C) When $r_{A_x} = 0$, then the disparity is $D = r_{P_x}$ and the ratio is zero (**0 %**).

(D) For a given disparity value there are different ratios that can be associated to it , dependning on the value of $r_{P_x}$ .

(E) For a given value of $r_{P_x}$ , there is a continuum of ratios that depend upon the $D$ value.

(F) As the phenotypic correlation approaches zero, irrespective of the disparity value, the ratio assumes extreme values, showing a marked difference between $r_{A_x}$ and $r_{P_x}$.

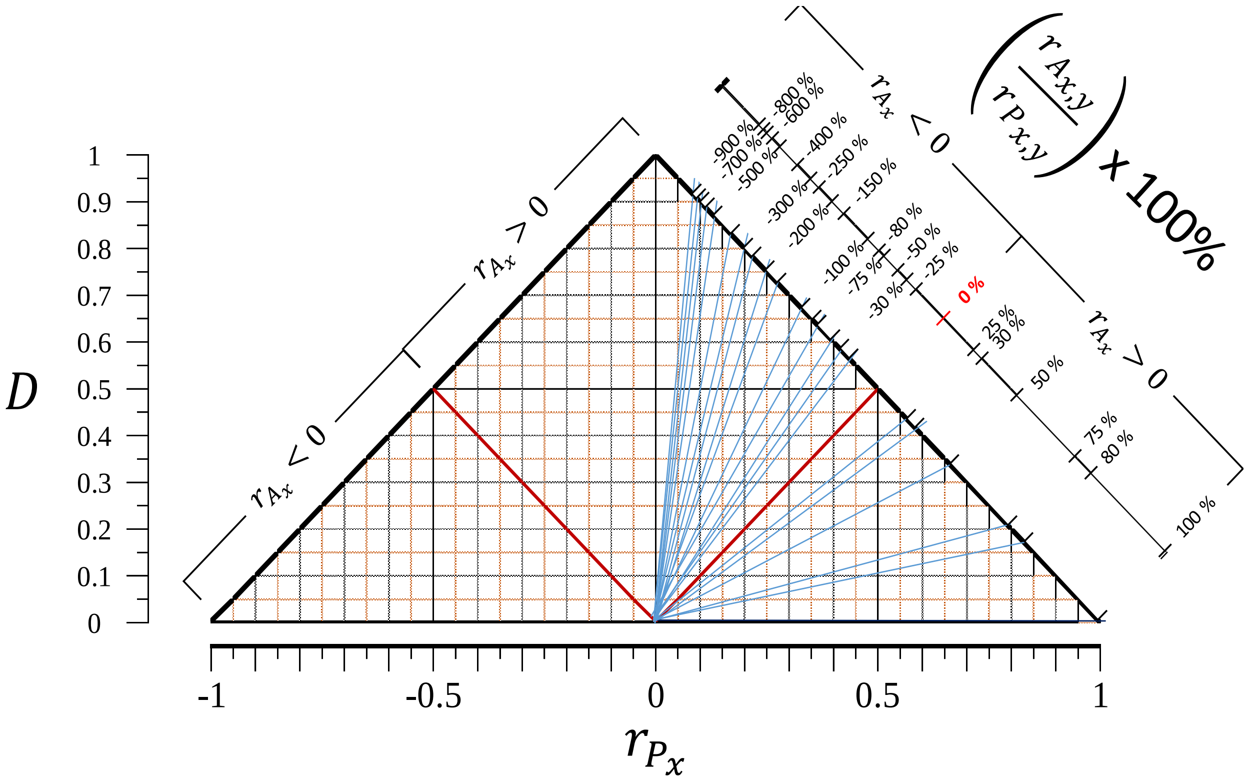

**Figure 7.7.4-1** Ratio of the genetic correlation over the phenotypic correlation as a function of the disparity index and the phenotypic correlation. Only one size is labeled. A mirror image of the lines apply at the left side of the triangle where the $r_{P_x} < 0$. The sign associated to the percentage indicates that $r_{A_x}$ has a different sign than the phenotypic correlation. At the right side ($r_{P_x} > 0$), negative sign percentages indicate that the $r_{A_x}$ is negative. At the left side ($r_{P_x} < 0$), negative sign indicates that $r_{A_x}$ is positive.

**Table 7.7.4-1**

Values of the ratio $k$, and parameters of the line that starts at the origin and reaches the side of the triangle at the point ( $r_{P_k}$ , $D_k$) having a slope $m$.

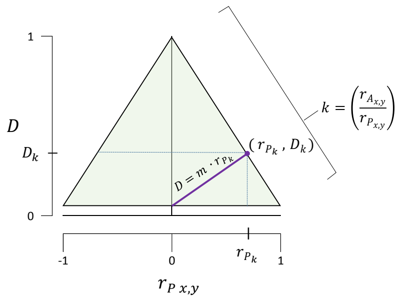

| $k = \left(\frac{r_{A_{x,y}}}{r_{P_{x,y}}}\right)$ | $m$ | $r_{P_{x,y\,k}}$ | $D_k$ | $k \cdot$ [illegible] |
|---|---|---|---|---|
| -10 | 11 | 0.083 | 0.917 | -1[illegible] |
| -9 | 10 | 0.091 | 0.909 | [illegible] |
| -8 | 9 | 0.100 | 0.900 | [illegible] |
| -7 | 8 | 0.111 | 0.889 | [illegible] |
| -6 | 7 | 0.125 | 0.875 | [illegible] |
| -5 | 6 | 0.143 | 0.857 | [illegible] |
| -4 | 5 | 0.167 | 0.833 | [illegible] |
| -3 | 4 | 0.200 | 0.800 | [illegible] |
| -2.5 | 3.5 | 0.222 | 0.778 | [illegible] |
| -2 | 3 | 0.250 | 0.750 | [illegible] |
| -1.5 | 2.5 | 0.286 | 0.714 | [illegible] |
| -1 | 2 | 0.333 | 0.667 | [illegible] |
| -0.8 | 1.8 | 0.357 | 0.643 | [illegible] |
| -0.75 | 1.75 | 0.364 | 0.636 | [illegible] |
| -0.5 | 1.5 | 0.400 | 0.600 | [illegible] |
| -0.3 | 1.3 | 0.435 | 0.565 | [illegible] |
| -0.25 | 1.25 | 0.444 | 0.556 | [illegible] |
| **0** | **1** | **0.500** | **0.500** | |
| 0.25 | 0.75 | 0.571 | 0.429 | |
| 0.3 | 0.7 | 0.588 | 0.412 | |
| 0.5 | 0.5 | 0.667 | 0.333 | |
| 0.75 | 0.25 | 0.800 | 0.200 | |

See below for a description of the algorithm used to create Figure 7.7.4-1.

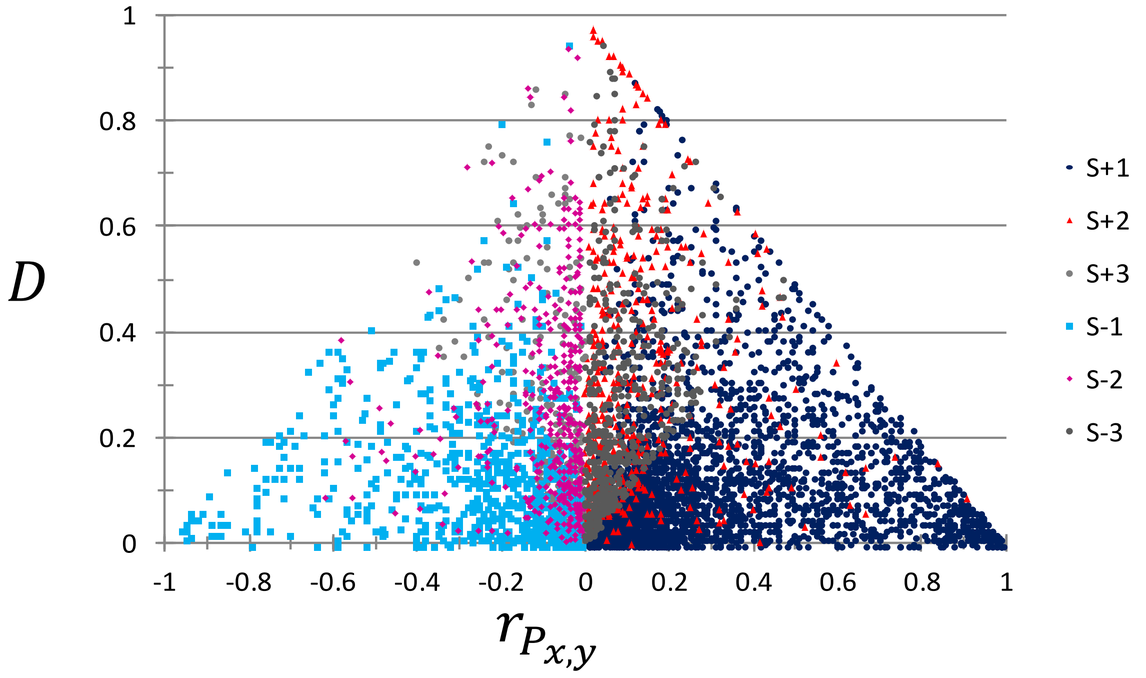

**Figure 7.7.4-2** Disparity data derived from genetic and phenotypic correlations collected from an extensive literature review $(n = 6280)$. A first impression points out to the fact that in the majority of the cases, the value of the genotypic correlation differ quite significantly from the value of the phenotypic correlation.

Algorithm to derive the line corresponding to a given ratio $k$ of the genetic correlation over the phenotypic correlation

**.1** Select a given value of the ratio $k = \left(\frac{r_{A_{x,y}}}{r_{P_{x,y}}}\right)$. If the phenotypic correlation has the same sign as the genetic correlation, then the maximum $k$ is 1. If the phenotypic correlation and the genetic correlation differ in sign, then $k$ can be from zero to 10 (above 10 is possible, but the value of $r_P$ will be extremenly small). For example, choosing a value of 5 means that $r_A$ is 5 times larger than , or that $r_A$ is 500% the value of $r_P$.

**.2** Determine the slope of the line $m = 1 - k$.

**.3** Determine the phenotypic correlation value $r_{P_k} = \frac{1}{1+|m|}$. (If working with the negative phenotypic correlation, multiply by -1.

**.4** Determine the disparity index $D_k$ .

**.5** Trace the line from the origin to the point $\left( r_{P_k} ,\ D_k \right)$. The line covers all the combinations of $\left( r_{P_{x,y}} ,\ D \right)$ that have a ratio equal to $k$.

### 8 Selection

The term $\sqrt{h_x^2\, h_y^2}\, r_{A_{x,y}}$ is a measure of coinheritance of a pair of phenotypic traits that is due to shared genetics. This expression has named *coheritability* by Nei (1960), and further studied by others (Searle 1961, Falconer 1960). It is in the last edition of *Principles of Quantitative Genetics* by Falconer and Mackay (1996) in which the term coheritability was reintroduced. The name is derived from the context of correlated response to selection. The formula for the expected response $R_X$ of a progeny in terms of phenotypic gain in trait $x$ due to selection of parents based on the same trait is

$$R_x = i\,(\,h_x^2\,)\,\sigma_{P_X} \qquad [\,8-1\,]$$

where $i$ is the intensity of selection, and $\sigma_{P_x}$ is the phenotypic standard deviation of trait $x$. In an analogous way, the expected response of correlated trait $x$ when parental selection is based one trait $y$ is:

$$R_{X|Y} = i\left(\sqrt{h_x^2 h_y^2}\, r_{A_{x,y}}\right)\sigma_{P_X} \qquad [\,8-2\,]$$

The term in brakets is the coheritability, which functions in an analogous manner as the heritability in equation [ 8-1 ].

Yet despite its implicit use, very little attention has been paid to understand the statistical and mathematical properties of coheritability. There is a persistent confusion in the scientific literature of what this term means or how should be calculated. Denoted as *coheritability* some reports present expressions that do not possess any consistent theoretical background, such as the mere ratio of additive genetic to phenotypic covariances $\frac{C_A}{C_P}$ (de Reggi 1972, Janssens 1979). The simple ratio $\frac{C_A}{C_P} = \frac{\sqrt{V_{Ax}V_{Ay}}r_A}{\sqrt{V_{Px}V_{Py}}r_P} = \frac{\sqrt{h_x^2\, h_y^2}\, r_{A_{xy}}}{r_P} = \frac{h_{x,y}}{r_P} \neq h_{xy}$, clearly is not a recognizable coheritability form (its bounds go from zero to infinity).

Most quantitative phenotypes are polygenic, and for these traits, selection is likely to act on many preexisting genetic variants of small effect. Detecting so called polygenic selection is challenging because selection acts on multiple loci simultaneously.

For the sake of completeness and to introduce symbols denoting estimators to be used later, this section starts with a brief explanation of the theory of mass selection and correlated response.

#### 8.1 Realized heritability

The mass selection method aims in choosing a subset of individuals from a original population to breed and generate the progeny (or offspring) generation. Individuals are ordered my their phenotypic value. Individuals whose phenotypic value are above a pre-specified value (i.e., truncation point), generally at an extreme of the distribution, are selected. Therefore, individuals are selected solely strictly by their rank judge by their phenotypic values.

Based on the assumption that the distribution of phenotypic values of the population follows a normal distribution $N(\bar{P}_o\,,\ \sigma_P^2)$, the observed mean phenotypic value of the original distribution $(\bar{P}_o\,)$, and the mean phenotypic value of the subset of individuals chosen $(\bar{P}_s)$ differ in the amount called the selection differential

$$S = (\bar{P}_s - \bar{P}_o).$$

$S$ depends on the percentage $k$ of individuals of the original population selected as parents, and the phenotypic standard deviation of the original population. The expected mean value of the selected individuals above the truncation point $t$ $(t > 0)$ is

$$E(x_s) = E(P \mid P\ >\ t\,) = \frac{h}{k}$$

and the variance of the selected individuals is

$$Var(P_s) = Var(P\ |P > t) = 1 - \frac{h}{k}\left(\frac{h}{k} - t\right)$$

where $h$ is the height (i.e. ordinate) of the standard normal curve at point $t$.

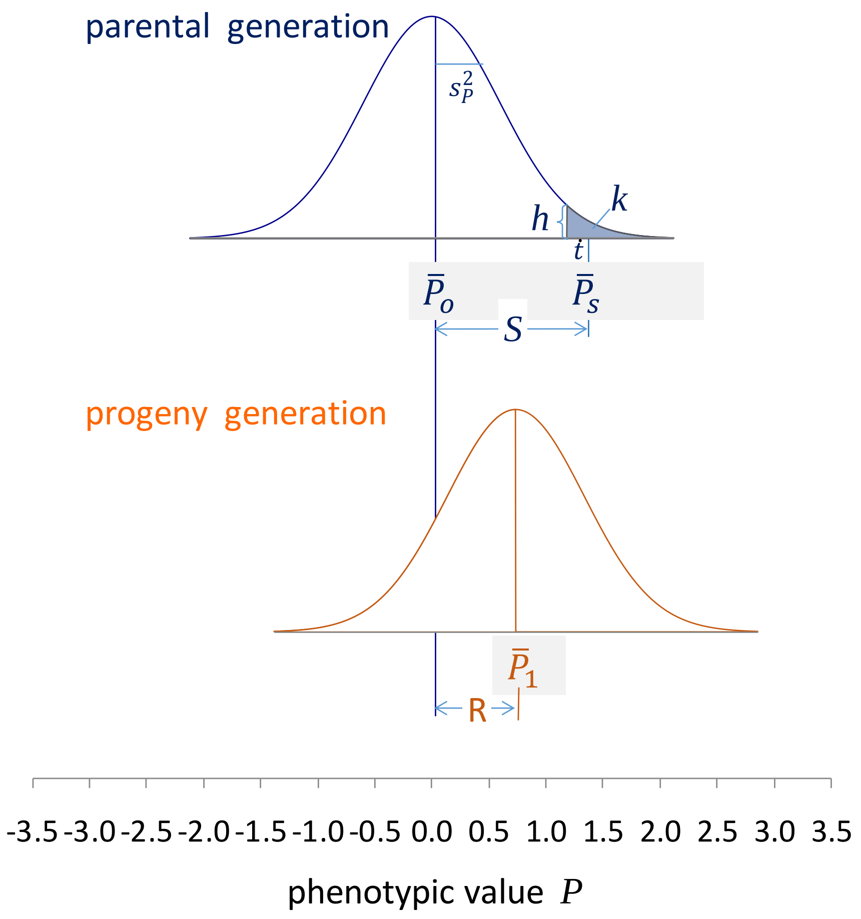

The expected response ( $R$ ) to mass selection for a desired trait is

$$R = h^2 S$$

When the progeny has been obtained and its mean phenotypic value $(\bar{P}_1)$ determined, the realized response to selection is calculated as

$$R_{realized} = \bar{P}_1 - \bar{P}_o$$

Given than the both the selection differential $S$ and the realized response $R_{realized}$ are known, it is then possible to determine the realized heritability of the trait as follows:

$$h^2_{realized} = \frac{R_{realized}}{S}$$

In practice, the cumulative response is plotted against the cumulated selection differential (Falconer and MacKay 1996, page 197ss), a a simple regression line is fitted. The slope of this line $(b)$ is a function of the realized heritability, such that $b = v \cdot h_r^2$. The coefficient $v = 1$ if the trait is measured in all individuals irrespective of their sex. If the measurements are obtained from either male or female, then $v = 0.5$, and the realized heritability in this case is obtained by doubling the slope.

#### 8.2 Realized coheritability

Using the correlated response of trait $y$ when selection was directed to trait $x$, a similar rationale can be used to determined the realized bivariate measures of genetic relationship.

By performing mass selection on an original population, based on the phenotypic value of trait $x$, using a specified selection differential $S_x$, both a direct response to selection for trait x $(R_x)$ as well as a correlated response in trait $y$ $(CR_{X|Y})$ are obtained.

( A ) If one round of selection has been performed, then the realized coheritability is the ratio of the correlated response on the selection differential,

$$h_{x,y\ realized} = \frac{CR_{X|Y}}{S_x}$$

If several rounds of selection have been done, then one can plot cumulative correlated response of trait $y$ on the cumulative selection differential of trait $x$, an a liner regression model fitted. The slope parameter $b$ (which can be positive or negative) becomes a function of the coheritability, such that:

$$h_{x,y\ realized} = \begin{cases} b & \text{if analyses employ measurements of all individuals irrespective of sex} \\ 2b & \text{if analyses employ measurements of either male or female individuals} \end{cases}$$

( B ) In the case of double selection experiments, in which one group of individuals is directly selected for the primary trait $x$ yielding $R_x$, and trait $y$ is assessed as a secondary (correlated) trait as $CR_{X|Y}$, and another group is directly selected for $y$ $(R_x)$, and $x$ becomes the secondary trait $CR_{Y|X}$. Both are plotted and the coheritabilities calculated as expounded above. It is expected that the two estimates should be numerically close, provided the theory of correlated response is adequate to describe the results, and no discernible asymmetry is found in the direct and correlated responses. The estimate of the realized coheritability is the geometric mean of the slope parameters of the regressions.

$$h_{x,y\ realized} = \begin{cases} \pm v \cdot \sqrt{\frac{|CR_{X|Y}}{|\ S_{C_Y}} \frac{CR_{Y|X}|}{S_{C_X}\ |}} & \text{for one single round of double selection} \\ \pm v \cdot \sqrt{b_{X|Y}\ b_{Y|X}} & \text{for several rounds of double selection} \end{cases}$$

where $S_{C_X}$ and $S_{C_Y}$ are the selection differentials used for direct selection of trait $x$ and $y$ respectively. The sign depends on the relationship observed in the plot of the data. Again, $v = 1$ when individuals are selected irrespective of sex, and $v = 0.5$ if measurements come from only one sex.

( C ) It both the realized heritabilities and genetic correlations (see below) have been estimated, then

$$h_{x,y\ realized} = \sqrt{h^2_{x\ realized}\ h^2_{y\ realized}}\ r_{A_{x,y\ realized}}$$

All the formulas presented are subject to large experimental variation and must be taken to indicate tendency rather than magnitude.

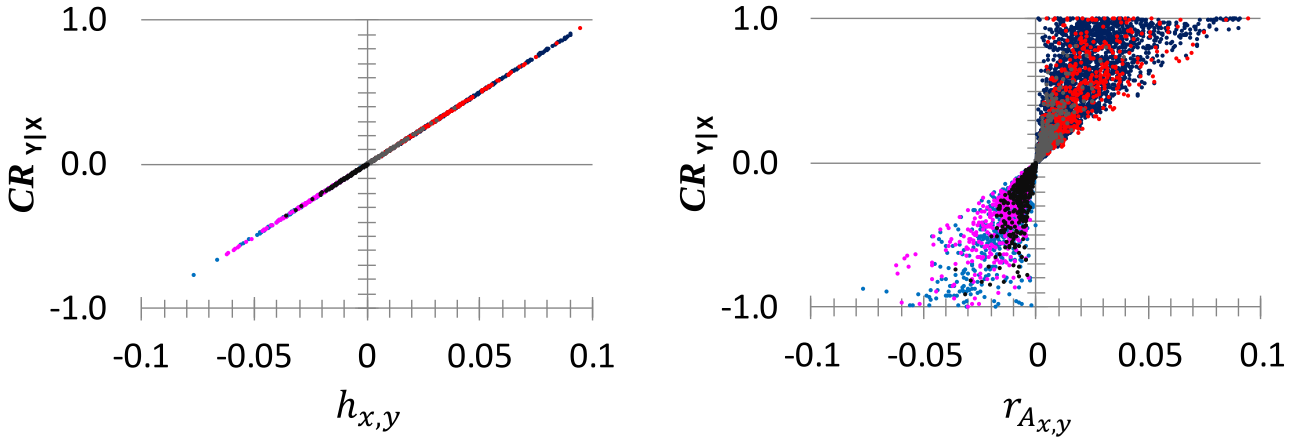

**Figure 8.2** The coheritability and the genetic correlation in relation to their capability to predict the correlated response to selection. The intensity of selection was set to an intensity of selection $i = 0.1$ and the phenotypic variance of the correlated trait standardized to $1$ ($n = 6288$).

#### 8.3 Realized genetic correlation

The realized genetic correlation between two traits is,

$$r_{A_{x,y\ realized}} = \begin{cases} \dfrac{CR_{Y|X}}{R_x} & \text{for single round of selection} \\ \sqrt{\dfrac{CR_{X|Y}}{R_x}\dfrac{CR_{Y|X}}{R_y}} & \text{for double selection experiment} \end{cases}$$

Note that the value of the correlated response may be either positive or negative.

#### 8.4 Relative efficiency of indirect selection with respect to direct selection

Observation of the response to selection of correlated traits may lead to consider that in certain circumstances it would be feasible to obtain more rapid and efficient advance of a desired character when applying selection for a secondary trait $y$ which is correlated to $x$. This means that to improve character $x$, selection should be applied to another (secondary) character $y$ and achieve progress through the correlated response of character $x$.

Indirect selection is advantageous under special conditions.

Let $R_x = i_x\, h_x^2\, \sigma_{P_x}$ be the direct response of the desired trait when selection is applied directly on character $x$.

Let $CR_{x|y} = i_y\, h_{xy}\, \sigma_{P_x}$ be the correlated response of character $x$ resulting from selection applied to secondary character $y$.

The relative efficiency (RE) of selection is the value of indirect selection relative to that of direct selection, and could be expressed as the ratio of expected responses, as follows

$$RE = \frac{CR_{x|y}}{R_x} = \frac{i_y\, h_{x,y}\, \sigma_{P_x}}{i_x\;\, h_x^2\;\, \sigma_{P_x}} = \frac{i_Y\, h_{x,y}}{i_x\;\, h_x^2}$$

Lerner and Cruden (1948 ) defined as relative efficiency of indirect selection the expression $\frac{h_Y r_A}{h_X}$ which is equivalent to

$$RE = \frac{h_{x,y}}{h_X^2}$$

this equation allows to determine conditions of when indirect selection of $x$ when selecting for $y$ would be greater than the direct response to selection for $x$ (see Figure 8.4),

$$i_Y\, h_{x,y} > i_x\;\, h_x^2$$

$$i_Y\, \sqrt{h_x^2\, h_y^2}\, r_{A_{xy}} > i_x\;\, h_x^2$$

$$i_Y\, \sqrt{h_y^2}\, r_{A_{xy}} > i_x\;\, \sqrt{h_x^2}$$

Inspection of this relationship shows that indirect selection would be greater if the heritability $h_y^2$ is greater than $h_x^2$, and the genetic correlation $r_{A_{xy}}$ is high. Cases like this would occur if trait $x$ is difficult to measure with precision but $y$ is not.

If the selection intensity is much greater for $y$ than $x$ would also result in a greater response of indirect selection. This would apply if $y$ were measurable in both sexes by $x$ measurable in only one sex.

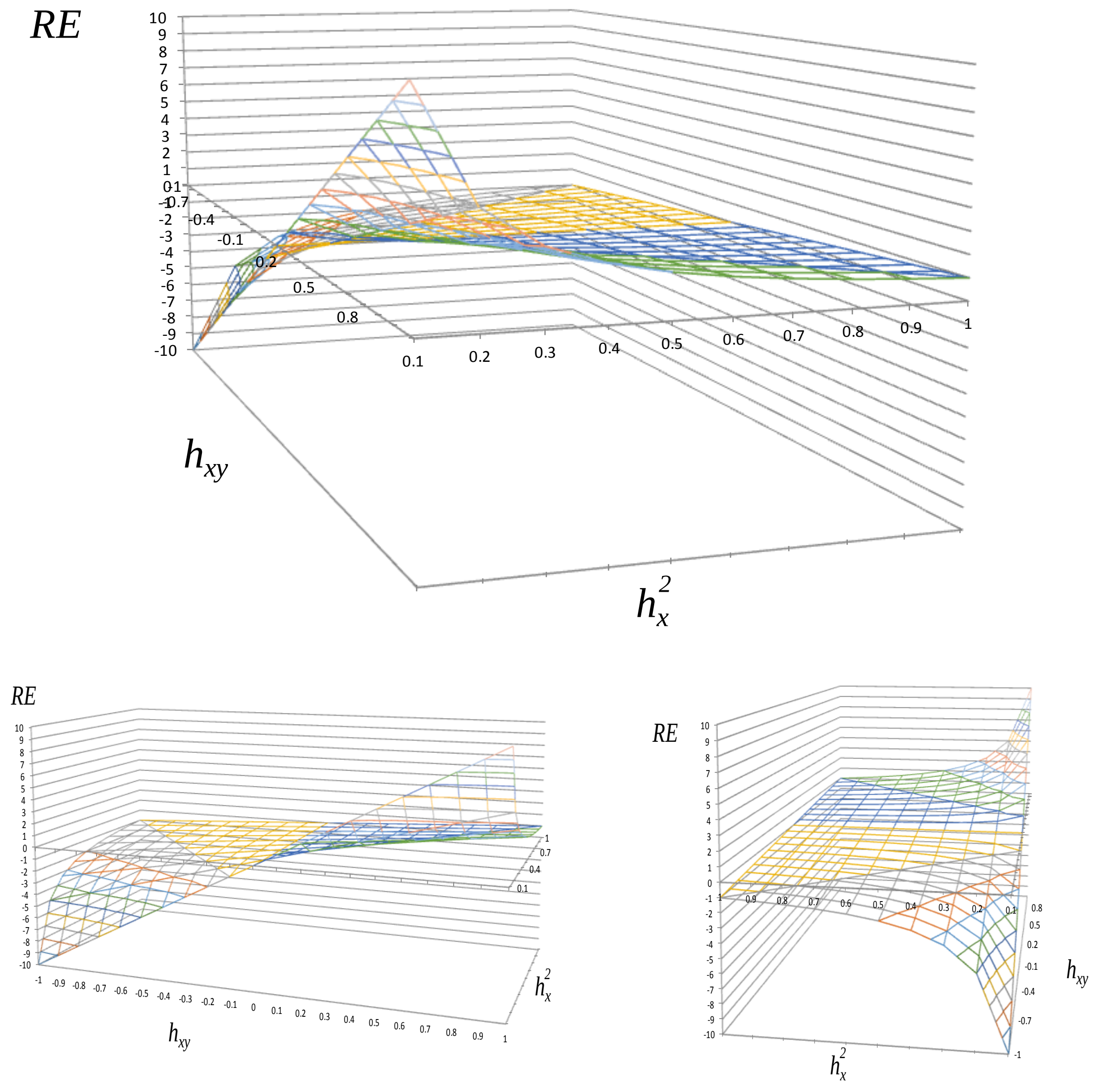

**Figure 8.4** Different views of the relationship between relative efficiency of selection as a function of the coheritability of the traits and the heritability of the directly selected trait.

#### 8.3 A measure of the intensity of GxE Interaction

##### 8.3.1 Type B coheritability and the intensity of GxE interaction S

It is well recognized by breeders that if the rank order and relative magnitude of phenotypic expression are the same across locations, it does not matter in which environment the parental selection is conducted. However, if the rank changes, it might be best to select in the environment in which the organisms will ultimately be reared. Falconer (1952) proposed that the magnitude of genotype-by-environment interaction (GxE) be quantified by the cross-environment genetic correlations, in which the same characters measured in two environments is considered to be two different characters. Type B genetic correlation (using Burdon's 1977 naming) is defined as

$$r_{B_{x\ 1,2}} = \frac{s_{x1,x2}}{\sqrt{s_{x1}^2\ s_{x2}^2}}$$

where $s_{x1,x2}$ refers to the genotypic covariance between the same trait $x$ in environments 1 and 2. The terms $s_{x1}^2$ and $s_{x2}^2$ denotes the estimated genetic variance for the trait $x$ in environments 1 and 2, respectively.

Nevertheless, the heritabilities displayed by the traits in the different environments are also important and informative, and they must be taken into account. Therefore, to have a better idea of the intensity of GxE, would be rather preferable to use the Type B coheritability of the trait, which is defined was defined in Section 3.2.2 as:

$$h_{B_{x\ 1,2}} = \sqrt{h_{x1}^2\ h_{x2}^2}\ r_{B_{x\ 1,2}}$$

$h_{Bx\ 1,2}$ is the coheritability between trait $x$ in environment 1 and 2, $h_{x\blacksquare}^2$ is the heritability of trait $x$ in a given environment (1=env1, 2=env2), and $r_{B_{x\ 1,2}}$ is the Type B-genetic correlation. The intensity of GxE is inversely related to the coheritability, $[GxE] \propto \frac{1}{h_{x1,x2}}$. When the intensity of GxE is high it means that the environment exert a larger influence on the phenotype manifested in their change of rank of the mean performance of the trait, and thus the coheritability is low. On the other hand, a high coheritability across sites shows that the genetic component is stable across environments. Cognizant that one cannot infer the underlying genetic architecture from an observed coheritability, because many different underlying causal pathways can generate the same pattern. Such patterns, however, might suggest causal hypotheses, yet caution must be exercised to make statements about causes of observed coheritabilities (Pigliucci 2005).

Here I present a measure of the intensity of the genotype-by-environment interaction, called S, based on the Type B coheritability of a trait exhibited by members of a genotype (i.e. family, clones, variety) in two distinct environments.

The measure S is defined as

$$S = \frac{1 - |h_{B_{x\,1,2}}|}{1 + |h_{B_{x\,1,2}}|} \qquad 0 < S < 1$$

Which can be plotted as a function of the absolute value of the Type coheritability $h_{B_{x\,1,2}}$ .

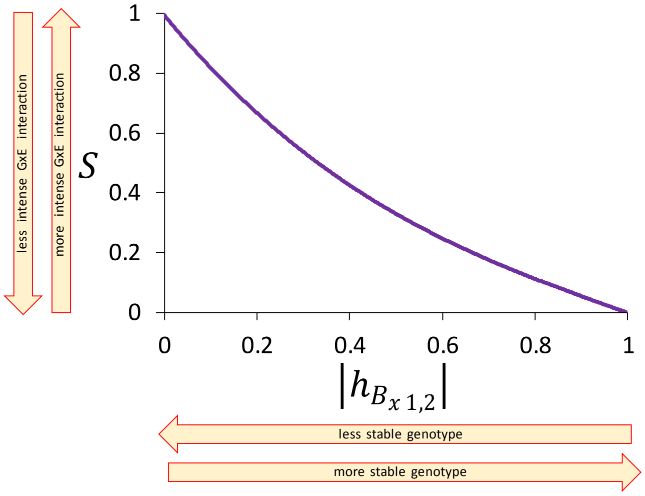

The graph illustrates the inverse relationship between intensity of genotype x environment interaction (S) and the coheritability of the same trait in two environments.

The low absolute value of the coheritability means that the breeding values of a trait displayed by a genotype in an environment are not linearly related to the breeding values of the same genotype in another environment, thus S would be high. High absolute value of the coheritability indicates that the genotype is stable across environments, and the intensity of S is reduced.

##### 8.3.2 Relationship of coheritabilities across environments. Case for two traits.

How the coheritabilities across environments relate to the coheritabilities between traits within an environment?

In the case that two traits are of interest in both environments, the Type B coheritability across sites of each single trait and the coheritability between both traits within each environment can be related, as follows,

Let $h_{B_{x\,1,2}} = \sqrt{h_{x1}^2\, h_{x2}^2}\, r_{B_{x\,1,2}}$ and $h_{B_{y\,1,2}} = \sqrt{h_{y1}^2\, h_{y2}^2}\, r_{B_{y\,1,2}}$ be the Type B coheritability for trait $x$ and $y$, respectively. Then its product is

$$h_{B_{x\,1,2}} \cdot h_{B_{y\,1,2}} = \sqrt{h_{x1}^2\, h_{x2}^2}\, r_{B_{x\,1,2}} \sqrt{h_{y1}^2\, h_{y2}^2}\, r_{B_{y\,1,2}} \qquad [\,8.3.2 - 1]$$

After some rearrangement

$$h_{B_{x\,1,2}} \cdot h_{B_{y\,1,2}} = \sqrt{h_{x1}^2\, h_{y1}^2} \sqrt{h_{x2}^2\, h_{y2}^2}\, r_{B_{x\,1,2}}\, r_{B_{y\,1,2}} \qquad [\,8.3.2 - 2]$$

the geometric means of the heritabilities $\sqrt{h_x^2\, h_y^2}$ can be replaced by $\frac{h_{x,y}}{r_{A_{x,y}}}$ employed now in equation 8.3.2-2], which yields,

$$h_{B_{x\,1,2}} \cdot h_{B_{y\,1,2}} = h_{x1,y1} h_{x2,y2} \frac{r_{B_{x\,1,2}}}{r_{A_{x1,y1}}} \frac{r_{B_{y\,1,2}}}{r_{A_{x2,y2}}} \qquad [\,8.3.2 - 2]$$

Therefore, the product of Type B coheritabilities of two traits is the product of the coheritabilities between the traits in each environment multiplied by the ratio of Type B correlation and the genetic correlation, for both tratits.

### 9 Comparing correlations and coheritabilities

#### 9.1 Types of comparisons

Testing single statistics (correlations, coheritability) was the purview of Section 5. This section deals with comparing two values of correlations or coheritabilities. A proper comparison of correlations between two groups must also involve comparing the groups based on other measures such as the response variable. For details of a method aiming to do this, see Tabesh et al. 2010.

Analogous to the statistical treatment concerning the comparison of correlations (Didenhofen and Musch 2015, ), we can use the same statistical tools to perform the comparison of coheritabilities. There are two cases that can be distinguished in the comparison. A test that of *independent coheritabilities* compares two values obtained from distinct individuals/samples. A test of *dependent coheritabilities* compares two values obtained from the same individuals/sample. It can be referred as either an *dependent, overlapping* test if it compares two coheritabilities with one trait in common, or *dependent, non-overlapping* test if it compares two coheritabilities involving different traits, none in common. Although different interpretation, these two cases of dependent testing are identical from a statistical perspective. An example may explain this better. If, in a forestry progeny trial, the coheritability between height growth and diameter is to be compared to the coheritability between height growth and wood volume, there is a common factor, namely, the variable height growth. This comparison is classified as *dependent* because it compares data obtained from the same group and *overlapping* because both coheritabilities have one factor in common. On the other hand, any of these will be compared to the coheritability between early and late wood density, then the comparison is classified as dependent and non-overlapping. In the case that the data comes from different groups or experiments, then it is classified as non-overlapping (see Example Box 9.1).

| | | Non-overlapping | Non-overlapping |
|---|---|---|---|
| Factor | Dependent | Two correlations or coheritabilities are calculated from the same sample without commom variables involved. | Two correlations or coheritabilities are calculated from the same sample with commom variables involved. |
| | Independent | Two correlations obtained from independent samples without common variable involved. | |

Example Box 9.1

**Dependent, overlapping**

| | | Trait 2 | | | | |
|---|---|---|---|---|---|---|
| | | **H1** 1 | **H3** 2 | **H6** 3 | **H9** 4 | **H12** 5 |
| Trait 1 **H1** | 1 | **1** | $h_{1,2}$ | $h_{1,3}$ | $h_{1,4}$ | $h_{1,5}$ |
| **H3** | 2 | | **1** | $h_{2,3}$ | $h_{2,4}$ | $h_{2,5}$ |
| **H6** | 3 | | | **1** | $h_{3,4}$ | $h_{3,5}$ |
| **H9** | 4 | | | | **1** | $h_{4,5}$ |
| **H12** | 5 | | | | | **1** |

**Problem:** To determine whether the genetic control of a phenotype in a juvenile age is the same as the same phenotype in a mature age.

**Question 1:** At what age is the juvenile phenotype significantly correlated to the phenotype at the mature age?

**Question 2:** At what age is the juvenile phenotype a better predictor of the phenotype at the mature age?

**Data collection:** Measurements of the trait regularly taken from the same individuals during development.

**Dependent, nonoverlapping**

| | | Trait 2 | | | | |
|---|---|---|---|---|---|---|
| | | **D1** 6 | **D3** 7 | **D6** 8 | **D9** 9 | **D12** 10 |
| Trait 1 **H1** | 1 | $h_{1,6}$ | $h_{1,7}$ | $h_{1,8}$ | $h_{1,9}$ | $h_{1,10}$ |
| **H3** | 2 | | $h_{2,7}$ | $h_{2,8}$ | $h_{2,9}$ | $h_{2,10}$ |
| **H6** | 3 | | | $h_{3,8}$ | $h_{3,9}$ | $h_{3,10}$ |
| **H9** | 4 | | | | $h_{4,9}$ | $h_{4,10}$ |
| **H12** | 5 | | | | | $h_{5,10}$ |

**Problem:** We look at whether the degree of genetic control of two traits remains the same or not at juvenile and mature stages.

**Question 1:** Is the correlation between two traits at juvenile and the mature age the same throughout development?

**Question 2:** Is the coheritability between two traits at a given age the same as in any other ages?

**Data collection:** Measurements of both traits regularly taken from the same individuals during development.

**Non-dependent, nonoverlapping**

| | | Generation | | |
|---|---|---|---|---|
| | | parental | progeny | |
| | | **D** 2 | **H** 3 | **D** 4 |
| Generation parental **H** | 1 | $h_{1,2}$ | $h_{1,3}$ | $h_{1,4}$ |
| parental **D** | 2 | **1** | $h_{2,3}$ | $h_{2,4}$ |
| progeny **H** | 3 | | **1** | $h_{3,4}$ |

**Problem:** We look at whether selection influences the degree of genetic control of two traits in two generations.

**Question 1:** What is the influence of selection on the genetic correlations between two traits?

**Question 2:** How selection influences the coinheritance of two traits?

**Data collection:** Measurements of both traits regularly taken from the same individuals at the same age in each generation.

To perform the statistical comparison of the magnitude of two coheritabilities, as well as the confidence interval for such comparison, the free software package *cocor* (Didenhofen and Munsch 2015, available online at http://comparingcorrelations.org) covers a broad range of tests including the comparisons of independent and dependent variables in overlapping or nonoverlapping cases. A thorough elaboration on statistical comparison of correlations (which do also apply to coheritabilities) is provided by the useful appendix accompanying the article (https://journals.plos.org/plosone/article?id=10.1371/journal.pone.0121945#sec010).

#### 9.2 The nature of the coheritability in terms of traits, environment and state

Another aspect pertinent to comparisons is the conceptual nature of the variables (i.e. correlations, coheritabilities) involved in a single correlation of coheritability measure. Table 9 defines the traits, expands the concept of environment, and introduces the concept of state

**Table 9** Conceptual considerations in bivariate measures of relationship such as correlations and coheritability

| | Trait | Environment | State |
|---|---|---|---|
| **Definition** | measure of a phenotypic character, or an aggregate of characters,observed in members of a sample or population. | it refers to any external or extrinsic factor that modulates, modifies, or influence the genetic control of the traits. | it refers to any intrinsic biological condition, quality, or event that influence the genetic control of the traits. |
| **Example** | continuous, discrete, complex | **spatial**: physical environmental, site.<br>**subject**: individual, cohort, family<br>**generational**: parent, progeny<br>**temporal**: circadian, year<br>**treatment**: before, during, after test | **developmental stage**: juvenile, adult.<br>**disease state**: unaffected, affected.<br>**sex:** male, female<br>**ethnic provenance, strain, biovar, clone** |

| | | | |
|---|---|---|---|
| different traits | same environment | same state | **case I** |
| | | different state | **case II** |
| | different environment | same state | **case III** |
| | | different state | **case IV** |
| same trait | same environment | same state | **case V** |
| | | different state | **case VI** |
| | different environment | same state | **case VII** |
| | | different state | **case VIII** |

**Case I** coheritability between two distinct traits, say insect resistance and the abundance of certain metabolite in the leaf (traits) in a localities with varying insect pest abundance (environment) and obtained in individual trees of certain age (state).

**Case II** coheritability between the concentration of two pollutants (traits) in a contaminated site (environment), evaluated in pregnant mothers and the fetuses (state)

**Case III** coheritability between two types of cholesterol evaluated before and after a diet change (environment) in the same individuals in a juvenile and a mature age (state).

**Case IV** coheritability between the abundance of two hormones (traits) measured in individuals when awake and sleeping (environment), in males and females (state).

**Case V** coheritability between wood volume growth (trait) measured in trees planted in the same site (environment) measured at distinct ages (state).

**Case VI** coheritability between concentrations of a pollutant in the blood (trait) measured in specific homes (environments) and among individuals before and after some treatment (state).

**Case VII** coheritability between height growth (trait) measured in individuals growing in different sites (environment) and from different clones (state).

**Case VIII** coheritability between pest resistance (trait) in different plantation sites (environment) and among resistant and susceptible genotypes.

Examples for some cases are show below.

##### Case VIII

same trait, same environment, same state

| | $H_{age2}$ | $H_{age3}$ | $H_{age4}$ |
|---|---|---|---|
| $H_{age1}$ | $h_{H_1H_2}$ | $h_{H_1H_3}$ | $h_{H_1H_4}$ |
| $H_{age2}$ | | $h_{H_2H_3}$ | $h_{H_2H_4}$ |
| $H_{age3}$ | | | $h_{H_3H_4}$ |

The H represents the trait, the numeral indicate the age in which the measurement was taken. The coheritability is represented by $h$ and the symbols of the traits as subscripts.

Measurement in the same individuals

$h_{H_1H_2}$ $h_{H_3H_4}$ dependent nonoverlapping

$h_{H_1H_4}$ $h_{H_3H_4}$ dependent overlapping

##### Case VII

same trait, different environment, same state

| | $H_{site2}$ | $H_{site3}$ | $H_{site4}$ |
|---|---|---|---|
| $H_{site1}$ | $h_{Hsite_1H_{site2}}$ | $h_{H_{site1}H_{site3}}$ | $h_{H_{site1}H_{site4}}$ |
| $H_{site2}$ | | $h_{H_{site2}H_{site3}}$ | $h_{H_{site2}H_{site4}}$ |
| $H_{site3}$ | | | $h_{H_{site3}H_{site4}}$ |

The H represents the trait, the subscript indicate the locality at which the measurement was taken. The coheritability is represented by $h$ and the symbols of the traits as subscripts.

Type B correlations coheritability

$h_{H_1H_3}$ $h_{H_2H_4}$ dependent nonoverlapping

$h_{H_1H_4}$ $h_{H_1H_3}$ dependent overlapping

if different organs in same individuals, dependent

different individuals yet belonging to the same family

#### Case VI

different traits, same environment, same state

| | D | E | F |
|---|---|---|---|
| H | $h_{HD}$ | $h_{HE}$ | $h_{HF}$ |
| J | | $h_{JE}$ | $h_{JF}$ |
| K | | | $h_{KF}$ |

$h_{HD}$ $h_{KF}$ dependent nonoverlapping

$h_{HF}$ $h_{KF}$ dependent overlapping

The H, J, K, D, E, F represent the trai. The coheritability is represented by $h$ and the symbols of the traits as subscripts.

#### Case II

different traits, same environment, different state

| | $D_j$ | $D_m$ |
|---|---|---|
| $H_j$ | $h_{(HD)_j}$ | |
| $H_m$ | | $h_{(HD)_m}$ |

$h_{(HD)_j}$ $h_{(HD)_m}$ independent

different individuals

The H and D represent the traits, the subscript j refers to a juvenile state, m to the mature state. Also applicable is that j represents the parental generation, and m the progeny generation. The coheritability is represented by $h$ and the symbols of the traits as subscripts.

#### Case III

different traits, different environment, same state

site 2

site 1

| | $D_{site1}$ | $D_{site2}$ |
|---|---|---|
| $H_{site1}$ | $h_{(HD)_{site1}}$ | |
| $H_{site2}$ | | $h_{(HD)_{site2}}$ |

$h_{(HD)_1}$ $h_{(HD)_2}$ (in)dependent

The H and D represent the traits, the subscript refers to a environments where the measurements were taken. Also applicable is that j represents the parental generation, and m the progeny generation. The coheritability is represented by $h$ and the symbols of the traits as subscripts.

### 10 Simulation of coheritability and coenvironmentability data

#### 10.1 Derivation of joint probability density function of simulated $\boldsymbol{h}_{xy}$ and $\boldsymbol{e}_{xy}$

Let $(X_1, X_2)$ be a bivariate random vector in which the $X_1$ and $X_2$ variables independently have a Uniform $\left(-\frac{1}{2}, \frac{1}{2}\right)$ distribution. The joint distribution of $X_1$ and $X_2$ is

$f_{X_1,X_2}(x_1, x_2) = f_{X_1}(x_1) \times f_{X_2}(x_2) = 1 \times 1 = 1$, for $-1 < x_1 < 1$ and $-1 < x_2 < 1$.

Now consider a new bivariate random $(H, E)$ vector be defined by

$H = g_1(X_1, X_2) = X_1 + X_2, \quad -1 \le H \le 1,$

$E = g_2(X_1, X_2) = X_1 - X_2, \quad -1 \le E \le 1$

Our goal is to find the joint distribution of $H$ and $E$ expressed in terms of $f_{X_1,X_2}(x_1, x_2)$. The set

$\mathcal{A} = \{(x_1, x_2) : f_{X_1,X_2}(x_1, x_2) > 0\}$, and

$\mathcal{B} = \{(h, e) : h = g_1(x_1, x_2) = x_1 + x_2;\ e = g_2(x_1, x_2) = x_1 - x_{2,},$ for $(x_1, x_2) \in \mathcal{A}$ $\}$

Thus, the joint pdf $f_{H,E}(h, e)$ will be positive on the set $\mathcal{B}$.

Verification of the 1-to-1 transformation

Here we demonstrate that $h = g_1(x_1, x_2)$ and $e = g_2(x_1, x_2)$ define a one-to-one transformation of $\mathcal{A}$ onto $\mathcal{B}$. The transformation is onto because of the definition of $\mathcal{B}$. Thus, for each $(h, e) \in \mathcal{B}$ there is only one $(x_1, x_2) \in \mathcal{A}$ such that $(h, e) = (g_1(x_1, x_2), g_2(x_1, x_2))$. Therefore, we aim to express $x_1$ and $x_2$ in terms of $h$ and $e$. Thus, we obtain the following inverse transformation:

$$x_1 = g_1^{-1}(h, e) = \frac{h + e}{2}$$

$$x_2 = g_2^{-1}(h, e) = \frac{h - e}{2}$$

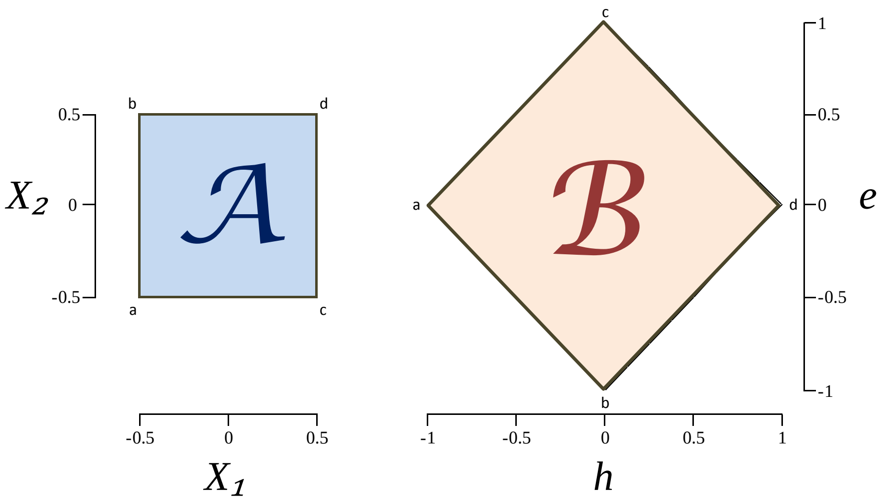

| | $X_1$ | $X_2$ |
|---|---|---|
| a | -0.5 | -0.5 |
| b | -0.5 | 0.5 |
| c | 0.5 | -0.5 |
| d | 0.5 | 0.5 |

| | $h$ <br> $X_1 + X_2$ | $e$ <br> $X_1 + X_2$ |
|---|---|---|
| a | -1 | 0 |
| b | 0 | -1 |
| c | 0 | 1 |
| d | 1 | 0 |

The tables below each graph show that the transformation is 1-to-1.

(5) Jacobian determinant of the transformation

$$J = \begin{vmatrix} \dfrac{\partial x_1}{\partial h} & \dfrac{\partial x_1}{\partial e} \\ \dfrac{\partial x_2}{\partial h} & \dfrac{\partial x_2}{\partial e} \end{vmatrix} = \begin{vmatrix} \dfrac{\partial}{\partial h}\left(\dfrac{h+e}{2}\right) & \dfrac{\partial}{\partial e}\left(\dfrac{h+e}{2}\right) \\ \dfrac{\partial}{\partial h}\left(\dfrac{h-e}{2}\right) & \dfrac{\partial}{\partial e}\left(\dfrac{h-e}{2}\right) \end{vmatrix} = = \begin{vmatrix} \dfrac{1}{2} & \dfrac{1}{2} \\ \dfrac{1}{2} & -\dfrac{1}{2} \end{vmatrix} = -0.5$$

We observe that $J$ is not identical to zero in $\mathcal{B}$. Then the joint pdf of $(H, E)$ is zero outside the set $\mathcal{B}$, and on the set $\mathcal{B}$ is given by

$$f_{H,E}(h, e) = f_{X_1,X_2}(g_1^{-1}(h, e),\ g_2^{-1}(h, e))\ |\ J\ |$$

$$= 1|-0.5\,|$$

$$f_{H,E}(h, e) = 0.5$$

Thus, $P\ [(H,E) \in \mathcal{B}] = P[(X_1, X_2) \in \mathcal{A}]$, and the probability distribution of $(H, E)$ is completely determined by the probability distribution of $(X_1, X_2)$.

Joint probability density function of the variables H and E.

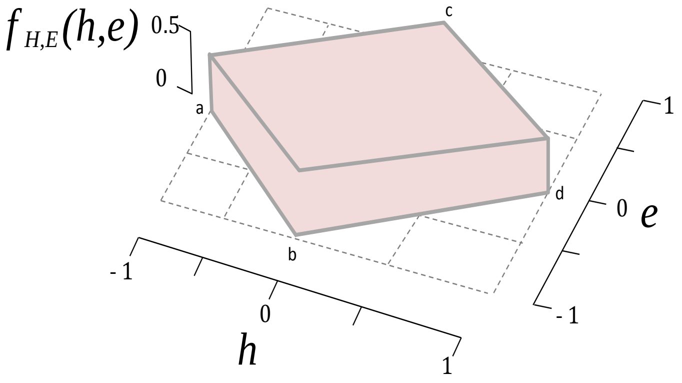

Results of generating simulated data using the model explained in this section.

The data $(n = 3000)$ obtained using a SAS program. Notice that the data allocates randomly throughout the 2DHER-field, all partitions have approximately similar content of data points.

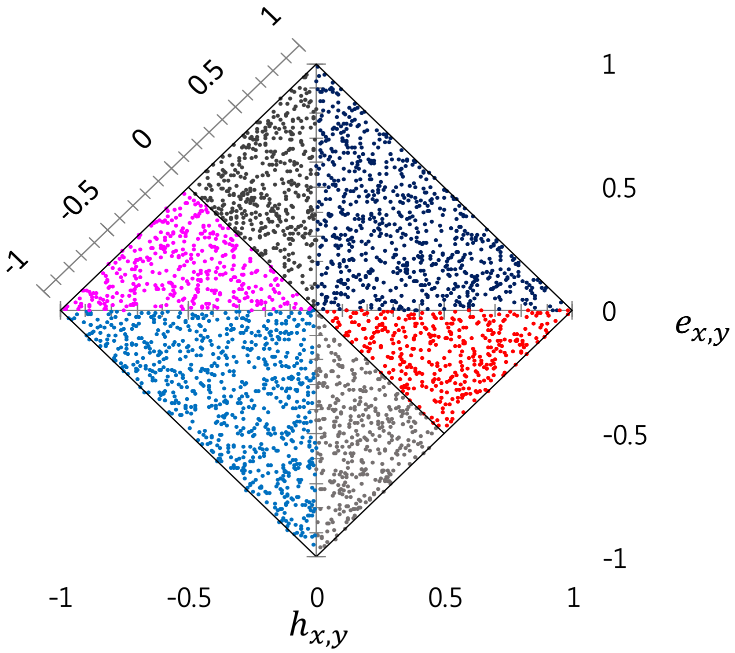

### 11 The Breeder's Equation

This section attempts to illustrate that role of the coheritability in the context of the multivariate form of the breeder's equation, particularly the $H = GP^{-1}$ component, includes terms of coheritability and correlations related in a complex manner, and would remain hidden were not derived explicitly. The equations for a 2x2 and a 3x3 matrices are presented here, this just to show the complex relationship when the covariance terms are not zero.

The breeder's equation measures within-generation response to selection. Predictions from the breeder's equation typically do not hold well in natural populations (Morrisey et al. 2010). Rollinson and Rowe (2015) discuss about the persistent directional selection on body size that is widely observed in wild populations, yet response to selection is generally not in the direction predicted by the breeder's equation. Stasis tends to dominate the temporal dynamic of traits in nature (Uyeda et al. 2010).

This lack of correspondence between expectation and observation may be due, in part, to an incomplete picture of all traits under selection (Pujol et al. 2018), or how the traits interact and correlate properly (Lande and Arnold 1983). Kruuk et al. (2008) acknowledge that the breeder's equation is too simplistic a representation to apply to studies of natural selection in heterogeneous environments. An alternative form of a breeder's equation is presented by Houchmandzadeh (2014).

The breeder's equation, in multivariate form, is given as: $\Delta z = GP^{-1}s$, where $\Delta z$ is a vector of the change in trait means between generations, $G$ is the genetic variance-covariance matrix, $P^{-1}$ is the inverse of the phenotypic variance-covariance matrix, and $s$ is the vector of selection differentials (or alternatively, $\Delta z = G\beta$ where $\beta$ is a vector of selection gradients (Lande and Arnold 1983). Matrix formulations involving phenotypic and genetic correlations are important and must possess certain characteristics. Methods to generate realistic correlation matrices have been proposed for simulation studies (Hardin et al. 2013, Numpacharoen and Atsawarunruagkit 2012).

#### 11.1 The 2x2 Breeder's equation matrix

Let $\boldsymbol{G}_{2x2}$ be the 2x2 (unstructured) genetic variance-covariance matrix

$$\boldsymbol{G}_{2x2} = \begin{bmatrix} V_{A_1} & C_{A_{1.2}} \\ C_{A_{1.2}} & V_{A_2} \end{bmatrix}$$

And $\boldsymbol{P}_{2x2}$ the 2x2 (unstructured) phenotypic variance-covariance matrix $\boldsymbol{P}_{2x2}$

$$\boldsymbol{P}_{2x2} = \begin{bmatrix} V_{P_1} & C_{P_{1.2}} \\ C_{P_{1.2}} & V_{P_2} \end{bmatrix}$$

whose determinant is $\boldsymbol{det}\ \boldsymbol{P}_{2x2} = V_{P_1} V_{P_2} - C_{P_{1,2}}^2$ or

$$\boldsymbol{det}\ \boldsymbol{P}_{2x2} = V_{P_1} V_{P_2} (1 - r_{P_{1,2}}^2)$$

Inverse of $\boldsymbol{P}$ is

$$\boldsymbol{P}_{2x2}^{-1} = \frac{1}{V_{P_1} V_{P_2} (1 - r_{P_{1,2}}^2)} \begin{bmatrix} V_{P_2} & -C_{P_{1.2}} \\ -C_{P_{1.2}} & V_{P_1} \end{bmatrix} = \frac{1}{(1 - r_{P_{1,2}}^2)} \begin{bmatrix} \frac{1}{V_{P_1}} & -\frac{r_{P_{1,2}}}{\sqrt{V_{P_1} V_{P_2}}} \\ -\frac{r_{P_{1,2}}}{\sqrt{V_{P_1} V_{P_2}}} & \frac{1}{V_{P_2}} \end{bmatrix}$$

Which we can use to derive the the 2x2 heritability-coheritability $\mathbf{H}_{2x2} = \boldsymbol{GP^{-1}}_{2x2}$

$$\mathbf{H}_{2x2} = \boldsymbol{GP^{-1}}_{2x2} = \frac{1}{\left(1 - r_{P_{1,2}}^2\right)} \begin{bmatrix} V_{A_1} & C_{A_{1.2}} \\ C_{A_{1.2}} & V_{A_2} \end{bmatrix} \begin{bmatrix} \frac{1}{V_{P_1}} & -\frac{r_{P_{1,2}}}{\sqrt{V_{P_1} V_{P_2}}} \\ -\frac{r_{P_{1,2}}}{\sqrt{V_{P_1} V_{P_2}}} & \frac{1}{V_{P_2}} \end{bmatrix}$$

$$= \frac{1}{(1 - r_{P_{1,2}}^2)} \begin{bmatrix} \frac{V_{A_1}}{V_{P_1}} - \frac{C_{A_{1,2}}}{\sqrt{V_{P_1} V_{P_2}}} r_{P_{1,2}} & \frac{C_{A_{1,2}}}{V_{P_2}} - \frac{V_{A_1}}{\sqrt{V_{P_1} V_{P_2}}} r_{P_{1,2}} \\ \frac{C_{A_{1,2}}}{V_{P_1}} - \frac{V_{A_2}}{\sqrt{V_{P_1} V_{P_2}}} r_{P_{1,2}} & \frac{V_{A_2}}{V_{P_2}} - \frac{C_{A_{1,2}}}{\sqrt{V_{P_1} V_{P_2}}} r_{P_{1,2}} \end{bmatrix}$$

Finally, the 2x2 heritability-coheritability matrix is

$$\mathbf{H}_{2x2} = \boldsymbol{GP^{-1}}_{2x2} = \frac{1}{\left(1 - r_{P_{1,2}}^2\right)} \begin{bmatrix} h_1^2 - h_{1,2}\, r_{P_{1,2}} & h_{1,2} \left( \sqrt{\frac{V_{P_1}}{V_{P_2}}} - \frac{r_{P_{1,2}}}{r_{A_{1,2}}} \sqrt{\frac{V_{A_1}}{V_{A_2}}} \right) \\ h_{1,2} \left( \sqrt{\frac{V_{P_2}}{V_{P_1}}} - \frac{r_{P_{1,2}}}{r_{A_{1,2}}} \sqrt{\frac{V_{A_2}}{V_{A_1}}} \right) & h_2^2 - h_{1,2}\, r_{P_{1,2}} \end{bmatrix}$$

Note that the diagonal elements consist of functions of heritability, coheritability, and phenotypic correlation. The off-diagonal elements involve the coheritabilities affected by raios of variances and covariances.

#### 11.2 The 3x3 Breeder's equation matrix

Let $\boldsymbol{G_{3x3}}$ be the 3x3 (unstructured) genetic variance-covariance matrix

$$\boldsymbol{G_{3x3}} = \begin{bmatrix} V_{A_1} & C_{A_{1.2}} & C_{A_{1.3}} \\ C_{A_{1.2}} & V_{A_2} & C_{A_{2.3}} \\ C_{A_{1.2}} & C_{A_{2.3}} & V_{A_3} \end{bmatrix}$$

And $\boldsymbol{P_{3x3}}$ the 3x3 (unstructured) phenotypic variance-covariance matrix

$$\boldsymbol{P_{3x3}} = \begin{bmatrix} V_{P_1} & C_{P_{1.2}} & C_{P_{1.3}} \\ C_{P_{1.2}} & V_{P_2} & C_{P_{2.3}} \\ C_{P_{1.2}} & C_{P_{2.3}} & V_{P_3} \end{bmatrix}$$

whose determinant is

$$\det \boldsymbol{P_{3x3}} = V_{P_1}V_{P_2}V_{P_3} + 2\, C_{P_{1.2}}C_{P_{1.3}}C_{P_{2.3}} - V_{P_1}C_{P_{2.3}} - V_{P_2}C_{P_{1.3}} - V_{P_3}C_{P_{1.2}}$$

Multiplying and dividing by $V_{P_1}V_{P_2}V_{P_3}$, it yields

$$\det \boldsymbol{P_{3x3}} = V_{P_1}V_{P_2}V_{P_3}(1 + 2r_{P_{1,2}}r_{P_{1.3}}r_{P_{2,3}} - r^2_{P_{1,2}}r^2_{P_{1,3}}r^2_{P_{2,3}})$$

The inverse of the phenotypic variance-covariance matrix is

$$\boldsymbol{P^{-1}_{3x3}} = \frac{1}{(1 + 2r_{P_{1,2}}r_{P_{1.3}}r_{P_{2,3}} - r^2_{P_{1,2}}r^2_{P_{1,3}}r^2_{P_{2,3}})}\begin{bmatrix} \frac{1}{V_{P_1}}(1 - r^2_{P_{2,3}}) & \frac{1}{\sqrt{V_{P_1}V_{P_2}}}(r_{P_{1.3}}r_{P_{2,3}} - r_{P_{1,2}}) & \frac{1}{\sqrt{V_{P_1}V_{P_3}}}(r_{P_{1.2}}r_{P_{2,3}} - r_{P_{1,3}}) \\ \frac{1}{\sqrt{V_{P_1}V_{P_2}}}(r_{P_{1.3}}r_{P_{2,3}} - r_{P_{1,2}}) & \frac{1}{V_{P_2}}(1 - r^2_{P_{1,3}}) & \frac{1}{\sqrt{V_{P_2}V_{P_3}}}(r_{P_{1.2}}r_{P_{1,3}} - r_{P_{2,3}}) \\ \frac{1}{\sqrt{V_{P_1}V_{P_3}}}(r_{P_{1.2}}r_{P_{2,3}} - r_{P_{1,3}}) & \frac{1}{\sqrt{V_{P_2}V_{P_3}}}(r_{P_{1.2}}r_{P_{1,3}} - r_{P_{2,3}}) & \frac{1}{V_{P_3}}(1 - r^2_{1,2}) \end{bmatrix}$$

Then $\mathbf{H_{3x3}} = \boldsymbol{GP^{-1}}_{\boldsymbol{3x3}}$ is

$\boldsymbol{GP^{-1}}_{\boldsymbol{3x3}}$

$$= \frac{1}{\det P_{3x3}}\begin{bmatrix} V_{A_1} & C_{A_{1.2}} & C_{A_{1.3}} \\ C_{A_{1.2}} & V_{A_2} & C_{A_{2.3}} \\ C_{A_{1.2}} & C_{A_{2.3}} & V_{A_3} \end{bmatrix}\begin{bmatrix} \frac{1}{V_{P_1}}(1 - r^2_{P_{2,3}}) & \frac{1}{\sqrt{V_{P_1}V_{P_2}}}(r_{P_{1.3}}r_{P_{2,3}} - r_{P_{1,2}}) & \frac{1}{\sqrt{V_{P_1}V_{P_3}}}(r_{P_{1.2}}r_{P_{2,3}} - r_{P_{1,3}}) \\ \frac{1}{\sqrt{V_{P_1}V_{P_2}}}(r_{P_{1.3}}r_{P_{2,3}} - r_{P_{1,2}}) & \frac{1}{V_{P_2}}(1 - r^2_{P_{1,3}}) & \frac{1}{\sqrt{V_{P_2}V_{P_3}}}(r_{P_{1.2}}r_{P_{1,3}} - r_{P_{2,3}}) \\ \frac{1}{\sqrt{V_{P_1}V_{P_3}}}(r_{P_{1.2}}r_{P_{2,3}} - r_{P_{1,3}}) & \frac{1}{\sqrt{V_{P_2}V_{P_3}}}(r_{P_{1.2}}r_{P_{1,3}} - r_{P_{2,3}}) & \frac{1}{V_{P_3}}(1 - r^2_{1,2}) \end{bmatrix}$$

Then $\mathbf{H}_{3x3}$ results in

$$\boldsymbol{GP^{-1}}_{\mathbf{3x3}} = \frac{1}{(1+2r_{P_{1,2}}r_{P_{1.3}}r_{P_{2,3}} - r_{P_{1,2}}^2 r_{P_{1,3}}^2 r_{P_{2,3}}^2)} \begin{bmatrix} h_1^2\left(1-r_{P_{2,3}}^2\right) + h_{1,2}(r_{P_{1,3}}r_{P_{2,3}} - r_{P_{1,2}}) + h_{1,3}(r_{P_{1,2}}r_{P_{2,3}} - r_{P_{1,3}}) & \frac{\sqrt{V_{P_1}}}{\sqrt{V_{P_2}}}\left[h_1^2(r_{P_{1,3}}r_{P_{2,3}} - r_{P_{1,2}}) + h_{1,2}\left(1-r_{P_{1,3}}^2\right) + h_{1,3}(r_{P_{1,2}}r_{P_{1,3}} - r_{P_{2,3}})\right] & \frac{\sqrt{V_{P_1}}}{\sqrt{V_{P_3}}}\left[h_1^2(r_{P_{1,2}}r_{P_{2,3}} - r_{P_{1,3}}) + h_{1,2}(r_{P_{1,2}}r_{P_{1,3}} - r_{P_{2,3}}) + h_{1,3}\left(1-r_{P_{1,2}}^2\right)\right] \\ \frac{\sqrt{V_{P_2}}}{\sqrt{V_{P_1}}}\left[h_{1,2}\left(1-r_{P_{2,3}}^2\right) + h_2^2(r_{P_{1,3}}r_{P_{2,3}} - r_{P_{1,2}}) + h_{2,3}(r_{P_{1,2}}r_{P_{2,3}} - r_{P_{1,3}})\right] & h_{1,2}(r_{P_{1,3}}r_{P_{2,3}} - r_{P_{1,2}}) + h_2^2\left(1-r_{P_{1,3}}^2\right) + h_{2,3}(r_{P_{1,2}}r_{P_{1,3}} - r_{P_{2,3}}) & \frac{\sqrt{V_{P_2}}}{\sqrt{V_{P_3}}}\left[h_{1,2}(r_{P_{1,2}}r_{P_{2,3}} - r_{P_{1,3}}) + h_2^2(r_{P_{1,2}}r_{P_{1,3}} - r_{P_{2,3}}) + h_{2,3}\left(1-r_{P_{1,2}}^2\right)\right] \\ \frac{\sqrt{V_{P_3}}}{\sqrt{V_{P_1}}}\left[h_{1,3}\left(1-r_{P_{2,3}}^2\right) + h_{2,3}(r_{P_{1,3}}r_{P_{2,3}} - r_{P_{1,2}}) + h_3^2(r_{P_{1,2}}r_{P_{2,3}} - r_{P_{1,3}})\right] & \frac{\sqrt{V_3}}{\sqrt{V_{P_2}}}\left[h_{1,3}(r_{P_{1,3}}r_{P_{2,3}} - r_{P_{1,2}}) + h_{1,2}\left(1-r_{P_{1,3}}^2\right) + h_3^2(r_{P_{1,2}}r_{P_{2,3}} - r_{P_{2,3}})\right] & h_{1,3}(r_{P_{1,2}}r_{P_{2,3}} - r_{P_{1,3}}) + h_{2,3}(r_{P_{1,2}}r_{P_{1,3}} - r_{P_{2,3}}) + h_3^2\left(1-r_{P_{1,2}}^2\right) \end{bmatrix}$$

To facilitate visualization, the $\mathbf{H}_{3x3}$ is presented below as a block matrix where the $\boldsymbol{GP^{-1}}_{\mathbf{3x3}}$ has been partition in one row-group and three col-groups (column vectors), namely $\mathbf{H}_{11}$ , $\mathbf{H}_{12}$ , $\mathbf{H}_{13}$.

$$\mathbf{H}_{3x3} = \boldsymbol{GP^{-1}}_{\mathbf{3x3}} = \frac{\mathbf{1}}{(1+2r_{P_{1,2}}r_{P_{1.3}}r_{P_{2,3}} - r_{P_{1,2}}^2 r_{P_{1,3}}^2 r_{P_{2,3}}^2)} [\, \mathbf{H}_{11} \;\; \mathbf{H}_{12} \;\; \mathbf{H}_{13} \,]$$

Where

$$\boldsymbol{H}_{11} = \begin{bmatrix} h_1^2(1-r_{P_{2,3}}^2) + h_{1,2}(r_{P_{1,3}}r_{P_{2,3}} - r_{P_{1,2}}) + h_{1,3}(r_{P_{1,2}}r_{P_{2,3}} - r_{P_{1,3}}) \\ \frac{\sqrt{V_{P_2}}}{\sqrt{V_{P_1}}}[h_{1,2}(1-r_{P_{2,3}}^2) + h_2^2(r_{P_{1,3}}r_{P_{2,3}} - r_{P_{1,2}}) + h_{2,3}(r_{P_{1,2}}r_{P_{2,3}} - r_{P_{1,3}})] \\ \frac{\sqrt{V_3}}{\sqrt{V_{P_1}}}[h_{1,3}(1-r_{P_{2,3}}^2) + h_{2,3}(r_{P_{1,3}}r_{P_{2,3}} - r_{P_{1,2}}) + h_3^2(r_{P_{1,2}}r_{P_{2,3}} - r_{P_{1,3}})] \end{bmatrix}$$

$$\boldsymbol{H}_{12} = \begin{bmatrix} \frac{\sqrt{V_{P_1}}}{\sqrt{V_{P_2}}}[h_1^2(r_{P_{1,3}}r_{P_{2,3}} - r_{P_{1,2}}) + h_{1,2}(1-r_{P_{1,3}}^2) + h_{1,3}(r_{P_{1,2}}r_{P_{1,3}} - r_{P_{2,3}})] \\ h_{1,2}(r_{P_{1,3}}r_{P_{2,3}} - r_{P_{1,2}}) + h_2^2(1-r_{P_{1,3}}^2) + h_{2,3}(r_{P_{1,2}}r_{P_{1,3}} - r_{P_{2,3}}) \\ \frac{\sqrt{V_{P_3}}}{\sqrt{V_{P_2}}}[h_{1,3}(r_{P_{1,3}}r_{P_{2,3}} - r_{P_{1,2}}) + h_{1,2}(1-r_{P_{1,3}}^2) + h_3^2(r_{P_{1,2}}r_{P_{2,3}} - r_{P_{2,3}})] \end{bmatrix}$$

$$\boldsymbol{H_{13}} = \begin{bmatrix} \frac{\sqrt{V_{P_1}}}{\sqrt{V_{P_3}}}\left[h_1^2\left(r_{P_{1,2}}r_{P_{2,3}} - r_{P_{1,3}}\right) + h_{1,2}\left(r_{P_{1,2}}r_{P_{1,3}} - r_{P_{2,3}}\right) + h_{1,3}\left(1 - r_{P_{1,2}}^2\right)\right] \\ \frac{\sqrt{V_{P_2}}}{\sqrt{V_{P_3}}}\left[h_{1,2}\left(r_{P_{1,2}}r_{P_{2,3}} - r_{P_{1,3}}\right) + h_2^2\left(r_{P_{1,2}}r_{P_{1,3}} - r_{P_{2,3}}\right) + h_{2,3}\left(1 - r_{P_{1,2}}^2\right)\right] \\ h_{1,3}\left(r_{P_{1,2}}r_{P_{2,3}} - r_{P_{1,3}}\right) + h_{2,3}\left(r_{P_{1,2}}r_{P_{1,3}} - r_{P_{2,3}}\right) + h_3^2\left(1 - r_{P_{1,2}}^2\right) \end{bmatrix}$$

In the case that the phenotypic matrix $\boldsymbol{P}$ has zero off-diagonal elements (no covariances), it becomes a diagonal matrix whose elements are the phenotypic variances of the traits, and further consider that the phenotypic variances have been standardized to 1, then the heritability-coheritability $\mathbf{H}$ matrices become

$$\mathbf{H}_{\boldsymbol{2x2}} = \boldsymbol{GP^{-1}}_{\boldsymbol{2x2}} = \begin{bmatrix} h_1^2 & h_{1,2} \\ h_{1,2} & h_2^2 \end{bmatrix}$$

and

$$\mathbf{H}_{\boldsymbol{3x3}} = \boldsymbol{GP^{-1}}_{\boldsymbol{3x3}} = \begin{bmatrix} h_1^2 & h_{1,2} & h_{1,3} \\ h_{1,2} & h_2^2 & h_{2,3} \\ h_{1,3} & h_{2,3} & h_3^2 \end{bmatrix}$$

The null phenotypic covariance between the traits implies that the phenotypic correlations between the traits is zero. As we have seen, this fact does not preclude the coheritability to have statistically significant values, and be of importance. In addition, under the condition $r_{P_{x,y}} = 0$, then $h_{x,y} = -e_{x,y}$ .

### 12 Supplementary Text

#### 12.1 Alternative conceptual frameworks to assess quantitative joint inheritance

The notion that a phenotype is by nature a composite of an organism's observable characters that are manifested in morphological, developmental, biochemical, behavioral or physiological properties, leads to think that a phenotype is inherently a multivariate concept. This has prompted researchers to find ways to express a multivariate form of heritability (Carper 2008).

Based on Lande (1979) proposal of a multivariate generalization of the Breeder's equation $\Delta\boldsymbol{z} = \boldsymbol{G}\boldsymbol{P}^{-1}\boldsymbol{s}$ (where $\boldsymbol{G}$ and $\boldsymbol{P}$ are the additive genetic and (co)variance matrices, respectively; and $\boldsymbol{s}$ the selection differential), Klingenberg and Leamy (2001) interpreted the dominant eigenvalue of $\boldsymbol{G}\boldsymbol{P}^{-1}$ as a multivariate heritability estimate. Myers et al. (2006) re-scaled the largest eigenvalue by the sum of the eigenvalues. These eigenvalue-based multivariate heritability measures have drawbacks mainly in the difficulty of estimating the standard errors of eigenvalues. In addition, as the number of traits increase, concomitantly increase the number of variance and covariance to be estimated, leading to an over parametrized model.

While imbalance is fairly easily accommodated in the multivariate setting, the inclusion of multiple traits greatly increases the number of variance and covariance components to be estimated, and consequently of computation time. The lack of reliable standard error estimates has led to the use of nonparametric resampling techniques in connection with multivariate analysis (Myers et al. 2006, Carper 2008).

Other forms have been proposed such as the ratio of the sum of the diagonal elements of $\boldsymbol{G}$ and $\boldsymbol{P}$, $\mathrm{h{=}tr}(\boldsymbol{G})/\mathrm{tr}(\boldsymbol{P})$ proposed by Klingender and Monteiro (2005). They recognize that the method ignores the covariation among traits, and the direction of the selection differential or the direction of variation in $\boldsymbol{G}$ and $\boldsymbol{P}$. Basset and De Jong (2011) used an Evolutionary Algorithm to express multivariate heritability of $\mathrm{M}$ traits as a metric function of $\boldsymbol{G}\boldsymbol{P}^{-1}$ namely, $h = m(\boldsymbol{G}\boldsymbol{P}^{-1}) = \sqrt[M]{\det(\boldsymbol{G}\boldsymbol{P}^{-1})}$. Guo et al. (2016) present two measures of coheritability between a pair traits using GWAs data in the framework of high dimensional linear models.

#### 12.2 Is there a "missing" coheritability?

In the framework of genome-wide association studies, the problem of a "missing" coheritability is expected, and estimates of coheritability of a pair of traits using GWAS data must be taken cautiously, under the view that methodological and statistical artifacts have a higher risk of influencing such estimates.

Gianola et al. (2015) rightly stated accurate predictions of complex traits and estimation of genomic correlations can be obtained from multivariate quantitative genetic analysis based on markers. But such determinations (SNPS heritability, correlations) cannot be confused with heritabilities and genetic correlations because of the discrepancy and distinctiveness of the sources that bring forth the data. Certainly in single-trait analyses, imperfect marker-QTL linkage disequilibrium results in missing heritability. In a multivariate context the problem is compounded by missing correlations. Therefore, caution is recommended in making pronouncements on causality when dealing with SNP heritabilities and genomic correlations involving complex traits. Speculating about genetic correlations in the study of complex traits, using marker data must be considered conjectural.

#### 12.3 Coheritability of threshold traits

This brief explanation of coheritability of threshold traits is based on Lee et al. (2011, 2012).

First, two models are presented that describe the phenotypic trait in its observed scale, and in an unobserved underlying liability scale.

**OBSERVED SCALE**

$$x = u_x + e_x \qquad y = u_y + e_y$$

where

The X and Y are the threshold traits defined as

$$x = \begin{cases} 1 & \text{for individuals having the condition } x \\ \square & \square \\ 0 & \text{for individuals not having the condition } y \end{cases}$$

$$Var(x) = K_x(1 - K_x)$$

$$y = \begin{cases} 1 & \text{for individuals having the condition } x \\ \square & \square \\ 0 & \text{for individuals not having the condition } y \end{cases}$$

$u_{\blacksquare}$ is the random additive genetic effects from aggregate marker data

$e_{\blacksquare}$ is the random residual

$$Var(y) = K_y\left(1 - K_y\right)$$

$K_x$ and $K_y$ represent the prevalence of the condition in the population

**LIABILITY SCALE**

$$l_x = g_x + e_{l\,x} \qquad l_y = g_y + e_{l\,y}$$

where

$l_{\blacksquare}$ is the liability phenotype corresponding to trait $\blacksquare$.
$l_{\blacksquare} \sim N(0\,,1)$ ,

$g_{\blacksquare}$ is the random additive genetic effect on the liability scale, $g_{\blacksquare} \sim N(0\,,\sigma^2_{g_{\blacksquare}})$ ,

$e_{l\,\blacksquare}$ is the random residual on the liability scale, $e_{\blacksquare} \sim N(0\,,\sigma^2_{e_{\blacksquare}})$ ,

A simple linear regression can be used as an approximate model of the relationship between the genetic value on the observed scale and the genetic value in the liability scale, for each trait

$$u_{\blacksquare} = c + b_{\blacksquare} g_{\blacksquare}$$

Where the regression parameter $b$ is defined as the covariance of the dependent variable $u$ divided by the variance of the independent variable $g$

$$b = \frac{Cov(u, g)}{\sigma_g^2} = z$$

Therefore,

$$u_{■} = z_{■} g_{■}$$

The genetic covariance between the additive values of $u_x$, and $u_y$ are related to the covariance to the additive values in the liability scale, as follows:

$$Cov(u_x, u_y) = Cov(z_x g_x, z_y g_y) = z_x z_y \, Cov(g_x, g_y)$$

We define the coheritability as the ratio of the genetic covariance divided by the square root of the product of the phenotypic variance of the traits.

Note that the $Cov(g_x, g_y) = h_{l_{x,y}}$ because the variance of the liability phenotype for each trait is unity.

Thus

$$Cov(u_x, u_y) = z_x z_y \, h_{l_{x,y}}$$

In the observed scale, the coheritability between threshold traits $x$ and $y$,

$$h_{o_{x,y}} = \frac{\sigma_{u_x, u_y}}{\sqrt{K_x(1 - K_x)\, K_y(1 - K_y)}}$$

The relationship between the coheritability in the observed ($h_{o_{x,y}}$) and liability ($h_{l_{x,y}}$) scales is,

$$h_{o_{x,y}} = \frac{z_x \, z_y \, h_{l_{x,y}}}{\sqrt{K_x(1 - K_x)\, K_y(1 - K_y)}}$$

In the liability scale, the coheritability between traits $x$ and $y$, is

$$h_{l_{x,y}} = \frac{h_{o_{x,y}} \sqrt{K_x(1 - K_x)\, K_y(1 - K_y)}}{z_x \, z_y}$$

#### 12.4 Partitions of the additive genetic covariance

The additive genetic covariance between two traits $x$ and $y$ can be partitioned into a pleiotropic component and a linkage component (Gardner and Latta 2007):

$$C_{G_{x,y}} = \underbrace{2\sum_{i=1}^{n} p_i\, q_i\, a_{ix}\, a_{iy}}_{pleiotropic} + \underbrace{2\sum_{j=1}^{l_1}\sum_{k\neq j}^{l_2} a_{jx}\, a_{ky}\, D_{jk}}_{linkage\ \ disequilibrioum}$$

More explicitly, for the case of a two biallelic, pleiotropic loci that are in linkage disequilibrium, the genetic covariance on two traits $(x, y)$ is

$$C_{G_{x,y}} = \underbrace{2\, p_1\, q_1\, a_{1x}\, a_{1y} + 2\, p_2\, q_2\, a_{2x}\, a_{2y}}_{pleiotropic} + \underbrace{2\ a_{1x}\, a_{2y}\, D_{12} + 2\ a_{1y}\, a_{2x}\, D_{12}}_{linkage\ \ disequilibrioum}$$

where $p$ is the allele frequency of the $i^{th}$ locus, $q = 1 - p$, $a_{1x}$ is the additive effect of the locus 1 on trait $x$, $D_{12}$ is the measure of linkage disequilibrium ($D_{jk}$ depends upon the recombination distance between loci).

Details regarding the pleiotropic component

Consider a single biallelic locus whose alleles $A_1$ and $A_2$ exert effects on two traits, namely trait $x$ and trait $y$. Using Falconer and Mackay's notation, we associate an arbitrarily-assigned value to the genotypes formed by random association of the alleles, whose frequencies in the population are $p$ for $A_1$, and $q$ for $A_2$ The objective is to determine a measure of an individual based upon the particular allele combination present in the locus, and transmitted to the progeny. The average effect of an allele on a trait is such value, and depends on the genotypic values $a$ and $d$. Table 12.4 shows the results of the derivation of the average effect of the alleles $A_1$ and $A_2$ on each of the traits $x$ and $y$, expressed as the mean deviation (from the population mean) of individuals which received that allele from one parent, the other allele come from a random parent from the population.

Figures 12.4-1 and 12.4-2 display the average effect of alleles when the

**Table 12.4** Average effect of alleles of a biallelic, pleotropic locus affecting traits $x$ and $y$

Allele in population: $A_1$ ($p$) spanning $A_1A_1$–$A_1A_2$; $A_2$ ($q$) spanning $A_1A_2$–$A_2A_2$

| | | | $A_1A_1$ | $A_1A_2$ | $A_2A_2$ | genotypic value | Mean genotypic value | Population genotypic (and phenotypic) mean to be deducted | Average effect of the allele on a trait |
|---|---|---|---|---|---|---|---|---|---|
| | | | $+a$ | $d$ | $-a$ | | | | |
| Trait $x$ | $A_1$ | $A_1A_1$ | $p$ | | | $+a_x$ | $pa_x + qd_x$ | $-[\,a_x(p-q) + 2d_x pq\,]$ | $q\alpha_x = q[a_x + d_x(q-p)]$ |
| | | $A_1A_2$ | | $q$ | | $d_x$ | | | |
| | $A_2$ | $A_1A_2$ | | $p$ | | $d_x$ | $-qa_x + pd_x$ | $-[\,a_x(p-q) + 2d_x pq\,]$ | $-p\alpha_x = -p[a_x + d_x(q-p)]$ |
| | | $A_2A_2$ | | | $q$ | $-a_x$ | | | |
| Trait $y$ | $A_1$ | $A_1A_1$ | $p$ | | | $+a_y$ | $pa_y + qd_y$ | $-[\,a_y(p-q) + 2d_y pq\,]$ | $q\alpha_y = q[a_y + d_y(q-p)]$ |
| | | $A_1A_2$ | | $q$ | | $d_y$ | | | |
| | $A_2$ | $A_1A_2$ | | $p$ | | $-a_y$ | $-qa_y + pd_y$ | $-[\,a_y(p-q) + 2d_y pq\,]$ | $-p\alpha_y = -p[a_y + d_y(q-p)]$ |
| | | $A_2A_2$ | | | $q$ | $d_y$ | | | |

We can see that in these derivations $\alpha_x = [a_x + d_x(q-p)]$ and $\alpha_y = [a_y + d_y(q-p)]$ are the average effect on a trait due to allele substitution in that locus.

| | | genotype frequency $f_G$ | Breeding value: Trait $x$ $B_x$ | Breeding value: Trait $y$ $B_y$ |
|---|---|---|---|---|
| genotype | $A_1A_1$ | $p^2$ | $2q\alpha_x$ | $2q\alpha_y$ |
| | $A_1A_2$ | $2pq$ | $(q-p)\alpha_x$ | $(q-p)\alpha_y$ |
| | $A_2A_2$ | $q^2$ | $-2q\alpha_x$ | $-2q\alpha_y$ |

The genotypic covariance is obtained by summing up the product of the genotype frecuency $f_G$ times the breeding values of trait $B_x$ and $B_y$ for each genotype, as follows:

$$C_{G_{x,y}} = \sum_{i \in S} \left(f_G B_x B_y\right)_i \qquad S = \{\, A_1A_1 \,,\, A_1A_2 \,,\, A_2A_2 \,\}$$

Genotypic covariance of breeding values of trait $x$ and trait $y$

$$C_{G_{x,y}} = p^2(2q\alpha_x)(2q\alpha_y) + 2qp(q-p)\alpha_x(q-p)\alpha_y + (-2p\alpha_x)(-2p\alpha_y)q^2$$

$$= 4p^2q^2\alpha_x\alpha_y + 2pq(q-p)^2\alpha_x\alpha_y + 4p^2q^2\alpha_x\alpha_y$$

$$= 2pq\alpha_x\alpha_y\,(2pq + (q-p)^2 + 2pq)$$

$$= 2pq\alpha_x\alpha_y(4pq + q^2 - 2pq + p^2)$$

$$= 2pq\alpha_x\alpha_y(p^2 + 2pq + q^2)$$

$$= 2pq\alpha_x\alpha_y$$

Therefore the genotypic covariance between two traits influenced by a single locus is a function of the average effect of gene substitution exerted on each trait and the frequency of the alleles in the population.

Genotypic covariance among traits $x$ and $y$ controlled by a biallelic locus (three equivalent expressions):

$$C_{G_{x,y}} = \begin{cases} 2pq\alpha_x\alpha_y \\ 2pq\,[a_x + d_x(q-p)][a_y + d_y(q-p)] \\ 2pq\,[a_xa_y + \left(a_xd_y + a_yd_x\right)(q-p) + d_xd_y(q-p)^2 \end{cases}$$

Notice that if dealing with the same trait, then $x = y$, it results in the familiar expression of the genotypic variance of the trait

Genotypic variance of a trait controlled by an biallelic locus

$$V_G = \begin{cases} 2pq\alpha^2 \\ 2pq\,[a + d(q-p)]^2 \\ 2pq\,[a^2 + 2ad(q-p) + d^2(q-p)^2] \end{cases}$$

Derivation of the average effect expressions of Table 12.4

Average effect of allele $A_1$ on trait *x*

$\alpha_{1x} = pa_x + qd_x - [\, a_x\,(p-q) + 2d_x\, pq \,]$
$2d_y\, pq \,]$

$= pa_x + a_x(p-q) - qd_x - 2pqd_x$

$= a_x(p-p+q) - qd_x(1-2p)$

$= qa_x + qd_x(q-p)$

$= q[a_x + d_x(q-p)]$

$= q\alpha_x$

Average effect of allele $A_1$ on trait *y*

$\alpha_{1y} = pa_y + qd_y - [\, a_y\,(p-q) +$

$= pa_y + a_y(p-q) - qd_y - 2pqd_y$

$= a_y(p-p+q) - qd_y(1-2p)$

$= qa_y + qd_y(q-p)$

$= q[a_y + d_y(q-p)]$

$= q\alpha_y$

Average effect of allele $A_2$ on trait *x*

$\alpha_{2x} = -qa_x + pd_x - [\, a_x\,(p-q) + 2d_x\, pq \,]$
$2d_y\, pq \,]$

$= -qa_x + pd_x - a_x\,(p-q) + 2d_x\, pq$
$2d_y\, pq$

$= -qa_x(q+p-q) - pd_x(1-2p)$

$= -pa_x - pd_x(p-q)$

$= -p[a_x + d_x(q-p)]$

$= -p\alpha_x$

Average effect of allele $A_2$ on trait *y*

$\alpha_{2y} = -qa_y + pd_y - [\, a_y\,(p-q) +$

$= -qa_y + pd_y - a_y\,(p-q) +$

$= -qa_y(q+p-q) - pd_y(1-2p)$

$= -pa_y - pd_y(p-q)$

$= -p[a_y + d_y(q-p)]$

$= -p\alpha_y$

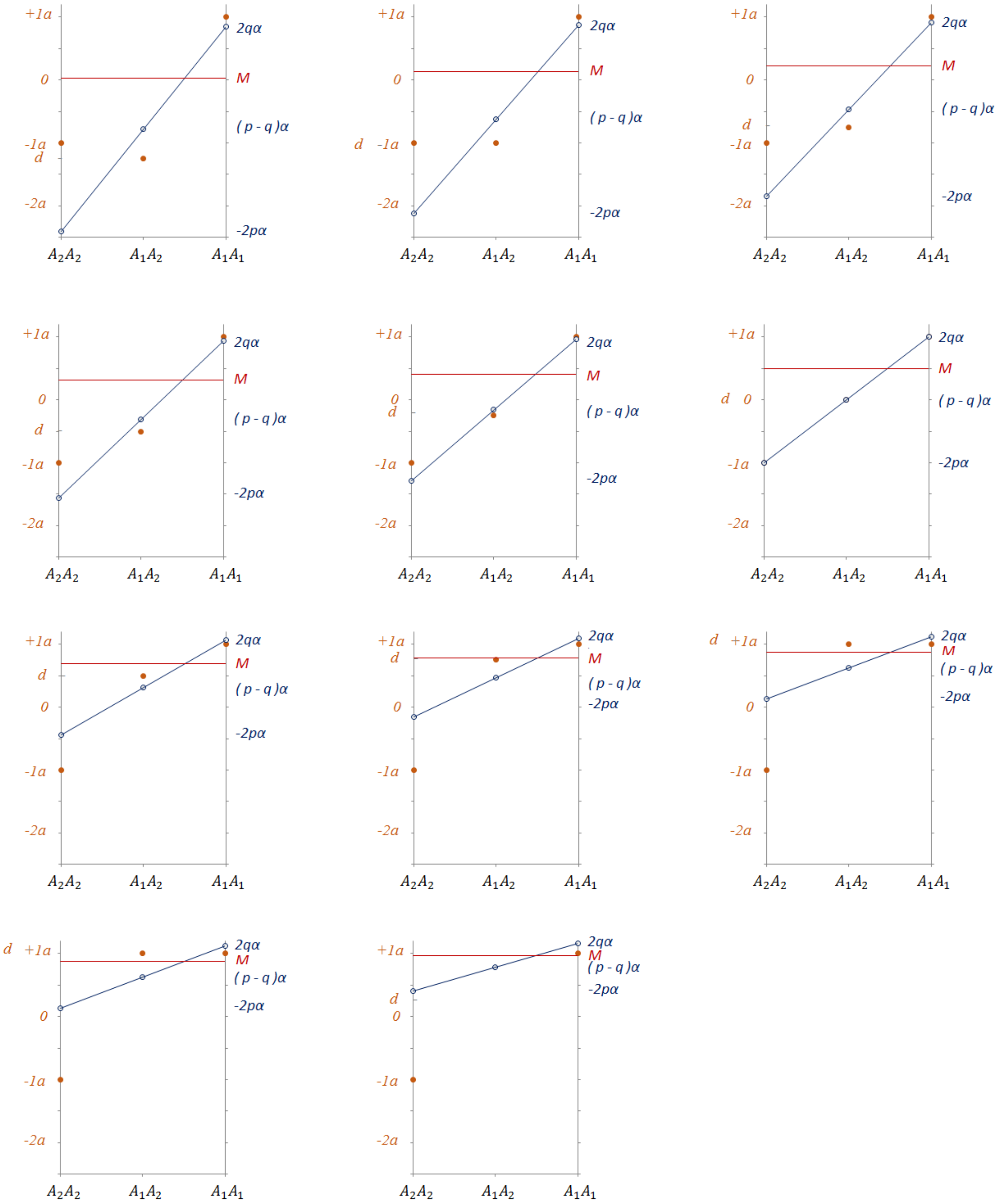

**Figure 12.4-1** Average effect of a allele on a single trait, under varying values of $d$. The allele frequency of $p = 0.25,\ q = 0.75$

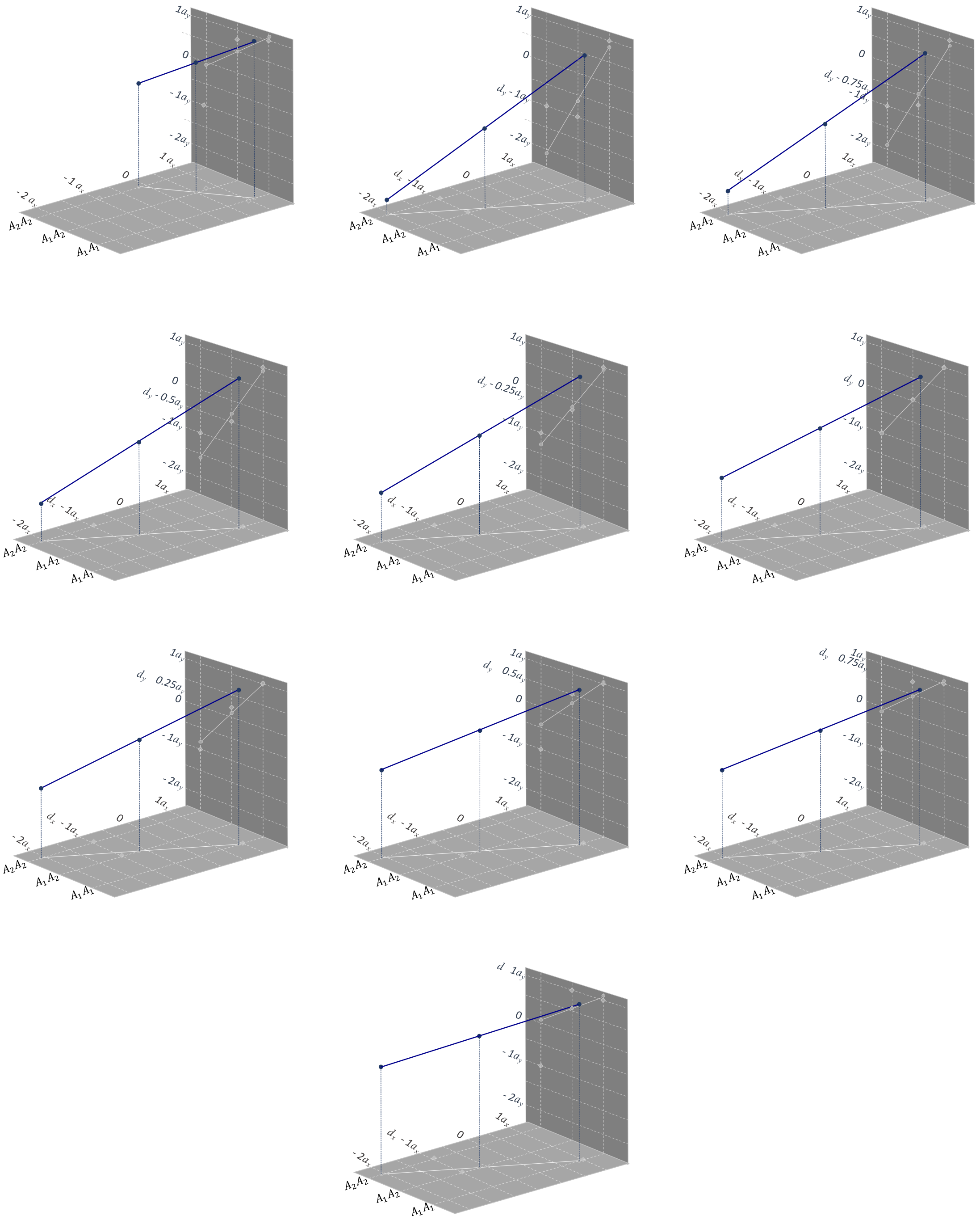

**Figure 12.4-2** Average effect of two alleles of a locus on two traits. The horizontal plane is for trait $x$ and has a fixed value of $d = -1.25a$, the vertical plane is the effect on trait y and has varying values of $d$. This is an extension to the Figure 12.4-1.

#### 12.5 An epidemiological bivariate measure gene-trait relationship

Here I present an extension of the concept positive predictive value, as generally used in epidemiology. The experimental scheme is to screen-test individuals for a particular genetic variant(s) that affects at least two traits. The data is meristic in nature, it uses count data to perform calculations. I called this measure bivariate positive predictive value, and is defined as,

$$bPPV = \frac{\begin{array}{c} Number\ of\ individuals\ with\ genetic\ variant\ \ V \\ exhibiting\ traits\ \ x\ \ and\ \ y \end{array}}{Number\ of\ individuals\ with\ genetic\ variant\ \ V}$$

The denominator includes all individuals with genetic variant $V$ that have both traits, and those having one of the traits, and those not presenting any of the traits. The bivariate positive predictive value of the genetic screening test is the probability of returning a positive result (i.e. the individual has a genetic variant in a locus of interest) correctly identifying those individuals possessing variant $V$ that do show both traits under consideration (see Trevethan 2017).

To illustrate this concept with hypothetical, yet realistic, examples.

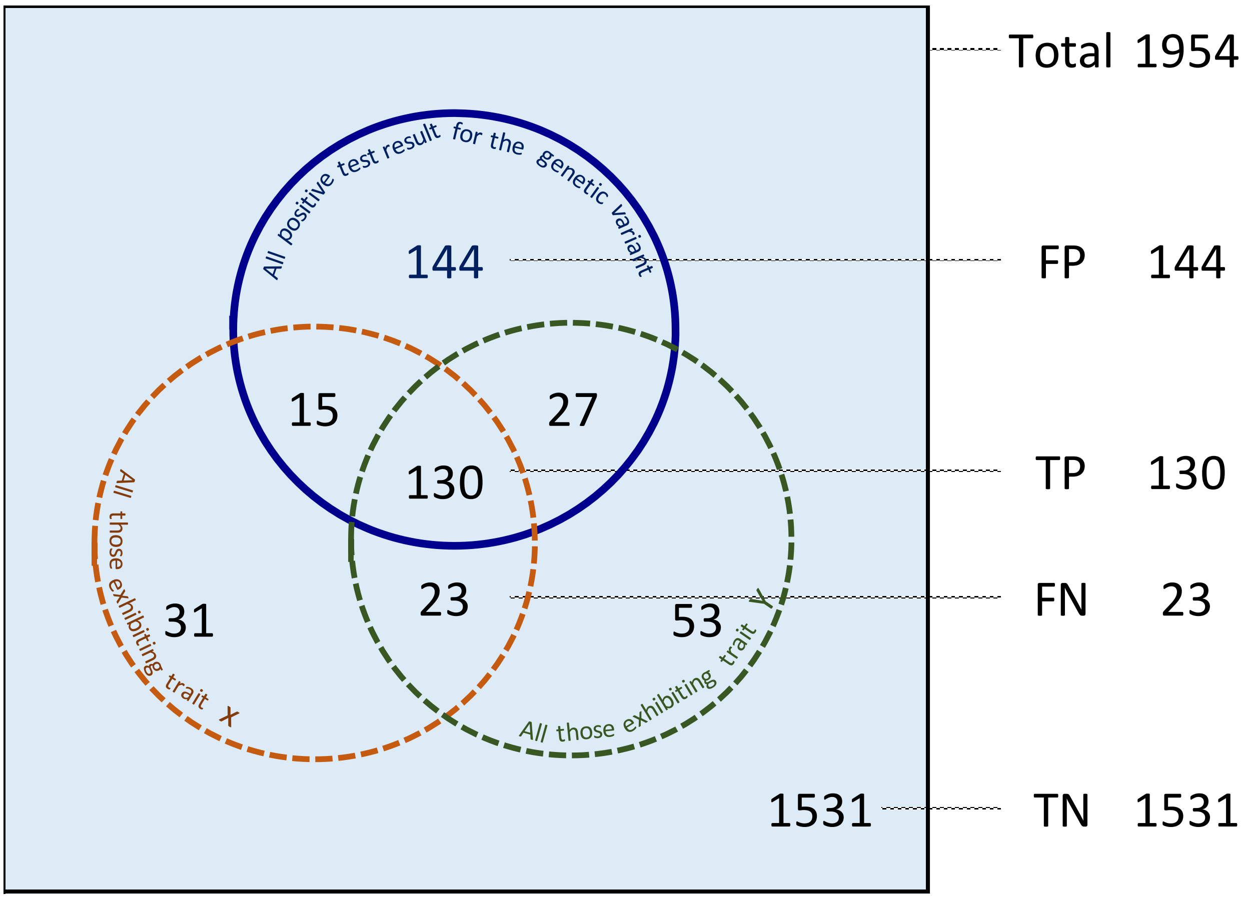

$$proportion\ of\ individuals\ exhibiting\ both\ traits = \frac{130 + 23}{1954} = 8\ \%$$

$$sensitivity = \frac{TP}{all\ those\ exhibiting\ both\ traits} = \frac{130}{130 + 23} = 85\ \%$$

$$specificity = \frac{TN}{all\ those\ not\ exhibiting\ both\ traits} = \frac{1531}{1531 + 31 + 15 + 144 + 27 + 53} = 85\ \%$$

$$\mathbf{bPPV} = \frac{TP}{all\ those\ with\ positive\ test\ result} = \frac{130}{130 + 15 + 27 + 144} = 41\ \%$$

**Figure 12.5-1** Determination of the bivariate positive predictive value (bPPV) of a genetic screening test. ( **FP**, False positive; **TP**, True Positive; **FN**, False Negative, **TN**, True Negative)

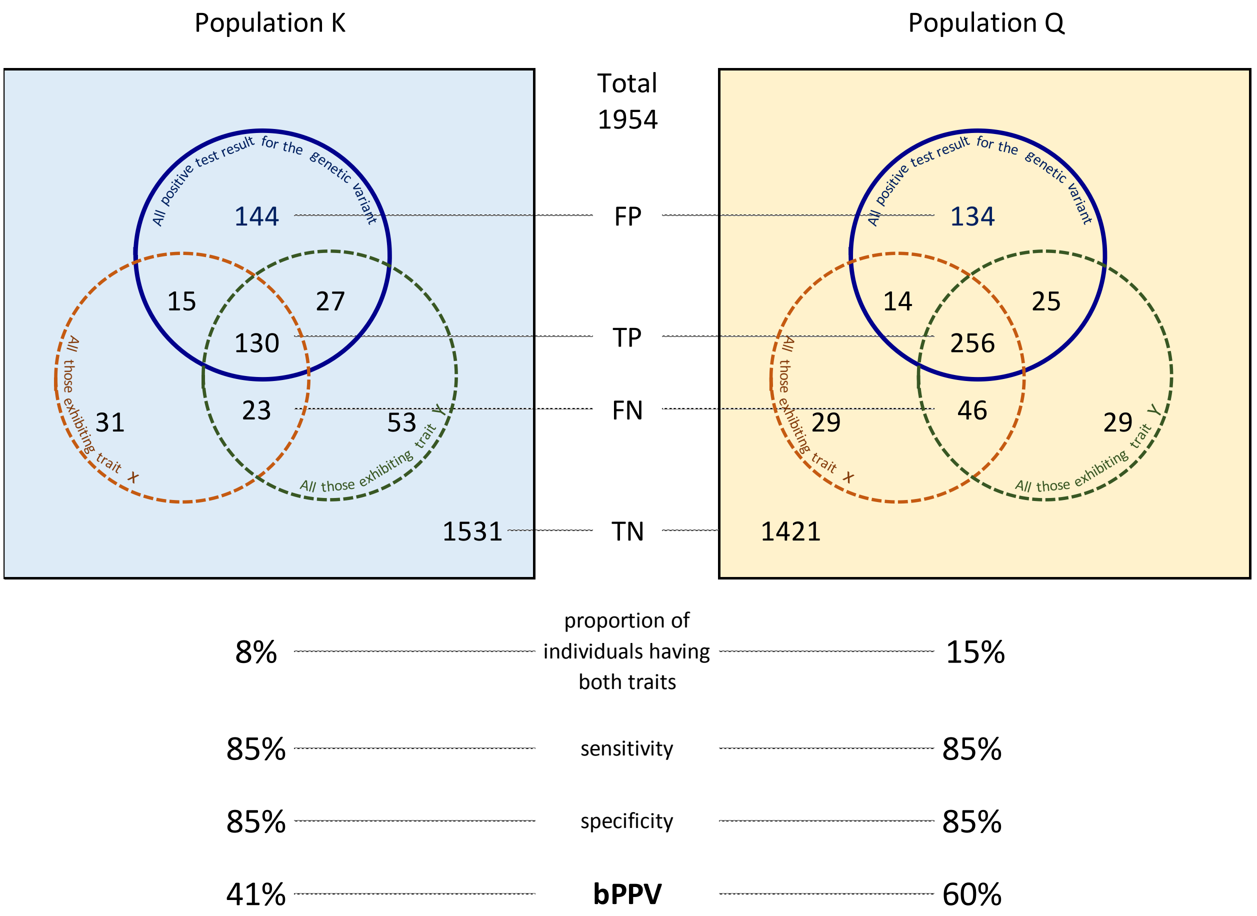

**Figure 12.5-1** Venn diagrams of on the determination of the bivariate positive predictive value (bPPV) of a genetic screening test in two populations of size n=1954. Both populations have a sensitivity of 85% and a specificity of 85%. The bivariate positive predictive value in population K is 41% and in population K is 60%. One of the reason for this difference is that only 8% of individual display both traits, whereas in population Q is 15%. ( **FP**, False positive; **TP**, True Positive; **FN**, False Negative, **TN**, True Negative)

#### Appendix 1. Literature sources of data use in the study

[L001] Aasmundstad T, Kongsro J, Wetten M, Dolvik NI, Vangen O. 2013. Osteochondrosis in pigs diagnosed with computed tomography: heritabilities and genetic correlations to weight gain in specific age intervals. Animal 7(10):1576-82. doi: 10.1017/S1751731113001158.

[L002] Abell, Caitlyn; Stalder, Kenneth J.; and Mabry, John W. (2011) "Genetic and Phenotypic Correlations for Maternal and Postweaning Traits from a Seedstock Swine Breeding System," *Animal Industry Report*: AS 657, ASL R2644.

[L003] Adolph, S.C., Hardin, J.S. 2007. Estimating phenotypic correlations: correcting for bias due to intraindividual variability. Functional Ecology 21:178-184.

[L004] Agrawal, A.A., Erwin, A.C., Cook, S.C. 2008. Natural selection on and predicted responses of ecophysiological traits of swamp milkweed (*Asclepias incarnata*). Journal of Ecology 96: 536–542 doi: 10.1111/j.1365-2745.2008.01365.x.

[L005] Akesson M, Bensch S, Hasselquist D, Tarka M, Hansson B 2008. Estimating Heritabilities and Genetic Correlations: Comparing the 'Animal Model' with Parent-Offspring Regression Using Data from a Natural Population. PLoS ONE 3(3): e1739. doi:10.1371/journal.pone.0001739

[L006] Alipanah, M., Deljo, J., Rokouie, M., Mohammadnia, R. 2013. Heritabilities and genetic and phenotypic correlations of egg quality traits in Khazak layers. Trakia Journal of Sciences 2:175-180.

[L007] Andrade, J.A.C., Filho J.B. M., 2006. Quantitative variation in the tropical maize population, ESAL-PBI. Sci. Agric. (Piracicaba, Braz.), v.65, n.2, p.174-182, March/April 2008.

[L008] Andrews C. P., Loeske E. B. Kruuk, Per T. Smiseth 2017. Evolution of elaborate parental care: phenotypic and genetic correlations between parent and offspring traits. *Behav Ecol* 2017; 28 (1): 39-48. doi: 10.1093/beheco/arw129

[L009] Araujo, M.R.A., Coulman, B. 2004. Genetic variation and correlation of agronomic traits in meadow bromegrass (*Bromus riparius* Rehm) clones. Ciência Rural, Santa Maria, v.34, n.2, p.505-510, mar-abr, 2004.

[L010] Arnold, S. 1981. Behavioral variation in natural populations. I. Phenotypic, genetic and environmental correlations between chemoreceptive responses to prey in the Garter snake, *Thamnophis elegans*. Evolution 35(3) 489-509.

[L011] Aslam ML, Bastiaansen JW, Crooijmans RP, Ducro BJ, Vereijken A, Groenen MA. 2011.Genetic variances, heritabilities and maternal effects on body weight, breast meat yield, meat quality traits and the shape of the growth curve in turkey birds. BMC Genet. 2011 Jan 25;12:14. doi: 10.1186/1471-2156-12-14.

[L012] Bhat, S.S., N. B. Singh, N. B. , Sankhyan,H. P. , Sharma, K. R. 2016. Variability studies for needle and wood traits of different half sib progenies of *Pinus roxburghii* Sargent. Physiol Mol Biol Plants 22(2):231–239. DOI 10.1007/s12298-016-0358-y

[L013] Baltunis, B.S., Brawer, J.T. 2010. Clonal stability in *Pinus radiata* across New Zealand and Australia. I. Growth and form traits. New Forests 40:305-322. doi 10.1007/s11056-010-9201-4.

[L014] Balzani, A., Cordell, H.J., Sutcliffe, E., Edwards, S. A. 2016. Heritability of udder morphology and colostrum quality traits in swine. J. Anim. Sci. 2016.94:3636–3644. doi:10.2527/jas2016-0458.

[L015 ] Barrosa,J., L.A. Velascoa, F.M. Winkler 2018. Heritability, genetic correlations and genotype by environment interactionsin productive traits of the Caribbean scallop, Argopecten nucleus (Mollusca:Bivalvia). Aquaculture 488:39-48.10.1016/j.aquaculture.2018.01.011

[L016] Bastarrachea, R.A., J.W. Kent, G. Rozada, et al. 2007. Heritability and genetic correlations of metabolic disease-related phenotypes in Mexico: Preliminary report from the GEMM Family study. Human Biology 79:121-129. doi 10.1353/hub.2007.0021.

[L017] Bedard, P.R., Hsu, C.S., Spangelo, L.P.S., Fejer, S.O., Rousselle G.L. 1971 Genetic, phenotypic and environmental correlations among 28 fruit and plant characters in the cultivated strawberry. Can. J. Gen. Cyt 13:470-479. doi: 10.1139/g71-071.

[L018] Benyamin, B., T. I. A. Sørensen, K. Schousboe, M. Fenger & P. M. Visscher & K. O. Kyvik 2007. Are there common genetic and environmental factors behind the endophenotypes associated with the metabolic syndrome? .Diabetologia 50:1880–1888. DOI 10.1007/s00125-007-0758-1

[L019] Bian, L., J. Shi, R. Zheng, J. Chen, H.X. Wu 2014. Genetic parameters and genotype-environment interactions of Chinese fir (*Cunninghamia lanceolata*) in Fujian province. Can. J. For. Res. 44:582-592. doi 10.1139/cjfr-2013-0427.

[L020] Bohlouhi, M., Alijami, S., Vaposhti, M.R. 2015. Genetic relationships among linear type traits and milk production traits of Holstein dairy cattle. Ann. Anim. Sci. 15(4)903-917.

[L021] Bolund, E.,H.Schielzeth, W. Forstmeier 2011.Correlates of male fitness in captive zebra finches - a comparison of methods to disentangle genetic and environmental effects. BMC Evolutionary Biology 2011 11:327.

[L022] Buchanan, J. W. , J. M. Reecy D. J. Garrick Q. Duan,D. C. Beitz, R. G. Mateescu 2015. Genetic parameters and genetic correlations among triacylglycerol and phospholipid fractions in Angus cattle.J. Anim. Sci. 2015.93:522–528. doi:10.2527/jas2014-8418.

[L023] Bouffier, L., P. Rozenberg, A. Raffin, A. Kremer 2008. Wood density variability in successive breeding populations of maritime pine. Can. J. For. Res. 38:2148-2158.

[L024] Buitenhuis, A. J., Rodenburg, T. B., Wissink,P. H. , Visscher,J., Koene,P. ,Bovenhuis,H. , Ducro, B. J., van der Poel, J. J. 2004. Genetic and Phenotypic Correlations Between Feather Pecking Behavior, Stress Response, Immune Response, and Egg Quality Traits in Laying Hens. Poultry Science 83:1077–1082. doi: 10.1093/ps/83.7.1077

[L025] Campo, J.L., M.G. Gil, S.G. Davila, I. Muñoz 2005. Estimation of heritability for fluctuating asymmetry in chickens by restricted maximum likelihood. Effects of age and sex. Poultry Science 84:1689-1697.

[L026] Canaza-Cayo, A. W., T. Huanca, J.P. Gutiérrez, P.A. Beltrán 2015. "Modelling of growth curves and estimation of genetic parameters for growth curve parameters in Peruvian young llamas (*Lama lama*)." *Small Ruminant Research* 130 (2015): 81-89.

[L027] Case, L.A., Wood, B.J., Miller, S.P. 2012. The genetic parameters of feed efficiency and its component traits in the turkey (*Meleagris gallopavo*). Genetics Selection Evolution 2012, 44:2.

[L028] Castañeda-Bustos, V.J., Montaldo H.H., Valencia-Posadas, M., Shepard, L., Perez-Elizalde, S., Hernandez-Mendo, O., Torres-Hernandez, G. 2017. Linear and nonlinear genetic relationships beyween type traits and productive life in US dairy goats. J. Dairy Sci. 100:1232-1245. Doi: 10.3168/jds.2016-11313.

[L029] Ceacero TM, Mercadante MEZ, Cyrillo JNdSG, Canesin RC, Bonilha SFM, de Albuquerque LG (2016) Phenotypic and Genetic Correlations of Feed Efficiency Traits with Growth and Carcass

Traits in Nellore Cattle Selected for Postweaning Weight. PLoS ONE 11(8): e0161366. doi:10.1371/journal.pone.0161366.

[L030] Chakraborty, S., H.K. Borah, B.K. Borah, D. Pathak, A.S.N. Zaman, B. K. Baruah, H. Kalita, B. Barman 2011. Genetic correlation and coheritability between $F_1$ and $F_2$ generations for quantitative traits in some crosses of green gram (*Vigna radiata* L. Wilczek). Acta Agronomica Hungarica 59(1)57-63. doi 10.1556/AAgr.59.2011.1.6.

[L031] Charmantier A., C. Perrins, R.H. McCleery, B.C. Sheldon 2006. Quantitative genetics of age at reproduction in wild swans: support for anatagonistic pleiotropy models of senescence. PNAS103:6587-6592.

[L032] Chen, X., Hawkings, B., Xie, C.Y., Ying, C.C. 2003. Age trends in genetic parameter and early selection of lodgepole pine provenances with particular reference to the Lambeth model. Forest Genetics 10:249-258.

[L033] Chen, Z.Q., B. Karlsson, T. Mörling, L. Olsson, E.J. Mellerowicz, H.X. Wu, S.O. Lundqvist, M.R.G Gil. 2016. Genetic analysis of fiber dimensions and their correlation with stem diameter and solid-wood properties in Norway spruce. Tree Genetics and Genomes 12:123. Doi: 10.1007/s11295-016-1065-0.

[L034] Chigezaa,G., Mashingaidzea, K., Shanahan, P. 2014. Combining ability,indirect and correlated response to selection for oil yield in sunflower(*Helianthus annuus*) under contrasting moisture environments. Field Crops Research 167:40–50.

[L035] Chiow, H.Y., J.C. Wyne 1983. Heritabilities and genetic correlations for yield and quality traits of advanced generation in a cross of peanut. Peanut Science 10:13-17.

[L036] Cho, K.H., Cho, C.I., Lee, J.H. · Park, K.D 2015. (Co)heritability of acetone and β-hydroxybutyrate concentrations in raw milk related to ketosis in Holsteins. Journal of the Korean Data & Information Science Society 2015, 26(4), 915–921. doi.org/10.7465/jkdi.2015.26.4.915 [korean]

[L037] Clapperton, M., AB Diack, O Matika, E J Glass, C D Gladney, M A Mellencamp, A Hoste, SC Bishop 2009. Traits associated with innate and adaptive immunity in pigs: heritability and associations with performance under different health status conditions. Genetics Selection Evolution 41:54. doi:10.1186/1297-9686-41-54.

[L038] Cole JB, Manyama M, Larson JR, Liberton DK, Ferrara TM, Riccardi SL, Li M, Mio W, Klein OD, Santorico SA, Hallgrímsson B, Spritz RA. 2017. Human Facial Shape and Size Heritability and Genetic Correlations. Genetics 205(2):967-978. doi: 10.1534/genetics.116.193185. Epub 2016 Dec 14.

[L039] Conner, J., Via, S. 1993. Patterns of Phenotypic and Genetic Correlations among Morphological and Life-HistoryTraits in Wild Radish, *Raphanus raphanistrum*. Evolution 47(2)704-711.

[L040] Cotter, S.C., L.E.B. Kruuk, K. Wilson 2004. Costs of resistance: genetic correlations and potential trade-offs in an insect immune system. J. Evol. Biol. 17:421-429. Doi 10.1046/j.1420-9101.2003.00655.x.

[L041] De Risio, L., J. Freeman, J., Lewis, T. 2016. Prevalence, heritability and genetic correlations of congenital sensorineural deafness and coat pigmentation phenotype in the English bull terrier. BMC Vet Res. 2016; 12: 146. doi: 10.1186/s12917-016-0777-6.

[L042] de Souza, V.A.B.d., Byrne, D.H. 1998. Heritability and phenotypic correlations, and predicted selection response of quantitative traits in peach: II. An analysis of several fruit traits. J. Amer. Soc. Hort. Sci 123(4)604-611

[L043] DeStephano, A., Seshadri, S., Beiser, A., Atwood, L.D., Massaro, J.M., Au, R., Wolf, P.A., DeCarli, C. 2009. Bivariate Heritability of Total and Regional Brain Volumes: The Framingham Study. Alzheimer Disease & Associated Disorders. 23(3):218-223. DOI: 10.1097/WAD.0b013e31819cadd8.

[L044] Diao, S., Hou, Y., Xie, Y., and Sun, X. 2016. Age trends of genetic parameters, early selection and family by site interactions for growth traits in *Larix kaempferi* open-pollinated families. BMC Genetics 17:104

[L045] Dochtermann, N.A., 2011. Testing Cheverud's conjecture for behavioral correlations and behavioral syndromes. Evolution 65-6: 1814–1820. doi:10.1111/j.1558-5646.2011.01264.x.

[L046] Donoghue, K.A., T. Bird-Gardiner, P.F. Arthur, R.M. Herd, R.F. Hegarty 2016. Genetic and phenotypic variance and covariance components for methane emission and postweaning traits in Agnus cattle. J. Anim. Sci. 94:1438-1445. doi: 10.2527/jas2015-0065.

[L047] Farias, W.J., Winkler, F.M., Brokordt, K.B. 2017. Genotype by environment interaction, heritabilities and genetic correlations for productive traits of *Haliotis rufescens*. Aquaculture doi: 10.1016/j.aquaculture.2017.02.030.

[L048] Ferreira, R. J. , S,F. M. Bonilha, F. M. Monteiro 2018. Evidence of negative relationship between female fertility and feed efficiency in Nellore cattle. Journal of Animal Science, sky276, 07 July 2018. doi.org/10.1093/jas/sky276.

[L049] Frigo, E., A. B. Samorè, D. Vicario, A. Bagnato, O. Pedron 2013. Heritabilities and Genetic Correlations of Body Condition Score and Muscularity with Productive Traits and their Trend Functions in Italian Simmental Cattle. Italian Journal of Animal Science, 12:2, e40, DOI: 10.4081/ijas.2013.e40.

[L050] Funk, W.C., Tyburczy, JA, Knudsen, K.L., Lindner, K.R., Allendorf, F,W. 2005. Genetic basis of variation in morphological and life-history traits of a wild population of pink salmon. Journal of Hederity 96(1)24-31. doi 10.1093/jhered/esi009.

[L051] Garcia D., V. Le Guen; C. R. Reis Mattos; P. de Souza Gonçalves, A. Clément-Demange 2002. Relationships between yield and some structural traits of the laticiferous system in *Hevea* clones resistant to South American leaf blight. *Crop Breeding and Applied Biotechnology, v. 2, n. 2, p. 307-318.*

[L052] Gaspar,M.J., Louzada, J.L., Aguiar, A., Almeida, M.H. 2008. Genetic correlations between wood quality traits of *Pinus pinaster* Ait. Ann. For. Sci. 65:703. doi: 10.1051/forest:2008054.

[L053] Gaspar,M.J., J.L. Louzada, M.E. Silva, A. Aguiar, M.H. Almeida 2008. Age trends in genetic parameters of wood density components in 46 half-sibling families of *Pinus pinaster*. Can. J. For. Res. 38:1470-1477.

[L054] Gezan, S.A., M.P. d. Carvalho, J. Sherrill 2017. Statistical methods to explore genotype-by-environment interaction for loblolly pine clonal trials. Tree Genetics and Genomes 13:1. doi: 10.1007/s1195-016-1081-0.

[L055] Glahn DC, Curran JE, Winkler AM, Carless MA, Kent JW Jr, Charlesworth JC, Johnson MP, Göring HH, Cole SA, Dyer TD, Moses EK, Olvera RL, Kochunov P, Duggirala R, Fox PT, Almasy L, Blangero J. 2012. High Dimensional Endophenotype Ranking in the Search for Major Depression Risk Genes. *Biological Psychiatry 71*(1)6–14. doi.org/10.1016/j.biopsych.2011.08.022

[L056] Guinez, R., J. E. Toro, S. Krapivka, A. C. Alcapan, P. A. Oyarzun 2017. Heritabilities and genetic correlation of shell thickness and shell length growth in a mussel, *Mytilus chilensis* (Bivalvia:Mytilidae). Aquaculture Research, 2017, 48, 1450–1457.doi:10.1111/are.12981.

[L057] Hadfield, J.D., Nutall, A., Ososrio, D., Owens, I.P.F. 2006. Testing the phenotypic gambit: phenotypic, genetic and environmental correlations of colour. Journal of Evolutionary Biology 20:549–557. doi:10.1111/j.1420-9101.2006.01262.x.

[L058] Haffray, P., Bugeon, J, Pincent, C., Chapuis, H., Mazeiraud, E., Rossignol, M.-N., Chatain, B., Vandeputte M, Dupont-Nivet, M. 2012. Negative genetic correlations between production traits and head or bony tissues in large all-female rainbow trout (*Oncorhynchus mykiss*). Aquaculture Vols. 368–369:145–152. doi 10.1164/rccm.201302-0263OC

[L059] Hai, P.H., Duong, L.A., Toan, N.Q., Ha, T.T.T. 2005.Genetic variation in growth, stem straightness, pilodyn and dynamic modulus of elasticity in second-generation progeny tests of *Acacia mangium* at three sites in Vietnam. New Forests (2015) 46:577–591. doi 10.1007/s11056-015-9484-6.

[L060] Hamzah, A., Nguyen, N.H., Mekkawy, W., Ponzoni, R.W., Khaw, H.L., Yee, H.Y., Bakar, K.R.A., Nor, S.A.M. 2016. Flesh characteristics: estimation of genetic parameters and correlated responses to selection for growth rate in the GIFT strain. Aquaculture Research 47:2139–2149. doi:10.1111/are.12667.

[L061] Hannrup, B., Ekberg, I. 1998. Age-age correlations for tracheid length and wood density in Pinus sylvestris. Can. J. For. Res. 28(9)1373-1379.

[L062] Hansen, T. F. , W. S. Armbruster, M. L. Carlson, C. Labon 2003. Evolvability and Genetic Constraint in *Dalechampia* Blossoms: Genetic Correlations and Conditional Evolvability. J. Exp. Zool 296B:23–39. DOI: 10.1002/jez.b.00014

[L063] S. Hatcher, K.D Atkins 2000. Breeding objectives which Include fleece weight and fibre diameter do not need fibre curvature. Asian-Aus. J. Anim. Sci. 13 Supplement July 2000 A:293-296.

[L064] Hatcher, S. , P. I. Hynd A,C , K. J. Thornberry, S. Gabb 2010. Can we breed Merino sheep with softer, whiter, more photostable wool? Animal Production Science 50:1089–1097.

[L065] Hein, P.R.G., J.M. Bouvet, E. Mandrou, P. Vigneron, B. Clair, G. Chaix. 2012. Age trends in microfibril angle inheritance and their genetic and environmental correlations with growth, density and chemical properties in *Eucalyptus urophylla* S.T. Blake wood. Annals of Forest Science 69:681-691. doi 10.1007/s13595-012-0186-3.

[L066] Hlusko, L.J., Mahaney, M.C. 2009. Quantitative Genetics, Pleiotropy, and Morphological Integration in the Dentition of *Papio hamadryas*. Evol Biol 36:5–18. DOI 10.1007/s11692-008-9048-1

[L067] Hong, Z., A. Fries, H.X. Wu 2014. High negative genetic correlations between growth traits and wood properties suggest incorporating multiple traits selection including economic weights for the future Scots pine breeding programs. Annals of Forest Science 71:463.472. doi: 10.1007/s13595-014-0359-3.

[L068] Horimoto, A.R.V.R. , J.B.S. Ferraz, J.C.C. Balieiro and J.P. Eler 2007. Phenotypic and genetic correlations for body structure scores (frame) with productive traits and index for CEIP classification in Nellore beef cattle. Genet. Mol. Res. 6 (1): 188-196

[L069] Hossein-Zadeh, N. G. 2015. Estimation of genetic relationships between growth curve parameters in Guilan sheep. J. Anim. Sci. Tech. 57:19. doi 10.1186/s40781-015-0052-6.

[L070] Hu, G., Gu, W., Bai, Q.L., Wang, B.Q. 2013. Estimation of genetic parameters for growth traits in a breeding program for rainbow trout (*Oncorhynchus mykiss*) in China. Genetics and Molecular Research 12 (2): 1457-1467. doi: 10.4238/2013.April.26.7.

[L071] Ikonen T., Morri S, Tyrisevä AM, Ruottinen O, Ojala M. 2004. Genetic and phenotypic correlations between milk coagulation properties, milk production traits, somatic cell count, casein content, and pH of milk. J Dairy Sci. 2004 Feb;87(2):458-67.

[L072] Ilatsia, E.D., Githinji, M.G., Muasya, T.K., Okeno, T.O., Kahi, A.K. 2008. Genetic parameter estimates for growth traits of Large White pigs in Kenya. South African Journal of Animal Science 38(3)166-173.

[L073] Jensen, H., Saether, B.-E., Ringsby, T.H., Tufto, J., Griffith, S.C., Ellengree, H. 2003. Sexual variation in heritability and genetic correlations of morphological traits in house sparrow (*Passer domesticus*). J. Evol. Biol. 16:1209-1307. Doi: 10.1046/j.1420-9101.2003.00614.x.

[L074] Johnson, S., Johnson, R. F. 1990. Environmental variation and quantitative genetic parameters in the feral pigeon, Columba livia. Biological Journal of the Linnean Society 40:321-332.

[L075] Kanai, M., M. Akiyama, A. Takahashi, N. Matoba, Y. Momozawa, M. Ikeda, N. Iwata, S. Ikegawa, M. Hirata, K. Matsuda, M. Kubo, Y. Okada, Y. Kamatani 2018. Genetic analysis of quantitative traits in the Japanese population links cell types to complex human diseases. Nature Genetics 50: 390–400. doi: 10.1038/s41588-018-0047-6.

[L076] Karasik, D., Serkalem Demissie,S., Zhou, Y., Lu, D., Broe, K.E., Bouxsein, M.L., Cupples, L.A., Kiel D.P. 2016. Heritability and Genetic Correlations for Bone Microarchitecture: The Framingham Study Families. Journal of Bone and Mineral Research 32(1):106–114. DOI: 10.1002/jbmr.2915

[L077] Kesbi, F.G., M. Eskandarinasab, A. Hassanabadi 2008. Estimation of genetic parameters for lamb weight at various ages in Mehaban sheep. Ital. J. Anim. Sci. 7:95-103.

[L078] Kjaer, J. B., Sørensen, P. 1997. Feather pecking behaviour in White Leghorns, a genetic study, British Poultry Science, 38:4, 333-341. doi: 10.1080/00071669708417999.

[L079] Knauer,M. T. , J. P. Cassady D. W. Newcom M. T. See 2014. Phenotypic and genetic correlations between gilt estrus, puberty, growth, composition, and structural conformation traits with first-litter reproductive measures. J. Anim. Sci. 2011. 89:935–942. doi:10.2527/jas.2009-2673.

[L080] Kogelmann L.J.A., H.N. Kadarmideen, T. Mark, P. Karlskov-Mortensen, C.S. Bruun, S. Cirera, M.J. Jacobson, C.B. Jorgensen, M. Fredholm 2013. A F2 pig resource population as a model for genetic studies of obesity and obsesity-rrelated diseases in humans: design and genetics parameters. Frontiers in Genetics 4(29). doi: 10.3389/fgene.2013.00029.

[L081] Kouassi A.B, Durel, C., Costa, F., Tartarini, S., van de Weg, E., Evans, K., Fernandez-Fernandez, F., Govan, C., Boudichevskaja, A., Dunemann, F., Antofie, A., Lateur, M., Stankiewicz-Kosyl, Soska, A., Tomala, K., Lewandowski, M., Rutkovski, K., Zurawicz, E., Guerra, W. , Laurens F. 2009. Estimation of genetic parameters and prediction of breeding values for apple fruit-quality traits using pedigreed plant material in Europe. Tree Genetics & Genomes Volume 5(4) 659–672. doi 10.1007/s11295-009-0217-x.

[L082] Lai, M, L. Dong, M. Yi, S. Sun, Y. Zhang, L. Fu, Z. Xu, L. Lei, C. Leng., L. Zhang 2017. Genetic variation, heritability and genotype x environment interactions of resin yield, growth traits and morphologic traits for *Pinus elliottii* at three progeny trials. Forests 8:409. doi 10.3390/f8110409.

[L083] Leng Y, Shi Y, Yu Q, Van Horn JD, Tang H, Li J, Xu W, Ge X, Tang Y, Han Y, Zhang D, Xiao M, Zhang H, Pang Z, Toga AW, Liu S. 2016. Phenotypic and Genetic Correlations Between the Lobar Segments of the Inferior Fronto-occipital Fasciculus and Attention. Sci Rep. 2016 (Sep 6) 6:33015. doi: 10.1038/srep33015.

[L084] Lenz, P., Auty, D., Achim, A., Beaulieu, J., Mackay, J. 2013. Genetic Improvement of White Spruce Mechanical Wood Traits—Early Screening by Means of Acoustic Velocity. Forests 2013, 4, 575-594; doi:10.3390/f4030575.

[L085] Li, C., O. Weng, J.-B. Chen, M. Li, C. Zhou, S. Chen, W. Zhou, D. Guo, C. Lu, J.-C. Chen, D. Xiang, S. Gan. 2017. Genetic parameters for growth and wood mechanical properties in *Eucalyptus cloeziana* F. Muell. New Forests 48: 33–49.

[L086] Lira, E.G., R.F. Ambile, M. Fagioli, A. P. L. Montalvao 2017. Genetic parameters, phenotypic, genotypic and environmental correlations and genetic variability on sunflower in the Brazilian savannah. Ciencia Rural 47:08 e20160719. Doi: 10.1590/0103-847cr20160719.

[L087] Liu Feng, Li Yangzhen , Du Min,Shao Changwei, Chen Songlin 2016. Analysis of phenotypic and genetic parameters for growthrelated traits in the half smooth tongue sole, *Cynoglossus* *semilaevis.* Chinese Journal of Oceanology and Limnology 34( 1) 163-169. Doi 10.1007/s00343-016-5044-y.

[L088] Lukowski, S.W., Lloyd-Jones, L.R., Holloway, A. 2017. Genetic correlations reveal the shared genetic architecture of transcription in human peripheral blood. Nature Communications 8: 483. doi 10.1038/s41467-017-00473-z.

[L089] Manjula,P., Park,H.-B.,Seo, D. Choi,N., Jin, S., Ahn,S.J., Heo,K.N., Kang,B.S., Lee,J.H. 2018. Estimation of heritability and genetic correlation of body weight gain and growth curve parameters in Korean native chicken. Asian-Australas J Anim Sci.31(1): 26–31. doi:10.5713/ajas.17.0179

[L090] Martínez , J.J., M. C. de Aranzamendi, E. H. Bucher 2018. Quantitative genetics in the monk parakeet (*Myiopsitta monachus*) from central Argentina: Estimation of heritability and maternal effects on external. PLoS ONE 13(8): e0201823. doi.org/10.1371/journal.pone.0201823

[L091] Matebesi,P.A., S.W.P. Cloete, and J.B. van Wyk 2009. Genetic parameter estimation of 16-month live weight and objectively measured wool traits in the Tygerhoek Merino flock. South African Journal of Animal Science 39(1)73-82.

[L092] McAdam, E.L., R. E. Vaillancourt, A. Koutoulis, S. P. Whittock 2014. Quantitative genetic parameters for yield, plant growth and cone chemical traits in hop (*Humulus lupulus L.)*. BMC Genetics 2014, 15:22. doi:10.1186/1471-2156-15-22.

[L093] Messina FJ, Fry JD. 2003. Environment-dependent reversal of a life history trade-off in the seed beetle Callosobruchus maculatus. J Evol Biol. 2003 May;16(3):501-9.

[L094] Miar Y, Plastow G, Bruce H, Moore S, Manafiazar G, et al. (2014) Genetic and Phenotypic Correlations between Performance Traits with Meat Quality and Carcass Characteristics in Commercial Crossbred Pigs. PLoS ONE 9(10): e110105. doi:10.1371/journal.pone.0110105.

[L095] Miguez-Soto, B., J. Fernandez-Lopez 2012. Genetic parameters and predicted selection responses for timber production traits in a *Castanea sativa* progeny trial: developing a breeding program. Tree Genetics and Genomes 8:409-423.

[L096] Mihai, G., Mirancea, I. 2016. Age trends in genetic parameters for growth and quality traits in Abies alba. Forest 9: 954-959. – doi: 10.3832/ifor1766-009 [online 2016-07-07].

[L097] Missanjo,E., Imbayarwo-Chikosi, V., Halimani, T. 2013. Estimation of Genetic and Phenotypic Parameters for Production Traits and Somatic Cell Count for Jersey Dairy Cattle in Zimbabwe. ISRN Veterinary Science Article ID 470585, 5 p. http:/ dx.doi.org/10.1155/2013/470585.

[L098] Missanjo, E., Matsumura, J. 2016. Genetic Improvement of wood properties in *Pinus kesiya Royle ex Gordon* for Sawn Timber Production in Malawi. Forests 7:253; doi:10.3390/f7110253.

[L099] Montaldo, H.H., Trejo, C., Lizana, C. 2017. Genetic parameters for milk yield and reproduction traits in the Chilean Dairy Overo Colorado cattle breed. Cien. Inv. Agr. 44(1):24-34. doi 10.7764/rcia.v44i1.1562.

[L100] Montes, C.S., Beaulieu, J., Hernandez, R.E. 2007. Genetic variation in wood shrinkage and its correlations with tree growth and wood density of *Calycophyllum spruceanum* at an early age in the Peruvian Amazon. Can. J. For. Res. 37:966-976.

[L101] Muñoz, M., Loretto; Jaju, Deepali; Voruganti, Saroja; Albarwani, Sulayma; Aslani, Afshin; Bayoumi, Riad; Al-Yahyaee, Said; Comuzzie, Anthony, G.; Millar, Philip, J.; Picton, Peter; Floras, John, S.; Nolte, Ilja; Hassan, Mohammed, O.; Snieder, Harold 2018. Heritability and genetic correlations of heart rate variability at rest and during stress in the Oman Family Study. Journal of Hypertension: July 2018 - Volume 36 - Issue 7 - p 1477–1485 doi: 10.1097/HJH.0000000000001715.

[L102] Mustiga, G. M. , S. A. Gezan, W. Phillips-Mora, A. Arciniegas-Leal, A. Mata-Quirós, J. C. Motamayor 2018. Phenotypic Description of *Theobroma cacao L.* for Yield and Vigor Traits From 34 Hybrid Families in Costa Rica Based on the Genetic Basis of the Parental Population.Front. Plant Sci., 19 June 2018 | https://doi.org/10.3389/fpls.2018.00808.

[L103] Narinc, D. , Aksoy, T., Karaman, E., Aygun, A., Firat, M.Z., Uslu, M.K. 2013. Japanese quail meat quality: Characteristics, heritabilities, and genetic correlations with some slaughter traits. Poultry Science 92 :1735–1744. doi 10.3382/ps.2013-03075

[L104] Navarro, A., Zamorano, M.J., Hildebrandt, S., Ginés, R., Aguilera, C., Afonso, J.M. 2009a. Estimates of heritabilities and genetic correlations for body composition traits and G×E interactions, in gilthead seabream (*Sparus auratus L.)*. Aquaculture 295:183–187.

[L105] Navarro, A., Zamorano, M.J., Hildebrandt, S., Ginés, R., Aguilera, C., Afonso, J.M. 2009b. Estimates of heritabilities and genetic correlations for growth and carcass traits in gilthead seabream (*Sparus auratus L.)* under artificial conditions. Aquaculture 289:225-230. Doi: 10.1016/j.aquaculture.2008.12.024.

[L106] Nguyen, N.H., J. Quinn, D. Powell, A. Elizur, N.P. Thoa, J. Nocillado, R. Lamont, C. Remilton, W. Knibb 2014. Heritability for body colour and its genetic association with morphometric traits in Banana shrimp (*Fenneropenaeus merguiensis*). BMC Genetics 15:132. doi: 10.1186/s12863-014-0132-5

[L107] Nieuwhof,G. J. , Conington,J. , Bunger, L. , Haresign, W., Bishop, S. C. 2008. Genetic and phenotypic aspects of foot lesion scores in sheep of different breeds and ages. Animal 2:9, pp 1289–1296. doi:10.1017/S1751731108002577.

[L108] Okamura T., Maeda K, Onodera W, Kadowaki H, Kojima-Shibata C, Suzuki E, Uenishi H, Satoh M, Suzuki K. 2015. Correlated responses of respiratory disease and immune capacity traits of Landrace pigs selected for *Mycoplasma pneumoniae* of swine (MPS) lesion. Anim Sci J. 2016 Sep;87(9):1099-105. doi: 10.1111/asj.12560. Epub 2015 Nov 26.

[L109] Oliveira, C.A.L., R.P.Riberio, G.M.Yoshida, N.M. Kunita, G.S. Rizzato, S.N. d. Oliveira, A.I. d. Santos, N.H. Nguyen 2016. Correlated changes in body shape after five generations of selection to improve growth rate in a breeding program for Nile tilapia *Oreochromis niloticus* in Brazil. J. Appl. Genetics 57:487-493. Doi: 10.1007/s13353-016-0338-5.

[L110] Pelletier, M.C., Henson, M., Boyton, S., Thomas, D., Vanclay, J.K. 2008. Genetic variation in shrinkage properties of *Eucalyptus pilularis* assessed using increment cores and test blocks. New Zealand Journal of Forestry Science 38(1): 194–210.

[L111] Pertile, S.F.N.; Zampar, A.; Petrini, J.; Gaya, L.d.G.; Rovadoscki, G. A.; Ramírez-Díaz, J.; Ferraz, J. B. S.; Michelan-Filho, T.; Mourão, G. B. 2014. Correlated responses and genetic parameters for performance and carcass traits in a broiler line. Revista Brasileira de Saude e Producao Animal 15(4):1006-1016.

[L112] Pfrender, M.E., Lynch, M. 2000. Quantitative Genetic Variation in Daphnia: Temporal Changes in Genetic Architecture. Evolution 54(5)1502-1509.

[L113] Pike, C., Montgomery, R.A. 2007. Genetic and phenotypic correlations among volume, wood specific gravity and foliar traits in white spruce (*Picea glauca (Moench) Voss*). Silva Genetica 64(4)159-170. doi 10.1515/sg-2015-0015.

[L114] Porth, I., Klapste, J., Skyba, O., Lai, B.S.K., Geraldes, A., Muchero, W., Tuskan, G.A., Douglas, C.J., El-Kassaby, Y.A., Mansfield, S.D. 2013. *Populus trichocarpa* cell wall chemistry and ultrastructure trait variation, genetic control and genetic correlations. New Phytologist 197: 777–790. doi: 10.1111/nph.12014.

[L115] Poveda, A., M.E. Ibañez, E. Rebato 2012. Heritability and genetic correlations of obesity-related phenotypes among Roma people. Annals of Human Biology 39(3):183–189. doi 10.3109/03014460.2012.669794.

[L116] Pratt, P.J., Moser,D. W. , Thompson, L. D., Jackson, S. P., Johnson, B. J., Garmyn, A. J., Miller M. F. 2014. The heritabilities, phenotypic correlations, and genetic correlations of lean color and palatability measures from longissimus muscle in beef cattle. J. Anim. Sci. 91:2931–2937. doi:10.2527/jas2012-5662

[L117] Ratcliffe, B., Hart, F.J., Klápště, J., Jaquish, B., Mansfield, S.D., El-Kassaby, Y.A. 2014 . Genetics of wood quality attributes in Western Larch. Annals of Forest Science (2014) 71:415–424. DOI 10.1007/s13595-013-0349-x.

[L118] Rincón, J., Zambrano, J., Echeverri, J. 2015. Estimation of genetic and phenotypic parameters for production traits in Holstein and Jersey from Colombia. Rev.MVZ Córdoba 20(Supl):4962-4973.

[L119] Rodenburg, T. B., Buitenhuis, A.J., Ask, B., Uitdehaag, K. A., Koene, P., van der Poel, J. J., van Arendonk, J. A. M., Bovenhuis, H. 2004. Genetic and Phenotypic Correlations Between Feather Pecking and Open-Field Response in Laying Hens at Two Different Ages. Behavior Genetics 34(4):407–415.

[L120] Roshanfekr, H., Berg, P., Mohammadi, K., Mohamadi, E.M. 2015. Genetic parameteres and genetic gains for reproductive traits of Arabi sheep. Biotechnology in Animal Husbandry 31(1)23-36. doi 10.2298/BAH1501023R.

[L121] Rowan, D.D., M. B. Hunt, P. A. Alspach, C. J. Whitworth, N. C. Oraguzie 2009. Heritability and genetic and phenotypic correlations of apple (*Malus × domestica*) Fruit volatiles in a genetically diverse breeding population. J. Agric. Food Chem., 2009, 57 (17), pp 7944–7952. doi 10.1021/jf901359r.

[L122] Sampson J.N., W.A. Wheeler, M. Yeager, et al. 2015. Analysis of heritability and shared heritability based on Genome-wide association studies for 13 cancer types. JNCI Natl Cancer Inst 107(12)djv279. Doi 10.1093/jnci/djv279.

[L123] Shen, K.K., V. Doré, S. Rose, J.Fripp, K.L. McMahon, G.I.de Zubicaray, N.G. Martin, P.M. Thompson, M.J. Wright, O. Salvado 2016. Heritability and Genetic Correlation Between the Cerebral Cortex and Associated White Matter Connections. Hum Brain Mapp. 37(6):2331–2347. doi10.1002/hbm.23177.

[L124] Shiferaw, W., Balcha, A., Mohammed, H. 2012. Genetic Variation for Grain Yield and Yield Related Traits in Tef [*Eragrostis tef (Zucc.)Trotter*] under Moisture Stress and Non-Stress Environments. American Journal of Plant Sciences, 2012, 3, 1041-1046 http://dx.doi.org/10.4236/ajps.2012.38124.

[L125] Simmons, L.W., Kotiaho, J.S. 2002. Evolution of Ejaculates: Patterns of Phenotypic and Genotypic Variation and Condition Dependence in Sperm Competition Traits. Evolution 56(8)1622-1631.

[L126] Sodini, S.M., K. E. Kemper, N. R. Wray, M. Trzaskowski 2018. Comparison of Genotypic and Phenotypic Correlations: Cheverud's Conjecture in Humans. Genetics Early online May 8, 2018; https://doi.org/10.1534/genetics.117.300630.

[L127] Spieker,E.A., P. Kochunov, L.M. Rowland, E. Sprooten, A.M. Winkler, Olvera, R.L., Almasy, L., R. Duggirala, P. T. Fox, J. Blangero, D. C. Glahn, J. E. Curran 2015. Shared genetic variance between obesity and white matter integrity in Mexican Americans. Front. Genet., 13 February 2015. doi 10.3389/fgene.2015.00026.

[L128] Stackpole, D.J., Vaillancourt, R.E., de Aguigar, M., Potts, B.M. 2010. Age trends in genetic parameters for growth and wood density in *Eucalyptus globulus*. Tree Genetics & Genomes 6:179–193. doi 10.1007/s11295-009-0239-4.

[L129] Stephens, J.M., P.A. Alspach ,R.A. Beatson, C. Whinefield, E. J. Buck 2012. Genetic Parameters and Development of a Selection Index for Breeding Red Raspberries for Processing. J. Am. Soc. Hort. Sci. 137(4)236-242.

[L130] Sun, X., Lei,S.F., Deng, F.Y., Wu, S., Papacian,C., Hamilton, J. 2006. Genetic and Environmental Correlations between Bone Geometric Parameters and Body Compositions. Calcif Tissue Int 79:43-49. DOI: 10.1007/s00223-006-0041-3.

[L131] Svensson, J.C., S.E.McKeand, H.L. Allen, R.G. Campbell 1999. Genetic Variation in Height and Volume of Loblolly Pine Open-Pollinated Families During Canopy Closure. Silvae Genetica 48(3–4)204-208.

[L132] Taylor, R.W., Boon, A.K., Dantzer, B., Reale, D., Humphries, M.M., Boutin, S., Gorrell, J.C., Coltman, D.W., McAdam, A.G. 2012. Low heritabilities, but genetic and maternal correlations between red squirrel behaviours. J. Evol. Biol. doi: 10.1111/j.1420-9101.2012.02456.x.

[L133] Tian Yongsheng, Xu Tianjun, Liang You, Chen Songlin 2011. Estimates of genetic and phenotypic parameters for weight and length in *Paralichthys olivaceus* (Temminck et Schlegel). *Acta Oceanol. Sin*ica 30(6)58-64. DOI: 10.1007/s13131-011-0161-0

[L134] Torres-Vázquez, J.A., Valencia-Posadas, M., Castillo-Juárez, H., Montaldo, H.H. 2009. Genetic and phenotypic parameters of milk yield, milk composition and age at first kidding in Saanen goats from Mexico. Livestock Science 126 (2009) 147–153. doi:10.1016/j.livsci.2009.06.008

[L135] Traglia, M., Croen, L.A., Lyall, K., Windham, G.C., Kharrazi, M., DeLorenze, G.N., Torres, A.R., Weiss, L.A. Independent maternal and fetal genetic effects on midgestational circulating levels of environmental pollutants. G3 7:1287-1299. doi 10.1534/g3.117.39784.

[L136] Ukrainetz, N.K., Kang, K.Y., Aitken, S.N., Stoehr, M., Mansfield, S.D. 2008. Heritability and phenotypic and genetic correlations of coastal Douglas-fir (*Pseudotsuga menziensii*) wood quality traits. Can. J. For. Res. 38:1536-1546. doi 10.1139/X07-234.

[L137] Vallas, M., Bovenhuis, H., Kaart, T., Pärna, K., Kiiman, H., Pärna, E. 2010. Genetic parameters for milk coagulation properties in Estonian Holstein cows. J. Dairy Sci. 93 :3789–3796. doi: 10.3168/jds.2009-2435.

[L138] Vandeputte, M., Puledda, A., Tyran, A.S., Bestin, A., Coulombet, C., Bajek, A., Baldit, G., Vergnet, A., Allal, A, Bugeon, J., Haffray, P. 2017. Investigation of morphological predictors of fillet and carcass yield in European sea bass (Dicentrarchus labrax) for application in selective breeding. Aquaculture 470 (2017) 40–49.

[L139] Vattikuti S, Guo J, Chow CC 2012. Heritability and Genetic Correlations Explained by Common SNPs for Metabolic Syndrome Traits. PLoS Genet 8(3):e1002637. doi:10.1371/journal.pgen.1002637.

[L140] Vergara OD, Ceron-Muñoz MF, Arboleda EM, Orozco Y, Ossa GA. 2009. Direct genetic, maternal genetic, and heterozygosity effects on weaning weight in a Colombian multibreed beef cattle population. J Anim Sci. 87(2):516-21. doi: 10.2527/jas.2007-0636. Epub 2008 Oct 24.

[L141] Verweij, N., van de Vegte, Y.J., van der Harst, P. 2018. Genetic study links components of the autonomous nervous system to heart-rate profile during exercise. Nature Communications 9:898. doi 10.1038/s41467-018-03395-6.

[L142] Volckaert, A.M., Bart Hellemans, Costas Batargias, Bruno Louro, Cécile Massault, Jeroen K J Van Houdt, Chris Haley, Dirk-Jan de Koning and Adelino VM Canario 2012. Heritability of cortisol response to confinement stress in European sea bass *Dicentrarchus labrax*. Genetics Selection Evolution 44:15.

[L143] Volker, P.W., C.A. Dean, W.N. Tibbits, I.C. Ravenwood 1990. Genetic parameters and gains expected from selection in Eucalyptus globulus in Tasmania. Silvae Genetica 39(1)18-21.

[L144] Wang K, Gaitsch H, Poon H, Cox NJ, Rzhetsky AQ 2017 Classification of common human diseases derived from shared genetic and environmental determinants. Nature Genetics 49:1319–1325.

[L145] Wang, M., Kong, J., Meng, X., Luan, S., Luo, K., Sui, J., Chen, B., Cao, J., Shi, X. 2017. Evaluation of genetic parameters for growth and cold tolerance traits in *Fenneropenaeus chinensis* juveniles. PLoS ONE 12 (8): e0183801. doi 10.1371/journal.pone.0183801.

[L146] Wang W, Zhao LJ, Liu YZ, Recker RR, Deng HW. 2006. Genetic and environmental correlations between obesity phenotypes and age at menarche. Int J Obes (Lond) 30(11):1595-600. Epub 2006 Mar 28.

[L147] Willmore, K.E., C.C. Roseman, J. Rogers, J. M. Cheverud, J. T. Richtsmeier 2009. Comparison of Mandibular Phenotypic and Genetic Integration between Baboon and Mouse. Evol Biol. 2009 March ; 36(1): 19–36. doi:10.1007/s11692-009-9056-9.

[L148] Wolcott, M. L., D. J. Johnston, S. A. Barwick, N. J. Corbet and P. J. Williams 2014. The genetics of cow growth and body composition at first calving in two tropical beef genotypes. Animal Production Science 54:37–49. doi 10.1071/AN12427.

[L149] Woo, J.G., Morrison, J.A., Stroop, D.M., Aronson-Friedman, L., Martin, L.J. 2014. Genetic architecture of lipid traits changes over time and differs differs by race: Princeton Lipid follow-up study. J. Lipid Res. 55(7)1515-1524.

[L150] Wu, T., Boezen,H.M., Postma,D.S., Los,H. , Postmus, P.E., Snieder, H., Boomsmae, D.I. 2010. Genetic and environmental influences on objective intermediate asthma phenotypes in Dutch twins. Eur Respir J 2010; 36: 261–268. DOI: 10.1183/09031936.00123909.

[L151] Wu, T., Treiber, F.A., H. Snider, H. 2013. Genetic influence on blood pressure and underlying hemodynamics measured at rest and during stress. Psychosom Med. 75(4): 404–412. doi:10.1097/PSY.0b013e31828d3cb6

[L152] Xu, L., Li, Q., Yu, H., Kong, L. 2017. Estimates of Heritability for Growth and Shell Color Traits and Their Genetic Correlations in the Black Shell Strain of Pacific Oyster *Crassostrea gigas*. Mar Biotechnol (2017) 19:421–429. DOI 10.1007/s10126-017-9772-6.

[L153] Yang, T.-L. , L.-J. Zhao, Y.-J. Liu, J.-F. Liu, R. R. Recker, H.-W. Deng 2006. Genetic and Environmental Correlations of Bone Mineral Density at Different Skeletal Sites in Females and Males. Calcif Tissue Int 78:212_217. DOI: 10.1007/s00223-005-0267-5.

[L154] Yang, Yan-Jun , V.Dvornyk, W.-X. Jian, S.-M. Xiao, H.-W. Deng 2005. Genetic and environmental correlations between bone phenotypes and anthropometric indices in Chinese. Osteoporos Int 16: 1134–1140. DOI 10.1007/s00198-004-1825-9

[L155] Yau, M.S., Demissie,S., Zhou, Y., Anderson, D.E., Lorbergs, A.L., Kiel, D.P., Allaire, B.T., Yang, L., Cupples, L. A., Travison, T.G., Bouxsein, M.L., Karasik, D., Samelson, E.J. 2016. Heritability of Thoracic Spine Curvature and Genetic Correlations with Other Spine Traits: The Framingham Study. Journal of Bone and Mineral Research 31(12):2077–2084. Doi 10.1002/jbmr.2925.

[L156] Zahid, M.A., Akhter, M., Sabar, Z Zaheen, M. and Awan, T. 2006. Correlation and path analysis studies of yield and economic traits in Basmati rice (*Oryza sativa L.)*. Asian J. Plant Sciences 5(4):643-645.

[L157] Zamudio, F., R. Baettyg, A. Vergara, F. Guerra, P. Rozenberg 2002. Genetic trends in wood density and radial growth with cambial age in a radiate pine progeny test. Am. For Sci. 59:541-549. Doi 10.1051/forest.2002039.

[L158] Zeng, L.H., Zhang, Q., He, B.X., Lian, H.M., Cai, Y.L., Wang, Y.S., Luo, M. 2013. Age trends in genetic parameters for growth and resin-yielding capacity in Masson pine. Silva Genetica 62:7-17.

[L159] Zhang, L.-C. , Ning, Z.-H., Xu, G.-Y. , Hou, Z.-C., Yang, N. 2005. Heritabilities and Genetic and Phenotypic Correlations of Egg Quality Traits in Brown-Egg Dwarf Layers. Poultry Science 84:1209–1213.

[L160] Zhang, Z; Zhu, L.; Sandler, J.; Friedenberg, S.S.; Egelhoff, J.; Williams, A.M.; Dykes, N.L.; Hornbuckle, W.; Krotscheck, U.; Moise, N.S.; Lust, G.; Todhunter, R.J. 2009. Estimation of heritabilities, genetic correlations, and breeding values of four traits that collectively define hip dysplasia in dogs. AJVR 70(4):483-492.

[L161] Zhao, H., Zeng, C., Wan, S. 2016. Estimates of Heritabilities and Genetic Correlations for Growth and Gonad Traits in Blunt Snout Bream, *Megalobrama amblycephala*. J. World Aquaculture Soc. 47(1):139-146. doi: 10.1111/jwas.12246.

### Appendix 2. Data Validation

Criteria for data validation

In order to ensure the quality of the data that are input, validation rules must be used beforehand analyses are carried out to check for fitness, accuracy, and consistency of the data. The validation logic included simple range and constraint validation (range of parameter estimates within their domains), structured validation (involves operations to check the data for consistency to mathematical relationships).

<u>Domain condition</u>

This range criteria establish that

heritabilities $h^2 \in [0\,,+1\,]$

correlations $r_{\blacksquare x,y} \in [-1\,,+1\,]$

coheritability $h_{x,y} \in [-1\,,+1\,]$

coenvironmentability $e_{x,y} \in [-1\,,+1\,]$

It, therefore, checks that the values of the parameter estimators are numerically within their domains.

<u>Boundary Condition</u>

The criterion states that

$$|h_{x,y}| + |e_{x,y}| \leq 1$$

It verifies that the calculated coheritabilities and coenvironmentabilities are located within the boundaries of the field due to their satisfying all the following conditions: $r_{P_{x,y}} = h_{x,y} + e_{x,y}$, and $-1 < r_{P_{x,y}} < 1$, and $-1 < h_{x,y} < 1$, and $-1 < e_{x,y} < 1$.

Disparity Condition

This structured validation criterion involves the disparity $D = \left| r_{P_{x,y}} - r_{A_{x,y}} \right|$ between the phenotypic and genetic correlations.

$$\left| r_{P_{x,y}} \right| + D \leq 1$$

This procedure checks that the phenotypic and genetic correlations maintain a congruent relationship as expressed in the equation $r_{P_{x,y}} = \sqrt{h_x^2\, h_y^2}\; r_{A_{x,y}} + \sqrt{(1 - h_x^2)(1 - h_y^2)}\; r_{E_{x,y}}$.

Limit Check

The limit criteria are

$$|h_{x,y}| \leq \sqrt{h_x^2\, h_y^2}$$

$$|e_{x,y}| \leq \sqrt{(1 - h_x^2)(1 - h_y^2)}$$

To check that the coheritability and coenvironmentability values are always below or equal to what would be expected were the genetic or environmental correlation be equal to 1, respectively. That is, $|h_{x,y}| \leq \sqrt{h_x^2\, h_y^2}$ means that the factor $\sqrt{h_x^2\, h_y^2}$ which takes values between zero and positive 1, is multiplied by another variable $\left| r_{A_{x,y}} \right|$ which also varies in the same range, the result necessarily will be a number smaller than the factor itself.

### Flow chart of data management

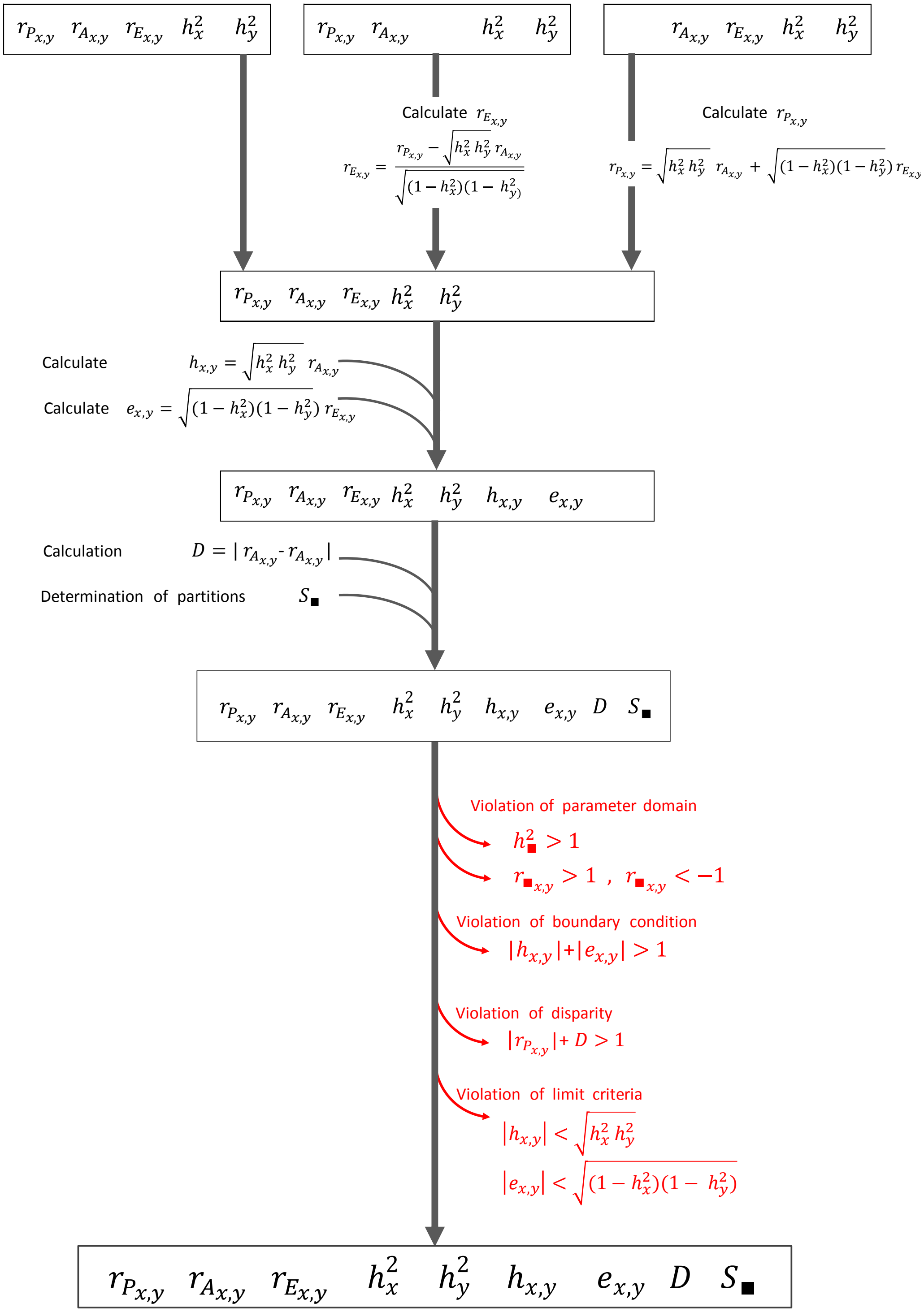

### Appendix 3. Statistical Analyses

#### Part A. Basic Statistics

**The UNIVARIATE Procedure**
**Variable: rP**

| Moments | | | |
|---|---|---|---|
| N | 6286 | Sum Weights | 6286 |
| Mean | 0.15800175 | Sum Observations | 993.199 |
| Std Deviation | 0.30906791 | Variance | 0.09552297 |
| Skewness | 0.4063202 | Kurtosis | 0.94320413 |
| Uncorrected SS | 757.289053 | Corrected SS | 600.361873 |
| Coeff Variation | 195.610433 | Std Error Mean | 0.00389822 |

| Basic Statistical Measures | | | |
|---|---|---|---|
| Location | | Variability | |
| Mean | 0.158002 | Std Deviation | 0.30907 |
| Median | 0.108000 | Variance | 0.09552 |
| Mode | 0.020000 | Range | 1.95200 |
| | | Interquartile Range | 0.32000 |

| Tests for Location: Mu0=0 | | | | |
|---|---|---|---|---|
| Test | Statistic | | p Value | |
| Student's t | t | 40.53173 | Pr > \|t\| | <.0001 |
| Sign | M | 1385 | Pr >= \|M\| | <.0001 |
| Signed Rank | S | 5559832 | Pr >= \|S\| | <.0001 |

| Extreme Observations | | | |
|---|---|---|---|
| Lowest | | Highest | |
| Value | Obs | Value | Obs |
| -0.96 | 2240 | 0.990 | 2064 |
| -0.95 | 4639 | 0.990 | 2067 |
| -0.95 | 2408 | 0.990 | 2212 |
| -0.95 | 1355 | 0.990 | 2511 |

#### The UNIVARIATE Procedure
#### Variable: rA

| Moments | | | |
|---|---|---|---|
| N | 6286 | Sum Weights | 6286 |
| Mean | 0.19502733 | Sum Observations | 1225.9418 |
| Std Deviation | 0.42936089 | Variance | 0.18435078 |
| Skewness | -0.0800828 | Kurtosis | -0.332983 |
| Uncorrected SS | 1397.7368 | Corrected SS | 1158.64464 |
| Coeff Variation | 220.154218 | Std Error Mean | 0.00541546 |

| Basic Statistical Measures | | | |
|---|---|---|---|
| Location | | Variability | |
| Mean | 0.195027 | Std Deviation | 0.42936 |
| Median | 0.160500 | Variance | 0.18435 |
| Mode | 0.010000 | Range | 2.00000 |
| | | Interquartile Range | 0.57000 |

**Note: The mode displayed is the smallest of 2 modes with a count of 69.**

| Tests for Location: Mu0=0 | | | | |
|---|---|---|---|---|
| Test | Statistic | | p Value | |
| Student's t | t | 36.01307 | Pr > \|t\| | <.0001 |
| Sign | M | 1184.5 | Pr >= \|M\| | <.0001 |
| Signed Rank | S | 4744694 | Pr >= \|S\| | <.0001 |

| Extreme Observations | | | |
|---|---|---|---|
| Lowest | | Highest | |
| Value | Obs | Value | Obs |
| -1 | 5222 | 1 | 4384 |
| -1 | 2754 | 1 | 4398 |
| -1 | 2753 | 1 | 5931 |
| -1 | 2750 | 1 | 6008 |

| Extreme Observations | | | |
|---|---|---|---|
| Lowest | | Highest | |
| Value | Obs | Value | Obs |
| -1 | 2748 | 1 | 6121 |

#### The UNIVARIATE Procedure

##### Variable: rE

| Moments | | | |
|---|---|---|---|
| N | 6286 | Sum Weights | 6286 |
| Mean | 0.14026039 | Sum Observations | 881.676821 |
| Std Deviation | 0.31501491 | Variance | 0.09923439 |
| Skewness | 0.31095032 | Kurtosis | 0.97735159 |
| Uncorrected SS | 747.352486 | Corrected SS | 623.68815 |
| Coeff Variation | 224.592918 | Std Error Mean | 0.00397323 |

| Basic Statistical Measures | | | |
|---|---|---|---|
| Location | | Variability | |
| Mean | 0.140260 | Std Deviation | 0.31501 |
| Median | 0.091000 | Variance | 0.09923 |
| Mode | 0.060000 | Range | 2.00000 |
| | | Interquartile Range | 0.32100 |

| Tests for Location: Mu0=0 | | | | |
|---|---|---|---|---|
| Test | Statistic | | p Value | |
| Student's t | t | 35.30134 | Pr > \|t\| | <.0001 |
| Sign | M | 1245 | Pr >= \|M\| | <.0001 |
| Signed Rank | S | 4947997 | Pr >= \|S\| | <.0001 |

| Extreme Observations | | | |
|---|---|---|---|
| Lowest | | Highest | |
| Value | Obs | Value | Obs |

| Extreme Observations | | | |
|---|---|---|---|
| Lowest | | Highest | |
| Value | Obs | Value | Obs |
| -1.000 | 2406 | 0.993 | 2402 |
| -0.982 | 2408 | 0.993 | 2403 |
| -0.981 | 6267 | 0.995 | 2212 |
| -0.980 | 2419 | 1.000 | 1838 |
| -0.971 | 6273 | 1.000 | 2445 |

**The UNIVARIATE Procedure**
**Variable: hxy**

| Moments | | | |
|---|---|---|---|
| N | 6286 | Sum Weights | 6286 |
| Mean | 0.07545434 | Sum Observations | 474.306 |
| Std Deviation | 0.17795416 | Variance | 0.03166768 |
| Skewness | 0.72275883 | Kurtosis | 2.61925239 |
| Uncorrected SS | 234.819846 | Corrected SS | 199.031398 |
| Coeff Variation | 235.843502 | Std Error Mean | 0.00224451 |

| Basic Statistical Measures | | | |
|---|---|---|---|
| Location | | Variability | |
| Mean | 0.075454 | Std Deviation | 0.17795 |
| Median | 0.049000 | Variance | 0.03167 |
| Mode | 0.000000 | Range | 1.71200 |
| | | Interquartile Range | 0.17100 |

| Tests for Location: Mu0=0 | | | | |
|---|---|---|---|---|
| Test | Statistic | | p Value | |
| Student's t | t | 33.61733 | Pr > \|t\| | <.0001 |
| Sign | M | 1190.5 | Pr >= \|M\| | <.0001 |
| Signed Rank | S | 4740642 | Pr >= \|S\| | <.0001 |

| Quantiles (Definition 5) | |
|---|---|
| Level | Quantile |

| Quantiles (Definition 5) | |
|---|---|
| Level | Quantile |
| 100% Max | 0.945 |
| 90% | 0.306 |
| 75% Q3 | 0.155 |
| 50% Median | 0.049 |
| 25% Q1 | -0.016 |
| 10% | -0.105 |
| 5% | -0.172 |
| 1% | -0.357 |
| 0% Min | -0.767 |

The UNIVARIATE Procedure
Variable: exy

| Moments | | | |
|---|---|---|---|
| N | 6286 | Sum Weights | 6286 |
| Mean | 0.0825867 | Sum Observations | 519.14 |
| Std Deviation | 0.19845122 | Variance | 0.03938289 |
| Skewness | 0.59411701 | Kurtosis | 2.2046199 |
| Uncorrected SS | 290.395492 | Corrected SS | 247.521432 |
| Coeff Variation | 240.294399 | Std Error Mean | 0.00250303 |

| Basic Statistical Measures | | | |
|---|---|---|---|
| Location | | Variability | |
| Mean | 0.082587 | Std Deviation | 0.19845 |
| Median | 0.053000 | Variance | 0.03938 |
| Mode | 0.000000 | Range | 1.74000 |
| | | Interquartile Range | 0.17900 |

| Tests for Location: Mu0=0 | | | | |
|---|---|---|---|---|
| Test | Statistic | | p Value | |
| Student's t | t | 32.99465 | Pr > \|t\| | <.0001 |
| Sign | M | 1237 | Pr >= \|M\| | <.0001 |
| Signed Rank | S | 4838707 | Pr >= \|S\| | <.0001 |

| Quantiles (Definition 5) | |
|---|---|
| Level | Quantile |
| 100% Max | 0.876 |
| 99% | 0.669 |
| 75% Q3 | 0.161 |
| 50% Median | 0.053 |
| 25% Q1 | -0.018 |
| 10% | -0.107 |
| 5% | -0.195 |
| 1% | -0.435 |
| 0% Min | -0.864 |

#### Part B. Multinomial Test

The FREQ Procedure

**Statistics for Table of partition**

**Chi-Square Test**
**for Specified Proportions**

| | |
|---|---|
| **Chi-Square** | 9.3370 |
| **DF** | 5 |
| **Pr > ChiSq** | 0.0964 |

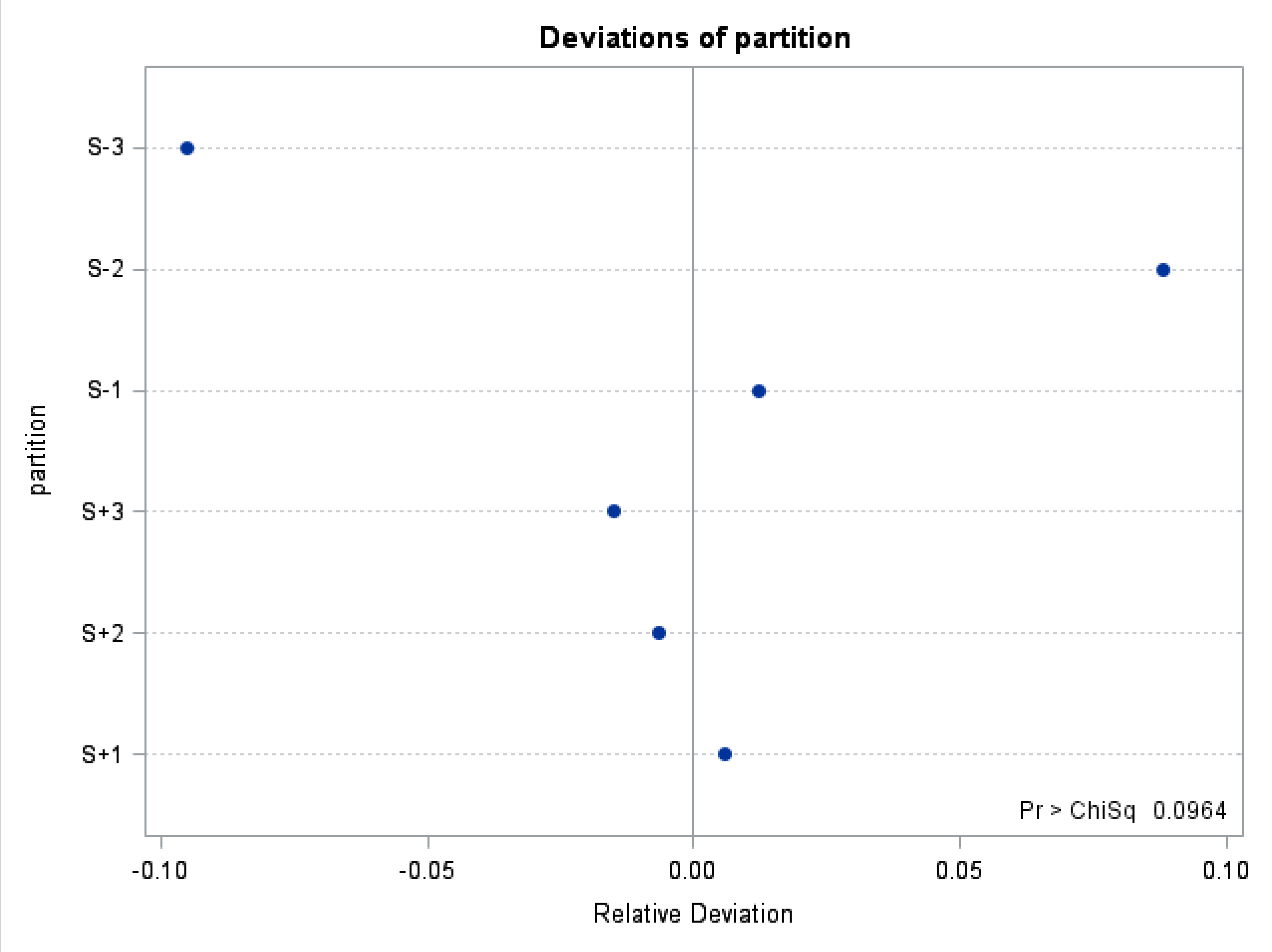

#### Part C. Model 1: Multiple Regression Analysis

**rP=hxy exy -ALL**

**The REG Procedure**
**Model: MODEL1**
**Dependent Variable: rP**

| | |
|---|---|
| Number of Observations Read | 6290 |
| Number of Observations Used | 6290 |

**Analysis of Variance**

| Source | DF | Sum of Squares | Mean Square | F Value | Pr > F |
|---|---|---|---|---|---|
| Model | 2 | 604.38562 | 302.19281 | 2.498E7 | <.0001 |
| Error | 6287 | 0.07605 | 0.00001210 | | |
| Corrected Total | 6289 | 604.46167 | | | |

| | | | |
|---|---|---|---|
| Root MSE | 0.00348 | R-Square | 0.9999 |
| Dependent Mean | 0.15790 | Adj R-Sq | 0.9999 |
| Coeff Var | 2.20262 | | |

**Parameter Estimates**

| Variable | DF | Parameter Estimate | Standard Error | t Value | Pr > \|t\| |
|---|---|---|---|---|---|
| Intercept | 1 | -0.00003174 | 0.00004923 | -0.64 | 0.5191 |
| hxy | 1 | 1.00008 | 0.00026117 | 3829.29 | <.0001 |
| exy | 1 | 0.99983 | 0.00023442 | 4265.11 | <.0001 |

**Collinearity Diagnostics**

| Number | Eigenvalue | Condition Index | Proportion of Variation | | |
|---|---|---|---|---|---|
| | | | Intercept | hxy | exy |
| 1 | 1.80892 | 1.00000 | 0.13712 | 0.14292 | 0.14240 |
| 2 | 0.63233 | 1.69137 | 0.86067 | 0.15532 | 0.21786 |

| Collinearity Diagnostics | | | | | |
|---|---|---|---|---|---|
| | | | Proportion of Variation | | |
| Number | Eigenvalue | Condition Index | Intercept | hxy | exy |
| 3 | 0.55875 | 1.79929 | 0.00222 | 0.70176 | 0.63974 |

**The REG Procedure**
**Model: MODEL1**
**Dependent Variable: rP**
**partition=S+1**

| | |
|---|---|
| Number of Observations Read | 3311 |
| Number of Observations Used | 3311 |

| Analysis of Variance | | | | | |
|---|---|---|---|---|---|
| Source | DF | Sum of Squares | Mean Square | F Value | Pr > F |
| Model | 2 | 227.80716 | 113.90358 | 1.439E9 | <.0001 |
| Error | 3308 | 0.00026185 | 7.915681E-8 | | |
| Corrected Total | 3310 | 227.80742 | | | |

| | | | |
|---|---|---|---|
| Root MSE | 0.00028135 | R-Square | 1.0000 |
| Dependent Mean | 0.34818 | Adj R-Sq | 1.0000 |
| Coeff Var | 0.08081 | | |

| Parameter Estimates | | | | | |
|---|---|---|---|---|---|
| Variable | DF | Parameter Estimate | Standard Error | t Value | Pr > \|t\| |
| Intercept | 1 | 0.00000626 | 0.00000814 | 0.77 | 0.4420 |
| hxy | 1 | 0.99997 | 0.00003262 | 30658.4 | <.0001 |
| exy | 1 | 1.00002 | 0.00002787 | 35877.0 | <.0001 |

#### The REG Procedure
#### Model: MODEL1
#### Dependent Variable: rP
#### partition=S+2

| | |
|---|---|
| Number of Observations Read | 617 |
| Number of Observations Used | 617 |

| Analysis of Variance | | | | | |
|---|---|---|---|---|---|
| Source | DF | Sum of Squares | Mean Square | F Value | Pr > F |
| Model | 2 | 9.75769 | 4.87885 | 6.839E7 | <.0001 |
| Error | 614 | 0.00004380 | 7.133518E-8 | | |
| Corrected Total | 616 | 9.75773 | | | |

| | | | |
|---|---|---|---|
| Root MSE | 0.00026709 | R-Square | 1.0000 |
| Dependent Mean | 0.11935 | Adj R-Sq | 1.0000 |
| Coeff Var | 0.22378 | | |

| Parameter Estimates | | | | | |
|---|---|---|---|---|---|
| Variable | DF | Parameter Estimate | Standard Error | t Value | Pr > \|t\| |
| Intercept | 1 | 0.00000866 | 0.00001802 | 0.48 | 0.6310 |
| hxy | 1 | 1.00000 | 0.00008550 | 11695.6 | <.0001 |
| exy | 1 | 0.99987 | 0.00020235 | 4941.33 | <.0001 |

#### The REG Procedure

**Model: MODEL1**
**Dependent Variable: rP**

**partition=S+3**

| | |
|---|---|
| Number of Observations Read | 367 |
| Number of Observations Used | 367 |

| Analysis of Variance | | | | | |
|---|---|---|---|---|---|
| Source | DF | Sum of Squares | Mean Square | F Value | Pr > F |
| Model | 2 | 2.51740 | 1.25870 | 2.451E7 | <.0001 |
| Error | 364 | 0.00001869 | 5.135616E-8 | | |
| Corrected Total | 366 | 2.51742 | | | |

| | | | |
|---|---|---|---|
| Root MSE | 0.00022662 | R-Square | 1.0000 |
| Dependent Mean | -0.07991 | Adj R-Sq | 1.0000 |
| Coeff Var | -0.28358 | | |

| Parameter Estimates | | | | | |
|---|---|---|---|---|---|
| Variable | DF | Parameter Estimate | Standard Error | t Value | Pr > \|t\| |
| Intercept | 1 | -0.00004385 | 0.00001926 | -2.28 | 0.0234 |
| hxy | 1 | 1.00029 | 0.00026374 | 3792.79 | <.0001 |
| exy | 1 | 0.99986 | 0.00014283 | 7000.37 | <.0001 |

The REG Procedure
Model: MODEL1
Dependent Variable: rP
partition=S-1

| | |
|---|---|
| Number of Observations Read | 880 |
| Number of Observations Used | 880 |

| Analysis of Variance | | | | | |
|---|---|---|---|---|---|
| Source | DF | Sum of Squares | Mean Square | F Value | Pr > F |
| Model | 2 | 37.41069 | 18.70535 | 4.02E8 | <.0001 |
| Error | 877 | 0.00004080 | 4.652576E-8 | | |
| Corrected Total | 879 | 37.41073 | | | |

| | | | |
|---|---|---|---|
| Root MSE | 0.00021570 | R-Square | 1.0000 |
| Dependent Mean | -0.24035 | Adj R-Sq | 1.0000 |
| Coeff Var | -0.08974 | | |

| Parameter Estimates | | | | | |
|---|---|---|---|---|---|
| Variable | DF | Parameter Estimate | Standard Error | t Value | Pr > \|t\| |
| Intercept | 1 | 0.00002168 | 0.00001123 | 1.93 | 0.0539 |
| hxy | 1 | 1.00011 | 0.00006963 | 14362.4 | <.0001 |
| exy | 1 | 1.00002 | 0.00005087 | 19656.8 | <.0001 |

The REG Procedure
Model: MODEL1
Dependent Variable: rP
partition=S-2

| | |
|---|---|
| Number of Observations Read | 473 |
| Number of Observations Used | 473 |

| Analysis of Variance | | | | | |
|---|---|---|---|---|---|
| Source | DF | Sum of Squares | Mean Square | F Value | Pr > F |
| Model | 2 | 4.74065 | 2.37033 | 3.301E7 | <.0001 |
| Error | 470 | 0.00003375 | 7.181444E-8 | | |
| Corrected Total | 472 | 4.74069 | | | |

| | | | |
|---|---|---|---|
| Root MSE | 0.00026798 | R-Square | 1.0000 |
| Dependent Mean | -0.08663 | Adj R-Sq | 1.0000 |
| Coeff Var | -0.30934 | | |

| Parameter Estimates | | | | | |
|---|---|---|---|---|---|
| Variable | DF | Parameter Estimate | Standard Error | t Value | Pr > \|t\| |
| Intercept | 1 | -0.00003170 | 0.00002128 | -1.49 | 0.1370 |
| hxy | 1 | 1.00001 | 0.00012337 | 8105.59 | <.0001 |
| exy | 1 | 1.00037 | 0.00025993 | 3848.65 | <.0001 |

**The REG Procedure**
**Model: MODEL1**
**Dependent Variable: rP**
**partition=S-3**

| | |
|---|---|
| Number of Observations Read | 562 |
| Number of Observations Used | 562 |

| Analysis of Variance | | | | | |
|---|---|---|---|---|---|
| Source | DF | Sum of Squares | Mean Square | F Value | Pr > F |
| Model | 2 | 3.74976 | 1.87488 | 2.68E7 | <.0001 |
| Error | 559 | 0.00003911 | 6.995705E-8 | | |
| Corrected Total | 561 | 3.74980 | | | |

| | | | |
|---|---|---|---|
| Root MSE | 0.00026449 | R-Square | 1.0000 |
| Dependent Mean | 0.08616 | Adj R-Sq | 1.0000 |
| Coeff Var | 0.30697 | | |

| Parameter Estimates | | | | | |
|---|---|---|---|---|---|
| Variable | DF | Parameter Estimate | Standard Error | t Value | Pr > \|t\| |
| Intercept | 1 | 0.00006554 | 0.00002007 | 3.26 | 0.0012 |
| hxy | 1 | 0.99986 | 0.00026281 | 3804.57 | <.0001 |
| exy | 1 | 0.99957 | 0.00013691 | 7301.15 | <.0001 |

#### Part D.  Model 2: Multiple Regression Analysis

##### rP=rA rE - ALL

**The REG Procedure**
**Model: MODEL1**
**Dependent Variable: rP**

| | |
|---|---|
| Number of Observations Read | 6290 |
| Number of Observations Used | 6290 |

| Analysis of Variance | | | | | |
|---|---|---|---|---|---|
| Source | DF | Sum of Squares | Mean Square | F Value | Pr > F |
| Model | 2 | 559.23457 | 279.61728 | 38869.5 | <.0001 |
| Error | 6287 | 45.22710 | 0.00719 | | |
| Corrected Total | 6289 | 604.46167 | | | |

| | | | |
|---|---|---|---|
| Root MSE | 0.08482 | R-Square | 0.9252 |
| Dependent Mean | 0.15790 | Adj R-Sq | 0.9252 |
| Coeff Var | 53.71456 | | |

| Parameter Estimates | | | | | |
|---|---|---|---|---|---|
| Variable | DF | Parameter Estimate | Standard Error | t Value | Pr > \|t\| |
| Intercept | 1 | 0.00264 | 0.00121 | 2.19 | 0.0289 |
| rA | 1 | 0.39036 | 0.00285 | 137.05 | <.0001 |
| rE | 1 | 0.56490 | 0.00388 | 145.62 | <.0001 |

#### rP=rA rE -PARTITION

The REG Procedure
Model: MODEL1
Dependent Variable: rP
partition=S+1

| | |
|---|---|
| Number of Observations Read | 3311 |
| Number of Observations Used | 3311 |

| Analysis of Variance | | | | | |
|---|---|---|---|---|---|
| Source | DF | Sum of Squares | Mean Square | F Value | Pr > F |
| Model | 2 | 214.72230 | 107.36115 | 27141.6 | <.0001 |
| Error | 3308 | 13.08512 | 0.00396 | | |
| Corrected Total | 3310 | 227.80742 | | | |

| | | | |
|---|---|---|---|
| Root MSE | 0.06289 | R-Square | 0.9426 |
| Dependent Mean | 0.34818 | Adj R-Sq | 0.9425 |
| Coeff Var | 18.06363 | | |

| Parameter Estimates | | | | | |
|---|---|---|---|---|---|
| Variable | DF | Parameter Estimate | Standard Error | t Value | Pr > \|t\| |
| Intercept | 1 | -0.01006 | 0.00194 | -5.19 | <.0001 |
| rA | 1 | 0.39393 | 0.00448 | 87.93 | <.0001 |
| rE | 1 | 0.59921 | 0.00511 | 117.36 | <.0001 |

### rP=rA rE -PARTITION

The REG Procedure
Model: MODEL1
Dependent Variable: rP
partition=S+2

| | |
|---|---|
| Number of Observations Read | 617 |
| Number of Observations Used | 617 |

| Analysis of Variance | | | | | |
|---|---|---|---|---|---|
| Source | DF | Sum of Squares | Mean Square | F Value | Pr > F |
| Model | 2 | 2.92511 | 1.46256 | 131.43 | <.0001 |
| Error | 614 | 6.83262 | 0.01113 | | |
| Corrected Total | 616 | 9.75773 | | | |

| | | | |
|---|---|---|---|
| Root MSE | 0.10549 | R-Square | 0.2998 |
| Dependent Mean | 0.11935 | Adj R-Sq | 0.2975 |
| Coeff Var | 88.38666 | | |

| Parameter Estimates | | | | | |
|---|---|---|---|---|---|
| Variable | DF | Parameter Estimate | Standard Error | t Value | Pr > \|t\| |
| Intercept | 1 | 0.00804 | 0.00825 | 0.97 | 0.3302 |
| rA | 1 | 0.25936 | 0.01665 | 15.57 | <.0001 |
| rE | 1 | 0.01197 | 0.02959 | 0.40 | 0.6859 |

#### rP=rA rE -PARTITION

The REG Procedure
Model: MODEL1
Dependent Variable: rP
partition=S+3

| | |
|---|---|
| Number of Observations Read | 367 |
| Number of Observations Used | 367 |

| Analysis of Variance | | | | | |
|---|---|---|---|---|---|
| Source | DF | Sum of Squares | Mean Square | F Value | Pr > F |
| Model | 2 | 0.97356 | 0.48678 | 114.77 | <.0001 |
| Error | 364 | 1.54386 | 0.00424 | | |
| Corrected Total | 366 | 2.51742 | | | |

| | | | |
|---|---|---|---|
| Root MSE | 0.06513 | R-Square | 0.3867 |
| Dependent Mean | -0.07991 | Adj R-Sq | 0.3834 |
| Coeff Var | -81.49611 | | |

| Parameter Estimates | | | | | |
|---|---|---|---|---|---|
| Variable | DF | Parameter Estimate | Standard Error | t Value | Pr > \|t\| |
| Intercept | 1 | -0.03867 | 0.00543 | -7.12 | <.0001 |
| rA | 1 | 0.11735 | 0.02188 | 5.36 | <.0001 |
| rE | 1 | 0.30134 | 0.01989 | 15.15 | <.0001 |

### rP=rA rE -PARTITION

The REG Procedure
Model: MODEL1
Dependent Variable: rP
partition=S-1

| | |
|---|---|
| Number of Observations Read | 880 |
| Number of Observations Used | 880 |

| Analysis of Variance | | | | | |
|---|---|---|---|---|---|
| Source | DF | Sum of Squares | Mean Square | F Value | Pr > F |
| Model | 2 | 33.38106 | 16.69053 | 3632.45 | <.0001 |
| Error | 877 | 4.02967 | 0.00459 | | |
| Corrected Total | 879 | 37.41073 | | | |

| | | | |
|---|---|---|---|
| Root MSE | 0.06779 | R-Square | 0.8923 |
| Dependent Mean | -0.24035 | Adj R-Sq | 0.8920 |
| Coeff Var | -28.20244 | | |

| Parameter Estimates | | | | | |
|---|---|---|---|---|---|
| Variable | DF | Parameter Estimate | Standard Error | t Value | Pr > \|t\| |
| Intercept | 1 | -0.00809 | 0.00367 | -2.20 | 0.0277 |
| rA | 1 | 0.36761 | 0.01029 | 35.73 | <.0001 |
| rE | 1 | 0.60166 | 0.01215 | 49.54 | <.0001 |

### rP=rA rE -PARTITION

The REG Procedure
Model: MODEL1
Dependent Variable: rP
partition=S-2

| | |
|---|---|
| Number of Observations Read | 473 |
| Number of Observations Used | 473 |

| Analysis of Variance | | | | | |
|---|---|---|---|---|---|
| Source | DF | Sum of Squares | Mean Square | F Value | Pr > F |
| Model | 2 | 1.11413 | 0.55706 | 72.20 | <.0001 |
| Error | 470 | 3.62656 | 0.00772 | | |
| Corrected Total | 472 | 4.74069 | | | |

| | | | |
|---|---|---|---|
| Root MSE | 0.08784 | R-Square | 0.2350 |
| Dependent Mean | -0.08663 | Adj R-Sq | 0.2318 |
| Coeff Var | -101.39814 | | |

| Parameter Estimates | | | | | |
|---|---|---|---|---|---|
| Variable | DF | Parameter Estimate | Standard Error | t Value | Pr > \|t\| |
| Intercept | 1 | -0.00466 | 0.00796 | -0.59 | 0.5582 |
| rA | 1 | 0.21788 | 0.01982 | 10.99 | <.0001 |
| rE | 1 | -0.03615 | 0.02960 | -1.22 | 0.2226 |

The REG Procedure
Model: MODEL1
Dependent Variable: rP
partition=S-3

| | |
|---|---|
| Number of Observations Read | 562 |
| Number of Observations Used | 562 |

| Analysis of Variance | | | | | |
|---|---|---|---|---|---|
| Source | DF | Sum of Squares | Mean Square | F Value | Pr > F |
| Model | 2 | 1.86161 | 0.93080 | 275.56 | <.0001 |
| Error | 559 | 1.88820 | 0.00338 | | |
| Corrected Total | 561 | 3.74980 | | | |

| | | | |
|---|---|---|---|
| Root MSE | 0.05812 | R-Square | 0.4965 |
| Dependent Mean | 0.08616 | Adj R-Sq | 0.4947 |
| Coeff Var | 67.45178 | | |

| Parameter Estimates | | | | | |
|---|---|---|---|---|---|
| Variable | DF | Parameter Estimate | Standard Error | t Value | Pr > \|t\| |
| Intercept | 1 | 0.02795 | 0.00447 | 6.25 | <.0001 |
| rA | 1 | 0.09673 | 0.01367 | 7.08 | <.0001 |
| rE | 1 | 0.35169 | 0.01511 | 23.28 | <.0001 |

#### Part E.  Simple regression of phenotypic correlation against a single factor

##### model rP=hxy ALL

**The REG Procedure**
**Model: MODELx1**
**Dependent Variable: rP**

| | |
|---|---|
| Number of Observations Read | 6290 |
| Number of Observations Used | 6290 |

| Analysis of Variance | | | | | |
|---|---|---|---|---|---|
| Source | DF | Sum of Squares | Mean Square | F Value | Pr > F |
| Model | 1 | 384.34137 | 384.34137 | 10979.2 | <.0001 |
| Error | 6288 | 220.12030 | 0.03501 | | |
| Corrected Total | 6289 | 604.46167 | | | |

| | | | |
|---|---|---|---|
| Root MSE | 0.18710 | R-Square | 0.6358 |
| Dependent Mean | 0.15790 | Adj R-Sq | 0.6358 |
| Coeff Var | 118.49176 | | |

| Parameter Estimates | | | | | |
|---|---|---|---|---|---|
| Variable | DF | Parameter Estimate | Standard Error | t Value | Pr > \|t\| |
| Intercept | 1 | 0.05364 | 0.00256 | 20.95 | <.0001 |
| hxy | 1 | 1.38262 | 0.01320 | 104.78 | <.0001 |

#### rP=exy -ALL

**The REG Procedure**
**Model: MODELx1**
**Dependent Variable: rP**

| | |
|---|---|
| Number of Observations Read | 6290 |
| Number of Observations Used | 6290 |

**Analysis of Variance**

| Source | DF | Sum of Squares | Mean Square | F Value | Pr > F |
|---|---|---|---|---|---|
| Model | 1 | 427.01299 | 427.01299 | 15131.5 | <.0001 |
| Error | 6288 | 177.44868 | 0.02822 | | |
| Corrected Total | 6289 | 604.46167 | | | |

| | | | |
|---|---|---|---|
| Root MSE | 0.16799 | R-Square | 0.7064 |
| Dependent Mean | 0.15790 | Adj R-Sq | 0.7064 |
| Coeff Var | 106.38845 | | |

**Parameter Estimates**

| Variable | DF | Parameter Estimate | Standard Error | t Value | Pr > \|t\| |
|---|---|---|---|---|---|
| Intercept | 1 | 0.04994 | 0.00229 | 21.78 | <.0001 |
| exy | 1 | 1.30811 | 0.01063 | 123.01 | <.0001 |

#### rP=rA -ALL

**The REG Procedure**
**Model: MODEL1**
**Dependent Variable: rP**

| | |
|---|---|
| Number of Observations Read | 6290 |
| Number of Observations Used | 6290 |

| Analysis of Variance | | | | | |
|---|---|---|---|---|---|
| Source | DF | Sum of Squares | Mean Square | F Value | Pr > F |
| Model | 1 | 406.69694 | 406.69694 | 12931.1 | <.0001 |
| Error | 6288 | 197.76473 | 0.03145 | | |
| Corrected Total | 6289 | 604.46167 | | | |

| | | | |
|---|---|---|---|
| Root MSE | 0.17734 | R-Square | 0.6728 |
| Dependent Mean | 0.15790 | Adj R-Sq | 0.6728 |
| Coeff Var | 112.31365 | | |

| Parameter Estimates | | | | | |
|---|---|---|---|---|---|
| Variable | DF | Parameter Estimate | Standard Error | t Value | Pr > \|t\| |
| Intercept | 1 | 0.04254 | 0.00246 | 17.32 | <.0001 |
| rA | 1 | 0.59191 | 0.00521 | 113.71 | <.0001 |

#### rP=rE -ALL

**The REG Procedure**
**Model: MODEL1**
**Dependent Variable: rP**

| | |
|---|---|
| Number of Observations Read | 6290 |
| Number of Observations Used | 6290 |

| Analysis of Variance | | | | | |
|---|---|---|---|---|---|
| Source | DF | Sum of Squares | Mean Square | F Value | Pr > F |
| Model | 1 | 424.12059 | 424.12059 | 14787.9 | <.0001 |
| Error | 6288 | 180.34108 | 0.02868 | | |
| Corrected Total | 6289 | 604.46167 | | | |

| | | | |
|---|---|---|---|
| Root MSE | 0.16935 | R-Square | 0.7017 |
| Dependent Mean | 0.15790 | Adj R-Sq | 0.7016 |
| Coeff Var | 107.25201 | | |

| Parameter Estimates | | | | | |
|---|---|---|---|---|---|
| Variable | DF | Parameter Estimate | Standard Error | t Value | Pr > \|t\| |
| Intercept | 1 | 0.04250 | 0.00234 | 18.19 | <.0001 |
| rE | 1 | 0.82326 | 0.00677 | 121.61 | <.0001 |

#### Part F. Simple Regression of rA on rP

##### Phenotypic correlation as predictor of genetic correlation

**The REG Procedure**
**Model: MODEL1**
**Dependent Variable: rA**

| | |
|---|---|
| Number of Observations Read | 6290 |
| Number of Observations Used | 6290 |

| Analysis of Variance | | | | | |
|---|---|---|---|---|---|
| Source | DF | Sum of Squares | Mean Square | F Value | Pr > F |
| Model | 1 | 781.01307 | 781.01307 | 12931.1 | <.0001 |
| Error | 6288 | 379.78362 | 0.06040 | | |
| Corrected Total | 6289 | 1160.79669 | | | |

| | | | |
|---|---|---|---|
| Root MSE | 0.24576 | R-Square | 0.6728 |
| Dependent Mean | 0.19490 | Adj R-Sq | 0.6728 |
| Coeff Var | 126.09348 | | |

| Parameter Estimates | | | | | |
|---|---|---|---|---|---|
| Variable | DF | Parameter Estimate | Standard Error | t Value | Pr > \|t\| |
| Intercept | 1 | 0.01542 | 0.00348 | 4.43 | <.0001 |
| rP | 1 | 1.13670 | 0.01000 | 113.71 | <.0001 |

#### Part G. Test of heterogeneity of regression coefficients

Results of the test that the regression coefficients of Model 1 (involving the coheritability and coenvironmentability as regressors) are not different than zero

**rP=hxy exy -ALL**

**The REG Procedure**
**Model: MODEL1**

| Test test00 Results for Dependent Variable rP | | | | |
|---|---|---|---|---|
| Source | DF | Mean Square | F Value | Pr > F |
| Numerator | 2 | 299.12757 | 4.221E9 | <.0001 |
| Denominator | 6207 | 7.086589E-8 | | |

Results of the test that the regression coefficients of Model 1 (involving the coheritability and coenvironmentability as regressors) are not different than unity

**rP=hxy exy -ALL**

**The REG Procedure**
**Model: MODEL1**

| Test test01 Results for Dependent Variable rP | | | | |
|---|---|---|---|---|
| Source | DF | Mean Square | F Value | Pr > F |
| Numerator | 2 | 4.57815E-9 | 0.06 | 0.9374 |
| Denominator | 6207 | 7.086589E-8 | | |

#### Test of Heterogenetity of slopes by partition model 1

##### The GLM Procedure

##### Dependent Variable: rP

| Source | DF | Sum of Squares | Mean Square | F Value | Pr > F |
|---|---|---|---|---|---|
| Model | 12 | 598.2551448 | 49.8545954 | 7.033E8 | <.0001 |
| Error | 6197 | 0.0004393 | 0.0000001 | | |
| Corrected Total | 6209 | 598.2555841 | | | |

| R-Square | Coeff Var | Root MSE | rP Mean |
|---|---|---|---|
| 0.999999 | 0.166491 | 0.000266 | 0.159914 |

| Source | DF | Type I SS | Mean Square | F Value | Pr > F |
|---|---|---|---|---|---|
| hxy | 1 | 381.6300961 | 381.6300961 | 5.384E9 | <.0001 |
| exy | 1 | 216.6250481 | 216.6250481 | 3.056E9 | <.0001 |
| hxy*partition | 5 | 0.0000002 | 0.0000000 | 0.64 | 0.6681 |
| exy*partition | 5 | 0.0000004 | 0.0000001 | 1.02 | 0.4038 |

| Parameter | Estimate | | Standard Error | t Value | Pr > \|t\| |
|---|---|---|---|---|---|
| Intercept | 0.0000078329 | | 0.00000561 | 1.40 | 0.1625 |
| hxy | 0.9997105089 | B | 0.00025964 | 3850.43 | <.0001 |
| exy | 0.9998122234 | B | 0.00010991 | 9096.53 | <.0001 |
| hxy*partition S+1 | 0.0002540093 | B | 0.00026188 | 0.97 | 0.3321 |
| hxy*partition S+2 | 0.0002920603 | B | 0.00027023 | 1.08 | 0.2798 |
| hxy*partition S+3 | 0.0005000960 | B | 0.00040304 | 1.24 | 0.2147 |
| hxy*partition S-1 | 0.0003553349 | B | 0.00027009 | 1.32 | 0.1884 |
| hxy*partition S-2 | 0.0004238301 | B | 0.00027890 | 1.52 | 0.1287 |
| hxy*partition S-3 | 0.0000000000 | B | . | . | . |
| exy*partition S+1 | 0.0002021176 | B | 0.00011064 | 1.83 | 0.0678 |
| exy*partition S+2 | 0.0000516191 | B | 0.00022539 | 0.23 | 0.8189 |
| exy*partition S+3 | 0.0002783650 | B | 0.00017860 | 1.56 | 0.1191 |
| exy*partition S-1 | 0.0001797528 | B | 0.00012612 | 1.43 | 0.1541 |
| exy*partition S-2 | 0.0004022520 | B | 0.00026639 | 1.51 | 0.1311 |
| exy*partition S-3 | 0.0000000000 | B | . | . | . |

#### Test of Heterogenetity of slopes by partition model 2

##### The GLM Procedure

##### Dependent Variable: rP

| Source | DF | Sum of Squares | Mean Square | F Value | Pr > F |
|---|---|---|---|---|---|
| Model | 12 | 566.8512326 | 47.2376027 | 9321.37 | <.0001 |
| Error | 6197 | 31.4043515 | 0.0050677 | | |
| Corrected Total | 6209 | 598.2555841 | | | |

| R-Square | Coeff Var | Root MSE | rP Mean |
|---|---|---|---|
| 0.947507 | 44.51616 | 0.071188 | 0.159914 |

| Source | DF | Type I SS | Mean Square | F Value | Pr > F |
|---|---|---|---|---|---|
| rA | 1 | 406.6296609 | 406.6296609 | 80240.0 | <.0001 |
| rE | 1 | 148.6597054 | 148.6597054 | 29334.9 | <.0001 |
| rA*partition | 5 | 3.5007162 | 0.7001432 | 138.16 | <.0001 |
| rE*partition | 5 | 8.0611501 | 1.6122300 | 318.14 | <.0001 |

| Parameter | Estimate | | Standard Error | t Value | Pr > \|t\| |
|---|---|---|---|---|---|
| Intercept | -.0067448116 | | 0.00159477 | -4.23 | <.0001 |
| rA | 0.0511668388 | B | 0.01525943 | 3.35 | 0.0008 |
| rE | 0.4257233458 | B | 0.01474941 | 28.86 | <.0001 |
| rA*partition S+1 | 0.3388508491 | B | 0.01622625 | 20.88 | <.0001 |
| rA*partition S+2 | 0.2303500082 | B | 0.01747176 | 13.18 | <.0001 |
| rA*partition S+3 | 0.0170564459 | B | 0.02716135 | 0.63 | 0.5300 |
| rA*partition S-1 | 0.3184734467 | B | 0.01765384 | 18.04 | <.0001 |
| rA*partition S-2 | 0.1628277772 | B | 0.01849852 | 8.80 | <.0001 |
| rA*partition S-3 | 0.0000000000 | B | . | . | . |
| rE*partition S+1 | 0.1717135371 | B | 0.01563650 | 10.98 | <.0001 |
| rE*partition S+2 | -.4235432624 | B | 0.02471607 | -17.14 | <.0001 |
| rE*partition S+3 | -.0673930605 | B | 0.02460769 | -2.74 | 0.0062 |
| rE*partition S-1 | 0.1769540581 | B | 0.01952937 | 9.06 | <.0001 |
| rE*partition S-2 | -.4605367454 | B | 0.02774716 | -16.60 | <.0001 |
| rE*partition S-3 | 0.0000000000 | B | . | . | . |

Part H. Correlation between the squared disparity and sampling variance of the genetic correlation

| Simple Statistics | | | | | | |
|---|---|---|---|---|---|---|
| **Variable** | **N** | **Mean** | **Std Dev** | **Sum** | **Minimum** | **Maximum** |
| **D2** | 3234 | 0.06406 | 0.12350 | 207.16323 | 0 | 0.96040 |
| **V_rA** | 3241 | 0.04516 | 0.15385 | 146.36436 | 0 | 3.79080 |

**Pearson Correlation Coefficients**
**Prob > |r| under H0: Rho=0**
**Number of Observations**

| | **D2** | **V_rA** |
|---|---|---|
| **D2** | 1.00000<br><br>3234 | 0.17093<br><.0001<br>3233 |
| **V_rA** | 0.17093<br><.0001<br>3233 | 1.00000<br><br>3241 |

#### Part I. Wilcoxon Rank Test of the squared disparity and the sampling variance of the genetic correlation

**The NPAR1WAY Procedure**

| Wilcoxon Scores (Rank Sums) for Variable $D^2$ Classified by Variable partition | | | | | |
|---|---|---|---|---|---|
| **partition** | **N** | **Sum of Scores** | **Expected Under H0** | **Std Dev Under H0** | **Mean Score** |
| **S+1** | 1862 | 2539773.00 | 3011785.00 | 26240.6879 | 1364.00269 |
| **S+2** | 251 | 608568.00 | 405992.50 | 14206.0000 | 2424.57371 |
| **S+3** | 171 | 344470.00 | 276592.50 | 11881.7354 | 2014.44444 |
| **S-1** | 454 | 607333.50 | 734345.00 | 18444.1501 | 1337.73899 |
| **S-2** | 192 | 449557.50 | 310560.00 | 12546.9608 | 2341.44531 |
| **S-3** | 304 | 681293.00 | 491720.00 | 15494.5485 | 2241.09539 |
| Average scores were used for ties. | | | | | |

| Kruskal-Wallis Test | |
|---|---|
| **Chi-Square** | 647.5628 |
| **DF** | 5 |
| **Pr > Chi-Square** | <.0001 |

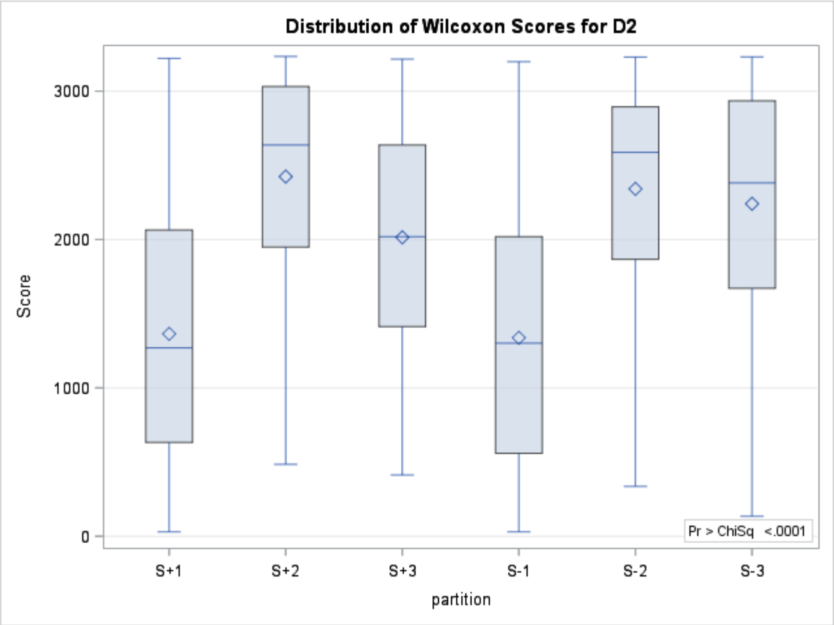

The NPAR1WAY Procedure

| Wilcoxon Scores (Rank Sums) for Variable Var( rA ) Classified by Variable partition | | | | | |
|---|---|---|---|---|---|
| **partition** | **N** | **Sum of Scores** | **Expected Under H0** | **Std Dev Under H0** | **Mean Score** |
| **S+1** | 1865 | 2477918.00 | 3023165.0 | 26324.8233 | 1328.64236 |
| **S+2** | 251 | 507142.50 | 406871.0 | 14236.0428 | 2020.48805 |
| **S+3** | 171 | 349740.00 | 277191.0 | 11906.4988 | 2045.26316 |
| **S-1** | 455 | 796619.50 | 737555.0 | 18501.7617 | 1750.81209 |
| **S-2** | 194 | 436730.00 | 314474.0 | 12634.3796 | 2251.18557 |
| **S-3** | 305 | 685511.00 | 494405.0 | 15550.5149 | 2247.57705 |
| Average scores were used for ties. | | | | | |

| Kruskal-Wallis Test | |
|---|---|
| **Chi-Square** | 496.6782 |
| **DF** | 5 |
| **Pr > Chi-Square** | <.0001 |

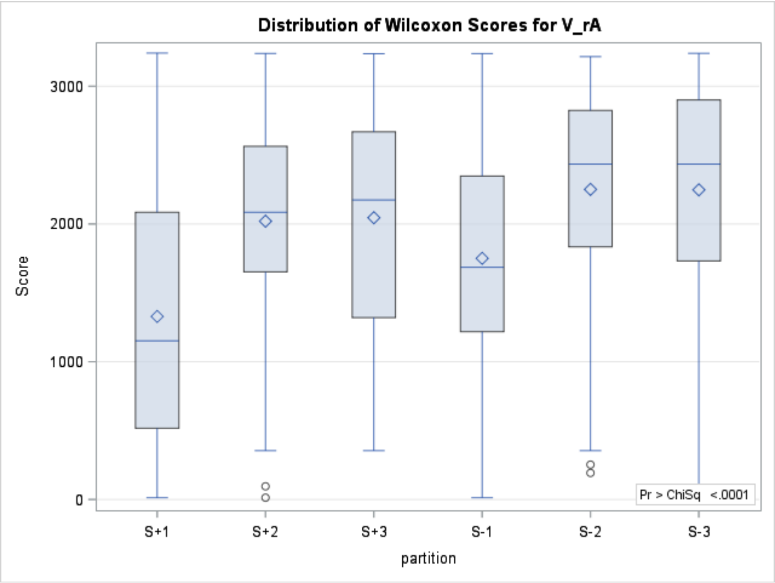

**Results:**

**Wilcoxon signed rank test with continuity correction**

Test statistic V : 3138479

p-value : 2.971110e-31

There are ties in the ranks, so the exact p-value cannot be computed. The p-value shown is inexact but fairly robust when the number of ties is small.

null hypothesis $\mu_0 = 0.0$

alternative hypothesis: two sided, $\hat{\mu} \neq \mu_0$

95% confidence interval : 0.0056 --------- 0.0088

sample estimate of pseudo-median $\hat{\mu}$ : 0.00711298393513

Results from http://astatsa.com/WilcoxonTest/Result/

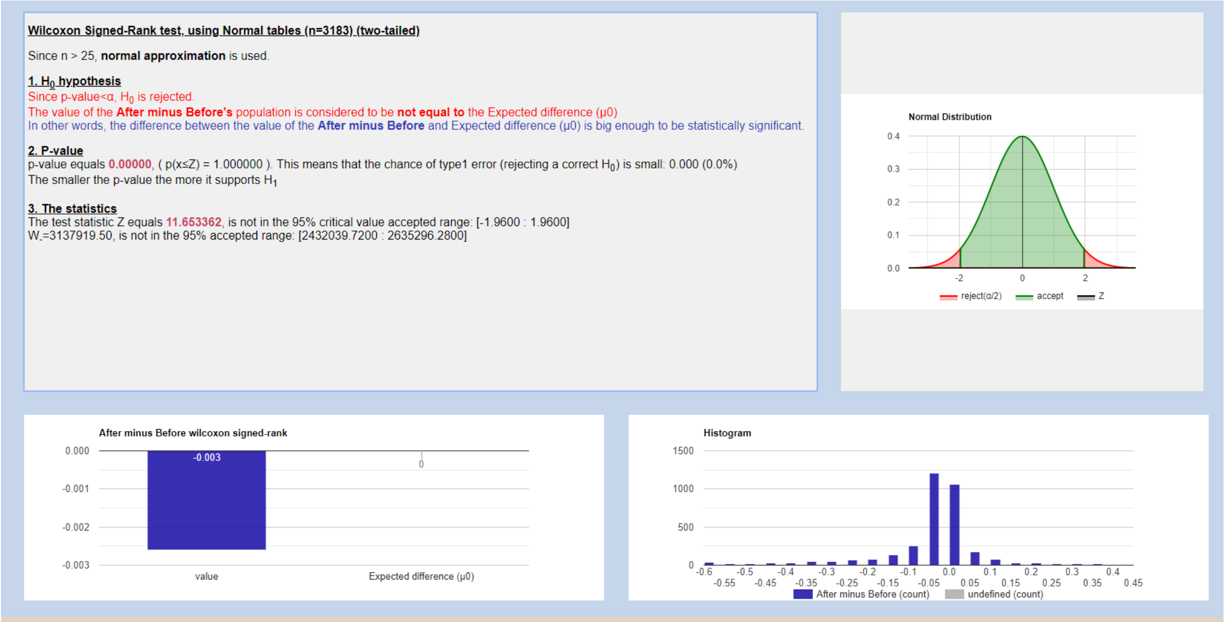

Results from [http://www.statskingdom.com/175wilcoxon_signed_ranks.html](http://www.statskingdom.com/175wilcoxon_signed_ranks.html)

**Figure S1. Abundance, dispersion and distribution of coheritability, coenvironmentability, and phenotypic correlation, overall and by partition. A single datum determined by $\{\ h_{x,y}\ ,e_{x,y}\ ,r_{P_{x,y}}\ \}$ is represented by a dot in the scatter plot.**

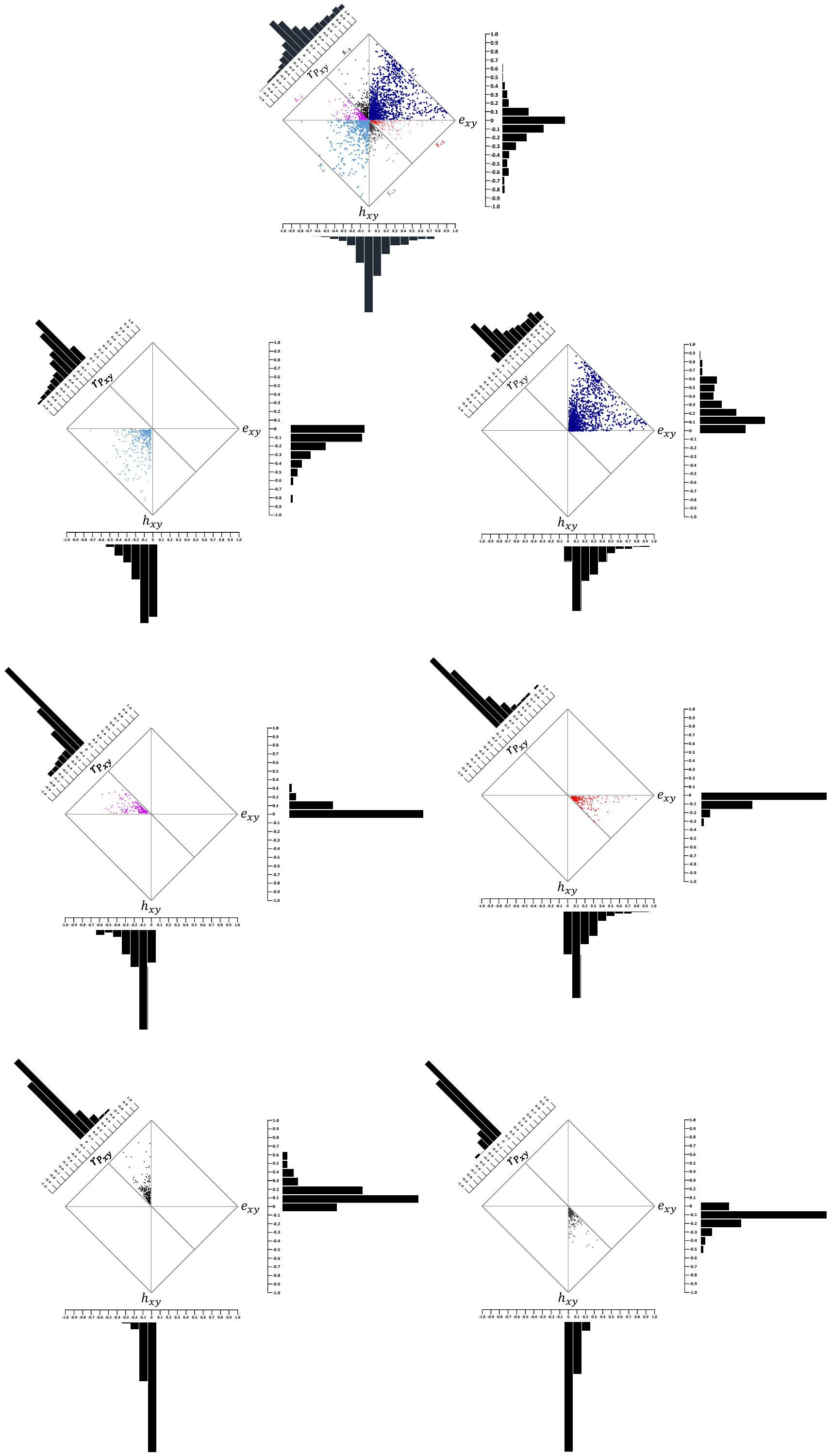

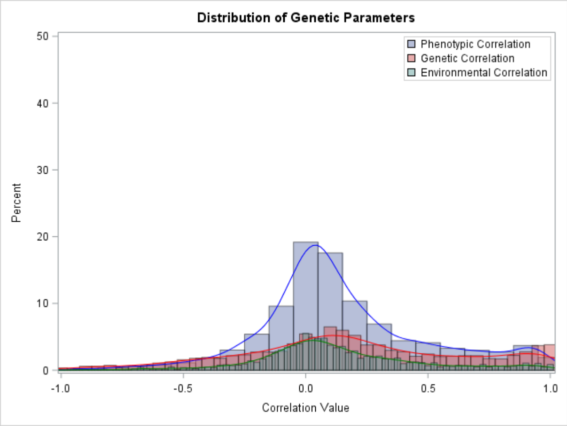
Distribution of Genetic Parameters
Phenotypic Correlation
Genetic Correlation
Environmental Correlation
Percent
Correlation Value

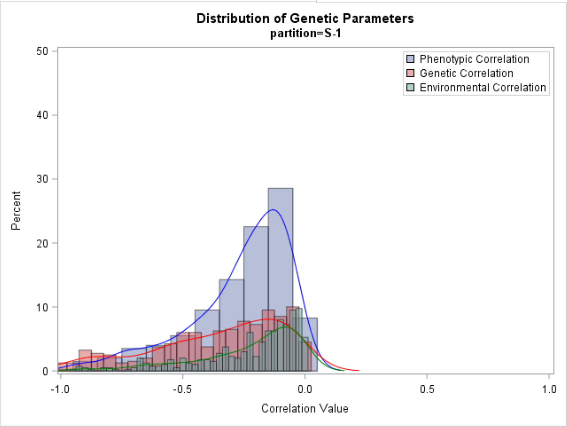
Distribution of Genetic Parameters
partition=S-1
Phenotypic Correlation
Genetic Correlation
Environmental Correlation
Percent
Correlation Value

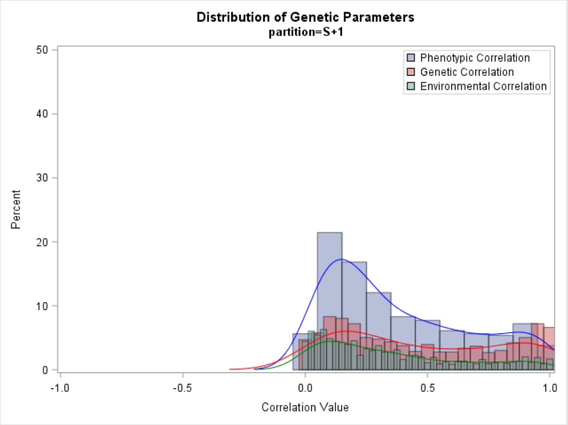
Distribution of Genetic Parameters
partition=S+1
Phenotypic Correlation
Genetic Correlation
Environmental Correlation
Percent
Correlation Value

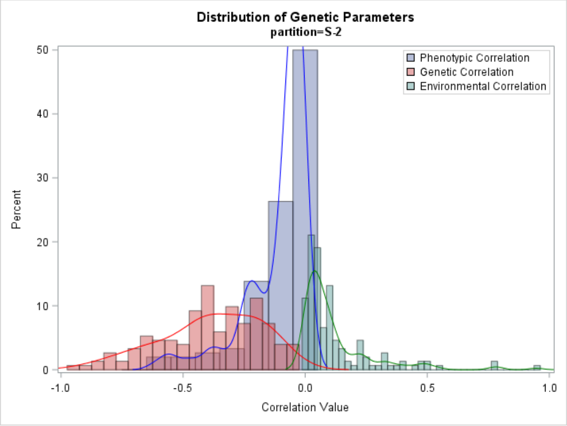
Distribution of Genetic Parameters
partition=S-2
Phenotypic Correlation
Genetic Correlation
Environmental Correlation
Percent
Correlation Value

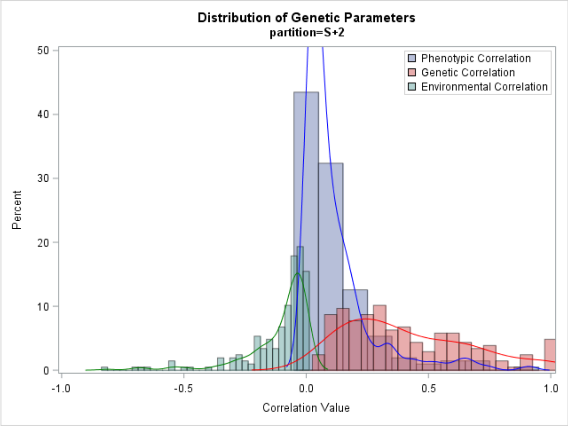
Distribution of Genetic Parameters
partition=S+2
Phenotypic Correlation
Genetic Correlation
Environmental Correlation
Percent
Correlation Value

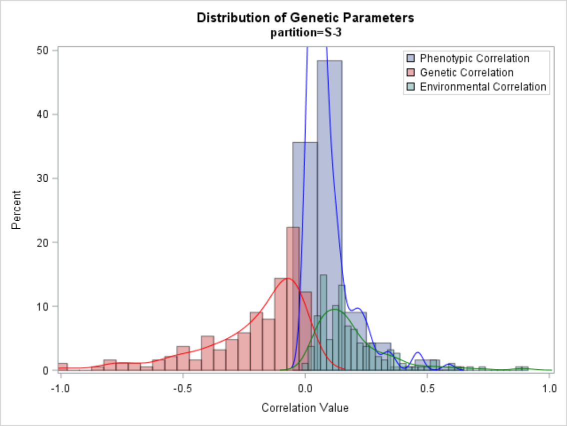
Distribution of Genetic Parameters
partition=S-3
Phenotypic Correlation
Genetic Correlation
Environmental Correlation
Percent
Correlation Value

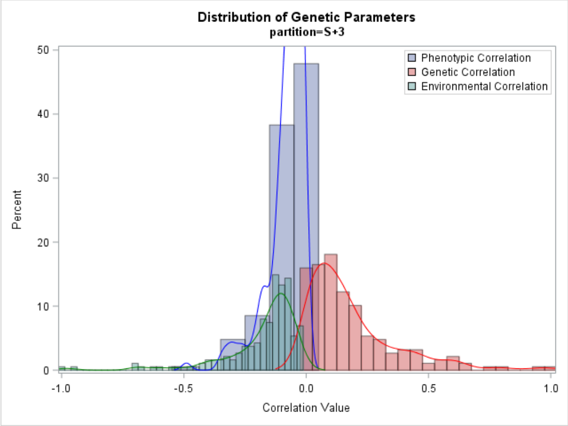
Distribution of Genetic Parameters
partition=S+3
Phenotypic Correlation
Genetic Correlation
Environmental Correlation
Percent
Correlation Value

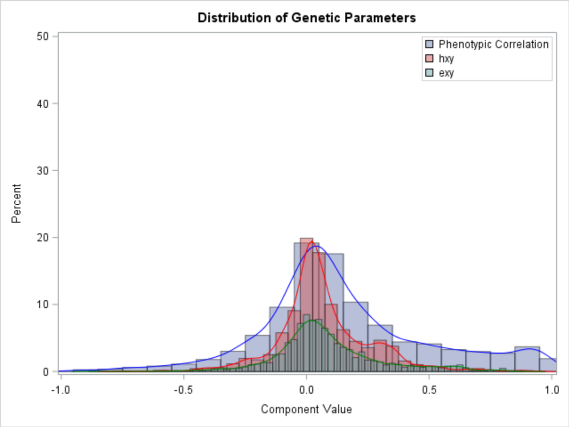
Distribution of Genetic Parameters
Phenotypic Correlation
hxy
exy
Percent
Component Value

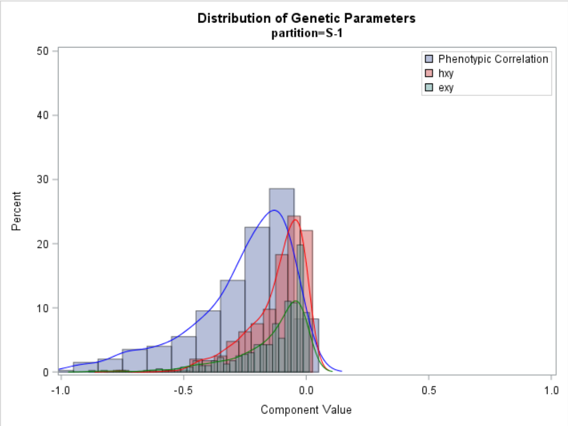
Distribution of Genetic Parameters
partition=S-1
Phenotypic Correlation
hxy
exy
Percent
Component Value

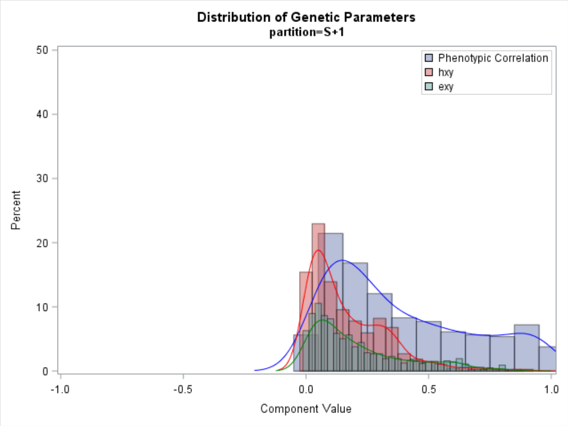
Distribution of Genetic Parameters
partition=S+1
Phenotypic Correlation
hxy
exy
Percent
Component Value

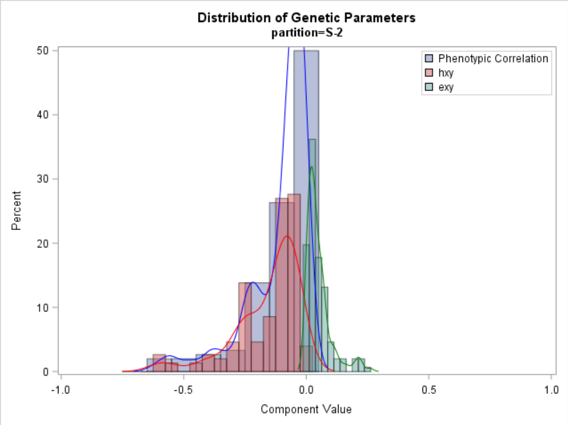
Distribution of Genetic Parameters
partition=S-2
Phenotypic Correlation
hxy
exy
Percent
Component Value

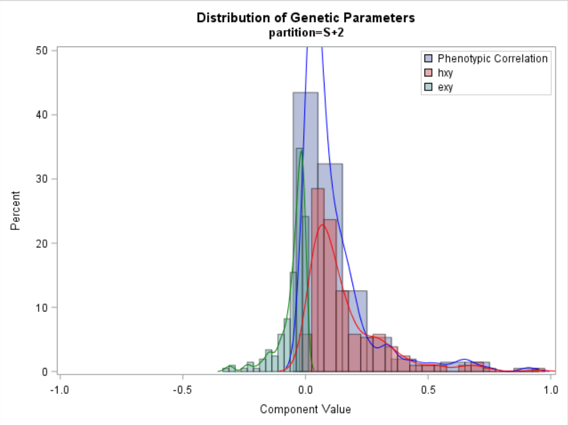
Distribution of Genetic Parameters
partition=S+2
Phenotypic Correlation
hxy
exy
Percent
Component Value

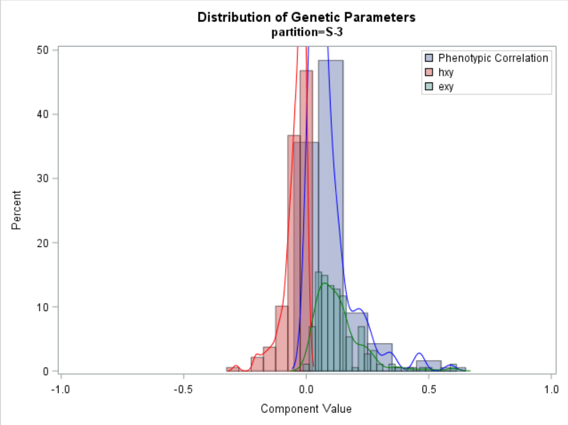
Distribution of Genetic Parameters
partition=S-3
Phenotypic Correlation
hxy
exy
Percent
Component Value

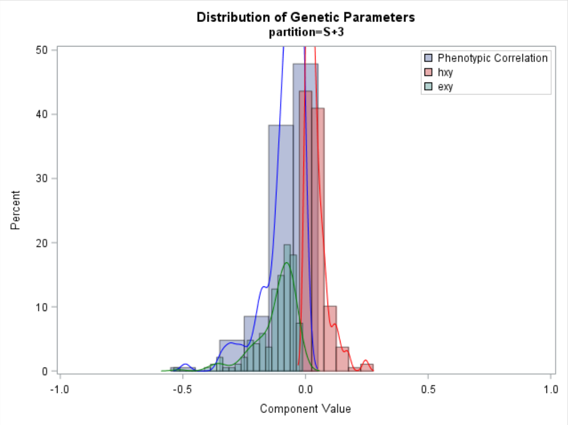
Distribution of Genetic Parameters
partition=S+3
Phenotypic Correlation
hxy
exy
Percent
Component Value

#### Figure S2. Scatter plot and regression of genetic correlation (A) and environmental correlation (B) against the phenotypic correlation. (C) Regression of environmental correlation on genetic correlation.

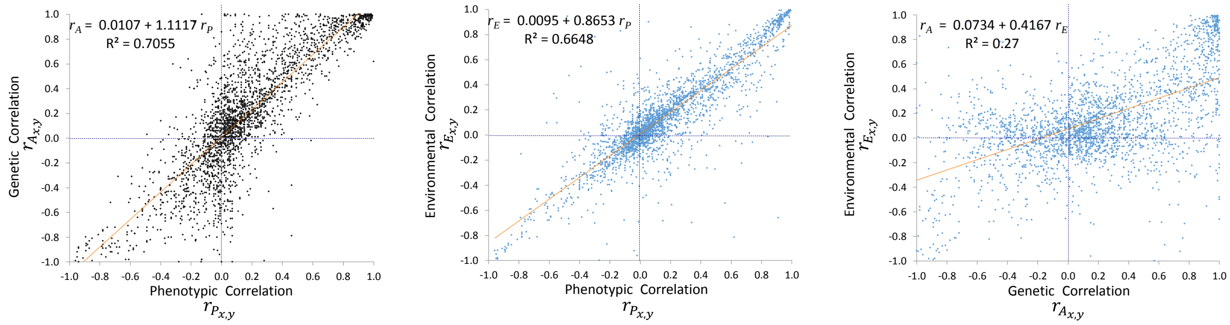

#### Figure S3. Disparity Index by partition

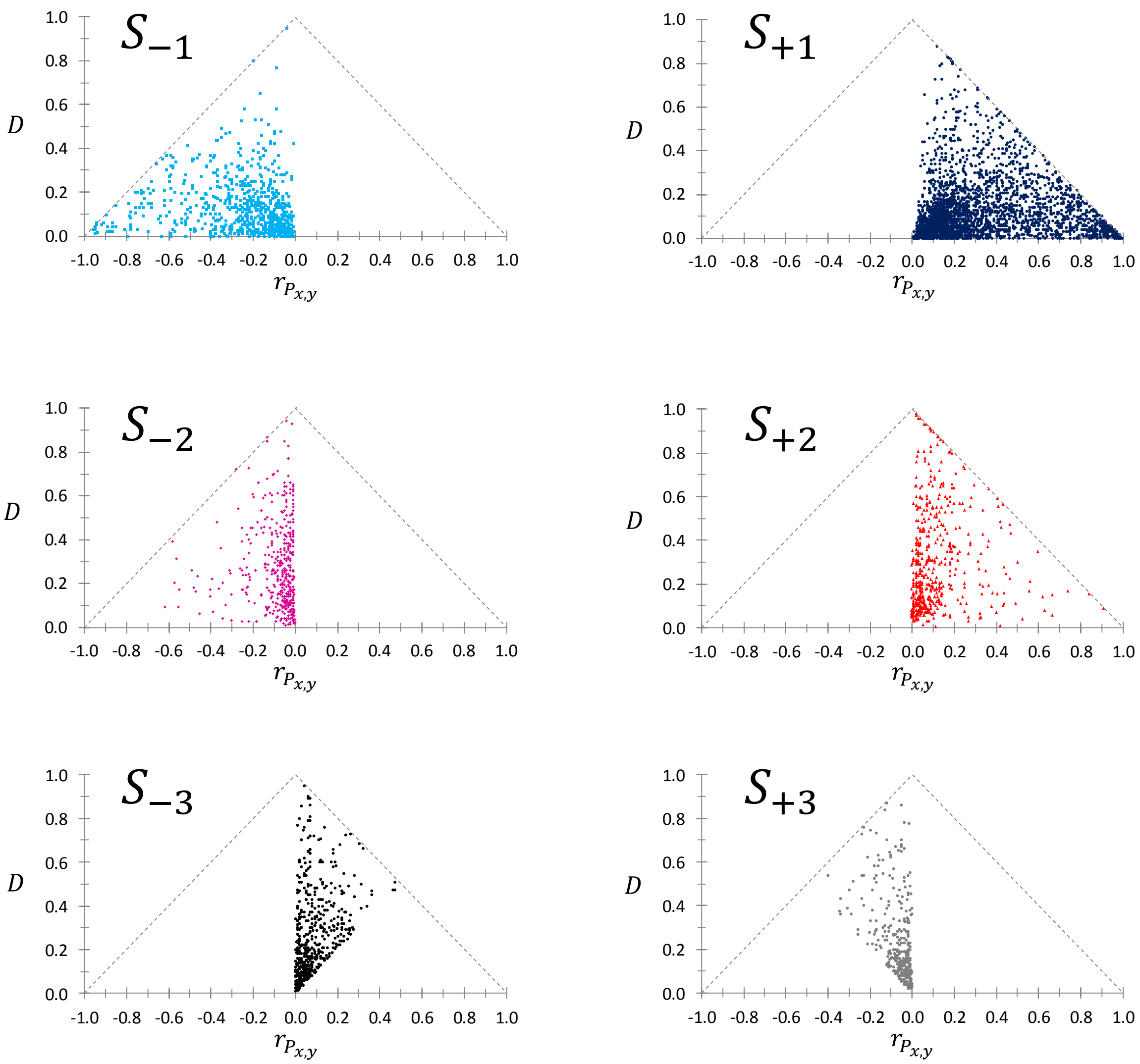

#### Figure S4. Supplementary Examples

Pick et al. (2016) studied the role of egg size as a maternal effector (a mother's trait causing maternal effects). Despite the positive association between egg size and fitness, a significant amount of additive genetic variation is maintained in natural populations, which lead to an evolutionary stasis in egg size. The authors hypothesized that this variation is kept by genetic constraints due to shared genetics among maternal resource investment on egg components. They assessed this hypothesis by performing artificial selection in a captive population of Japanese quail (*Coturnix japonica*). They found effectively that the high maternal egg investment line produced eggs with more albumen, yolk and shell mass than low-line eggs.

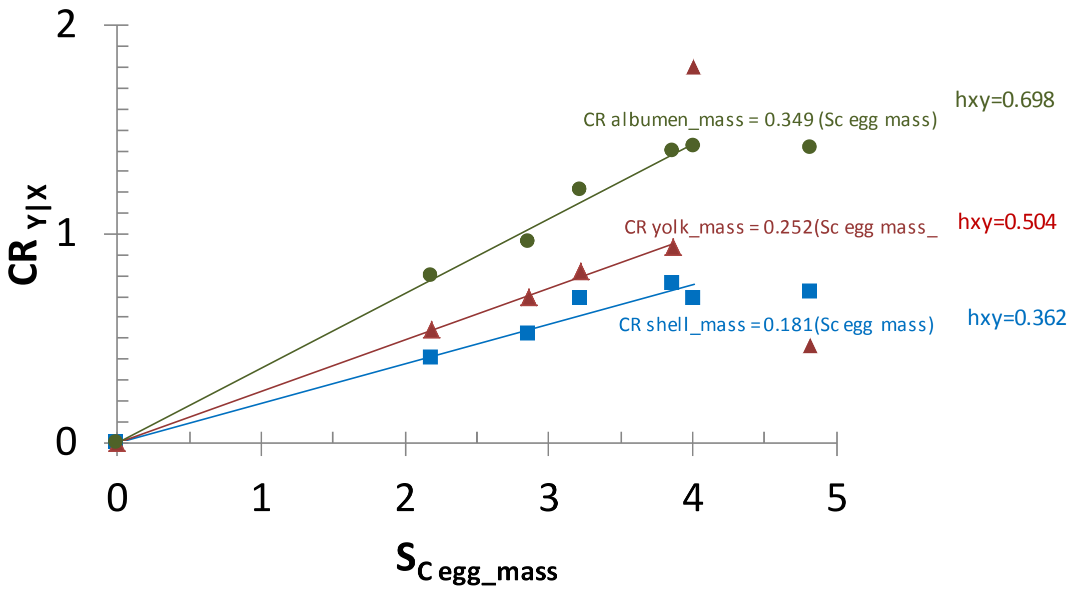

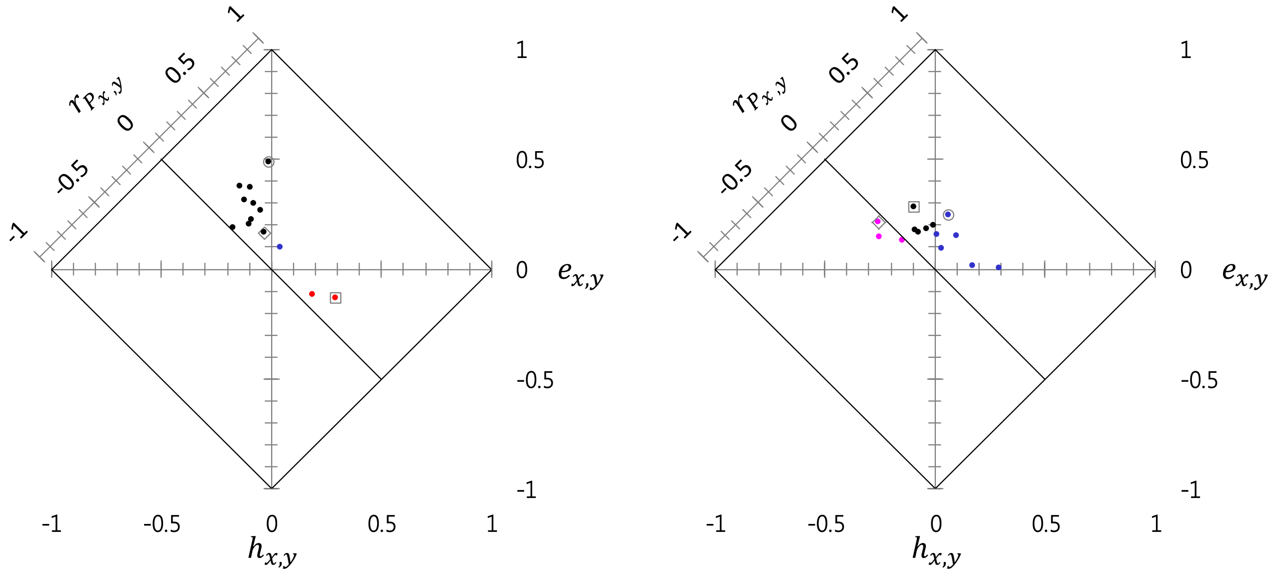

Figure S4-1. Coheritability, coenvironmentability and phenotypic correlation from pairwise analyses of morphological traits in house sparrow, *Passer domesticus*. (A) male, (B) female. The pair of traits highlighted are: tarsus length-bill length (square), tarsus length-body mass (circle), bill length-body mass (rhomboid). (Data source: Jensen et al. 2003). Notice the comparison of the phenotypic correlations of tarsus length and bill length in males ($r_m = 0.161, n = 407$) and females ($r_f = 0.185, n = 364$) were similar (Ho: $r_m = r_f$ was retained based on Zou's confidence interval of the difference -0.1609 and 0.1136 which included zero), the relative contribution of the coheritabilities and coenvironmentabilities were dramatically different in each sex. The coheritability (in males 0.295, in females -0.095) and and coenvironmentability (0.134 males, 0.280 females) were significantly different between each other. This tells of a differential genetic and environmental influence on these traits by sex.

#### Figure S5  Coheritability surface as a function of the heritabilities of the traits given a genetic correlation.

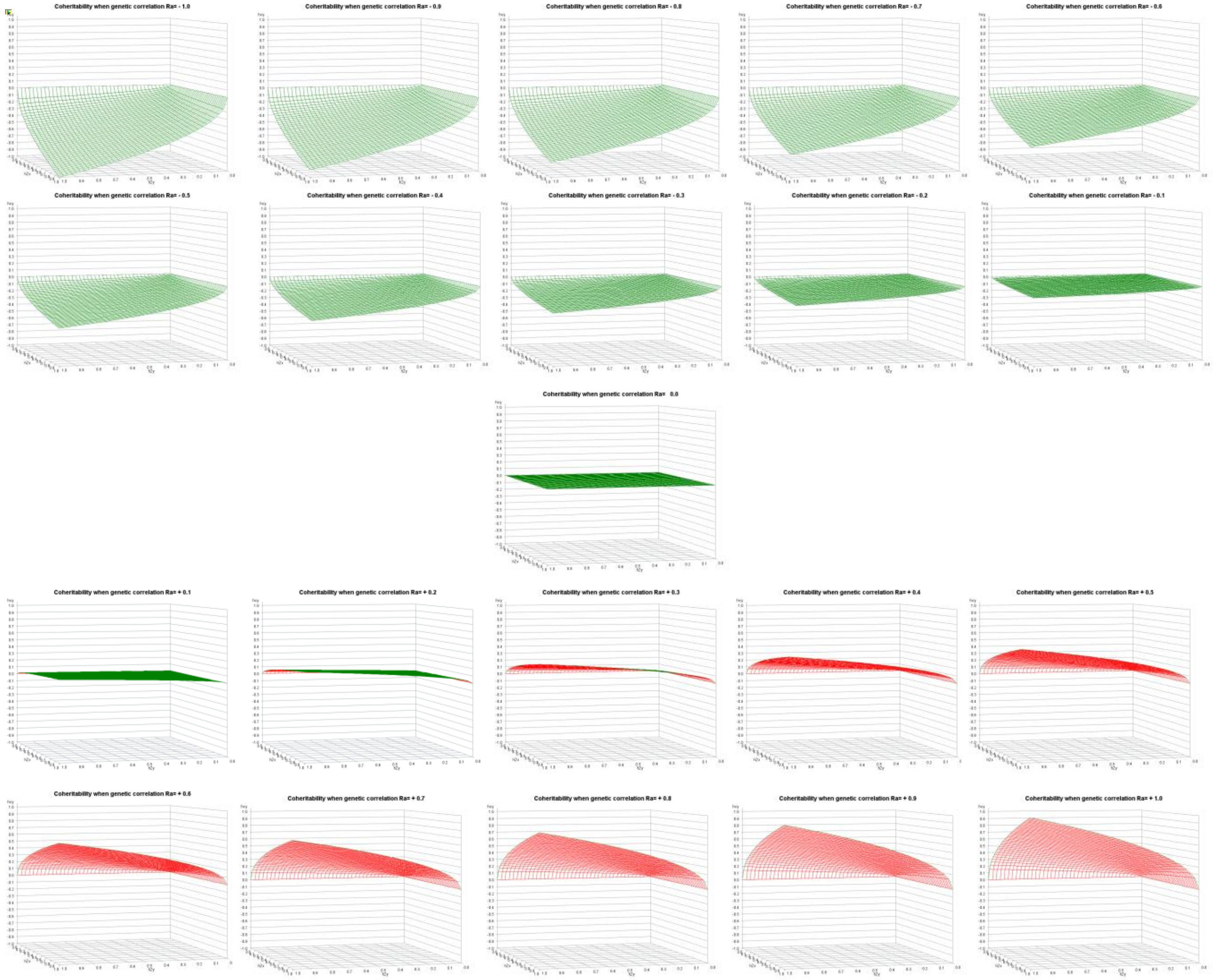

### Table T2. Definitions

| | heritability<br>$h^2$ | environmentability<br>$e^2$ | genetic correlation<br>$r_{A_{xy}}$ | environmental correlation<br>$r_{E_{xy}}$ | phenotypic correlation<br>$r_{P_{xy}}$ | coheritability<br>$h_{xy}$ | coenvironmentability<br>$e_{xy}$ |
|---|---|---|---|---|---|---|---|
| definition | • Proportion of the total phenotypic variance that is attributed to variability of breeding values. | • Residual proportion of the total phenotypic variance that is attributed to factors not accounted by breeding values. | Linear association between breeding values of two traits. Degree to which two characters are influenced by same loci.<br>• | | • Linear association between phenotypic values of two traits. | • Proportion of total phenotypic variability of two traits attributed to the additive covariance of the traits. | • Proportion of total phenotypic variability of two traits attributed to non genetic covariance of the traits. |
| | $h^2 = \frac{V_{A_x}}{V_{P_x}}$ | $e^2 = 1 - h^2$ | $r_{A_{x,y}} = \frac{C_{A_{x,y}}}{\sqrt{V_{A_x} V_{A_y}}}$ | $r_{E_{x,y}} = \frac{r_{P_{x,y}} - h_{xy}}{\sqrt{(1-h_x^2)(1-h_y^2)}}$ | $r_{P_{x,y}} = \frac{C_{P_{x,y}}}{\sqrt{V_{P_x} V_{P_y}}}$<br>$= \sqrt{h_x^2 h_y^2}\ r_{A_{x,y}} + \sqrt{(1-h_x^2)(1-h_y^2)}\ r_{E_{x,y}}$ | $h_{xy} = \frac{C_{A_{x,y}}}{\sqrt{V_{P_x} V_{P_y}}}$<br>$= \sqrt{h_x^2 h_y^2}\ r_{A_{x,y}}$ | $e_{xy} = r_{P_{x,y}} - h_{xy}$<br>$= \sqrt{(1-h_x^2)(1-h_y^2)}\ r_{E_{x,y}}$ |
| data | • breeding values of the trait<br>• population with known genetic structure | | • breeding values of each traits | • residual | • phenotypic values of each trait | • heritabilities of each trait, and genetic correlation, or<br>• phenotypic variances, and additive genetic covariance | • heritabilities of each trait, and environmental correlation, or<br>• phenotypic variances, and environmental covariance |
| range , domain | $0 < h^2 < 1$ | $0 < e^2 < 1$ | $-1 < r_{A_{xy}} < 1$ | $-1 < r_{E_{xy}} < 1$ | $-1 < r_{P_{xy}} < 1$ | $-1 < h_{xy} < 1$ | $-1 < e_{xy} < 1$ |
| cause | • degree of resemblace between relatives<br>• | •<br>•<br>• | • pleiotropic action of genes<br>• linkage disequilibrium<br>• | | •<br>•<br>• | •<br>•<br>• | •<br>•<br>• |
| graphical depiction | 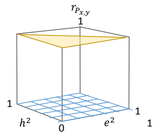  | | 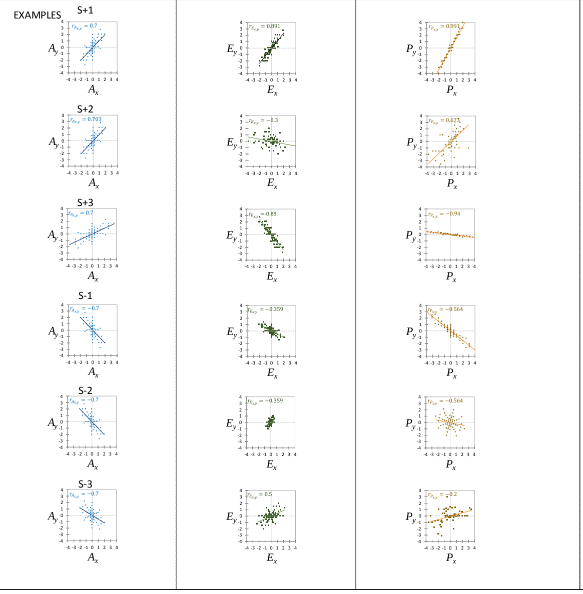  | | | 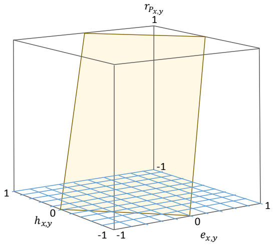  | |

### Appendix 5. Hypergeometric function

$_2F_1$ is the generalized hypergeometric function

$$_2F_1(a_1 , a_2 \,;\, b_1; x) \;=\; \sum_{m=0}^{\infty} \frac{(a_1)_m\,(a_2)_m}{(b_1)_k}\,\frac{x^m}{m\,!}$$

where $(q)_m$ denotes the rising Pochhammer symbol

$$(q)_m = \begin{cases} 1 & m = 0 \\ q(q-1)\,.\,.\,.\,(q+m-1) & m > 0 \end{cases}$$

| | **Hypergeometric function** $_2F_1(\,a_1 ,\, a_2 \,;\, b_1;\, x) = \sum_{m=0}^{\infty} \frac{(a_1)_m\,(a_2)_m}{(b_1)_k}\,\frac{x^m}{m\,!}$ | **Pochhammer function** $\frac{(a_1)_0 = (a_2)_0}{(b_1)_0}$ | **Hypergeometric function associated to the density distribution of the sample correlation coefficient** $_2F_1\left(\frac{1}{2}\,,\frac{1}{2};\, n-\frac{1}{2};\frac{1+\rho r}{2}\right)$ |
|---|---|---|---|
| *m = 0* | $\frac{(a_1)_0(a_2)_0}{(b_1)_0}\frac{x^0}{0\,!}$ | $(a_1)_0 = (a_2)_0 = \left(\frac{1}{2}\right)_0 = \mathbf{1}$<br>$(b_1)_0 = \left(n-\frac{1}{2}\right)_0 = \mathbf{1}$ | 1 |
| *m = 1* | $\frac{(a_1)_1(a_2)_1}{(b_1)_1}\frac{x^1}{1!} = \frac{a_1\,a_2}{b_1}\frac{x^1}{1\,!}$ | $(a_1)_1 = (a_2)_1 = \left(\frac{1}{2}\right)_1 = \mathbf{\frac{1}{2}}$<br>$(b_1)_1 = \left(n-\frac{1}{2}\right)_1 = \left[n-\frac{1}{2}\right] = \mathbf{\frac{1}{2}\,(2n-1)}$ | $\frac{\frac{1}{2}\cdot\frac{1}{2}}{\frac{(2n-1)}{2}}\left(\frac{1+\rho r}{2}\right)^1\frac{1}{1} = \mathbf{\frac{1\,(1+\rho r)}{4\,(2n-1)}}$ |
| *m = 2* | $\frac{(a_1)_2(a_2)_2}{(b_1)_2}\frac{x^2}{2!} = \frac{a_1(a_1+1)a_2(a_2+1)}{b_1(b_1+1)}\frac{x^2}{2\,!}$ | $(a_1)_2 = (a_2)_2 = \left(\frac{1}{2}\right)_2 = \frac{1}{2}\cdot\left[\frac{1}{2}+1\right] = \mathbf{\frac{3}{4}}$<br>$(b_1)_2 = \left(n-\frac{1}{2}\right)_2 = \left[n-\frac{1}{2}\right]\left[n+\frac{1}{2}\right] = \mathbf{\frac{1}{4}\,(2n-1)(2n+1)}$ | $\frac{\frac{3}{2}\cdot\frac{3}{2}}{\frac{(2n-1)(2n+1)}{4}}\left(\frac{1+\rho r}{2}\right)^2\frac{1}{2} = \mathbf{\frac{9}{32}\frac{(1+\rho r)^2}{(2n-1)(2n+1)}}$ |
| *m = 3* | $\frac{(a_1)_3(a_2)_3}{(b_1)_3}\frac{x^3}{3!} = \frac{a_1(a_1+1)(a_1+2)a_2(a_2+1)(a_2+2)}{b_1(b_1+1)(b_1+2)}\frac{x^3}{3\,!}$ | $(a_1)_3 = (a_2)_3 = \left(\frac{1}{2}\right)_3 = \frac{1}{2}\cdot\left[\frac{1}{2}+1\right]\left[\frac{1}{2}+2\right] = \mathbf{\frac{15}{8}}$<br>$(b_1)_3 = \left(n-\frac{1}{2}\right)_3 = \left[n-\frac{1}{2}\right]\left[n+\frac{1}{2}\right]\left[n+\frac{3}{2}\right]$<br>$= \mathbf{\frac{1}{8}\,(2n-1)(2n+1)(2n+3)}$ | $\frac{\frac{15}{8}\cdot\frac{15}{8}}{\frac{(2n-1)(2n+1)(2n+3)}{8}}\left(\frac{1+\rho r}{2}\right)^3\frac{1}{6} = \mathbf{\frac{225}{384}\frac{(1+\rho r)^3}{(2n-1)(2n+1)(2n+3)}}$ |
| *m = 4* | $\frac{(a_1)_4(a_2)_4}{(b_1)_4}\frac{x^4}{4!} = \frac{a_1(a_1+1)(a_1+2)(a_1+3)a_2(a_2+1)(a_2+2)(a_2+3)}{b_1(b_1+1)(b_1+2)(b_1+3)}\frac{x^4}{4\,!}$ | $(a_1)_4 = (a_2)_4 = \left(\frac{1}{2}\right)_4 = \frac{1}{2}\cdot\left[\frac{1}{2}+1\right]\left[\frac{1}{2}+2\right]\left[\frac{1}{2}+3\right] = \mathbf{\frac{105}{16}}$<br>$(b_1)_4 = \left(n-\frac{1}{2}\right)_4 = \left[n-\frac{1}{2}\right]\left[n+\frac{1}{2}\right]\left[n+\frac{3}{2}\right]\left[n+\frac{5}{2}\right]$<br>$= \mathbf{\frac{1}{16}\,(2n-1)(2n+1)(2n+3)(2n+5)}$ | $\frac{\frac{105}{16}\cdot\frac{105}{16}}{\frac{(2n-1)(2n+1)(2n+3)(2n+5)}{16}}\left(\frac{1+\rho r}{2}\right)^4\frac{1}{24}$<br>$= \mathbf{\frac{11025}{6144}\frac{(1+\rho r)^4}{(2n-1)(2n+1)(2n+3)(2n+5)}}$ |
| *m = 5* | $\frac{(a_1)_5(a_2)_5}{(b_1)_5}\frac{x^5}{5!}$<br>$= \frac{a_1(a_1+1)(a_1+2)(a_1+3)(a_1+4)a_2(a_2+1)(a_2+2)(a_2+3)(a_2+4)}{b_1(b_1+1)(b_1+2)(b_1+3)(b_1+4)}\frac{x^5}{5\,!}$ | $(a_1)_5 = (a_2)_5 = \left(\frac{1}{2}\right)_5 = \frac{1}{2}\cdot\left[\frac{1}{2}+1\right]\left[\frac{1}{2}+2\right]\left[\frac{1}{2}+3\right]\left[\frac{1}{2}+4\right] = \mathbf{\frac{945}{32}}$<br>$(b_1)_5 = \left(n-\frac{1}{2}\right)_5 = \left[n-\frac{1}{2}\right]\left[n+\frac{1}{2}\right]\left[n+\frac{3}{2}\right]\left[n+\frac{5}{2}\right]\left[n+\frac{7}{2}\right]$<br>$= \mathbf{\frac{1}{32}\,(2n-1)(2n+1)(2n+3)(2n+5)(2n+7)}$ | $\frac{\frac{945}{32}\cdot\frac{945}{32}}{\frac{(2n-1)(2n+1)(2n+3)(2n+5)(2n+7)}{32}}\left(\frac{1+\rho r}{2}\right)^5\frac{1}{120}$<br>$= \mathbf{\frac{893025}{122880}\frac{(1+\rho r)^5}{(2n-1)(2n+1)(2n+3)(2n+5)(2n+7)}}$ |
| *m = 6* | $\frac{(a_1)_6(a_2)_6}{(b_1)_6}\frac{x^6}{6!}$<br>$= \frac{a_1(a_1+1)(a_1+2)(a_1+3)(a_1+4)a_2(a_2+1)(a_2+2)(a_2+3)(a_2+4)(a_2+5)}{b_1(b_1+1)(b_1+2)(b_1+3)(b_1+4)(b_1+5)}\frac{x^6}{6\,!}$ | $(a_1)_6 = (a_2)_6 = \left(\frac{1}{2}\right)_6 = \frac{1}{2}\cdot\left[\frac{1}{2}+1\right]\left[\frac{1}{2}+2\right]\left[\frac{1}{2}+3\right]\left[\frac{1}{2}+4\right]\left[\frac{1}{2}+5\right] = \mathbf{\frac{10395}{64}}$<br>$(b_1)_6 = \left(n-\frac{1}{2}\right)_6 = \left[n-\frac{1}{2}\right]\left[n+\frac{1}{2}\right]\left[n+\frac{3}{2}\right]\left[n+\frac{5}{2}\right]\left[n+\frac{7}{2}\right]\left[n+\frac{9}{2}\right]$<br>$= \mathbf{\frac{1}{64}\,(2n-1)(2n+1)(2n+3)(2n+5)(2n+7)(2n+9)}$ | $\frac{\frac{10395}{64}\cdot\frac{10396}{64}}{\frac{(2n-1)(2n+1)(2n+3)(2n+5)(2n+7)(2n+9)}{64}}\left(\frac{1+\rho r}{2}\right)^6\frac{1}{720}$<br>$= \mathbf{\frac{108056025}{2949120}\frac{(1+\rho r)^6}{(2n-1)(2n+1)(2n+3)(2n+5)(2n+7)(2n+9)}}$ |
| *m > 7* | $\sum_{m=7}^{\infty}\frac{(a_1)_m\,(a_2)_m}{(b_1)_k}\,\frac{x^m}{m\,!}$ | | $O\,(n^{-8})$ |

### Appendix 6. Delta Method

Taylor Expansion of a trivariate function around its means

$$f(x,y,z) = f(x_o,y_o,z_o) + f_x^1(x_o,y_o,z_o)(x-x_o) + f_y^1(x_o,y_o,z_o)(y-y_o) + f_z^1(x_o,y_o,z_o)(z-z_o)$$

where $x_o, y_o, z_o$ are constants (considered here as means). The first term is the actual estimate of the function. Terms of the form $f_{\blacksquare}^1(x_o, y_o, z_o)$ are the first-order derivatives evaluated around $x_o, y_o, z_o$.

Expected value

$$E[f(x,y,z)] = E[f(x_o,y_o,z_o) + f_x^1(x_o,y_o,z_o)(x-x_o) + f_y^1(x_o,y_o,z_o)(y-y_o) + f_z^1(x_o,y_o,z_o)(z-z_o)]$$

$$= E[f(x_o,y_o,z_o)] + f_x^1(x_o,y_o,z_o)E(x-x_o) + f_y^1(x_o,y_o,z_o)E(y-y_o) + f_z^1(x_o,y_o,z_o)E(z-z_o)$$

Since the expected value of a variable around its mean is zero,

$$E[f(x,y,z)] = f(x_o,y_o,z_o)$$

Variance

Here we define the variance of a variable as the squared deviations around its mean

$$Var[f(x,y,z)] = E(f(x,y,z) - E[f(x,y,z)])^2$$

$$= [f(x_o,y_o,z_o) + f_x^1(x_o,y_o,z_o)(x-x_o) + f_y^1(x_o,y_o,z_o)(y-y_o) + f_z^1(x_o,y_o,z_o)(z-z_o) - f(x_o,y_o,z_o)]^2$$

$$= [f_x^1(x_o,y_o,z_o)(x-x_o) + f_y^1(x_o,y_o,z_o)(y-y_o) + f_z^1(x_o,y_o,z_o)(z-z_o)]^2$$

$$= E[f_x^1(x_o,y_o,z_o)(x-x_o) + f_y^1(x_o,y_o,z_o)(y-y_o) + f_z^1(x_o,y_o,z_o)(z-z_o)]^2$$

Squaring the terms within the brackets, we obtain

$$= E\Big[\big(f_x^1(x_o,y_o,z_o)\big)^2(x-x_o)^2 + \big(f_y^1(x_o,y_o,z_o)\big)^2(y-y_o)^2 + \big(f_z^1(x_o,y_o,z_o)\big)^2(z-z_o)^2$$

$$+ 2 f_x^1(x_o,y_o,z_o) f_y^1(x_o,y_o,z_o)(x-x_o)(y-y_o)$$

$$+ 2 f_x^1(x_o,y_o,z_o) f_z^1(x_o,y_o,z_o)(x-x_o)(z-z_o)$$

$$+ 2 f_y^1(x_o,y_o,z_o) f_z^1(x_o,y_o,z_o)(y-y_o)(z-z_o)\Big]$$

Taking the expectation of the terms involving variables

$$= \big(f_x^1(x_o,y_o,z_o)\big)^2 E(x-x_o)^2 + \big(f_y^1(x_o,y_o,z_o)\big)^2 E(y-y_o)^2 + \big(f_z^1(x_o,y_o,z_o)\big)^2 E(z-z_o)^2$$

$$+ 2 f_x^1(x_o,y_o,z_o) f_y^1(x_o,y_o,z_o)E[(x-x_o)(y-y_o)]$$

$$+ 2 f_x^1(x_o,y_o,z_o) f_z^1(x_o,y_o,z_o)E[(x-x_o)(z-z_o)]$$

$$+ 2 f_y^1(x_o,y_o,z_o) f_z^1(x_o,y_o,z_o)E[(y-y_o)(z-z_o)]$$

Finally, the variance of the function can be approximated

$$Var[\,f(\,x\,,y\,,z\,)\,] = \left(f_x^1(x_o\,,y_o\,,z_o\,)\right)^2 Var(x) + \left(f_y^1(x_o\,,y_o\,,z_o\,)\right)^2 Var(y) + \left(f_z^1(x_o\,,y_o\,,z_o\,)\right)^2 Var(z)$$

$$+\,2\,f_x^1(x_o\,,y_o\,,z_o\,)\,f_y^1(x_o\,,y_o\,,z_o\,)Cov(x,y)$$

$$+\,2\,f_x^1(x_o\,,y_o\,,z_o\,)\,f_z^1(x_o\,,y_o\,,z_o\,)Cov(x,z)$$

$$+\,2\,f_y^1(x_o\,,y_o\,,z_o\,)\,f_z^1(x_o\,,y_o\,,z_o\,)Cov(y,z)$$

| | | Estimator | | | |
|---|---|---|---|---|---|
| | | phenotypic correlation | coheritability | coenvironmentability | heritability |
| function | f = $f(\,x_0\,,y_0\,,z_0\,)$ | $r_{P_{x,y}} = \frac{s_{P_{x,y}}}{\sqrt{s_{P_x}^2\,s_{P_y}^2}}$ | $h_{x,y} = \sqrt{h_x^2\,h_y^2\,r_{A_{x,y}}}$ | $e_{x,y} = \sqrt{(1-h_x^2)(1-\,h_y^2)}\,r_{E_{x,y}}$ | $h^2 = \frac{V_A}{V_P}$ |
| elements | $x_0$ | $s_{P_{x,y}}$ | $h_x^2$ | $h_x^2$ | $V_A$ |
| | $y_0$ | $s_{P_x}^2$ | $h_x^2$ | $h_x^2$ | $V_P$ |
| | $z_0$ | $s_{P_y}^2$ | $r_{A_{x,y}}$ | $r_{E_{x,y}}$ | |
| first-order derivatives | $f_x^1 = \left.\frac{\delta\,\mathrm{f}}{\delta\,x}\right]_{x_0,y_0,z_0}$ | $\frac{\delta\,r_{P_{x,y}}}{\delta\,\,s_{P_{x,y}}} = (\,s_{P_x}^2\,s_{P_y}^2\,)^{-\frac{1}{2}}$ | $\frac{\delta\,h_{x,y}}{\delta\,h_x^2} = \frac{1}{2}\,h_y^2\,r_{A_{x,y}}\,(h_x^2 h_y^2)^{-\frac{1}{2}}$ | $\frac{\delta\,e_{x,y}}{\delta\,h_x^2} = -\frac{(1-h_y^2)}{2\sqrt{(1-h_x^2)(1-h_y^2)}}\,r_{E_{x,y}}$ | $\frac{\delta\,h^2}{\delta\,V_A} = \frac{1}{V_P}$ |
| | $f_x^1 = \left.\frac{\delta\,\mathrm{f}}{\delta\,y}\right]_{x_0,y_0,z_0}$ | $\frac{\delta\,r_{P_{x,y}}}{\delta\,s_{P_x}^2} = -\,\frac{1}{2}(\,s_{P_{x,y}}\,s_{P_y}^2)(\,s_{P_x}^2\,s_{P_y}^2)^{-\frac{3}{2}}$ | $\frac{\delta\,h_{x,y}}{\delta\,h_y^2} = \frac{1}{2}\,h_x^2\,r_{A_{x,y}}\,(\,h_x^2\,h_y^2)^{-\frac{1}{2}}$ | $\frac{\delta\,e_{x,y}}{\delta\,h_y^2} = -\frac{(1-h_x^2)}{2\sqrt{(1-h_x^2)(1-h_y^2)}}\,r_{E_{x,y}}$ | $\frac{\delta\,h^2}{\delta\,V_P} = -\,\frac{V_A}{V_P}$ |
| | $f_x^1 = \left.\frac{\delta\,\mathrm{f}}{\delta\,z}\right]_{x_0,y_0,z_0}$ | $\frac{\delta\,r_{P_{x,y}}}{\delta\,s_{P_y}^2} = -\,\frac{1}{2}(\,s_{P_{x,y}}\,s_{P_x}^2)(\,s_{P_x}^2\,s_{P_y}^2\,)^{-\frac{3}{2}}$ | $\frac{\delta\,h_{x,y}}{\delta\,r_{A_{x,y}}} = (\,h_x^2\,h_y^2)^{\frac{1}{2}}$ | $\frac{\delta\,e_{x,y}}{\delta\,r_{E_{x,y}}} = \sqrt{(1-h_x^2)(1-h_y^2)}$ | |

Note on the Taylor expansion of a function involving three variables

Taylor series are polynomials that approximate functions. For functions of three variables, the Taylor series depends upon first, second, third, etc. partial derivatives at some point $(x_o , y_o , z_o)$, which involve taking sequential derivatives with respect to the same or different variables. The following is the expansion of the function up to the fourth-order.

$$f(x , y , z) = f(x_o , y_o , z_o )$$

$$+ \frac{1}{1!}\left[ f_x^1(x_o , y_o , z_o )(x - x_o) + f_y^1(x_o , y_o , z_o )(y - y_o) + f_z^1(x_o , y_o , z_o )(z - z_o)\right]$$

$$+\frac{1}{2!}\left[ f_{xx}^2(x_o , y_o , z_o )(x - x_o)^2 + 2f_{xy}^2(x_o , y_o , z_o )(x - x_o)(y - y_o) + 2f_{xz}^2(x_o , y_o , z_o )(x - x_o)(z - z_o) + 2f_{yy}^2(x_o , y_o , z_o )(y - y_o)^2 + 2f_{yz}^2(x_o , y_o , z_o )(y - y_o)(z - z_o) + 2f_{zz}^2(x_o , y_o , z_o )(z - z_o)^2 \right]$$

$$+ \frac{1}{3!} \left[ f_{xxx}^3(x_o , y_o , z_o )(x - x_o)^3 + 3f_{xxy}^3(x_o , y_o , z_o )(x - x_o)^2(y - y_o) + 3f_{xxz}^3(x_o , y_o , z_o )(x - x_o)^2(z - z_o) + 3f_{yyx}^3(x_o , y_o , z_o )(y - y_o)^2(x - x_o) + f_{yyy}^3(x_o , y_o , z_o )(y - y_o)^3 + 3f_{yyz}^3(x_o , y_o , z_o )(y - y_o)^2(z - z_o) + 3f_{zzx}^3(x_o , y_o , z_o )(z - z_o)^2(x - x_o) + 3f_{zzy}^3(x_o , y_o , z_o )(z - z_o)^2(y - y_o) + f_{zzz}^3(x_o , y_o , z_o )(z - z_o)^3 + 6f_{xyz}^3(x_o , y_o , z_o )(x - x_o)(y - y_o)(z - z_o) \right]$$

$$+ \frac{1}{4!} \left[f_{xxxx}^4(x_o , y_o , z_o )(x - x_o)^4 + 4f_{xxxy}^4(x_o , y_o , z_o )(x - x_o)^3(y - y_o) + 6f_{xxyy}^4(x_o , y_o , z_o )(x - x_o)^2(y - y_o)^2 + 4f_{xxxz}^4(x_o , y_o , z_o )(x - x_o)^3(z - z_o) + 4f_{xyyy}^4(x_o , y_o , z_o )(x - x_o)(y - y_o)^3 + 12f_{xxyz}^4(x_o , y_o , z_o )(x - x_o)^2(y - y_o)(z - z_o) + 6f_{xxzz}^4(x_o , y_o , z_o )(x - x_o)^2(z - z_o)^2 + 12f_{xyyz}^4(x_o , y_o , z_o )(x - x_o)(y - y_o)^2(z - z_o) + 12f_{xyzz}^4(x_o , y_o , z_o )(x - x_o)(z - z_o)(y - y_o)^2 + 4f_{xzzz}^4(x_o , y_o , z_o )(x - x_o)(z - z_o)^3 + f_{yyyy}^4(x_o , y_o , z_o )(y - y_o)^4 + 4f_{yyyz}^4(x_o , y_o , z_o )(y - y_o)^3(z - z_o) + 6f_{yyzz}^4(x_o , y_o , z_o )(y - y_o)^2(z - z_o)^2 + 4f_{yzzz}^4(x_o , y_o , z_o )(y - y_o)(z - z_o)^3 + f_{zzzz}^4(x_o , y_o , z_o )(z - z_o)^4 \right]$$

$$+ O(n^{-5})$$

### Appendix 7. Web resources

The URLs of online resources relevant to this work are presented below:

---

#### Testing the significance of correlations

*https://www.psychometrica.de/correlation.html*

1. Comparison of correlations from independent samples. 2. Comparison of correlations from dependent samples. 3. Testing linear independence (Testing against 0). 4. Testing correlations against a fixed value. 5. Calculation of confidence intervals of correlations. 6. Fisher-Z-Transformation. 7. Calculation of the weighted mean of a list of correlations. 8. Transformation of the effect sizes r, d, f, Odds Ratio and eta square. 9. Calculation of Linear Correlations.

Citation

Lenhard, W. & Lenhard, A. 2014 . *Hypothesis Tests for Comparing Correlations*. Available: Bibergau (Germany): Psychometrica. DOI: 10.13140/RG.2.1.2954.1367
https://www.psychometrica.de/correlation.html.

---

#### The confidence interval of rho (the correlation coefficient)

*http://vassarstats.net/rho.html*

1. Calculation of the 0.95 and 0.99 confidence intervals for rho, based on the Fisher r-to-z transformation. The values of *r* and *n* are required. Note that the confidence interval of rho is symmetrical around the observed r only with large values of n.
2. *- Free, full-length, and interactive statistics textbook. It is a companion site of "VassarStats: Web Site for Statistical Computation"*

Citation
Lowry, R 2015. Concepts and Applications of Inferential Statistics.
*http://vassarstats.net/textbook/*

---

#### Confidence Interval

http://www.psyctc.org/cgi-bin/R.cgi/CI_correln.R

---

#### Sample size calculators for designing clinical research

*http://www.sample-size.net/correlation-sample-size*

1. Total sample size required to determine whether a correlation coefficient differs from zero.

Citation
Hulley SB, Cummings SR, Browner WS, Grady D, Newman TB 2013 . Designing clinical research : an epidemiologic approach. 4th ed. Philadelphia, PA: Lippincott Williams & Wilkins. Appendix 6C, page 79.

#### Fisher r-to-Z transformation

http://vassarstats.net/tabs_rz.html

For any particular value of r, the Pearson product-moment correlation coefficient, this section will perform the Fisher r-to-z transformation according to the formula

$$Z_r = \frac{1}{2}\, ln\left(\frac{1+r}{1-r}\right)$$

If a value of N is entered (optional), it will also calculate the standard deviation of $z_r$ as

$$\sigma_{Z_r} = \frac{1}{n-3}$$

#### Cocor Comparing correlations

*http://comparingcorrelations.org*

1. Tests for comparing two correlations based on independent groups: Fisher's z; Zou's confidence interval.

2. Tests for comparing two correlations based on dependent groups with overlapping variables: Pearson and Filon's; Hotelling's; Williams; Olkin's; Dunn and Clark's; Hendrickson's modification of Williams'; Steiger's modification of Dunn and Clark's;Hittner's modification of Dunn and Clark's;Zou's.

3. Tests for comparing two correlations based on dependent groups with nonoverlapping variables: Pearson and Filon's; Dunn and Clark's; Steiger's modification of Dunn and Clark's; Raghunathan, Rosenthal, and Rubin's modification of Pearson and Filon's; Silver, Hittner, and May's modification of Dunn and Clark's; Zou's confidence interval.

Citation:
Diedenhofen B, Musch J 2015 . cocor: A Comprehensive Solution for the Statistical Comparison of Correlations. PLoS ONE 10(4): e0121945. doi:10.1371/journal.pone.0121945. http://journals.plos.org/plosone/article/file?id=10.1371/journal.pone.0121945&type=printable

Package documentation
http://comparingcorrelations.org/repo/pckg/cocor/cocor.pdf

Useful online blogs and information

#### Fisher's transformation of the correlation coefficient

*https://blogs.sas.com/content/iml/2017/09/20/fishers-transformation-correlation.html?utm_source=feedburner&utm_medium=feed&utm_campaign=Feed%3A+TheDoLoop+%28The+DO+Loop%29*

By Rick Wicklin on The DO Loop |September 20, 2017

This article shows that Fisher's "z transformation," which is z = arctanh(r), is a normalizing transformation for the Pearson correlation of bivariate normal samples of size N. The transformation converts the skewed and bounded sampling distribution of r into a normal distribution for z. The standard error of the transformed distribution is 1/sqrt(N-3), which does not depend on the correlation.

One can download the SAS program that creates all the graphs in this article.

---

#### Simulate correlations by using the Wishart distribution

*https://blogs.sas.com/content/iml/2017/10/11/simulate-correlations-wishart-distribution.html*

By Rick Wicklin on The DO Loop October 11, 2017

There is a simpler ways to simulate the correlation estimates: One can directly simulate from the Wishart distribution. Each draw from the Wishart distribution is a sample covariance matrix for a multivariate normal sample of size N. If you convert that covariance matrix to a correlation matrix, you can immediately extract the off-diagonal elements.

---

#### Coverage probability of confidence intervals: A simulation approach

*https://blogs.sas.com/content/iml/2016/09/08/coverage-probability-confidence-intervals.html*

By Rick Wicklin on The DO Loop

The article uses the SAS DATA step and Base SAS procedures to estimate the coverage probability of the confidence interval for the mean of normally distributed data. This discussion is based on Section 5.2 (p. 74–77) of Simulating Data with SAS.

---

http://keisan.casio.com/exec/system/1359533867

http://www.emathhelp.net/calculators/probability-statistics/
